# Weak magnetic field effects in biology are measurable—accelerated *Xenopus* embryogenesis in the absence of the geomagnetic field

**DOI:** 10.1101/2024.10.10.617626

**Authors:** Alessandro Lodesani, Geoff Anders, Lykourgos Bougas, Tobias Lins, Dmitry Budker, Peter Fierlinger, Clarice D. Aiello

**Affiliations:** Quantum Biology Institute, Los Angeles, USA; Leverage, USA; Johannes Gutenberg-Universität Mainz, Mainz, Germany; Technische Universität München, Munich, Germany; Helmholtz Institute Mainz, Mainz, Germany; GSI Helmholtzzentrum für Schwerionenforschung GmbH, Darmstadt, Germany; Department of Physics, University of California, Berkeley, USA

## Abstract

Despite decades of reports of weak magnetic field effects in biology across the tree of life and on a broad range of cell types, the evidence to date remains met with skepticism. To remedy this, we present open-data, large-scale, and varied morphological evidence that *Xenopus laevis* embryo development is accelerated in a well-engineered, environmentally-calibrated hypomagnetic field of less than 1 nT. These data imply that basal tadpole physiology can sense and react to the absence of Earth’s minute magnetic field of approximately 50 *µ*T. The effect is significant, as demonstrated by a variety of statistical measures. As no definitive biophysical mechanism has been identified to account for its occurrence, this study raises the question of which mechanism provides the most plausible explanation. How that question is answered may have implications in a variety of fields, including human health, behavioral ecology, and space exploration.

## 1. Summary

The biological effects of weak magnetic fields, such as Earth’s geomagnetic field (*∼* 50 *µ*T), have long been reported, but remain poorly understood and met with skepticism [1, 2]. This scientific stalemate is due both to the absence of a definitive biophysical mechanism to explain such effects, and to experimental shortcomings. First, the rationale has been that the biological effects of weak magnetic fields are negligible because the field strengths are thought to be too small to trigger a biophysical sensing mechanism. Second, large-scale, thoroughly environmentally controlled data are lacking, and existing results are often ambiguous regarding whether the effects are truly attributable to magnetic fields or other environmental factors. However, the reality of a given effect can be established absent a known mechanism with a sufficiently well-designed experiment. Here, we demonstrate that the development of *Xenopus laevis* embryos is significantly accelerated when they are raised in a hypomagnetic environment (*<* 1 nT) shielded from Earth’s geomagnetic field. Our large-scale dataset and analysis code are publicly available. After extensive calibration of environmental conditions, our assessment is that the origin of the effects is the absence of Earth’s magnetic field. Our main finding is thus that the basal physiology of a non-migratory species like *Xenopus laevis* significantly reacts to a magnetic shift on the order of 50 *µ*T, with effects observed as early as one day post-fertilization. These results are sufficiently robust to advance the scientific consensus that weak magnetic fields indeed exert measurable biological effects. We also raise the question of which underlying mechanism could best explain our observations. Answering this could have far-reaching implications across fields such as human health, behavioral ecology, and space exploration.

## 2. Introduction

Weak magnetic field effects in biology have been reported for many years [1]. Reported effects are wide-ranging, from the up- and down-regulation of cell proliferation [2], to DNA repair [3], to cellular respiration and metabolism [4, 5], to cytoskeleton configuration [6, 7], and ion channel functioning [8]. There are also established migratory phenomena, such as the ability of birds to navigate at dusk, which are presumed to depend on Earth’s magnetic field [9], which is itself very weak.

Nevertheless, these magnetic effects have remained controversial. First, weak magnetic fields (we consider shifts of *∼* 1 mT, on the order of a cell phone’s field [10], or smaller to be ‘weak’ for the purposes of this paper) are broadly not expected to be strong enough to have any noticeable effect on biological organisms. Second, the magnitude of reported effects does not vary directly with field intensity, yielding uncertainty about the underlying mechanism. The reality of a given effect can be established absent a known mechanism with a sufficiently well-designed experiment. However, until now the studies of weak magnetic field effects in biology have suffered from shortcomings, ranging from small sample sizes to inadequately controlled conditions.

In our experiment, we raise *Xenopus laevis* frog embryos in a carefully constructed hypomagnetic chamber. The chamber, in combination with compensation coils, subtracts the Earth’s magnetic field (of approximately 50 *µ*T), yielding a hypomagnetic condition of less than 1 nT, close to zero. We examine over 2,750 tadpoles over 10 batches. The first seven batches have *Xenopus* embryos, raised in individual wells, divided randomly between the hypomagnetic condition and a control condition, which is a deactivated cell incubator with comparable light, humidity, and temperature conditions. Of these, the first three batches were used to identify a plausible hypothesis, namely, that *Xenopus* embryos raised in a hypomagnetic condition experience accelerated development; the remaining four batches test this hypothesis. We then raise three further batches of embryos in a positive control, where the Earth’s magnetic field is recreated inside the hypomagnetic chamber, and embryos are again divided between the hypomagnetic chamber and the deactivated incubator.

Using a variety of objective morphological measures, we find significant differences between the tadpoles raised in the hypomagnetic condition and the regular and positive controls. Differences, which pertain to both the shape and color of the embryos, are visible upon inspection and are confirmed by a variety of statistical tests, where the role of human judgment is reduced as much as possible. The differences appear on day 1 post-fertilization and are present on days 2 and 3 post-fertilization; on day 3, the embryos are preserved in formalin. The differences we observe are consistent with those observed by other labs, both those that studied the effects of minutely increasing and decreasing the intensity of magnetic fields experienced by *Xenopus* embryos. All of our data has been made publicly available.

This experiment solidifies direct observational evidence for weak magnetic field effects in a particular biological system. Since we remove a pre-existing magnetic field, rather than applying a new one, it follows that *Xenopus* development is impacted by Earth’s magnetic field. *Xenopus* is not migratory, so there is no reason to expect that it would evolve mechanisms to navigate using Earth’s magnetic field, especially as early as day 1 post-fertilization. There are also no indications of magnetite, as in magnetotactic bacteria, or conduction channels, like those in sharks and rays. This raises the question of the mechanism and biological significance of the effect, which we speculate on only briefly.

Historically, our experiment bears some resemblance to that of Francis Hauksbee [11], who in the early eighteenth century used a sophisticated air pump to create a superior vacuum condition for studying electricity. In this case, rather than an air pump, we use a mu-metal and copper-shielded box; rather than bringing air pressure close to zero, we bring the magnetic field close to zero; and rather than confirming the electrical nature of light emitted by frictional processes, we confirm that weak magnetic field effects in biology are measurable.

## 3. The experiment is well-controlled, with magnetic field environment as the only significant variable

We chose *Xenopus laevis* for our experiments because it is a widely used model organism in embryology, with straightforward husbandry and embryos that can be raised at room temperature (Supplementary Section SI1). Frogs are mostly non-migratory, with known exceptions of magnetic geofield sensing during pond changes [12]; we see no clear evolutionary advantage for *Xenopus laevis* in being able to sense Earth’s weak magnetic field early in embryogenesis. Our selection of *Xenopus* is thus based on how common it is, rather than on any desired organismal trait.

For three days, we raise each tadpole individually in the wells of 24- or 48-well plates, with a total of over 2,750 tadpoles. This setup disrupts chemical communication between tadpoles, completely prevents cross-contamination, and, most importantly, allows us to monitor the morphological development of each individual. All plates are prepared at the same time and filled with 0.1x Marc’s Modified Ringer’s (MMR) solution (Supplementary Section SI2) from the same bottle. After *in vitro* fertilization (Supplementary Section SI2) and egg randomization (Supplementary Section SI3), embryos are placed in either a hypomagnetic or control environment (*i.e.*, exposed to Earth’s ambient magnetic field) and left undisturbed for 24 hours, with the plates’ plastic lids on. Imaging is performed approximately every 24 hours post-fertilization (p.f.) for three days (Supplementary Section SI4). Day 1 and 2 imaging sessions are usually completed in well under an hour, since the embryos are not motile, while day 3 tadpoles are fixed in formalin prior to imaging. Typical images for each day are displayed in panel c) of Fig. 1; the entirety of our image database, along with all developed code, is organized and accessible in the project’s GitHub depository (a guide is found in Supplementary Section SI69). We perform one *in vitro* fertilization per week, with the resulting tadpole populations referred to as ‘batches’ (*e.g.*, batch B2 is the second batch; for a batch overview, see Supplementary Section SI5).

**Fig. 1:**
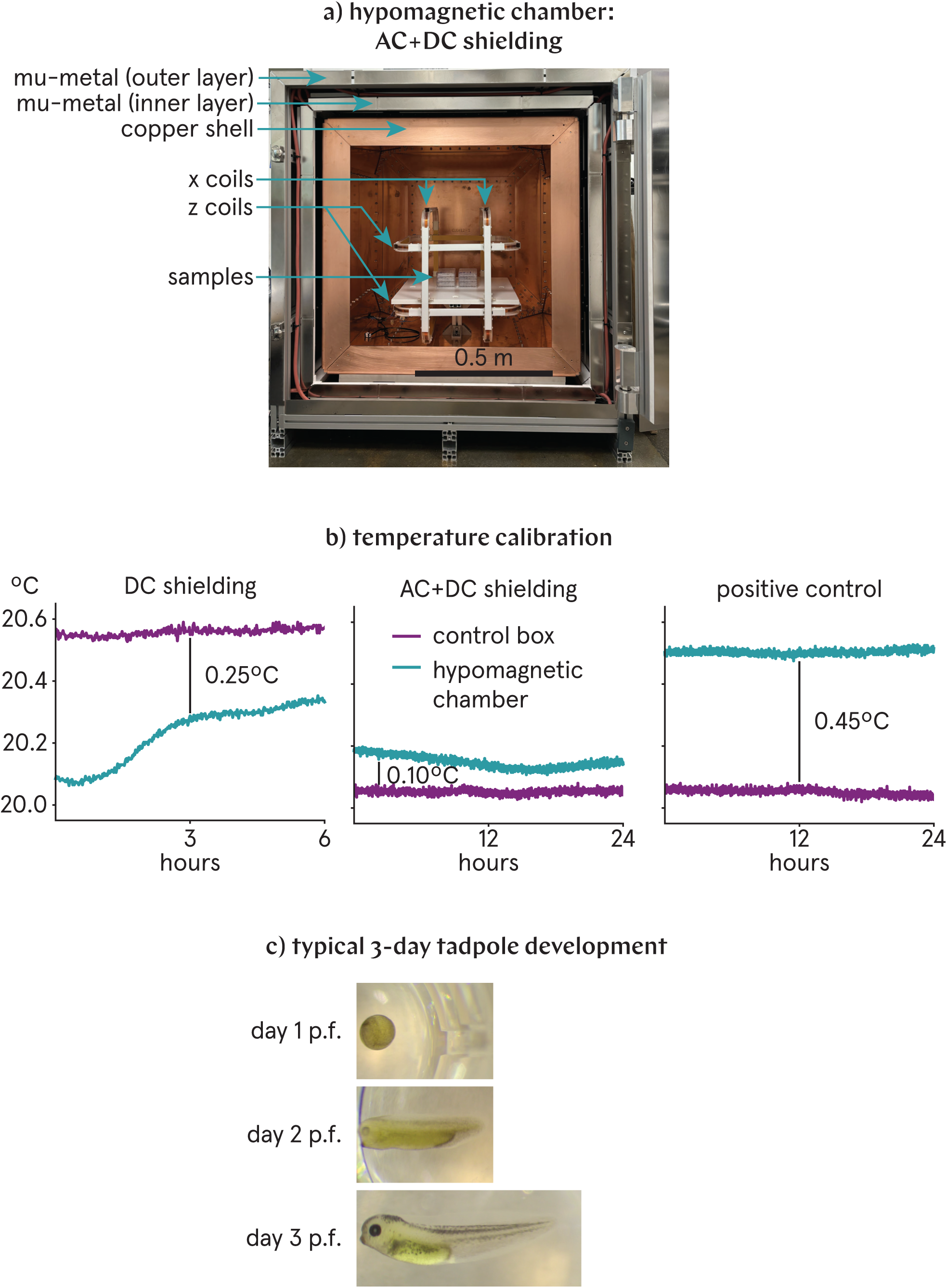
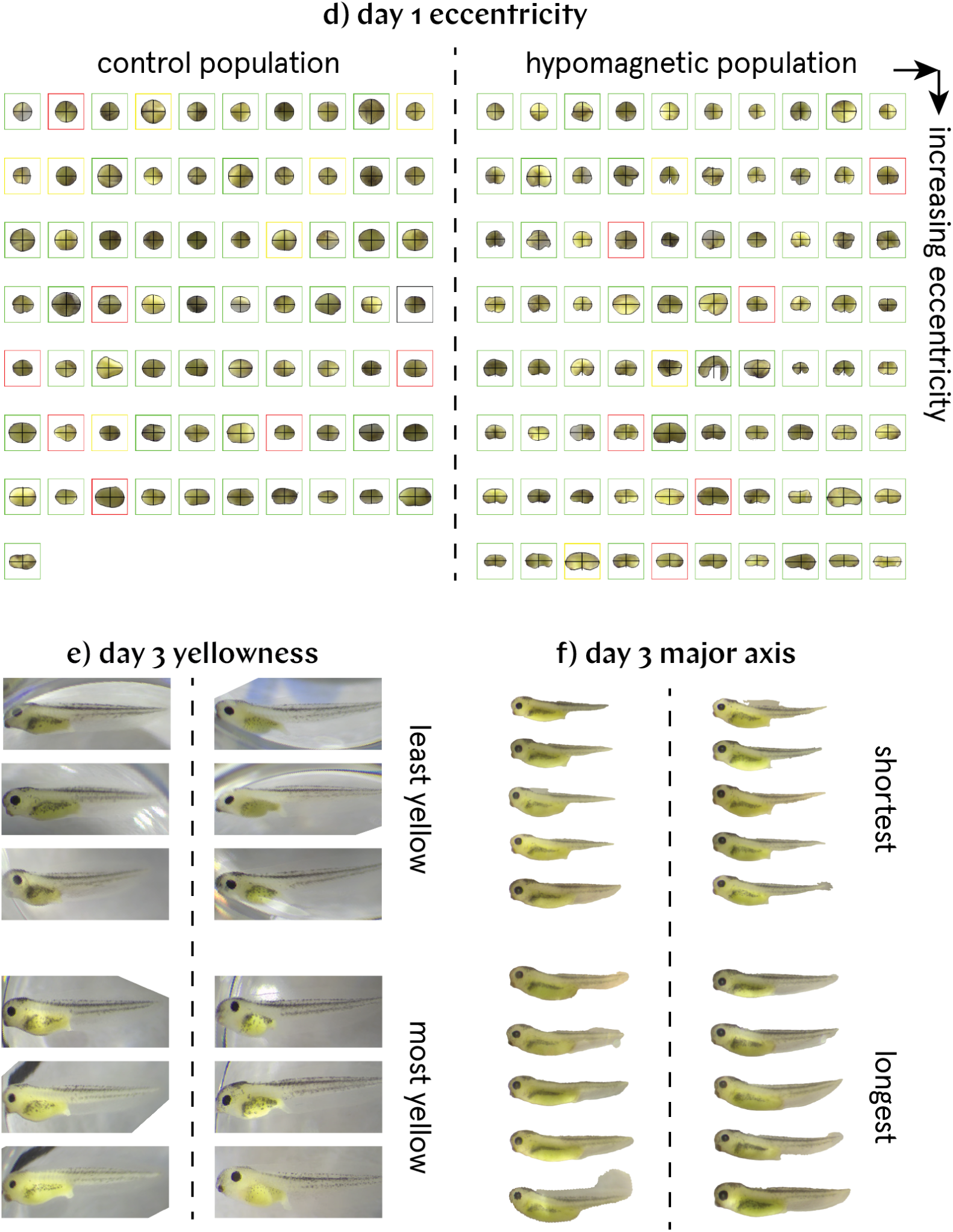
Environmental conditions are well-controlled, and the differences between control and hypomagnetic populations are qualitatively evident. a) The hypomagnetic chamber is equipped with both AC (copper shell) and DC (mu-metal) shielding, and x and z coils for positive control runs; the sample placement is shown. b) Temperature calibration reveals nearly identical temperatures within the hypomagnetic chamber and the control box, with a maximum temperature difference of 0.45°C during positive control runs. c) Representative images depict the 3-day development of individually tracked tadpoles, totaling over 2,750 individuals. d) Day 1 images of batch B2, ordered by increasing eccentricity, show that the hypomagnetic population exhibits significantly higher eccentricity. e) Least and most yellow tadpoles of batch B2 on day 3; those from the hypomagnetic population exhibit noticeably less yellow pigmentation. f) Shortest and longest tadpoles of batch B7 on day 3; the hypomagnetic population tends to be longer, suggesting accelerated developmental progression.

The hypomagnetic chamber, depicted in panel a) of Fig. 1, is a multi-layered metal enclosure designed to block ambient magnetic fields primarily through passive shielding. Its residual magnetic field is measured at less than 1 nT. Details on the chamber and the magnetic calibration procedure are provided in Supplementary Section SI9. Between batches B5 and B6, the chamber, which initially shielded only against DC and low-frequency fields using mu-metal, was upgraded with a permanent copper shell to block AC fields as well. Calibration curves thus refer to either the initial configuration (‘DC shielding’) or the final configuration (‘AC+DC shielding’). Tadpole rates of embryogenesis inside the chamber before and after copper shielding installation are similar, adding strength to the proposition that the observed biological effects are due to DC field shielding. Tadpoles in the control group are kept inside a closed, but switched off, cell incubator. This environment is referred to as the ‘control box’ (see Supplementary Section SI10); Earth’s weak DC magnetic field permeates the control box. Batches B1–B7 are regular experimental runs; batches denoted +1–3 are positive control runs, where a magnetic field mimicking Earth’s is artificially applied inside the hypomagnetic chamber (see Supplementary Section SI11).

Environmental factors that are known to perturb embryogenesis were recorded for the two environments. Sensors were excluded from the experimental environments during runs to prevent interference with the ambient magnetic field, so all calibrations were conducted separately under carefully replicated conditions. A thorough discussion about the effects of environmental factors, including a comparison of humidity, pressure, and light intensity and spectrum is provided in Supplementary Section SI12. Light levels are low, with 0.0873 *±* 0.0003 lux in the control box and 0.0751 *±* 0.0002 lux in the hypomagnetic chamber. Curves of temperature are presented in panel b) of Fig. 1.

Temperature is a critical factor regulating the speed of *Xenopus* embryogenesis: Warmer temperatures accelerate development [13]. Before installation of the AC shielding (batches B1–5), the temperature was up to half a degree lower in the hypomagnetic chamber than in the control box. After the AC shielding was added (batches B6–7), this temperature difference shifted slightly, with the hypomagnetic chamber now being marginally warmer by 0.10–0.15°C. In positive control runs (batches +1–3), the heat generated by the coils further increased the temperature difference to approximately 0.45°C. All those figures are conservative; actual temperature differences are likely less pronounced due to thermalization with the lab environment (details in Supplementary Section SI12).

After thoroughly assessing temperature, light levels, humidity, and pressure between the hypomagnetic chamber and the control box, we conclude that any observed significant effects are most likely due to the difference in magnetic field conditions between them.

In our experiments, we detect significant variability in the developmental speed of embryos across different weekly batches. This variability does not appear to arise solely from the use of different females as the egg source every week (see Supplementary Section SI6). Although not extensively documented in the literature—with the exception of one comprehensive study [14], which reaches a similar conclusion—we suspect that such variability is a common occurrence that remains largely unreported; we have reported and quantified it in a way that we believe is adequate and meets or exceeds the standards of the field.

Despite the large batch-to-batch variability, we can qualitatively distinguish the control from the hypomagnetic population as early as day 1 p.f. These observations are evident in panels d) through f) of Fig. 1. First, day 1 p.f. embryos in the hypomagnetic chamber exhibit greater eccentricity, suggesting accelerated development. d) depicts all day 1 images of batch B2 arranged by increasing eccentricity. By 24 hours p.f., most embryos in the hypomagnetic population have progressed beyond the circular-shaped egg stages, exhibiting a more elongated, bean-like shape. Day 1 images have frames color-coded based on the health assessment of the tadpoles by day 3 (see Supplementary Section SI7: Green indicates healthy tadpoles; orange indicates one health condition; red indicates multiple health conditions; and black indicates those that were deceased by day 1 p.f.). The images also show the major and minor axes of each embryo, as determined by the algorithm to calculate eccentricity described in Section 4. Second, hypomagnetic tadpoles appear paler in color in both day 2 and day 3 images. e) displays the three least and three most yellow healthy tadpoles for each condition from batch B2 by day 3, as identified by the algorithm detailed in Section 5. Third, hypomagnetic tadpoles tend to be longer and are typically at more advanced developmental stages compared to control tadpoles. f) highlights this point by depicting the five shortest and the five longest healthy tadpoles for each condition from batch B7 by day 3, after segmentation by the algorithm described in Section 6.

We then conducted the following gross morphological analysis, including positive control runs, to quantify the qualitative differences observed. Throughout our study, we aimed to minimize human intervention by ensuring that our image analysis is nearly fully automated (with a few exceptions detailed in Supplementary Section SI13). In addition, we propose that key morphological measures, derived from Nieuwkoop and Faber (NF) [15] stage-labeled images from reliable sources such as Xenbase [16], could be effectively used to characterize embryo developmental stages; we provide select examples to support this approach in the Supplementary Information (see Supplementary Figs. SI10, SI12, SI14, SI16, SI33, SI35, and SI37). We also aimed to provide multiple objective measures to corroborate each morphological observation. While the absolute magnitude of the effect depends on the measure chosen (for example, in the data of Section 4, a 20% increase in eccentricity corresponds roughly to a 1% decrease in solidity; both measures are good predictors of the same process of accelerated development, see Supplementary Figs. SI9 and SI15), we also consistently calculated a range of effect-size and effect-change metrics; these metrics are magnitude-insensitive because they standardize differences, being mostly reliable statistical indicators of the morphological distinctions between the control and hypomagnetic tadpole populations, independent of absolute scale. Our confidence in the presented results thus arises from a comprehensive analysis involving 19 distinct morphological measures, four effect-size metrics, and four effect-change metrics, applied across three days of imaging and tracking over 2,750 individual tadpoles.

## 4. The hypomagnetic field increases the eccentricity of day 1 embryos

At day 1 p.f., we observed frog embryos at various developmental stages, which we determined to range from NF stages 18 to 24 with the help of a standard table [15, 16]. During these stages, the embryos transition in shape from egg-like to bean-like and eventually to a more elongated, worm-like form.

To quantify our qualitative observation that frog embryos in hypomagnetic conditions exhibit accelerated development compared to control embryos already by this time point, we calculated morphological metrics assessing deviation from circularity—namely, eccentricity, elongation, roundness, and solidity. All four metrics consistently confirm our initial qualitative observation of advanced development under hypomagnetic conditions; they are detailed in Supplementary Section SI17, with a particular focus on eccentricity in the main text. Our code fits an ellipse to each day 1 p.f. image, defining the embryo’s eccentricity as 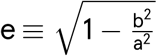, where b and a are the minor and major axes of the fitted ellipse, respectively.

Results for the eccentricity of day 1 images are presented in Fig. 2. A detailed guide on how to read the analysis plots is available in Supplementary Section SI74. Due to significant batch-to-batch variability, we present the normalized plots in the main text, where each batch is scaled relative to the average control value. The non-normalized data are presented in the Supplementary Information. In the figure, batches B1–7 correspond to experimental runs, while batches +1–3 represent positive control runs, where a magnetic field simulating Earth’s is applied within the hypomagnetic chamber (see Supplementary Section SI11). Throughout the plots, magnetic field conditions are denoted as C for control and H for hypomagnetic. The plots alternate between results for all tadpoles and for those classified as healthy based on the human scoring criterion described in Supplementary Section SI7.

**Fig. 2:**
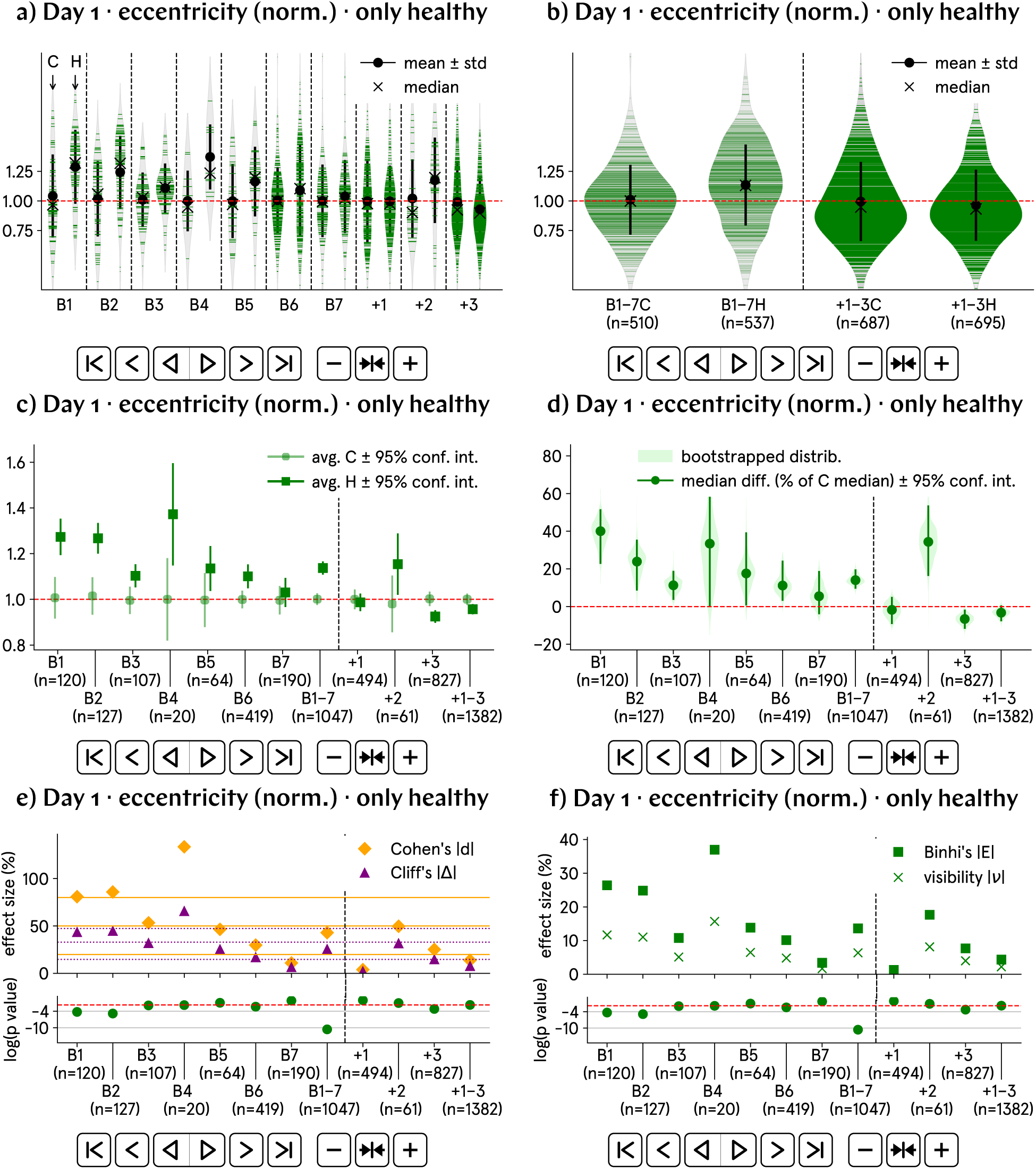
At day 1 p.f., the eccentricity of hypomagnetic embryos is approximately 20% higher than that of the control population. a) and b) Normalized eccentricity by batch and aggregate data; B1–7 represent experimental runs, and +1–3 correspond to positive control runs, in which an artificial magnetic field inside the hypomagnetic chamber simulates Earth’s. c) and d) Analysis of the population averages and medians further support the observed 20% increase in eccentricity. The abscissa points labeled B1–7 and +1–3 represent the aggregate values for the experimental and positive control runs, respectively. e) and f) Effect size metrics, namely Cohen’s d and Cliff’s Δ, and Binhi’s E and the visibility ν, confirm a medium to large effect, consistently reflecting the 20% level difference.

In the a) and b) violin plots, each tadpole’s value is represented by a horizontal line. The violin width reflects the kernel density estimation of the data distribution at that value. It is quantitatively evident that eccentricity is higher for tadpoles in hypomagnetic conditions, indicating a more elongated shape. In the non-normalized version of these plots, available in Supplementary Section SI19, we provide predicted tadpole stages based on elongation from a few stage-labeled reference images found in Xenbase [16].

The eccentricity in tadpoles exposed to hypomagnetic conditions exceeds that of control tadpoles by nearly 20%. Plot c) illustrates the average eccentricity for both control and hypomagnetic conditions across batches, along with their 95% confidence intervals (as detailed in Supplementary Section SI16). The standard deviation of the mean plotted in a) and b) reflects the variability or dispersion in the data; in contrast, the 95% confidence interval in c) emphasizes the accuracy of the estimate, indicating the range within which the true population mean is likely to fall with 95% confidence. The aggregate values for all experimental runs (B1–7) and all positive control runs (+1–3) are also shown. For normalized plots, each data point is adjusted relative to its corresponding control within the batch, while for non-normalized plots, the raw data is used despite batch-to-batch variability. Data aggregation is performed prior to statistical analysis.

Our dataset is generally non-parametric, so we also examine the median as a robust measure of central tendency. In plot d), we apply bootstrapping (explained in Supplementary Section SI16) to estimate the median difference in eccentricity between the control and hypomagnetic groups. The median eccentricity for tadpoles exposed to hypomagnetic conditions exceeds that of the control group by also nearly 20%.

In plots e) and f), we evaluate several measures of effect size (details in Supplementary Section SI14). Notably, both Cohen’s *d* and Cliff’s Δ (shown in e)), which are commonly used in biostatistics [17], tend to produce large and potentially overestimated values in our dataset. In contrast, Binhi’s *E* (shown in f)), a metric specifically designed for weak magnetic field effects in biology [1], more closely aligns with the net effects observed in plots c) and d); it estimates that the effect size of the hypomagnetic condition on eccentricity in day 1 embryos is again approximately 20%. Binhi’s *E* quantifies the effect size by comparing the difference in the groups’ means relative to their combined variability, accounting for both the magnitude and uncertainty of the measurements. We find that the visibility *ν* (also shown in f)) reliably yields a conservative estimate of effect size. Visibility *ν* is analogous to signal-to-noise, as it expresses the relative difference in the groups’ means as a percentage of their sum, providing a conservative estimate when the groups are close in magnitude. Below each plot, we also display the p value of a statistical test comparing the control and hypomagnetic populations (details in Supplementary Section SI74); points below the red dashed line indicate that the differences are statistically significant at the 0.01 level.

We provide a quantitative comparison of the observed accelerated development in terms of NF canonical stage numbers [15] in the Supplementary Information, using all day 1 measures of deviation from circularity—namely, eccentricity, elongation, roundness, and solidity (see Supplementary Figs. SI10, SI12, SI14, and SI16). These metrics were calculated from a few NF stage-labeled images sourced from Xenbase [16], offering an objective method to assess embryonic developmental stages.

The variability in major axis length for control embryos in day 1 p.f. images closely aligns with the 38% maximum difference reported in [14] for over 2,000 similar stage embryos. In contrast, the hypomagnetic embryos exhibit approximately 20% greater variability (see discussion in Supplementary Section SI63).

## 5. The hypomagnetic field decreases the yellowness of day 2 tadpoles

We qualitatively observed a difference in the yellow coloration of tadpoles which were exposed to hypomagnetic conditions. The yellow coloration in tadpoles is due to the presence of carotenoid and pteridine pigments [18] (one study also cites flavin pigments [19]), which absorb light in the blue spectrum and reflect yellow and orange wavelengths, resulting in the tadpoles’ characteristic yellow hue. After confirming that the ambient light environment was similar between the hypomagnetic and control groups (see Supplementary Fig. SI8 for light intensity and spectral comparison), we inquired whether the magnetic field itself could be directly related to these observed differences.

To quantify the color differences, we calculated several color metrics related to yellowness: L*a*b* yellowness, RGB yellowness, the Yellowness Index, and HSV yellowness. These metrics are described in detail in Supplementary Section SI26, with a primary focus on L*a*b* yellowness in the main text. All four metrics indicated distinct color development under hypomagnetic conditions. Although color differences are quantitatively evident in both day 2 and day 3 images, our analysis is exclusively for day 2 images. This is because day 3 images were taken after tadpole fixation with formalin, a process known to alter tissue color.

The L*a*b* color space [20] is designed for perceptual uniformity, meaning that small changes in objective color correspond to similar perceived differences. Unlike traditional color spaces such as RGB or CMYK, L*a*b* separates luminance from chromatic information, making it widely used in image processing for more accurate color representation. The b* channel, in particular, captures the blue-yellow axis, with positive values indicating increased yellowness. To simplify analysis, in Fig. 3 we present results for the b* channel—referred to as L*a*b* yellowness—in normalized plots, with each batch scaled relative to the average control value; the non-normalized data is presented in the Supplementary Information.

**Fig. 3:**
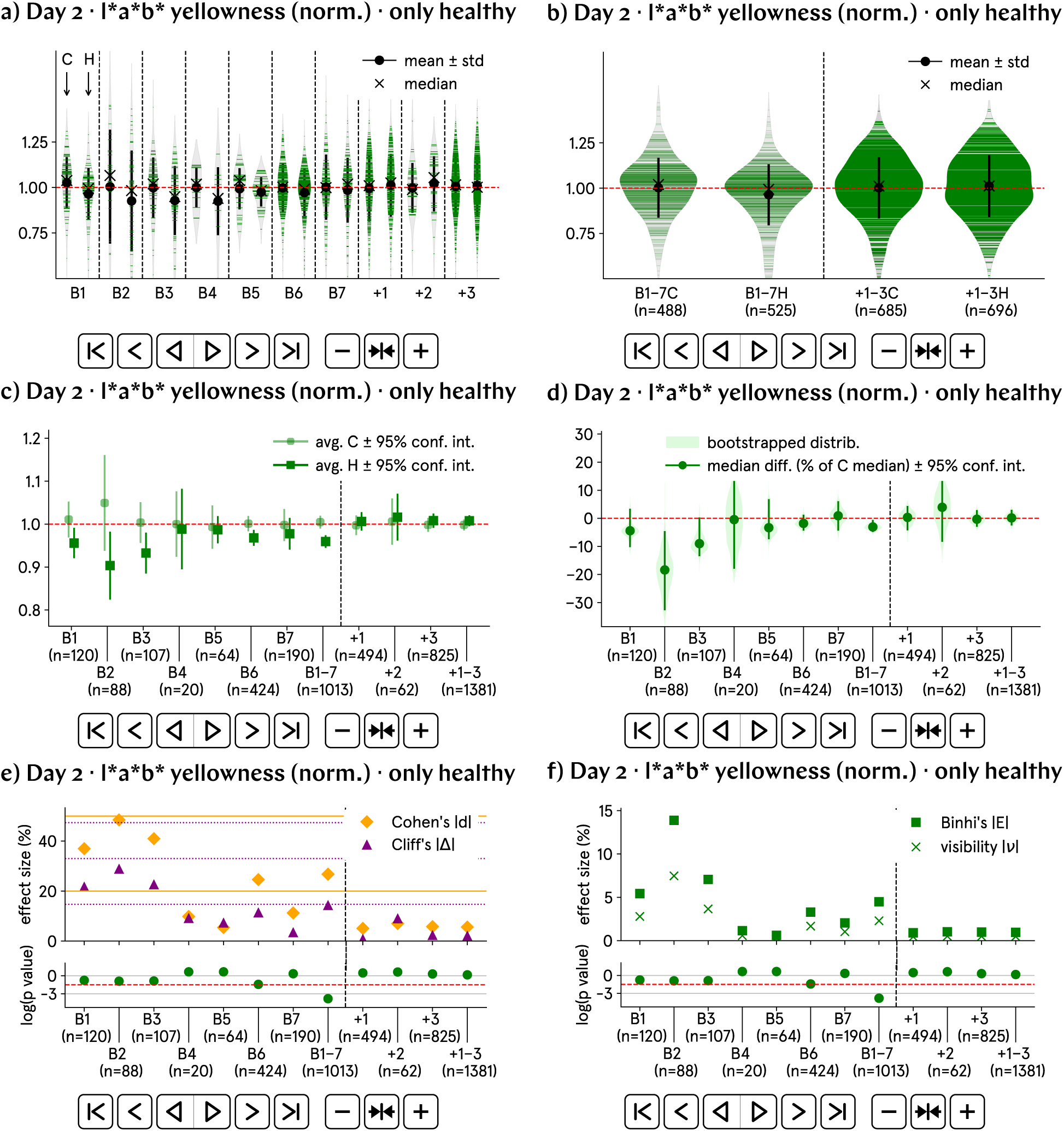
At day 2 p.f., the L*a*b* yellowness of hypomagnetic embryos is approximately 5% lower than that of the control population. a) and b) Normalized L*a*b* yellowness by batch and aggregate data; B1–7 represent experimental runs, and +1–3 correspond to positive control runs, in which an artificial magnetic field inside the hypomagnetic chamber simulates Earth’s. c) and d) Analysis of the population averages and medians further support the observed 5% decrease in L*a*b* yellowness. The abscissa points labeled B1–7 and +1–3 represent the aggregate values for the experimental and positive control runs, respectively. e) and f) Effect size metrics, namely Cohen’s d and Cliff’s Δ, and Binhi’s E and the visibility ν, confirm a small to medium effect, consistently reflecting the 5% level difference.

The violin plots a) and b) consistently show lower L*a*b* yellowness for tadpoles in hypomagnetic conditions. This effect, observed at the 5% level for day 2 images, is confirmed by both the average and bootstrapped median values plotted in c) and d), respectively, as well as by the magnitude of Binhi’s *E* in f). Importantly, no color difference is seen in the positive control tadpoles, which were raised inside the hypomagnetic box with an applied magnetic field mimicking Earth’s. This result supports a mechanism through which the observed color change is driven primarily by the altered magnetic field conditions.

To investigate the cause of reduced yellow pigmentation, we examine the sources of pteridine and carotenoid in *Xenopus* embryos.

Tadpoles are capable of synthesizing pteridines as early as NF stages 39–40 [21], which occur well after the 2 day p.f. mark; therefore, synthesized pteridines could not account for the observed color changes already visible in day 2 p.f. images.

Carotenoids are either deposited in the egg during oogenesis or ingested from the diet after hatching. Since tadpoles do not begin feeding until well after day 3 p.f., all carotenoids present at this stage must originate from the yolk. Carotenoid content is reduced in amphibians that have undergone pituitary gland removal, leading to paler coloration [19]. In *Xenopus* tadpoles, the pituitary gland begins to form around NF stages 24–25 [15], *i.e.*, before our day 2 images are captured. However, the gland only starts to secrete hormones between stages NF 46–48 (5–6 days p.f.) [15], which is beyond the time frame of our study. Consequently, we do not believe the color differences we observe are of hormonal origin.

The visible changes in color during early development might thus reflect the metabolic use of the yolk’s contents, including the pigments [22]. The specific concentrations of carotenoids and pteridines in *Xenopus* yolk are not well-documented and can vary depending on factors like maternal diet, environmental conditions, and species [21]. The major yolk protein, vitellogenin, is a lipoprotein that acts as a nutrient source for the developing embryo. While vitellogenin itself lacks color, it often binds to carotenoids, which contribute to the yellowish hue of the yolk [23]. As vitellogenin is processed into yolk proteins like lipovitellin and phosvitin during embryogenesis, and the yolk is gradually consumed, the yellow color fades due to the depletion of carotenoids. This process may explain the slightly decreased yellow hue we observe in hypomagnetic tadpoles, which exhibit morphological markers indicative of accelerated embryonic development.

We did not assess the levels of the more commonly studied eumelanin pigment [24, 25]. At day 2 p.f., eumelanin levels remained low. While the binarization procedure detailed in Supplementary Section SI26 improved the accuracy of yellow pigment assessment, it was less reliable as a good discriminator of dark pigments like eumelanin.

## 6. The hypomagnetic field increases the length of day 3 tadpoles

The most striking evidence of accelerated development is the increased length of day 3 tadpoles exposed to hypomagnetic field conditions.

Day 3 tadpole images were segmented, and their major axes were calculated (as detailed in Supplementary Sections SI13 and SI35). The results are displayed in Fig. 4, with each batch normalized to the average control value. We opt not to present non-normalized data due to slight differences in magnification across batches.

**Fig. 4:**
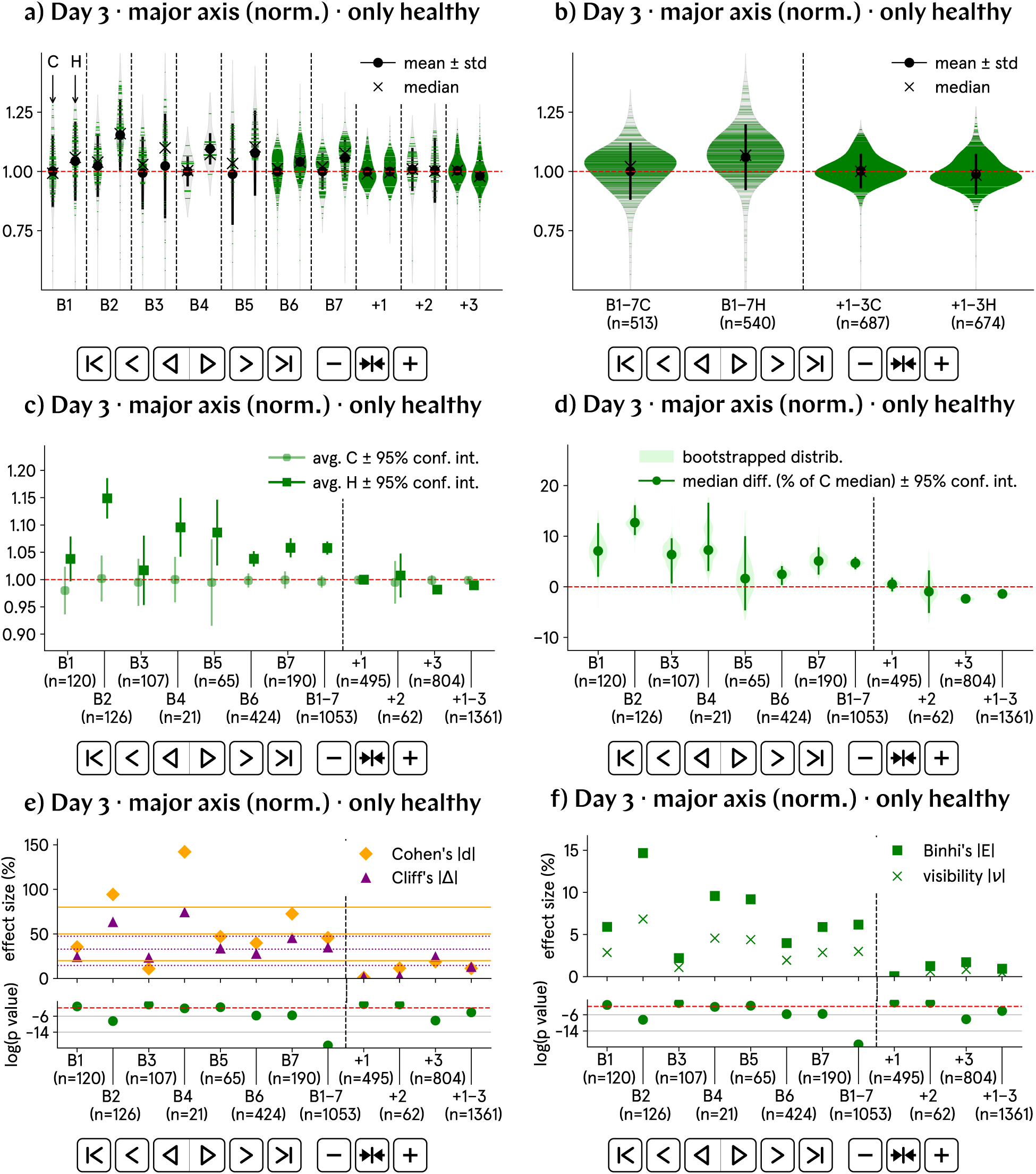
At day 3 p.f., the major axis of hypomagnetic embryos is approximately 7.5% higher than that of the control population. a) and b) Normalized major axis by batch and aggregate data; B1–7 represent experimental runs, and +1–3 correspond to positive control runs, in which an artificial magnetic field inside the hypomagnetic chamber simulates Earth’s. c) and d) Analysis of the population averages and medians further support the observed 7.5% increase in major axis. The abscissa points labeled B1–7 and +1–3 represent the aggregate values for the experimental and positive control runs, respectively. e) and f) Effect size metrics, namely Cohen’s d and Cliff’s Δ, and Binhi’s E and the visibility ν, confirm a medium effect, consistently reflecting the 7.5% level difference.

Plots a) and b) illustrate the distribution of major axis lengths, with b) showing a distinct and quantitatively different violin shape for the hypomagnetic population.

The mean, shown in c), and the bootstrapped median, in d), both indicate an approximately 7.5% increase in major axis length for the hypomagnetic population. This result aligns with Binhi’s *E* reported in f). Additionally, both Cohen’s *d* and Cliff’s Δ, plotted in e), classify this effect as medium in size. Notably, this effect is absent in the positive control tadpoles.

Additional morphological measures, namely perimeter, area, total curvature, solidity, elongation, and roundness (the latter reflecting the relationship between area and perimeter), as detailed in Supplementary Section SI35, consistently indicate more elongated and complex shapes in tadpoles exposed to the hypomagnetic conditions. These findings further reinforce the major axis results presented here.

We present a quantitative comparison of the observed accelerated development in terms of NF canonical stage numbers [15] in the Supplementary Information, using all day 3 measures that can be accurately reported in their non-normalized form (*i.e.*, without normalization by the control group average)—namely, solidity, elongation, and roundness (see Supplementary Figs. SI33, SI35, and SI37). Using NF stage-labeled images from Xenbase [16], we once more propose this as an objective approach for assessing embryonic development.

No evident difference in major axis is observed in day 2 images, as detailed in Supplementary Section SI63. We hypothesize that this is due to the reduced robustness of segmentation in day 2 images, likely caused by the pale coloration of the tadpoles, which may affect the accuracy of the major axis analysis. Additionally, the imaging conditions on day 2 were less controlled, with live tadpoles that were not manipulated, potentially leading to distortions or tadpoles being out of the focal plane. In contrast, on day 3, the fixed tadpoles were carefully positioned with tweezers to lie flat and on their sides, yielding more reliable measurements.

## 7. The hypomagnetic field alters the morphological progression of tadpoles between days 2 and 3

In Fig. 5, we compare the day 2 to 3 progression of four key morphological properties of tadpole development, namely major axis length, curvature, convexity, and solidity. A Z-score metric for effect change normalizes the data by accounting for variability, allowing us to quantify changes in each property in relation to the mean values for both days (see Supplementary Section SI15). A positive Z-score difference indicates a greater deviation from the mean on day 3 compared to day 2. Although Z-score differences are our primary measure, additional metrics such as effect change visibility, percentage change, and relative difference were also calculated; they are described in detail in Supplementary Sections SI15 and SI46, and corroborate the trends observed in the Z-score analysis.

**Fig. 5:**
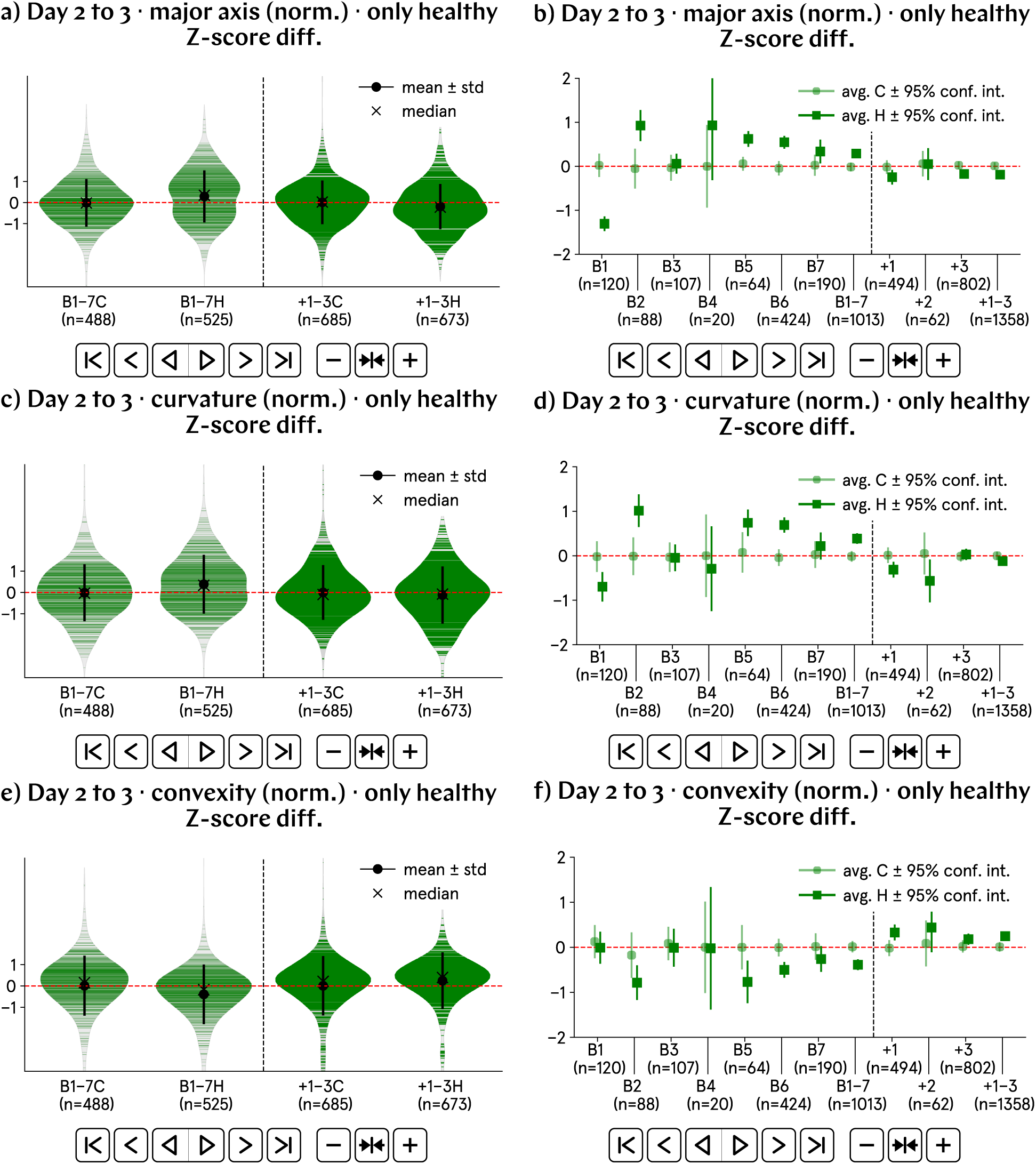
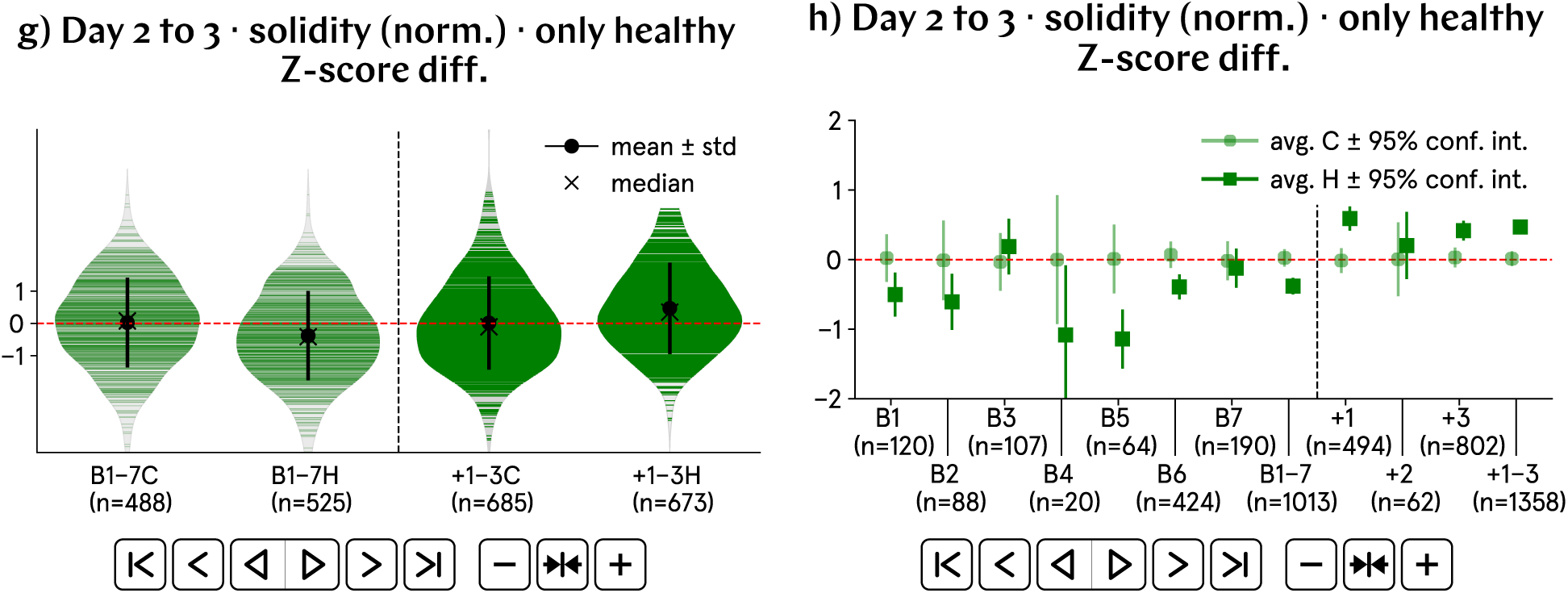
From day 2 to 3 p.f., the hypomagnetic conditions significantly influence the morphological progression of the tadpoles. This progression is quantified, for each population (control or hypomagnetic), through differences in Z-scores at days 2 and 3 p.f. For the progression of the four morphological properties studied, the control and hypomagnetic populations differ by approximately 0.5 units of standard deviation, indicating a significant shift in the hypomagnetic population’s data. The abscissa points labeled B1–7 and +1–3 represent the aggregate values for the experimental and positive control runs, respectively. a) and b) The major axis length in the hypomagnetic population exhibits a Z-score difference of approximately 0.5 higher than that of the control population. c) and d) A similar Z-score difference of about 0.5 higher for the total curvature is found in the hypomagnetic group. e) and f) In contrast, the convexity in the hypomagnetic population shows a Z-score difference around 0.5 lower than in the control group. g) and h) Solidity in the hypomagnetic population also has a Z-score difference nearly 0.5 lower than in the control. The higher Z-scores for major axis length and total curvature suggest accelerated morphological development under hypomagnetic conditions, while the lower Z-scores for convexity and solidity indicate a faster increase in shape complexity.

Plots a) and b) illustrate a predominantly positive Z-score difference for the major axis in the hypomagnetic group compared to the control. Plots c) and d) similarly reveal an overall positive Z-score difference for the total curvature, despite greater variability across batches. In contrast, convexity and solidity Z-score differences, depicted, respectively, in plots e) and f) and in plots g) and h), are lower for the hypomagnetic population. Across the four morphological properties analyzed, the Z-score difference metric reveals a divergence of approximately 0.5 units of standard deviation between the control and hypomagnetic populations, indicating a significant shift in the data associated with the hypomagnetic group.

The higher Z-score differences for major axis length and total curvature suggest accelerated morphological development under hypomagnetic conditions (as qualitatively observed in tadpole length and curvature differences between day 2 and 3 p.f. images in Fig. 1 c)). We interpret the average decrease in convexity and solidity Z-score differences as an indication of a faster increase in tadpole shape complexity. Lower convexity correlates with more indentations and protrusions, while lower solidity points to a more fragmented or irregular structure.

## 8. Discussion

After quantitatively analyzing over 8,000 images of nearly 3,000 viable tadpoles across 19 morphological measures, using four effect-size and four effect-change metrics, we conclude that magnetic fields as weak as the Earth’s can have a macroscopic impact on biological processes such as embryo development.

We have demonstrated the existence of a link between Earth’s minute DC magnetic field and basal tadpole physiology regulation that is present as early as one day p.f., for purposes likely *not* related to orientation, navigation, or homing. In the absence of the geomagnetic field, embryo development accelerates, a trend we observed and tracked over a three-day period in our experiments.

### Potential influence of magnetic field variations

We cannot entirely dismiss the possibility that variations in magnetic field strength—*i.e.*, those occurring when plates are inserted or removed from the hypomagnetic chamber—may have influenced the baseline physiology of the tadpoles, rather than the constant hypomagnetic environment. This unlikely scenario would require a lasting biological response to short-term magnetic field variations.

### Developmental response to the hypomagnetic field

It remains unclear whether *Xenopus* embryos sense Earth’s magnetic field for a specific developmental purpose, or if their physiological response to hypomagnetic conditions simply reflects a deviation from an evolutionary adaptation that is arguably optimal for the geophysical environment in which the species evolved. Nonetheless, it is striking that in the absence of the geomagnetic field, the tadpoles consistently exhibit accelerated developmental speed, rather than a broad or indiscriminate alteration in developmental rhythms. This uniform response suggests the existence of a coordinated mechanism controlling the rate of development that includes magnetic cues. In rare instances, our data might be better modeled as a two-state system, which is consistent with responding and non-responding populations. These cases correspond to plots of type b) in which the aggregate violin suggests two distinct peaks in width, indicating a bimodal distribution.

### Implications of accelerated embryogenesis

We speculate whether the observed accelerated development under hypomagnetic conditions could, in principle, increase the incidence of embryo abnormalities and/or carcinogenesis. While no such effect has been detected in our data, it is possible that faster growth inherently raises the risk of developmental errors, as it has been previously reported in studies on the biological effects of hypomagnetic fields [1, 6, 26]. For example, hypomagnetic conditions have been linked to disturbances in the mitotic spindle [6], aligning with previous findings that weak magnetic fields alter cytoskeleton dynamics [1, 2, 7]. Consistent with our results, there are two literature reports that link the application of a weak magnetic field (rather than the elimination of a weak magnetic field)—specifically, 25 *µ*T and 2 mT, albeit at low AC frequencies—with slowed development (rather than accelerated development) in *Xenopus* [27, 28].

### Pigmentation changes and carotenoid depletion

There is one literature report linking the exposure to a weak magnetic field (of strength 0.5 mT) to an increase in eumelanin (dark) pigmentation in tadpoles [29]; while our study focused on yellow pigments, that result mirrors our own findings. We hypothesized that the depletion of yolk carotenoids could explain the paler-yellow appearance observed in hypomagnetic tadpoles. Carotenoids are known to be degraded by reactive oxygen species [30]. Overall levels of reactive oxygen species have been documented to increase under hypomagnetic field environments [31, 32]; an elevated reactive oxygen species concentration may thus have contributed to the observed reduction in carotenoid levels.

The effects of weak magnetic fields in biology have often been overlooked, including in studies of native organismal behavior, human health, and space exploration, as well as in guidelines for electromagnetic exposure. The rationale has been that the biological effects of weak magnetic fields are negligible because the field strengths are thought to be too small to trigger a biophysical sensing mechanism. Given the results of this study, however, this reasoning is flawed. Further investigation is therefore warranted, especially the use of omics and molecular techniques, to identify the nature of changes likely to be caused by the presence or absence of weak magnetic fields.

The fact that living beings can respond to magnetic field shifts of this magnitude thus usually comes as a surprise. To explain our findings, we note that organisms are traditionally thought to sense weak DC magnetic fields via three mechanisms (see Supplementary Section SI77).

In a small minority of cases, most notably in magnetotactic bacteria, cells contain biogenic magnetite crystals, which allow the organism to respond to magnetic fields like a classical compass [33]; these crystals must be large enough to overcome thermal motion and align with Earth’s magnetic field [34, 35]. To date, there is only one report of ferrimagnetic material found in an amphibian, specifically in the newt [36]. Given that *Xenopus* is one of the most extensively studied biological model organisms worldwide, we find it unlikely, but not impossible, that such a magnetite-based magnetosensing mechanism is at play in this species, as cells containing large particles of magnetite would have likely been identified by now.

In another minority of instances, organisms possess specialized organs that function as conductive channels. When these channels intersect magnetic field lines, a voltage is induced according to the classical law of electromagnetic induction; such a voltage is then sensed by the organism. This is believed to explain how sharks and rays detect magnetic fields [37]. Given the structural complexity involved, we find it very unlikely that similar systems exist on day 1 p.f. *Xenopus* embryos, which contain fewer than 50,000 cells [38], yet are already capable of detecting the absence of Earth’s magnetic field. In addition, the detection of DC magnetic fields through this mechanism requires the organism to move and cross magnetic field lines, and *Xenopus* embryos are not mobile until at least day 2 p.f.

In most cases, however, scientists have been unable to explain how weak magnetic fields impact biology, especially because the effects do not necessarily get larger when the magnetic field strength is increased, and are broadly nonspecific, occurring across diverse species and cell types [1, 2, 5, 8]. A chemical explanation that aligns with these observations exists but remains unverified microscopically and in living cells: electron spin-dependent chemical reactions [9, 39–41]. These reactions bear striking similarities to the simplest form of room-temperature technological quantum sensing in the solid-state [42, 43] and, at least *in vitro*, can detect magnetic fields using electron spin superpositions—even at room temperature [44]. In this hypothesis, organisms ‘sense a magnetic field’ by detecting changes in the physiological concentrations of reaction products, as the reaction rate responds to the magnetic field.

This hypothesis, which we favor, would itself be a surprise. If further experiments demonstrate that these or similar findings [45, 46] are a result of quantum sensing-like chemical reactions happening within living cells, the implications are major (see Supplementary Section SI78 for details). This will mean, first, that electron spin superpositions survive the strongly decohering environment of the cell for long enough to be used for function (*e.g.*, to sense and physiologically react to Earth’s weak magnetic field); and second, that weak magnetic fields could, in principle, be quantum-engineered to influence the endogenous spin physics of biological matter, thereby enabling the up- and down-regulation of the machinery of the cell. While the effects reported so far remain largely nonspecific, enough knowledge of the spin physics within the specific proteins sustaining electron spin-dependent chemical reactions *in vivo* could, in principle, allow the targeted and predictable manipulation of physiological processes with weak magnetic fields.

## 9. Conclusion

We individually tracked the development of more than 2,750 *Xenopus laevis* embryos for three days under extremely carefully controlled magnetic field conditions. It is our assessment that other environmental factors such as temperature and light were controlled to a level where their variation would not be the origin of the observed biological effects. Through robust statistical analysis, we observed clear morphological changes in the tadpoles during experimental runs under hypomagnetic conditions of less than 1 nT; these largely disappear in the positive control runs.

We conclude that exposure to a hypomagnetic environment accelerates tadpole development in a measurable way. Therefore, *Xenopus laevis* embryos can detect and respond to the absence of Earth’s minute DC magnetic field of approximately 50 *µ*T as early as day 1 p.f.

Our data is publicly available and sufficiently robust to advance the scientific consensus that weak magnetic fields indeed exert measurable biological effects.

## 10. Author contribution

Following the CRediT taxonomy [47], each author confirms contribution of:

1. Alessandro Lodesani: conceptualization; methodology; investigation; data curation; validation; formal analysis; software; writing (original draft); writing (review and editing); and visualization;
2. Geoff Anders: conceptualization; methodology; writing (original draft); writing (review and editing); project administration; and funding acquisition;
3. Lykourgos Bougas: investigation; data curation; and formal analysis;
4. Tobias Lins: investigation; data curation; and formal analysis;
5. Dmitry Budker: conceptualization; supervision; and funding acquisition;
6. Peter Fierlinger: conceptualization; investigation; supervision; and funding acquisition;
7. Clarice D. Aiello: conceptualization; methodology; data curation; validation; formal analysis; software; writing (original draft); writing (review and editing); visualization; supervision; project administration; and funding acquisition.

## 11. Acknowledgements

We wholeheartedly thank: Michael Levin, from Tufts University, for suggesting the original idea of the experiment to P. F.; Arielle Gabalski for help with initial lab setup; Gabriel E. Bertolesi, from the University of Calgary, for first bringing to our attention the differences in developmental speed between the two tadpole groups; Vera D. Aiello, from the University of São Paulo, for first bringing to our attention the color differences between the two tadpole groups; Marko Horb and Nikko-Ideen Shaidani, from the National Xenopus Resource at the Marine Biological Laboratory, for early tadpole assessment and continuous frog husbandry support; Gabriel E. Bertolesi, from the University of Calgary, Paul Blank, from the NIH, Douglas Brash, from Yale University, Richard Fuisz, Maria Ingaramo and Andrew York, from Calico, Rod Kunz, Divya Shastry, and Alexandra Wrobel, from MIT-Lincoln Laboratory, Sergey Polyakov, from NIST, and Graham Timmins, from the University of New Mexico, for their valuable feedback on the manuscript; James Wiles, from the Wolfram Institute, for insights into image segmentation; and Jennilyn and Bret Gaitan, Oliver Carefull, and Melinda Bradley for project support.

We acknowledge the use of ChatGPT for improving the readability of the text and assisting with writing and debugging python code.

This experiment was partially funded by grants from the Faggin Foundation and Nevin Freeman.

Two authors (C. D. A. and G. A.) partially funded the conducted research. The remaining authors have no conflicts of interest to declare.

The [Quantum Biology Institute](https://www.quantumbiology.org/) is a California non-profit (501(c)(3) status pending) focused research organization [48] that performs basic research underpinning the quantum biology field in an open-science fashion. It is part of the [Quantum Biology Ecosystem](https://www.quantumbiology.org/).

## Supplementary Information

### SI1. Frog husbandry

For each batch, two wild-type *Xenopus laevis* adult female frogs were used for egg harvesting. The frogs were sourced from the National Xenopus Resource at the Marine Biological Laboratory in Woods Hole, Massachusetts, and acclimated in the Los Angeles laboratory for at least two weeks. They were housed in fish tanks equipped with mechanical, chemical, and biological filtration systems. The tanks were filled with tap water. The pH of the water was adjusted to 7.4–7.8 using sodium bicarbonate, chlorine was neutralized with sodium thiosulfate, and reef salts were added to optimize water hardness to an optimal value of around 1600 (in units of mg/L of CaCO*_3_*). Nitrifying bacteria were used to maintain optimal nitrate and nitrite levels. Water quality was monitored at least four times per week, and partial or total water changes were performed periodically. The frogs were fed twice weekly with Nasco frog brittle.

No individual frog was used more than once to provide eggs for the experiment. Following egg harvesting, the two females were euthanized using an anesthetic overdose, ensuring a humane and efficient process, as described in [49]. The frogs were placed in a water bath containing 0.5% MS222, buffered to pH 7.0, for 30 minutes. Afterward, unresponsiveness was confirmed through a foot pinch test.

### SI2. Egg harvesting and fertilization protocol

Embryos were generated using the *in vitro* fertilization method detailed in [50], summarized here for convenience. Two days prior to fertilization, each adult female frog was primed with an injection of 50 U of pregnant mare serum gonadotropin (PMSG) in 0.5 mL phosphate buffer saline (PBS, density of 100 U/mL). The following evening (after 5 pm), the frogs were boosted with an injection of 500 U of human chorionic gonadotropin (hCG) in 0.5 mL PBS (density of 1000 U/mL). The next morning, eggs were collected in a Petri dish by physically squeezing the frogs.

Adult male frog testes were sourced weekly from the National Xenopus Resource and stored at 4*◦* C.

Before detailing the fertilization protocol, the required solutions and their preparation are as follows.

#### Marc’s Modified Ringer’s (MMR) solution

To prepare 1 L of 10x MMR, mix the following with 1 L of deionized (DI) water:

1. g NaCl (1 M)
2. g KCl (20 mM)
3. g MgSO_4_·7H_2O_ (10 mM)
4. g CaCl_2_·2H2_O_ (20 mM)
5. g Hepes (50 mM)

Adjust the pH to 7.4–7.8 using NaOH. Dilute to 1x MMR and 0.1x MMR as needed (1:10 and 1:100 ratios with DI water).

#### 2% L-cysteine

Dissolve 2 g of L-cysteine in 100 mL of 0.1x MMR and adjust the pH to *∼* 8.0 with NaOH.

For each fertilization procedure, half a testis was crushed just before egg collection in 1 mL of 1x MMR solution using a 1.5 mL plastic centrifuge tube and a plastic pestle.

Immediately after collection, the eggs were evenly spread as a single layer in a dish using a plastic pipette, and the testis solution was applied uniformly before being gently mixed. After allowing the mixture to sit for 5 minutes, the dish was flooded with 0.1x MMR and mixed again to ensure thorough coverage.

Twenty minutes later, the 0.1x MMR solution was removed and the dish flooded with the 2% L-cysteine solution. The eggs were left to sit for 5 minutes, and were periodically mixed with a plastic pipette to remove the jelly coat. Once the eggs clumped together, the cysteine was removed, and the dish was washed 3–5 times with 0.1x MMR before being flooded again with fresh 0.1x MMR.

### SI3. Egg randomization procedure

After fertilization, viable eggs were identified using a stereo microscope. Healthy eggs display a dark upper half and a small lighter dot indicating sperm entry. Poor-quality eggs appear swollen and pale.

Eggs were sorted into 24- or 48-well plates, with each well containing 1 mL of 0.1x MMR for the 24-well plates and 0.75 mL for the 48-well plates. One egg was placed per well to prevent cross-contamination and to enable the tracking of individual tadpoles over the three days of the experiment.

Two plates were filled simultaneously, alternating one egg per plate. Once filled, one plate was immediately placed in the hypomagnetic chamber and the other in the control box, both covered with plastic lids. This process continued until all viable eggs were sorted. The embryos, then at the 1- or 2-cell stage, were left undisturbed for 24 hours until data collection on the first day of the experiment.

### SI4. Daily image acquisition

One image of each tadpole was captured daily for three days, approximately at every 24 hours post-fertilization, with control images always taken first. Imaging for all plates was typically completed within one hour; the imaging time per plate was usually well under 10 minutes. Imaging was conducted one plate at a time, with only one plate removed from its environment at any given moment; each plate was promptly returned to its environment immediately after the imaging process.

While magnification was consistent for all images taken on a given day, slight variations between days were possible due to the analog magnification control.

Day 1 and day 2 images were taken from live tadpoles, whereas day 3 images were captured after the tadpoles were euthanized in formalin and fixed. The exception to that are day 3 pictures from batches B1–3, which were also taken with live tadpoles. After day 3 imaging, each tadpole was stored individually in labeled 1.5 mL centrifuge tubes for potential follow-up work. Samples are available upon request.

### SI5. Each batch at a glance

Relevant parameters for each batch can be found summarized in the table below:

**Table SI1:** Details for each batch. B1–7 are experimental runs; +1–3 are positive control runs.

| # | Fertilization | Temp. | Shielding | Res. field | Day 3 formalin? | Observation |
| --- | --- | --- | --- | --- | --- | --- |
| B1 | 2024/05/01 | ambient | DC | $\lesssim 10$ nT | no | 0.30–0.50°C ↓ in hypo |
| B2 | 2024/05/19 | ambient | DC | < 1 nT | no | 0.30–0.50°C ↓ in hypo |
| B3 | 2024/05/30 | ambient | DC | < 1 nT | no | 0.30–0.50°C ↓ in hypo |
| B4 | 2024/07/08 | lab AC, 70°F | DC | < 1 nT | yes | 0.30–0.50°C ↓ in hypo |
| B5 | 2024/07/24 | lab AC, 70°F | DC | < 1 nT | yes | 0.30–0.50°C ↓ in hypo |
| B6 | 2024/08/05 | lab AC, 70°F | AC+DC | < 1 nT | yes | 0.10–0.15°C ↑ in hypo |
| B7 | 2024/08/19 | lab AC, 70°F | AC+DC | < 1 nT | yes | 0.10–0.15°C ↑ in hypo |
| +1 | 2024/08/27 | lab AC, 70°F | AC+DC | $\sim 30$ μT | yes | 0.45°C ↑ in hypo |
| +2 | 2024/09/02 | lab AC, 70°F | AC+DC | $\sim 30$ μT | yes | 0.45°C ↑ in hypo |
| +3 | 2024/09/09 | lab AC, 70°F | AC+DC | $\sim 30$ μT | yes | 0.45°C ↑ in hypo |

‘AC’ in the Temperature (‘Temp.’) column means ‘air conditioning’; ‘AC’ in the Shielding column means ‘alternating current’. ‘Res. field’ stands for ‘residual field’ inside the hypomagnetic chamber. The column ‘Day 3 formalin?’ refers to whether day 3 images were taken after fixation with formalin.

Batch B1 was run without active compensation inside the hypomagnetic chamber (see Supplementary Section SI9), yielding a residual magnetic field of *<* 10 nT.

Before installation of the AC shielding inside the hypomagnetic chamber, the temperature was up to half a degree lower in the hypomagnetic chamber than in the control box. After the AC shielding was added, this temperature difference shifted slightly, with the hypomagnetic chamber now being marginally warmer (details in Supplementary Section SI12).

### SI6. Control: The effect of a particular female frog as egg source is not significant

To illustrate the high variability in the experiments, batch B6 was designed to control for the influence of eggs from different females on the results. Eggs from two different females were fertilized using sperm from the same male frog, with approximately 100 eggs from each female placed under each magnetic field condition. This setup allows for a robust comparison, as any differences observed are likely attributable to the female source, providing a strong control for the potential influence of a particular female on the results.

The violin plots of Supplementary Fig. SI1 display kernel density estimates for various morphological properties over the three days of the experiment; data obtained from embryos of each female are separated by the dashed black line. For each female, a Mann-Whitney [17] p value is provided when significant (here, p smaller than 0.01); otherwise, ‘n.s.’ indicates no significance. Differences between eggs from different females in the control condition are evaluated using the Kruskal-Wallis test [17], and the p value is displayed if it is smaller than 0.01. We only compare control groups across females, as it is not appropriate to compare the hypomagnetic condition between females due to differing effects that might arise from the environmental conditions themselves.

It is reassuring that nearly all morphological properties used to differentiate the control and hypomagnetic populations—except for roundness on day 1—show no statistically significant differences within the control group based on the female frog. This consistency within the control condition suggests that the observed differences between control and hypomagnetic populations are not influenced by variability among female frogs.

We hypothesize that eggs with either higher or lower overall health may exhibit varying susceptibility to the hypomagnetic environment, potentially leading to differential developmental effects. Further studies focusing on the initial health of the eggs and the health of the mother could provide valuable insights into this phenomenon.

**Fig. SI1:**
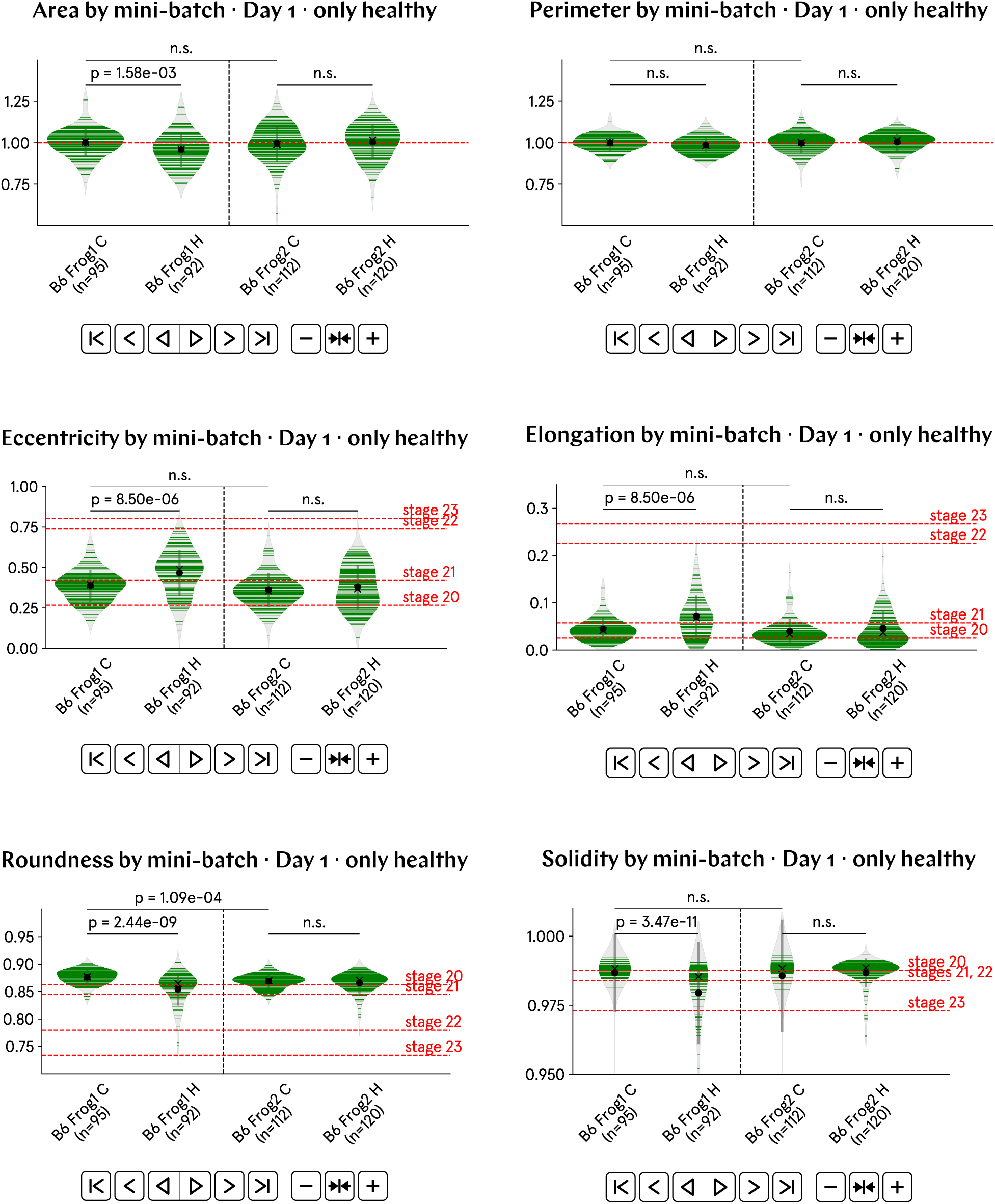
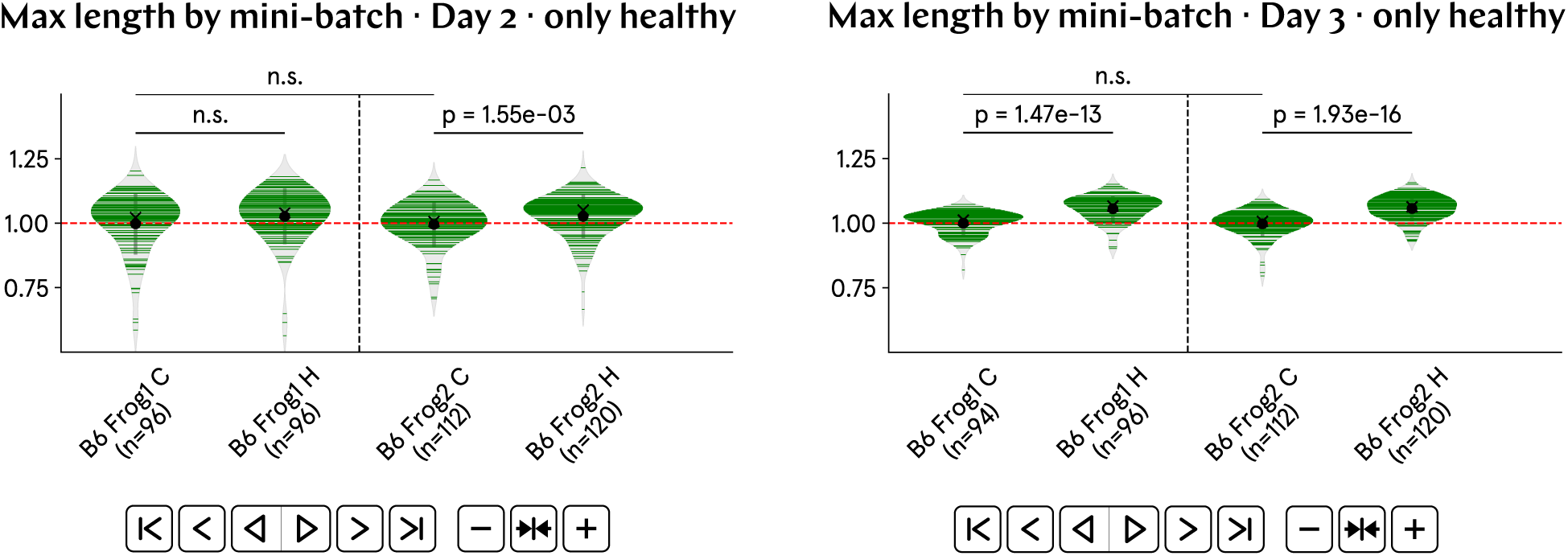
Maternal variability does not account for batch variability. Across the three days of the experiment, nearly all morphological properties examined above show no statistically significant differences within the control group, based on the female frog that provided the eggs. This consistency suggests that the observed distinctions between control and hypomagnetic populations are largely independent of maternal variability. Moreover, the week-to-week variability between batches cannot be solely attributed to different females providing the eggs.

### SI7. Control: The effect of magnetic field conditions on the overall health scores of the tadpoles is not significant

To evaluate the impact of the hypomagnetic condition on overall tadpole health, we implemented the only human-made scoring system used in this study. Each tadpole received a score: 1 for healthy, 2 for tadpoles with one morphological defect or adverse condition, 3 for those with multiple defects or adverse conditions, and 4 for tadpoles deceased by day 1 p.f. Descriptions of defects, if any are present, are provided above the tadpole images in the publicly available plate posters, which can be found in the GitHub folder 10 Posters and are detailed in Supplementary Section SI71.

Contrary to previous findings in the literature (*e.g.*, [6]), our results described below do not show that tadpoles raised under hypomagnetic conditions are statistically different from control tadpoles in terms of overall health based on the scoring scale described above. In other words, aside from an accelerated development, we do not observe that tadpoles raised in the hypomagnetic chamber are less healthy or exhibit a higher rate of abnormalities compared to the control group.

Based on the health scoring, the results displayed in Supplementary Fig. SI2 suggest that control and hypomagnetic tadpole populations exhibit remarkably similar health levels. Additionally, the positive control populations show better overall health compared to the tadpoles in the experimental runs; we attribute this puzzling fact to the high variability across batches.

We conducted *χ^2^* tests on the health-scored tadpole populations. For the aggregated data of experimental batches B1–7, the contingency Table SI2 yields a *χ^2^* statistic of 4.52 and a non-significant p value of 0.21. For the aggregated positive control runs +1–3, the contingency Table SI3 yields a *χ^2^* statistic of 0.85 with a non-significant p value of 0.84. Additionally, *χ^2^* tests for individual batches revealed no p values below 0.01. All these results strongly suggest that the control and hypomagnetic populations are not statistically different in terms of health scores.

We now raise two key points. First, we speculate whether accelerated development, despite our present observations, could in principle lead to an increased occurrence of abnormalities. While we do not observe this effect in our current data, accelerated growth might inherently increase the likelihood of developmental errors. Second, we consider the potential influence of having disrupted chemical signaling among tadpoles by raising them in isolation within separate wells. Our controlled experimental setup could have masked the emergence of developmental abnormalities whose cues are chemical in origin, or of toxic chemicals that increase mortality. To address this second point, we performed additional experiments with tadpoles raised communally in Petri dishes (see Supplementary Section SI8 below). Our findings, however, show that raising tadpoles together for three days did not significantly alter the rate of abnormalities either. Further studies might explore whether more extended periods of communal rearing or larger population sizes could reveal a stronger role for chemical signaling in tadpole development under hypomagnetic conditions.

**Table SI2:** Contingency table for experimental batches B1–7.

| health score vs. cond. | control | hypomagnetic |
| --- | --- | --- |
| 1 | 513 | 541 |
| 2 | 55 | 55 |
| 3 | 52 | 62 |
| 4 | 268 | 230 |

**Table SI3:** Contingency table for positive control runs +1–3.

**Table SI2:** Contingency table for experimental batches B1–7.
| health score vs. cond. | control | hypomagnetic |
| --- | --- | --- |
| 1 | 687 | 698 |
| 2 | 35 | 34 |
| 3 | 24 | 19 |
| 4 | 50 | 46 |

**Fig. SI2:**
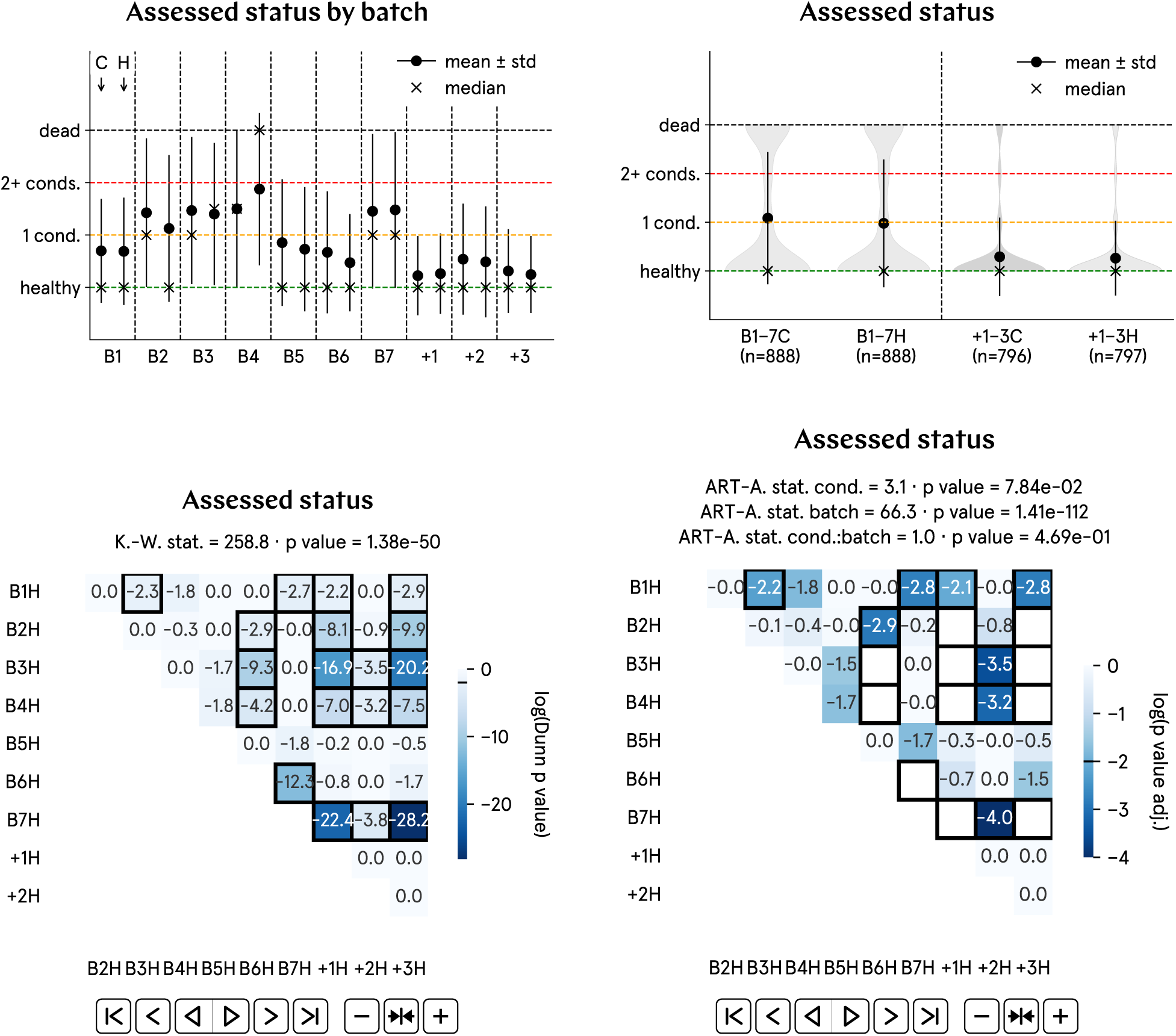
Magnetic field conditions do not significantly alter the health of the tadpoles. The human-made health scoring reveals that control and hypomagnetic tadpole populations exhibit remarkably similar health levels. A score of 1 is given for healthy tadpoles; of 2 for tadpoles with one morphological defect or adverse condition; of 3 for those with multiple defects or adverse conditions; and of 4 for tadpoles deceased by day 1 p.f.

### SI8. Control: The effect of magnetic field conditions on the mortality rates of communally raised tadpoles is not significant

We opted to raise the tadpoles individually in separate wells to mitigate the higher mortality rate commonly observed in tadpoles raised communally. In shared environments, contaminants produced by sick tadpoles or decomposing debris from dead tadpoles can spread. By isolating each tadpole, we aimed to prevent cross-contamination and ensure more controlled, consistent experimental conditions, which we believe were crucial for obtaining reliable data to support our conclusions.

For batches B5 and B6, in addition to the tadpoles in the well plates, we also raised tadpoles communally in Petri dishes. The images for these Petri dishes can be found in GitHub folder 10 Posters, with the label ‘PD’.

Statistical tests comparing mortality rates between the two conditions in Petri dishes revealed no significant differences. For instance, in batch B5, both the log-rank test [17] and the Cox proportional hazards model analysis [17] showed no statistically significant differences between the survival curves of the control and hypomagnetic groups, with p values of 0.81 and 0.84, respectively. Similar results were obtained when the health score described in Supplementary Section SI7 was applied to the Petri dish populations.

Based on our findings, we conclude that raising tadpoles communally does not increase the mortality rate in tadpoles exposed to hypomagnetic conditions. The higher abnormality rates reported in previous studies for amphibians raised without Earth’s geomagnetic field [6, 26, 51, 52] thus remain unexplained in light of our data.

### SI9. Description: hypomagnetic chamber

The hypomagnetic chamber was custom-designed by one of the authors (P. F.) and manufactured by Magnetic Shields Ltd. It was originally delivered with two layers of mu-metal, providing shielding against low-frequency fields (*<* 100 Hz) but being less effective at blocking high-frequency electromagnetic radiation; we refer to this configuration as having ‘DC shielding’, which is depicted in Supplementary Fig. SI3. Data for batches B1–5 were obtained using this configuration. Between batches B5 and B6, an additional copper layer was permanently installed to also shield against high-frequency electromagnetic radiation; we refer to this configuration of the hypomagnetic chamber as having ‘AC+DC shielding’.

The residual magnetic field inside the hypomagnetic chamber was measured with the tri-axial optically-pumped magnetometer QuSpin QZFM. This magnetometer has an intrinsic offset from the absolute zero that can slowly drift over a period of weeks (we measured it to shift by about 500 pT over the course of 6 weeks). For this reason, since our goal was to get as close as possible to zero residual field, the magnetometer offset was measured every two weeks and accounted for in the magnetic field calibration. The offset was calculated by performing a field measurement several times (usually 8 to 12) in the center of the hypomagnetic chamber, each time rotating the magnetometer by 180°. In this way, taking the x axis as an example and calling *B*_x_ the actual residual magnetic field, we should be measuring *B*_x_ and -*B*_x_ in an alternate fashion. Because of the intrinsic offset, the actual measurement will have a component *B*_offset x_ that does not depend on the direction the sensor is facing. This makes the value read by the magnetometer *B*_meas x+_ = *B*_x_ + *B*_offset x_ and *B*_meas x_- = *−B*_x_ + *B*_offset x_ when measuring along +x and -x, respectively. After the field value was measured a few times, the offset was computed using the average values of *B*_meas x+_ and *B*_meas x-_; assuming the true value *B*_x_ does not change, the offset is thus estimated as *B*_offset x_ = (*B*_meas x+ avg_ + *B*_meas x- avg_)/2. Once the offset was found, the final estimation of the magnetic field value was computed using *B*_x_ = *B*_meas x+_ *− B*_offset x_. The same procedure was applied to find the offset for the other axes. In our case, the offsets were estimated to be *B*_offset x_ = *−* 1492 pT, *B*_offset y_ = 257 pT and *B*_offset z_ = 3949 pT. In this work, whenever we mention a residual magnetic field *B*, it is calculated using the formula 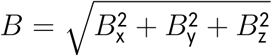, thus accounting for the intrinsic offset of the magnetometer.

To minimize the residual magnetic field, degaussing should in principle be performed each time the hypomagnetic chamber is open and closed. This process involves applying a large, low-frequency AC current to a cable that encircles the chamber’s edges. As the current is gradually reduced to zero, small, randomly oriented magnetic domains form within the mu-metal layers, ensuring a minimal net residual magnetic field inside the chamber. Using this method, residual fields as low as 4 nT have been measured at the center of the chamber. However, in our experiment, we chose not to degauss the chamber after each opening to avoid exposing the tadpoles to the strong AC field generated during the procedure. In addition, as shown in Supplementary Fig. SI4, the residual magnetic field changes significantly after the hypomagnetic chamber is opened following degaussing; opening and closing the chamber door is necessary whenever tadpoles are moved in or out for imaging. This occurs mostly because the inner mu-metal layer becomes partially magnetized by exposure to Earth’s magnetic field. However, we observed that, after 4 to 5 openings following degaussing, the magnetic field stabilized and no longer changed significantly. Therefore, before each experimental run, we performed degaussing, followed by several door openings and closings, before conducting the final field calibration.

The residual field inside of the hypomagnetic chamber was calibrated over a volume of 16’ *×* 16’ *×* 16’, as represented schematically in Supplementary Fig. SI5 a). The resulting field map is depicted in b) ; the 3D interactive version of this plot is available as [an HTML file](https://github.com/Quantum-Biology-Institute/Weak-magnetic-field-effects-in-biology-are-measurable/blob/main/11%20Calibration%20data/Field%20uniformity/calibration_2024_05_16.html) in the GitHub repository for this work (see Supplementary Sections SI69 and SI75). The measurement was taken after opening the door of the hypomagnetic chamber several times after performing degaussing. The calibration shows that, while the field is not very uniform over the large volume inspected, the area towards the center and back of the wooden shelf is very uniform over the size required for a few plates to be placed. For this reason, we decided to position the plates always close to the center of the wooden shelf and slightly towards the back.

As a last step in the reduction of the residual magnetic field, an active compensation system was designed and built by one of the authors (A. L.). As shown in Supplementary Fig. SI3, the compensation for the y axis consists of four single-turn coils that travel the perimeter of the chamber (taken as a section of the xz plane); this coil system was installed by Magnetic Shields Ltd. Compensation along the x and z axes was achieved by building the coils depicted in panel a) of Fig. 1. These coils have a Helmholtz geometry [53], ensuring good uniformity of the produced field; their support was constructed entirely out of plastic, including the screws, to ensure no stray magnetic field was introduced inside the hypomagnetic chamber. Each coil support in this system holds both one single-turn coil used for active compensation of the residual nT-level magnetic field, and one 10-turn coil used to replicate Earth’s magnetic field inside the hypomagnetic chamber for the positive control runs (see Supplementary Section SI11).

Active compensation was performed using a QuSpin Low Noise Coil Driver and the single-turn coil set. With this setup, it is possible to control the magnetic field value in each axis with a precision of *<* 0.1 nT. However, due to the slight fluctuations in magnetic field inside the hypomagnetic chamber caused by the opening and closing of the door, and the gradual drift in the magnetometer intrinsic offset, we can only confidently state that the residual magnetic field remained below 1 nT throughout the experiment.

**Fig. SI3:**
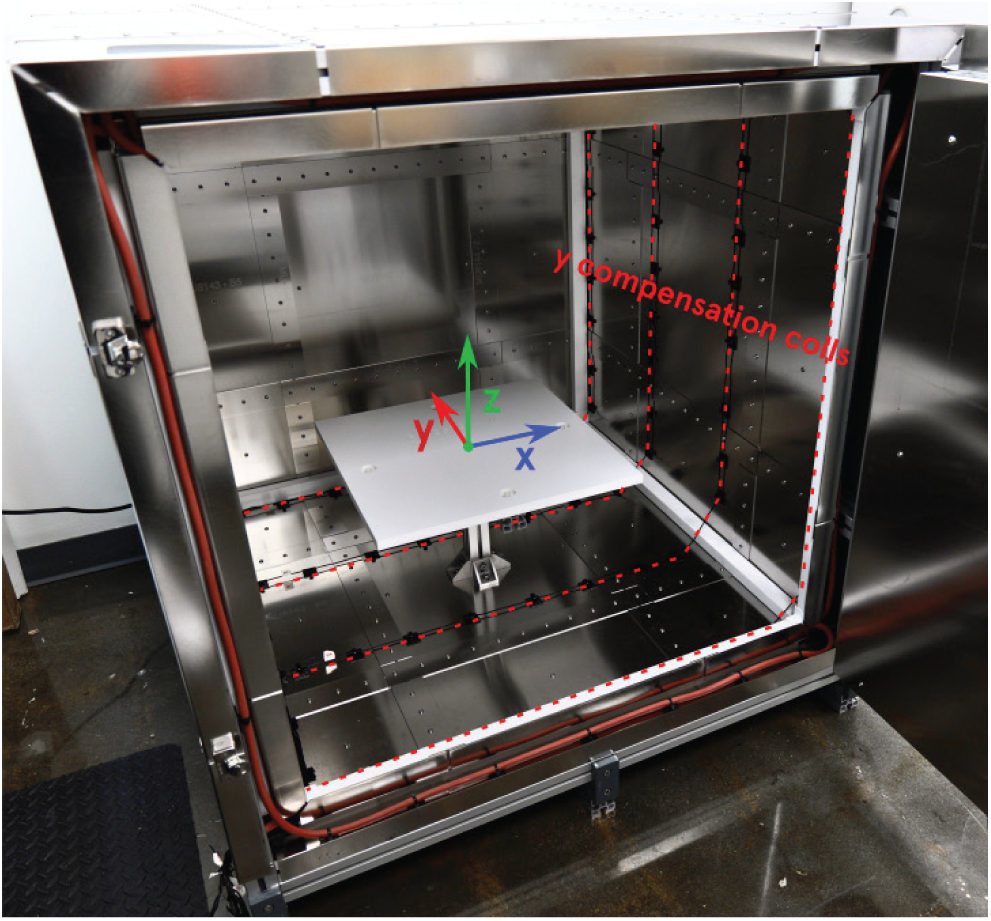
Hypomagnetic chamber with DC shielding only. This configuration has two layers of mu-metal. Also shown are y coils for magnetic field compensation, and the white wooden shelf where the samples are placed.

**Fig. SI4:**
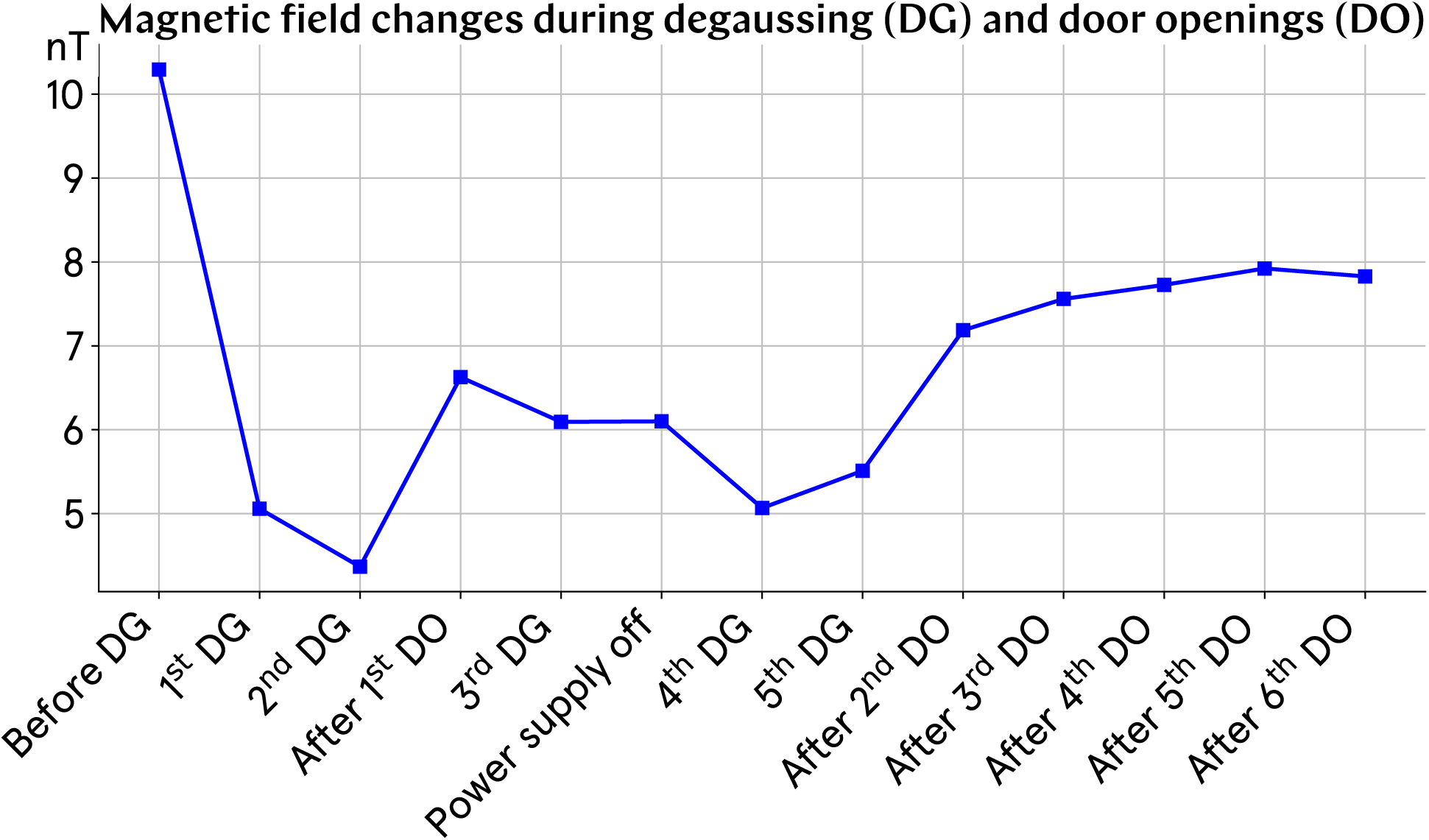
Degaussing and door opening influence on the residual magnetic field inside the hypomagnetic chamber. The field was measured at the center of the wooden shelf.

**Fig. SI5:**
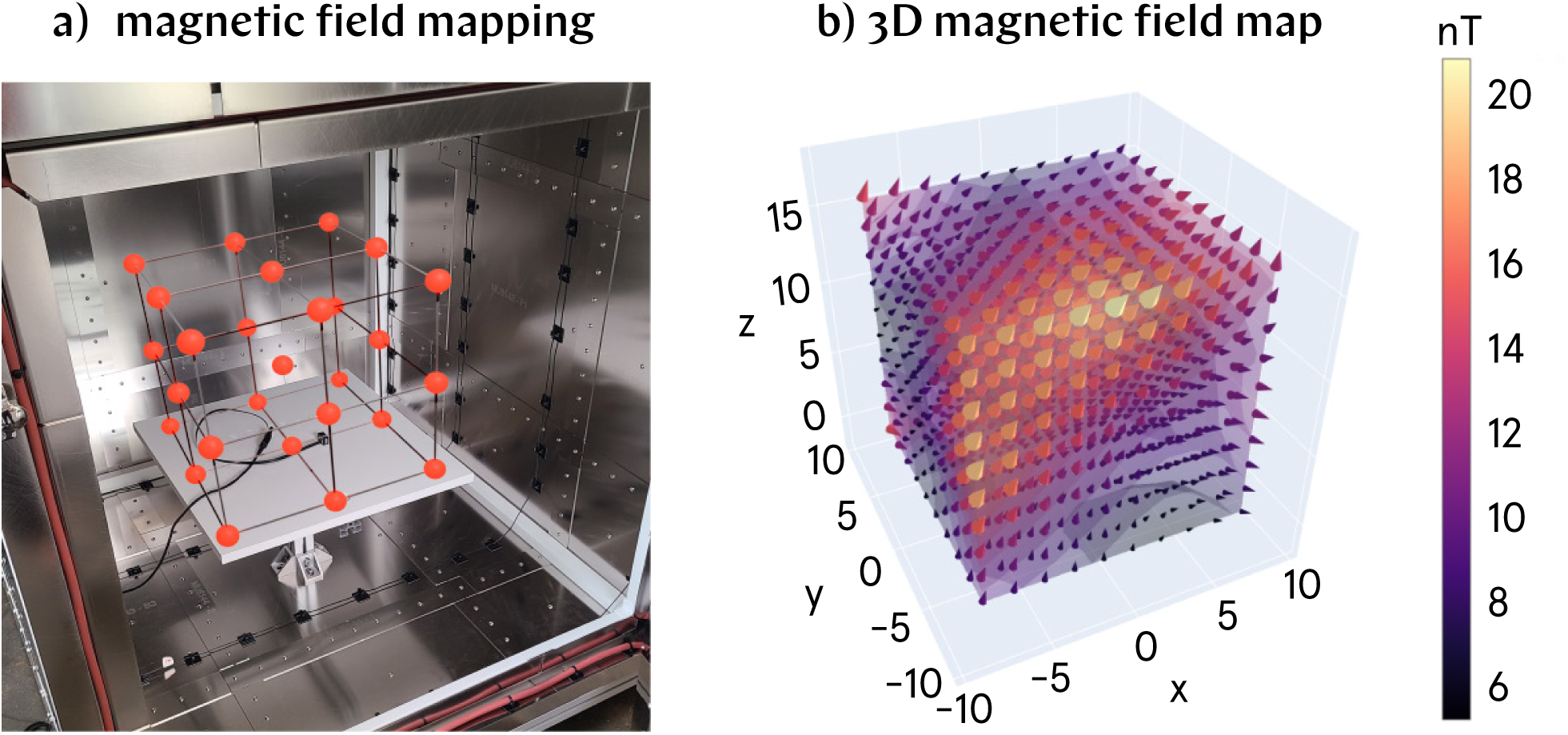
Residual field mapping and calibration. a) The magnetic field was mapped over a volume of 16’ 16’ 16’ and the residual field was measured in each of the points represented. b) The mapped field shows the field uniformity and can be inspected in the 3D interactive HTML plot available in the GitHub repository (see text for details). The photo in a) is taken with the same perspective as the plot in b).

### SI10. Description: control box

We chose to use an inactive (*i.e.*, switched off, but not unplugged) cell incubator (Forma Scientific CO_2_ Water Jacketed Incubator) already available in the lab as the control environment for this experiment, as shown in Supplementary Fig. SI6, which includes a sample placement overview. The internal glass door is also closed when the control samples are inside. This ‘control box’ exhibits similar environmental conditions to those of the hypomagnetic chamber, as confirmed by the data presented in Supplementary Fig. SI7.

Earth’s weak DC magnetic field, along with any ambient AC magnetic fields, are not blocked by the control box. While the control box is switched off, it is positioned against a wall with electrical connections and is approximately 1 m from the nearest switched on electronic device. We find it unlikely that measurable stray AC magnetic fields were generated within the control box due to the minimal AC current drawn from the wall for the following reason: Our most sensitive magnetometer, the QuSpin QZFM, with a bandwidth that includes 60 Hz, detected no differences at the nT level during calibration of the hypomagnetic chamber when current sources were alternately plugged in to the wall or unplugged while switched off. This strongly suggests that the control box being plugged in did not introduce any detectable AC magnetic fields.

The DC magnetic field environment inside the control box was measured by a Lakeshore F71 Teslameter to be 30.2 *µ*T in magnitude (with in-plane components of 12.2 and 7.7 *µ*T, and an axial component of 26.6 *µ*T).

We do not believe that there were measurable stray AC fields from the fact that the control box was plugged to the wall for the following reason. Our most sensitive magnetometer, the QuSpin QZFM, has a bandwidth including 60 Hz; during calibration of the hypomagnetic chamber magnetic field conditions, the QuSpin QZFM could not detect differences at the nT level when our current sources were plugged/unplugged from the wall.

**Fig. SI6:**
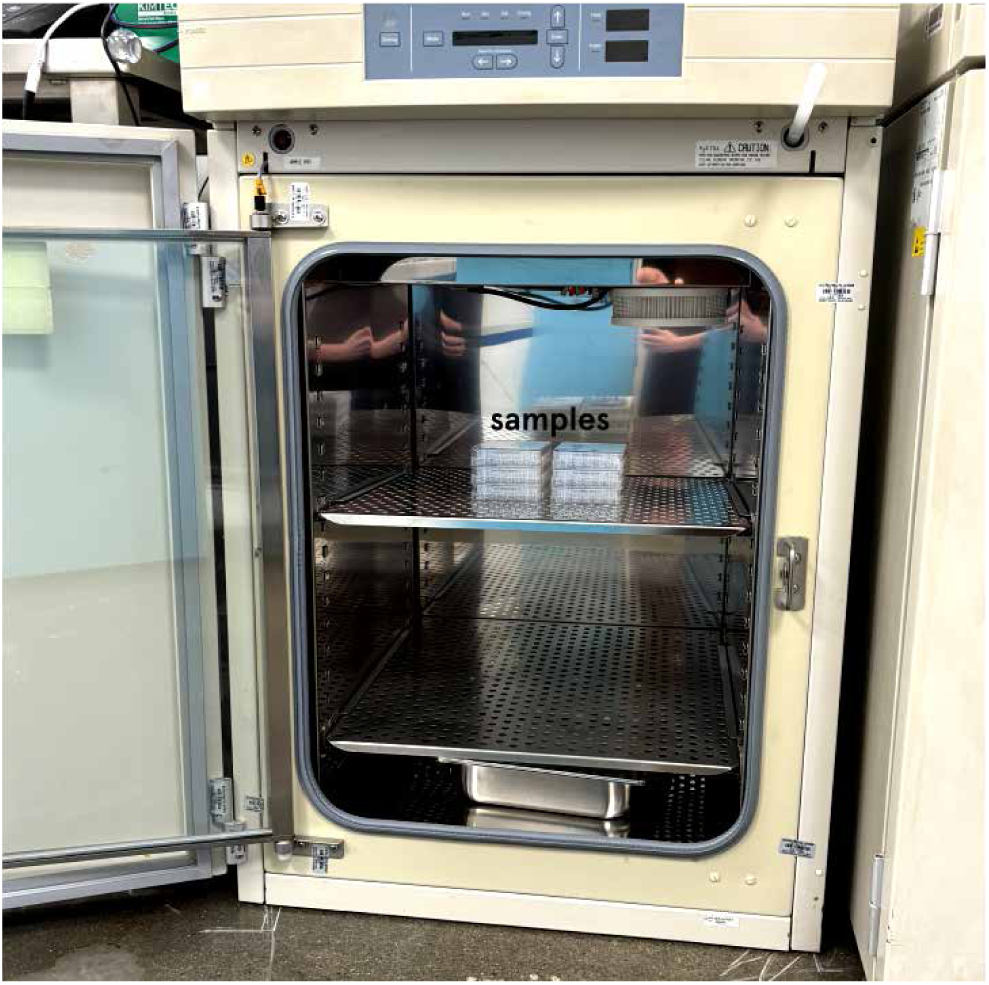
The control box is an incubator that is switched off, and that has a similar environment to the hypomagnetic chamber. Also shown is the metallic shelf where the samples are placed.

### SI11. Positive control run procedure

In the positive control runs, the magnetic field of the control box (detailed in Supplementary Section SI10) was replicated inside the hypomagnetic chamber, maintaining the same orientation relative to the plane in which the plates containing the tadpole embryos were positioned. This ensures that any potential effects related to the direction of the magnetic field are eliminated.

Two Jesverty SPS-3010 DC power supplies were used to drive current into a two-dimensional, 10-turn Helmholtz coil system. The current intensities used for reproducing Earth’s magnetic field inside the hypomagnetic chamber were 0.62 A through the x coils, and 1.28 A through the z coils shown in Fig. 1 a), for a total power consumption of 2.5 W. The dissipated power is responsible for the slightly higher temperature measured in the hypomagnetic chamber during the positive control runs, as discussed below in Supplementary Section SI12.

### SI12. Comparison: Environmental conditions in the hypomagnetic chamber and the control box are similar

Supplementary Fig. SI7 presents a comparison of humidity and pressure conditions between the hypomagnetic chamber and the control box, alongside the temperature curves from the main text. The curves were recorded for the hypomagnetic chamber with only DC shielding (shown in a) panels) and with both DC and AC shielding (shown in b) panels). As part of the positive control run procedure (described in Supplementary Section SI11), we also calibrated humidity and pressure for these runs (shown in c) panels).

The measurements shown in Supplementary Fig. SI7 were acquired simultaneously in both the control box and the hypomagnetic chamber using a pair of Element-A sensors from Elemental Machines. Placing sensors inside the environments during experimental runs would have interfered with the magnetic field conditions. As a result, all calibrations for temperature, humidity, pressure, and light were performed separately from the experiments, with careful replication of the experimental conditions. These calibration curves were recorded after the sensors had been inside their respective environments for over 24 hours with the doors closed.

#### SI12.1. Temperature

It can take up to a full day for the temperature to stabilize after the doors have been closed; in other words, starting with the control box and the hypomagnetic chamber initially in thermal equilibrium with the lab, whose temperature is measured at approximately 20.15°C when the air conditioning is set to 70°F, it can take up to a full day for the temperatures to reach the equilibrium conditions shown in Supplementary Fig. SI7. To minimize the temperature difference between the environments, both the hypomagnetic chamber and the control incubator were left open for at least a day before each experimental run began. Additionally, the doors of both environments were opened multiple times daily to remove the plates for measurement. Considering these factors, the actual temperature difference during the experiment was likely less pronounced than what is shown in the calibration curves.

In the first five batches of the experiment (B1–5), with only DC shielding, the temperature in the hypomagnetic chamber was measured to be 0.30–0.50°C lower than in the control box. This particular calibration was performed when the lab air conditioning was off; this explains why the control box’s temperature, at 20.6°C, is slightly higher than in the other calibration curves, at 20.1°C, which were taken with the lab air conditioning on and set to 70°F. The slightly higher temperature in the control box reinforces our argument that accelerated development occurs in the hypomagnetic chamber, even when its temperature is slightly lower than that of the control box. This calibration was cut short due to sensor malfunction, lasting only 6 hours, and we were unable to conduct a more thorough calibration before the permanent installation of the AC shielding. After the copper shell was added, the temperature in the hypomagnetic chamber was found to be slightly higher than in the control box by about 0.10–0.15°C. When simulating Earth’s magnetic field inside the hypomagnetic chamber for the positive control runs, the heat generated by the coils further increased the temperature difference to approximately 0.45°C. This aggregate temperature trend could not have influenced our results, which demonstrate faster development in the hypomagnetic populations with both DC and AC+DC shielding, and no significant difference in the positive control runs. This further supports the hypothesis that the absence of Earth’s magnetic field accelerates *Xenopus laevis* embryo development.

A simplistic Q_10_ model is sometimes used to provide a rough estimate on how embryogenesis changes with temperature; the Q_10_ coefficient describes how the rate of a physiological process is altered upon a 10°C increase in temperature:

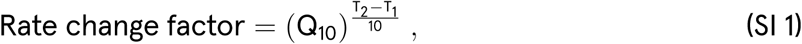

with the temperatures T_1_ and T_2_ in oC. For amphibians, Q_10_ is typically between 2 and 3 [13]. A Q_10_ of 2 (respectively, of 3) means the development rate doubles (triples) with a 10°C increase. Our temperature differential is at most approximately 0.50°C; for a Q_10_ of 2 (3), this translates into approximately 50 (144) minutes of accelerated development per day that could be attributable to the temperature difference between the hypomagnetic chamber and the control box. However, we believe that a development speed-up of one to two hours per day cannot fully explain the significant developmental changes we have observed.

Unfortunately, there is no published data that extends beyond the first 24 hours p.f. [13] to provide a comparison of developmental stages as a function of time during *Xenopus* embryogenesis; the existing tables are for up to 22 hours p.f. at 18°C and up to 24 hours at 22°C. When assessed for development, our images taken at 24 hours p.f., for tadpoles raised at the nominal lab temperature of approximately 20.15°C, fall within the expected range of NF stages 18 to 24 [15].

#### SI12.2. Humidity

Humidity influences the moisture content of the surrounding air, which in turn affects water balance and gas exchange in embryos. For batches B1 through B5, the humidity in the control box and in the hypomagnetic chamber differs by only approximately 1%, which is negligible, with the hypomagnetic chamber being slightly more humid. For batches B6 and B7, however, humidity is approximately 7% higher in the control box, meaning that evaporation inside the hypomagnetic chamber, at 60% RH, is nominally about 18% higher than in the control box, at 67% RH (evaporation rate *E*_RH_ ∝ (100 - RH); relative evaporation increase 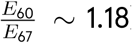). Condensation on the inner surfaces of the lids was rarely observed when the plates were removed for imaging from either the hypomagnetic chamber or the control box. While the plastic lids on the plates reduce overall evaporation, the relative difference between the two humidity conditions should still persist. Lower humidity can lead to faster heat loss due to evaporation, which acts as a cooling process by absorbing heat and could effectively slow developmental processes by lowering the temperature in the embryos’ immediate environment. This cooling effect would be expected to slow the development of the tadpoles in the hypomagnetic chamber. Contrary to this expectation, we observe accelerated development in that population. Overall, given the small humidity difference in most batches, we expect these effects to be minimal, with the larger humidity difference in B6 and B7 potentially having a more noticeable impact on development. During positive control runs, humidity is 4% higher in the control box (equivalently, the evaporation is nominally 12% higher in the hypomagnetic chamber).

To thoroughly test the effects of humidity, we prepared two 48-well plates with 0.75 mL of 0.1x MMR following the same procedure used for the experiments. The plates were left undisturbed with their lids on, one in the control box and one in the hypomagnetic chamber (AC+DC shielding configuration, with active compensation) for three days; the lab air conditioning was on and set to 70°F. The initial and final weights were compared to assess the impact of humidity on evaporation. The initial weights of the plates, before inserting them in the different environments, were *m*_H,i_ = 104.646 g and *m*_C,i_ = 105.064 g in the plate going to the hypomagnetic chamber and to the control box, respectively. The final weights after the plates were left untouched for three days were *m*_H,f_ = 104.216 g and *m*_C,f_ = 104.636 g. By taking into account the initial weight of the empty plates, which were *m*_H,p_ = 69.820 g and *m*_C,p_ = 69.835 g, we can estimate the total and relative evaporated solution. For the plate in the hypomagnetic chamber, the total evaporated volume is *m*_H,evap_ = *m*_H,i_ *− m*_H,f_ = 0.430 g, while the relative evaporated volume is *m*_H,evap%_ = 100 *· m*_H,evap_/(*m*_H,i_ *− m*_H,p_) = 1.23%. As a comparison, the total evaporated volume for the plate in the control condition is *m*_C,evap_ = *m*_C,i_ *− m*_C,f_ = 0.428 g, while the relative evaporated volume is *m*_C,evap%_ = 100 *· m*_C,evap_/(*m*_C,i_ *− m*_C,p_) = 1.21%. Thus, the measured percentage difference in evaporation (calculated as (*m*_H,evap_ *−m*_C,evap_)/*m*_C,evap_ *∼* 0.4%) is two orders of magnitude smaller than the expected nominal evaporation difference (18%). Similar to our argument for temperature, this further supports the claim that the actual humidity differences during the experiment were likely less significant than indicated by the calibration curves.

For completeness, we also monitored pH changes. The 0.1x MMR solution used to fill the plates in the weight test above had an initial pH of 7.27. After three days in the plates within the different environments, the pH was measured as pH_H_ = 7.02 and pH_C_ = 7.01—once again, a negligible difference.

Considering the minimal differences in evaporated mass and pH between the plates, it is highly unlikely that variations in humidity had any significant impact on our biological observations.

#### SI12.3. Pressure

Pressure influences the partial pressure of gases such as oxygen and carbon dioxide, which in turn affects gas exchange in embryos. The pressure difference between the control box and the hypomagnetic chamber is approximately 1 hPa, relative to a baseline of around 1000 hPa, with the hypomagnetic chamber slightly higher. However, normal atmospheric pressure fluctuations (*e.g.*, due to weather changes), which we observe in our 24-hour data, often exceed this difference. Therefore, a 1 hPa variation is unlikely to significantly affect gas exchange, fluid dynamics, or other key developmental processes in our water-dwelling embryos.

As the tadpoles are immersed in liquid and their plates are covered with lids, the slight variations in pressure reported above are unlikely to have any significant impact.

#### SI12.4. Light intensity and spectrum

Although both the hypomagnetic chamber and the control box have small openings, all 24-hour light measurements taken with Element-A sensors (conducted concurrently with the measurements shown in Supplementary Fig. SI7) consistently registered 0 lux, indicating that light levels were below the sensors’ nominal detection threshold of 0.01 lux. We then used two identical Adafruit TSL2591 High Dynamic Range Digital Light Sensors (nominal detection threshold of 188 *µ*lux) to measure the illuminance depicted in panels a) through c) of Supplementary Fig. SI8; to mitigate potential bias from one of the sensors, we alternated the sensors between the two environments and averaged the measurements. The control box has an average measured illuminance of 0.0873 *±* 0.0003 lux, and the hypomagnetic chamber an average measured illuminance of 0.0751 *±* 0.0002 lux; it is unclear why these values were not detected by the Element-A sensors. We also note that the Adafruit sensor readings remained nearly constant regardless of whether it is day or night, or whether the lab lights are on or off.

For comparison, the illuminance from a full moon on a clear night is around 0.1–0.3 lux, whereas starlight on a clear, moonless night provides about 0.0001 lux.

To compare the spectral characteristics of the light in the hypomagnetic chamber and the control box, we used the UV-sensitive Ocean Optics ST-UV-50 spectrometer, which covers a range of 185–650 nm and offers a resolution of 3.7 nm (full width at half maximum). Each spectrum was generated by averaging 100 measurements, each with a maximum integration time of 6 seconds. A dark spectrum was recorded with the spectrometer’s cap on for reference. Since spectrometers are highly directional devices, it is noteworthy that the spectra taken inside the control box were highly consistent, irrespective of the direction in which the spectrometer was pointed. In contrast, the hypomagnetic chamber, which has a 1’ hole vertically centered and penetrating all shielding layers, presented a challenge. When the spectrometer was placed centrally inside the chamber and pointed upwards toward the hole, an exceptionally strong light signal was recorded—likely unrepresentative of the actual light conditions within. To address this, we obtained two spectra by pointing the spectrometer in different directions (towards the front and back of the closed hypomagnetic chamber) and averaged the results to provide a more accurate depiction of the light environment. Both the control box and hypomagnetic chamber spectra were corrected by subtracting the dark spectrum. These results are shown in panel d) of Supplementary Fig. SI8. These spectra were recorded on the same day between 2 and 4 pm. Since our illuminance sensors showed no variation based on the time of day, the timing of these spectral measurements is likely irrelevant. The occasional negative values in the spectra are due to the dark spectrum recording higher values than the bright spectrum, likely indicating minimal activity at that wavelength and spectrometer orientation. The spectra are critical in illustrating that there is no such a thing as complete darkness, which is important given that one hypothesis to explain our findings involves the photoexcitation of chemical compounds. Many of these compounds are excited by near-UV photons, which are captured in the spectra. This is discussed further in Supplementary Section SI77.

Pigmentation in *Xenopus* is well-documented to respond to the background color of its environment, with most studies focusing on the regulation of eumelanin (dark) pigmentation [24]. However, there is emerging evidence suggesting that other pigment cells, such as the carotenoid- and pteridine-containing xanthophores, may also play a role in this adaptive response [54]. At least in some frogs, lightening takes up to six times as long as darkening [25]. The plates inside the hypomagnetic chamber were placed against a white wooden background, while those in the control box rested on a metallic grey surface. However, we believe this difference is negligible due to the very low light levels in both environments. Imaging was performed under ambient light and on a white surface; the imaging time per plate is under 10 minutes.

#### SI12.5. Air volume

For completeness, the inner dimensions of the control box are roughly 48 cm *×* 50 cm *×* 67 cm (width *×* depth *×* height), for an air volume of approximately 0.16 m3. The inner dimensions of the hypomagnetic chamber with only DC shielding were 100 cm *×* 100 cm *×* 100 cm, for an air volume of approximately 1.00 m3. After installation of the AC shielding, the dimensions are 88 cm *×* 88 cm *×* 88 cm, for an air volume of approximately 0.68 m3. However, the difference in air volume is inconsequential, as the tadpoles are housed in small water-filled plates covered with plastic lids, isolating them from the surrounding air environment.

**Fig. SI7:**
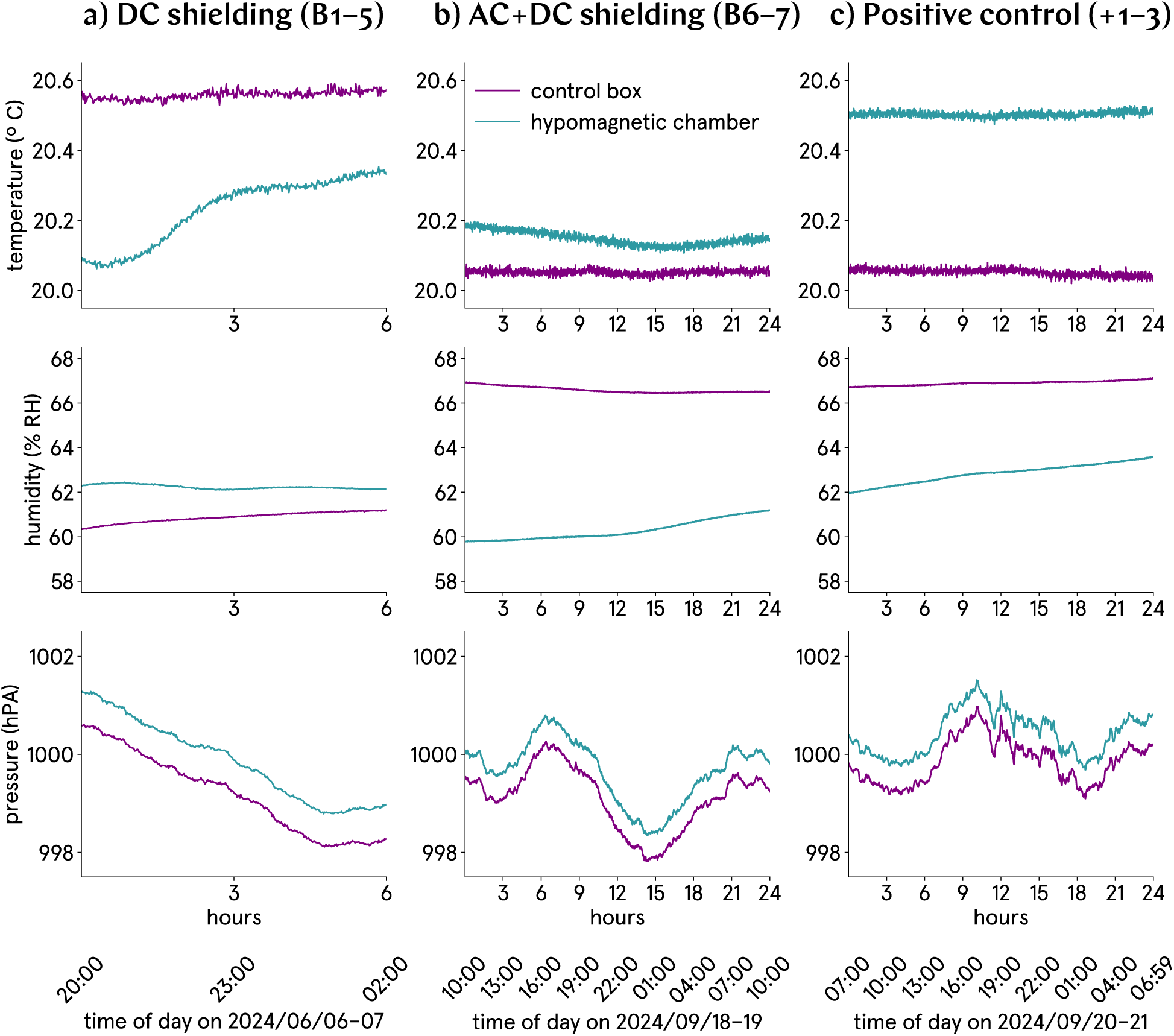
The temperature, humidity, and pressure levels are comparable between the hypomagnetic chamber and the control box. a) The DC shielding configuration of the hypomagnetic chamber was used for batches B1 through B5. a) The AC+DC shielding configuration of the hypomagnetic chamber was used for batches B6 through B7. c) The positive control runs +1 through +3 involved applying an artificial magnetic field inside the hypomagnetic chamber.

**Fig. SI8:**
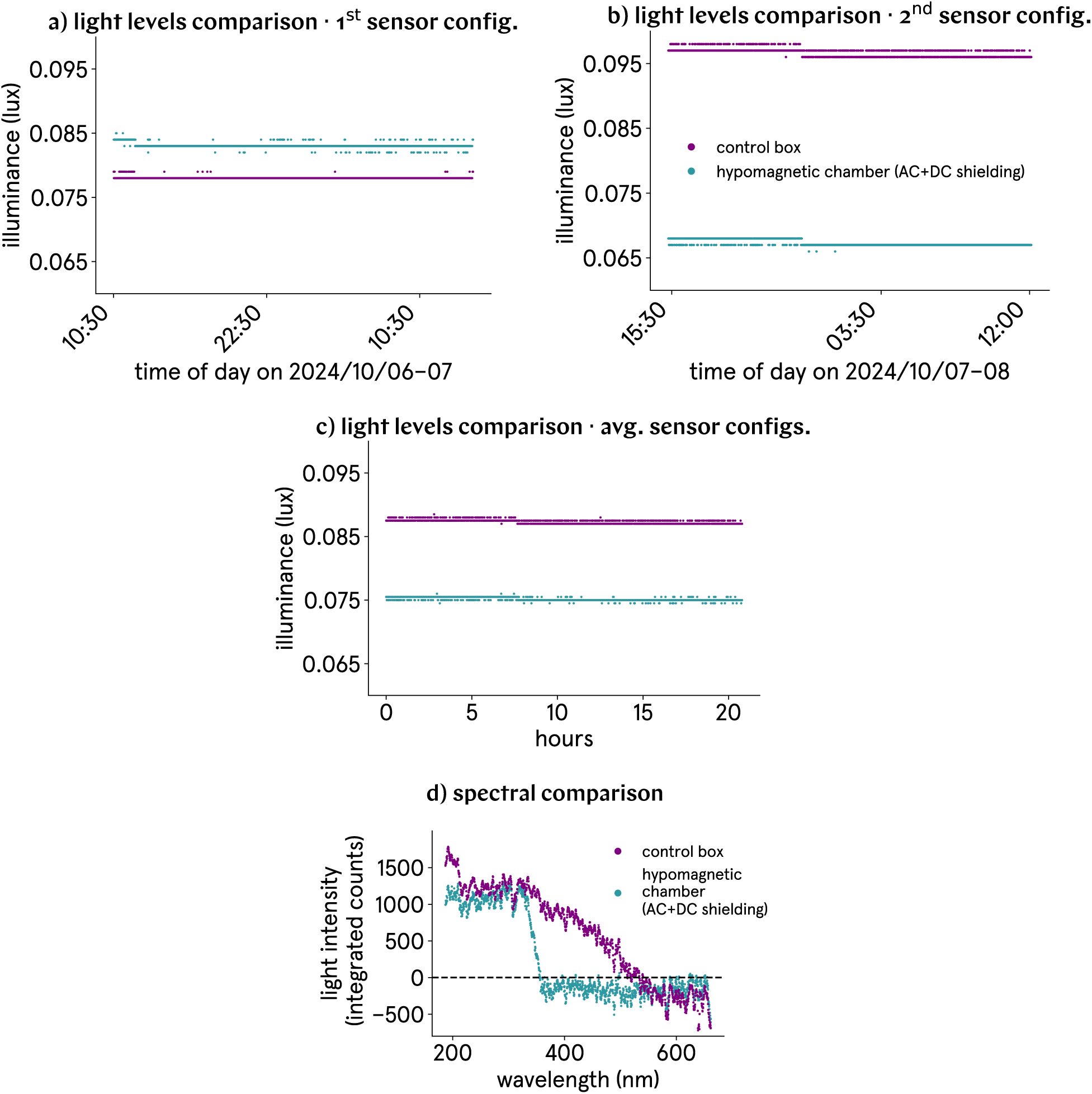
Light levels and spectra are comparable between the hypomagnetic chamber and the control box. a) and b) Light level measurements were simultaneously taken in both the hypomagnetic chamber and the control box using two identical sensors. To account for potential differences between the sensors, calibration curves were recorded over a period of approximately 24 hours, alternating the sensors between the two environments. c) The average illuminance for both environments is shown, based on the averaged recordings from each sensor. d) A spectral comparison between the environments reveals the presence of photons even in the UV range.

### SI13. Image segmentation

Raw.jpg images for the three days are automatically batch-segmented using Adobe Photoshop’s AI-assisted Select Object tool with default settings, followed by Select and Mask. This process isolates the object of interest against a transparent background; we save thus segmented images as.png. Photoshop’s tools outperformed both custom python code and ImageJ [55] plugins. The segmented images were then manually inspected; in cases where glare or unwanted objects were mistakenly chosen by the algorithm, they were cropped from the image and the Select Object action was repeated.

In a few instances where the tadpole eye was missed by the algorithm, it was manually added to the selection mask. For exceptional images containing multiple embryos, the leftmost one was consistently chosen.

In rare cases where the procedure failed repeatedly, images were manually segmented. The following is an exhaustive list of such images.

Day 1 images that required segmentation by hand (9 out of 2764 images):

1. B1CD1P3-13, glare
2. B1CD1P4-12, shadow
3. B1HD1P1-20, glare
4. B1HD1P2-23, glare
5. B2CD1P3-11, shadow
6. B2CD1P4-06, shadow
7. B2HD1P1-06, remove bubble
8. B2HD1P5-08, differentiate from debris
9. B5CD1P2-21, differentiate from debris

Day 2 images that required segmentation by hand (8 out of 2771 images):

1. B3CD2P4-14, shadow
2. B6HD2P1-24, glare
3. B6HD2P1-35, glare
4. B6HD2P2-24, glare
5. B6HD2P2-47, out of focus
6. B6HD2P5-44, out of focus
7. B7CD2P3-32, out of focus
8. B7CD2P4-14, out of focus

Day 3 images that required manual segmentation to accurately reflect their length (7 out of 2744 images):

1. B3CD3P1-10, tadpole color too light for contrast
2. B3CD3P5-03, tadpole color too light for contrast
3. B3HD3P2-09, glare
4. B3HD3P2-11, tadpole color too light for contrast
5. [B3HD2P2-12](https://github.com/Quantum-Biology-Institute/Weak-magnetic-field-effects-in-biology-are-measurable/blob/main/9%20D3%20images/B3/B3H/B3HD3/B3HD3P2/B3HD3P2-12.png), tadpole color too light for contrast
6. B3HD3P2-17, tadpole color too light for contrast
7. B3HD3P4-20, tadpole color too light for contrast

### SI14. Effect size analysis

Throughout the manuscript, we use four different measures of effect size; each assesses different aspects of the data. In the formulas for these measures, the following quantities appear:

n_C_ is the number of control tadpoles;
n_H_ is the number of hypomagnetic tadpoles;
{C_1_*, …,* C_nC_ } is the array of values of some property for control tadpoles;
{H_1_*, …,* H_nH_ } is the array of values of some property for hypomagnetic tadpoles;
*µ*_C_ is the mean value of some property for control tadpoles;
*µ*_H_ is the mean value of some property for hypomagnetic tadpoles;
*σ*_C_ is the standard deviation of some property for control tadpoles;
*σ*_H_ is the standard deviation of some property for hypomagnetic tadpoles;
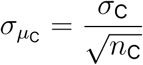 is the standard error of the mean of some property for control tadpoles; and
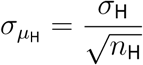 is the standard error of the mean of some property for hypomagnetic tadpoles.

The first measure is Cohen’s effect size *d* [56]:

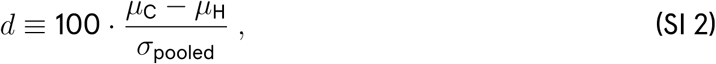

where the pooled standard deviation

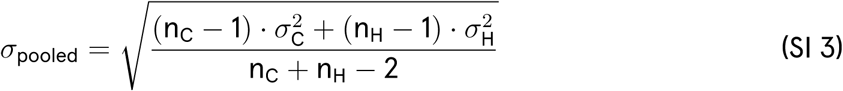

is a weighted average of the standard deviations of the two independent groups. Cohen’s formula expresses the difference between the means of the two groups as a percentage of the pooled standard deviation, which means it reflects how much the two distributions overlap, and which allows for a standardized comparison. Effect sizes are roughly classified as small (|d| ≳ 20%), medium (|d| ≳ 50%) and large (|d| ≳ 80%). If the variances of the control and hypomagnetic property under consideration are substantially different, Cohen’s *d* might not accurately reflect the magnitude of the effect, as it assumes equal variances between groups. When variability within groups is small and means are far apart, *d* will report a large effect size. However, even if the variance is low but the actual mean difference is small, Cohen’s *d* will still report a relatively large effect size due to the calculation based on pooled standard deviations.

The second measure is Cliff’s Δ [57] which, unlike Cohen’s *d*, does not assume a normal distribution of the data. It reflects the dominance of one group over the other. It is calculated as:

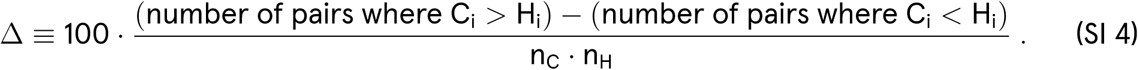

Effect sizes are roughly classified as small (|Δ| ≳ 15%), medium (|Δ| ≳ 33%) and large (|Δ| ≳ 47%).

The third measure *E*, also a percentage, was introduced by Binhi in the specific context of weak magnetic field effects in biology [1]:

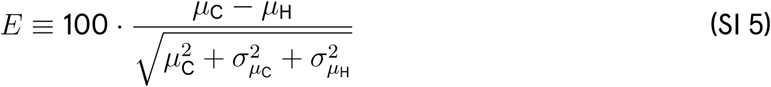

It considers not only the difference between the means but also the uncertainty in those means. Binhi’s *E* thus indicates how substantial the effect is relative to the combined variability and magnitude of the two groups. By incorporating the standard errors of the means, it accounts for the confidence in the observed differences. This measure is particularly useful when comparing groups with different sample sizes, as it adjusts for the reliability of the estimated means.

The fourth and final measure is the effect visibility *ν*, in percent:

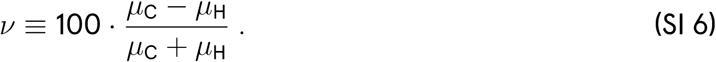

Effect visibility measures the relative difference between two means as a percentage of their sum. This measure is intuitive and akin to signal-to-noise, as it expresses the effect size in terms of how much one group deviates from the other relative to the total magnitude of the means. If the two means are close in value, or if the means are both large compared to their difference, the computed visibility will be smaller, regardless of the absolute difference. This measure is thus more conservative, especially when the groups are relatively similar in magnitude.

Both Cohen’s *d* and Cliff’s Δ emphasize the difference between the central tendencies of the groups and downplay variability in the data, which is why they can yield strong effect sizes when the groups are well separated. Differently from Cohen’s *d* and Cliff’s Δ, Binhi’s *E* and the visibility *ν* are more sensitive to the variability or relative similarity between groups; they are more direct comparisons of the mean difference relative to the combined magnitudes of the means or their variability. These are less influenced by the variability within each group and more by the relative difference in means. Taken together, these four measures reflect different aspects of our data.

### SI15. Effect change analysis

To compare the progression of morphological properties from day 2 to day 3, we use several measures. For each condition (control or hypomagnetic), we define:

D2 = {D2_1_*, …,* D2_n_} is the array of values of some property for day 2 images;
D3 = {D3_1_*, …,* D3_n_} is the array of values of the same property for day 3 images;
*µ*_D2_ is the mean value of the property on day 2;
*µ*_D3_ is the mean value of the property on day 3;
*σ*_D2_ is the standard deviation of the property on day 2; and
*σ*_D3_ is the standard deviation of the property on day 3.

The first measure, used in the main text, is the Z-score difference. A Z-score represents how many standard deviations a data point is from the mean. It is calculated as:

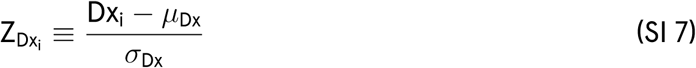

where x = *{*2, 3*}* represents day 2 or day 3, and i denotes the individual data point. The Z-score is unitless but should be understood as being expressed in units of standard deviation. It standardizes values by removing the mean and scaling by the standard deviation, allowing comparisons across different datasets or time points. To quantify effect change, we calculate the Z-scores for day 2 and day 3 datasets and then compute their difference:

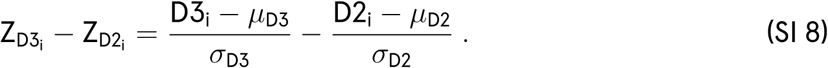

This comparison normalizes the differences between day 2 and day 3 by accounting for the variability in each dataset. A positive Z-score difference indicates that the day 3 value is further above the day 3 mean relative to the day 2 value’s distance from the day 2 mean, suggesting an increase in the measured property. The Z-score difference is also unitless and represents the relative change in terms of standard deviations.

The second measure is the effect change visibility *ν*, previously used to quantify effect size (Supplementary Section SI14), and defined as:

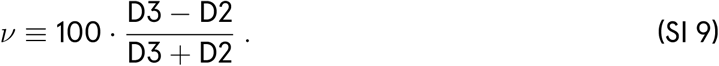

The third measure is the percentage change defined as:

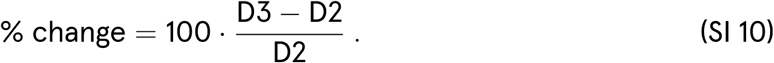

The fourth and final used measure is the relative difference in percent, which quantifies how different the two values are relative to their maximum:

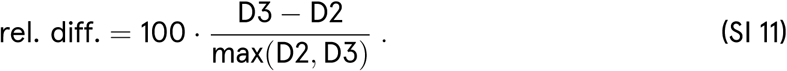

### SI16. Statistical analysis

#### SI16.1. Confidence interval calculation

Given a dataset *{*X_1_*, …,* X_n_*}* with sample size n, a mean *µ*, and a standard error of the mean *σ_µ_*, the confidence interval CI for the mean is calculated as:

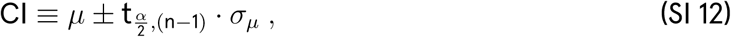

where 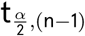 is the critical value from Student’s T-distribution with (n *−* 1) degrees of freedom, corresponding to the desired confidence level (1 *− α*) [17]. Throughout our manuscript, we use CI = 95 % (and hence *α* = 0.05).

Confidence intervals for the mean are used in plots of type c), described in Supplementary Section SI74.

#### SI16.2. Bootstrapping

Bootstrapping is a resampling technique. It involves repeatedly drawing samples with replacement from the original dataset distribution to create multiple ‘bootstrap samples’. A desired statistic (*e.g.*, median) can then be calculated for the samples. Bootstrapping helps estimate confidence intervals, standard errors, and other statistics when the underlying distribution is unknown or when sample sizes are small; it is non-parametric and doesn’t assume a specific data distribution.

Bootstrapping is utilized in plots of type d) (see Supplementary Section SI74), closely following the approach outlined in [58]. We employ 1000 bootstrap samples to estimate the 95% confidence interval and the distribution of the percentage difference between the medians of the control and hypomagnetic conditions. When the distribution of the observed properties is non-normal, as is most typical in our dataset, the median provides a more accurate representation of the central tendency of the sample compared to the mean.

#### SI16.3. Why we also report p values

In an effort to be as comprehensive as possible with our data analysis, we also report p values in several plots, most notably in plots of types e) and f) (for effect size) and g) through j) (for statistical tests), see Supplementary Section SI74.

When considered alongside the other evidence presented, we believe p values offer useful, complementary insight into whether the control and hypomagnetic populations are significantly different. However, it is important to note that p values are influenced not only by population differences but also by sample size: As sample size increases, precision improves (though this does not necessarily lead to greater accuracy). This contrasts with other metrics we report, such as effect size measures, which provide a more consistent assessment of population differences regardless of sample size.

#### SI16.4. T-test and Mann-Whitney test

In the bottom plot of plots of type e) and f), we report a p value that is obtained as follows.

A Kolmogorov-Smirnov normality test [17] checks if the control and hypomagnetic populations are normally distributed. If both populations have p values greater than 0.01 (suggesting normality), a T-test [17] is performed to compare the populations, and the obtained p value is reported. If at least one population fails the normality test, which is usually the case in our dataset, the Mann-Whitney test [17] is used instead, and its p value is reported. The Mann-Whitney test is a non-parametric method that compares medians when the data does not follow a normal distribution.

#### SI16.5. Kruskal-Wallis test and Dunn’s post-hoc test

The Kruskal-Wallis test [17] is a non-parametric statistical test used to compare three or more independent groups. In our case, we compare all batches within the same condition (control or hypomagnetic). Kruskal-Wallis is an alternative to the one-way analysis of variance (ANOVA) test [17] when the full data does not meet the assumption of normality, which is always the case in our dataset. The test ranks the data and assesses whether the median ranks across the groups are significantly different; it outputs a statistic (also known as ‘H-statistic’) and a p value. The H-statistic helps determine whether there are statistically significant differences between the medians of the groups; if it is large enough, it suggests that at least one of the groups is different from the others. The significance of the H-statistic is assessed by comparing it to a *χ^2^* distribution with (n *−* 1) degrees of freedom, where n is the number of groups; a significant p value suggests that at least one group’s median is different, but a post-hoc test is needed to pinpoint exactly which groups differ.

If a significant difference is found (*i.e.*, if the Kruskal Wallis p value is less than 0.01), we use Dunn’s post-hoc test [17] to determine where the differences lie between groups. The Dunn post-hoc test compares the ranks of data between all pairs of groups and adjusts for multiple comparisons using the Bonferroni correction. Notably, even though Dunn p values are reported pairwise, our python statistical routine analyzes the entire dataset for all batches for each condition, ensuring that the comparisons account for all available data.

The result of the Kruskal-Wallis test for the control and the hypomagnetic populations, in addition to a visual representation of the Dunn p value between pairs of batches, are found in plots of type g) and h).

#### SI16.6. ART-ANOVA test and Tukey’s post-hoc test

The Kruskal-Wallis H-statistic described above tests for differences between group distributions (specifically, ranks of the data); in contrast, the ART-ANOVA F-statistic examines the effects of factors on aligned and ranked data, including potential interaction effects.

ART-ANOVA stands for ‘Aligned Rank Transform-ANOVA’ [17], a non-parametric method used to analyze experiments when the normality assumption of traditional ANOVA is violated. It works by aligning the data based on the factors being tested, then applying a rank transformation before conducting an ANOVA. This approach allows for both main and interaction effects to be tested; here, we use ART-ANOVA tests to assess the extent to which both condition (control or hypomagnetic) *and* the individual batch have significant effects on the outcome of our experiments. A subset of the ART-ANOVA output are statistics (also known as ‘F-statistics’) and corresponding p values. The F-statistics represent different sources of variation in the data; in our data, the ‘condition’ F-statistic measures the effect of the primary factor (*i.e.*, control vs. hypomagnetic) on the outcome of the experiment; the ‘batch’ F-statistic assesses whether different batches have a significant effect on the results; and finally, the ‘interaction condition:batch’ F-statistic evaluates whether the effect of the condition depends on the specific batch, indicating an interaction between the two factors. To obtain the corresponding p values, the F-statistics are compared to the F-distribution, which depends on the number of groups and on sample size. The F-distribution table provides these critical values at various significance levels; we use the significance level of 0.01.

If one or more p values for the F-statistics is significant, a post-hoc test is used to identify which specific pairs of group means differ significantly. We employ Tukey’s post-hoc test [17], which pinpoints the specific groups with differences while controlling for the overall Type I error rate (the probability of finding a difference that does not exist). Notably, even though Tukey’s adjusted p values are reported pairwise, our python statistical routine analyzes the entire dataset for all batches and both conditions, ensuring that the comparisons account for all available data.

The results of the ART-ANOVA test for the control and the hypomagnetic populations, in addition to a visual representation of the adjusted p value between pairs of batches, are found in plots of type i) and j).

### SI17. Day 1 morphological measures: eccentricity, elongation, roundness, and solidity

At 1 day p.f., we observed frog embryos at a range of developmental stages, which we assessed to be from 18 to 24 [16]. To quantify our qualitative observation that frog embryos under hypomagnetic conditions are developmentally ahead of control frog embryos already in day 1 p.f., we turn to measures of morphological deviation from circularity. All the quantitative measures described below support our qualitative observation.

The first of such measures, the one used in the main text, is eccentricity. Eccentricity quantifies how much a shape deviates from being a perfect circle. Day 1 images were segmented and automatically fit to an ellipse by our image processing algorithm. The eccentricity of each day 1 image is then calculated as:

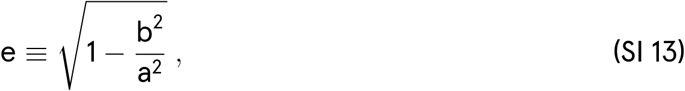

where b (a) is the shortest (longest) radius of the fitted ellipse. A value of *e* = 0 denotes a perfect circle; for 0 *<* e *<* 1, the increasing eccentricity indicates a more elongated shape. Our normalized (non-normalized, with predicted stages) analysis for eccentricity in day 1 images is found in Supplementary Fig. SI9 (Supplementary Fig. SI10).

The second of such measures is elongation, which quantifies how stretched out a shape is along its longest axis. The elongation of each day 1 image is calculated as:

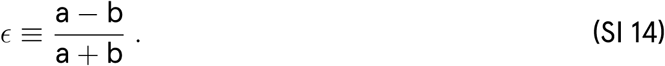

This measure behaves similarly to a normalized ratio of the axis lengths, indicating how stretched the ellipse is. The closer ɛ is to 1, the more elongated the shape becomes, while ɛ closer to 0 suggests a more circular shape. Our normalized (non-normalized, with predicted stages) analysis for elongation in day 1 images is found in Supplementary Fig. SI11 (Supplementary Fig. SI12).

The third measure of deviation from circularity is roundness, which compares the area of a shape to the area of a circle with the same perimeter. It is calculated as:

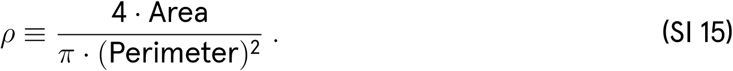

This formula is commonly used in image processing to quantify roundness, with values of *ρ* closer to 1 indicating a shape that is more circular. As the shape becomes more irregular or elongated, *ρ* decreases, indicating a deviation from circularity. Our normalized (non-normalized, with predicted stages) analysis for roundness in day 1 images is found in Supplementary Fig. SI13 (Supplementary Fig. SI14).

Finally, we use solidity, a measure of the compactness of a shape. It compares the area of the shape to the area of its convex hull, which is the smallest convex shape that can enclose the shape. Solidity is defined as:

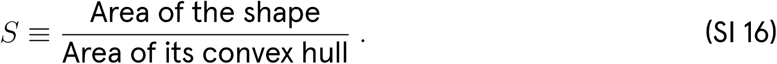

Solidity ranges from 0 to 1, with 1 indicating a shape with no concavities (*i.e.*, a convex shape), and lower values indicating that the shape is more irregular with more concavities. Our normalized (non-normalized, with predicted stages) analysis for solidity in day 1 images is found in Supplementary Fig. SI15 (Supplementary Fig. SI16).

Eccentricity, elongation, and roundness focus on the relationship between the shape’s axes or perimeter, while solidity involves the convex hull. All the above metrics compare the shape to a perfect circle or a convex shape to determine how closely it resembles those forms. Day 1 images, when ordered by decreasing solidity, show the strongest correlation with increasing developmental stage. As an example, Supplementary Fig. SI17 presents this ordering for all day 1 images in batch B2. In the main text, we selected eccentricity over solidity as it is a more intuitive measure.

The high variability of these measures across batches is noteworthy. Comparisons of the hypomagnetic environment’s effect are more easily interpreted in the normalized version of our plots, where each measure is adjusted relative to the control condition’s average.

The code output for day 1 images is identical to Fig. 1 d) , with each embryo being annotated with its programmatically determined major and minor axes.

### SI18. Day 1 eccentricity (norm.) results

**Fig. SI9:**
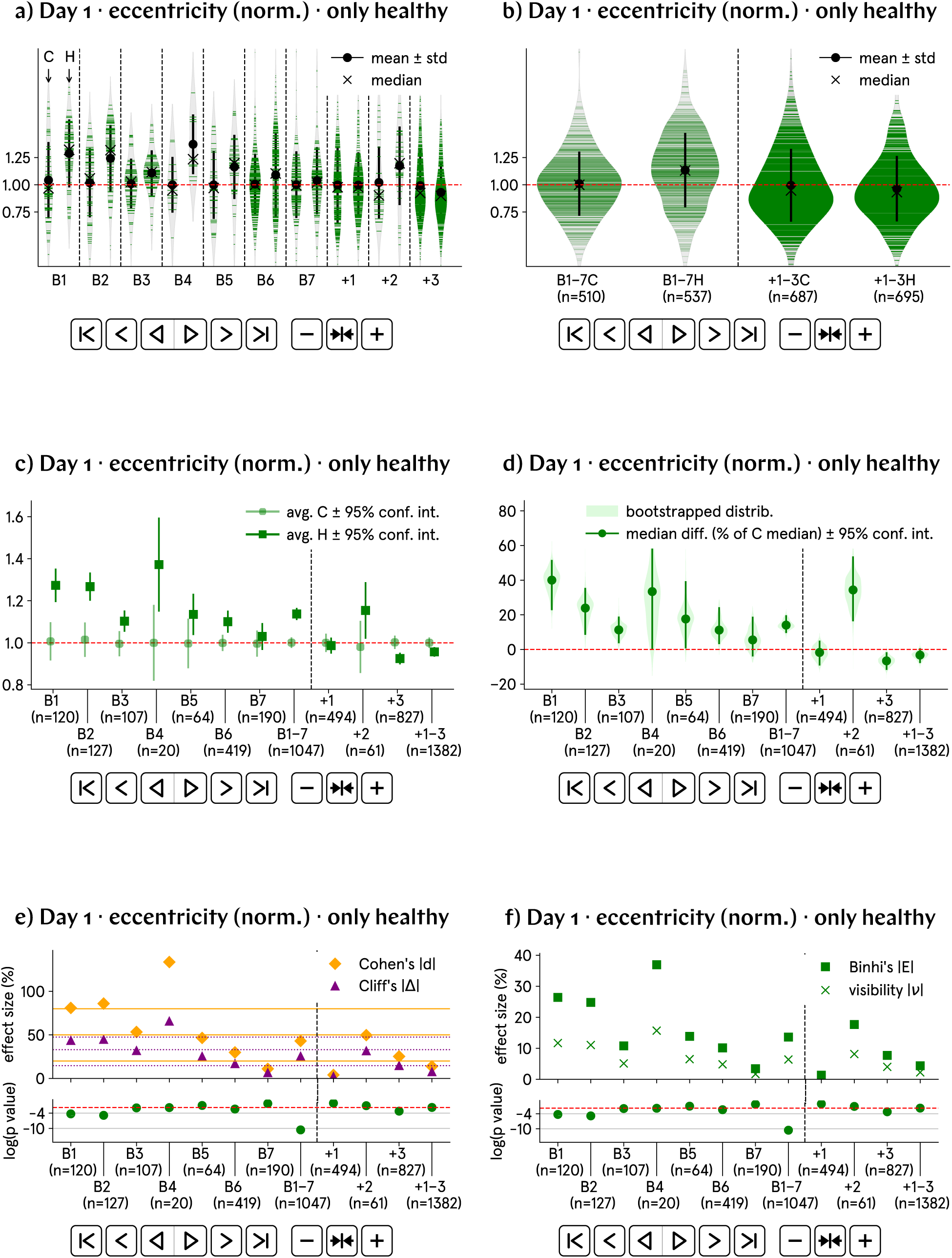
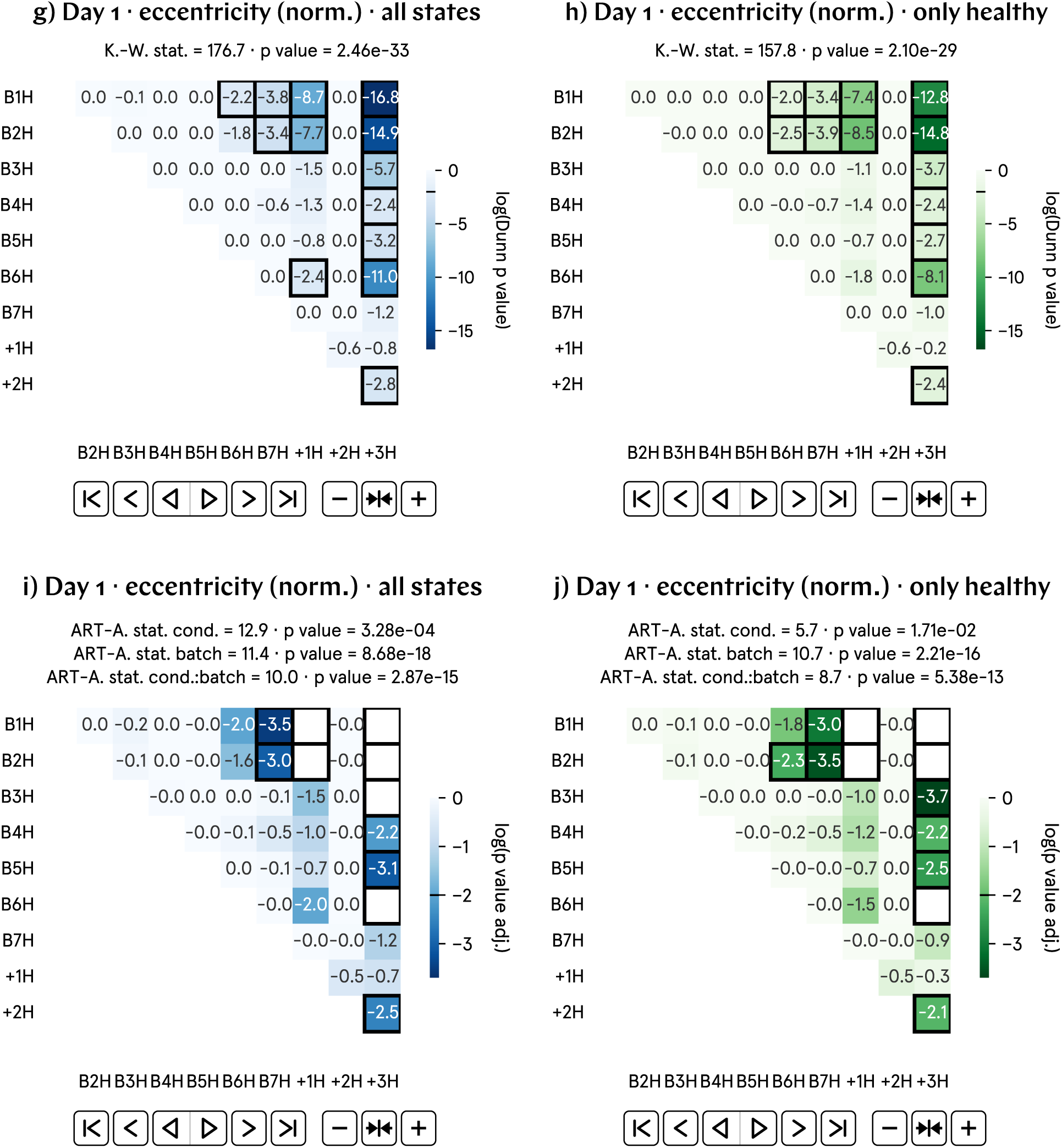
At day 1 p.f., the eccentricity of hypomagnetic embryos is approximately 20% higher than that of the control population. a) and b) Normalized eccentricity by batch and aggregate data; B1–7 represent experimental runs, and +1–3 correspond to positive control runs, in which an artificial magnetic field inside the hypomagnetic chamber simulates Earth’s. c) and d) Analysis of the population averages and medians further support the observed 20% higher eccentricity. e) and f) Effect size metrics, namely Cohen’s *d* and Cliff’s Δ, and Binhi’s *E* and the visibility *ν*, confirm a medium to large effect, consistently reflecting the 20% level difference. g) through j) Most statistically significant differences are found in the positive control tadpole populations raised in the hypomagnetic chamber, as expected.

### SI19. Day 1 eccentricity results

**Fig. SI10:**
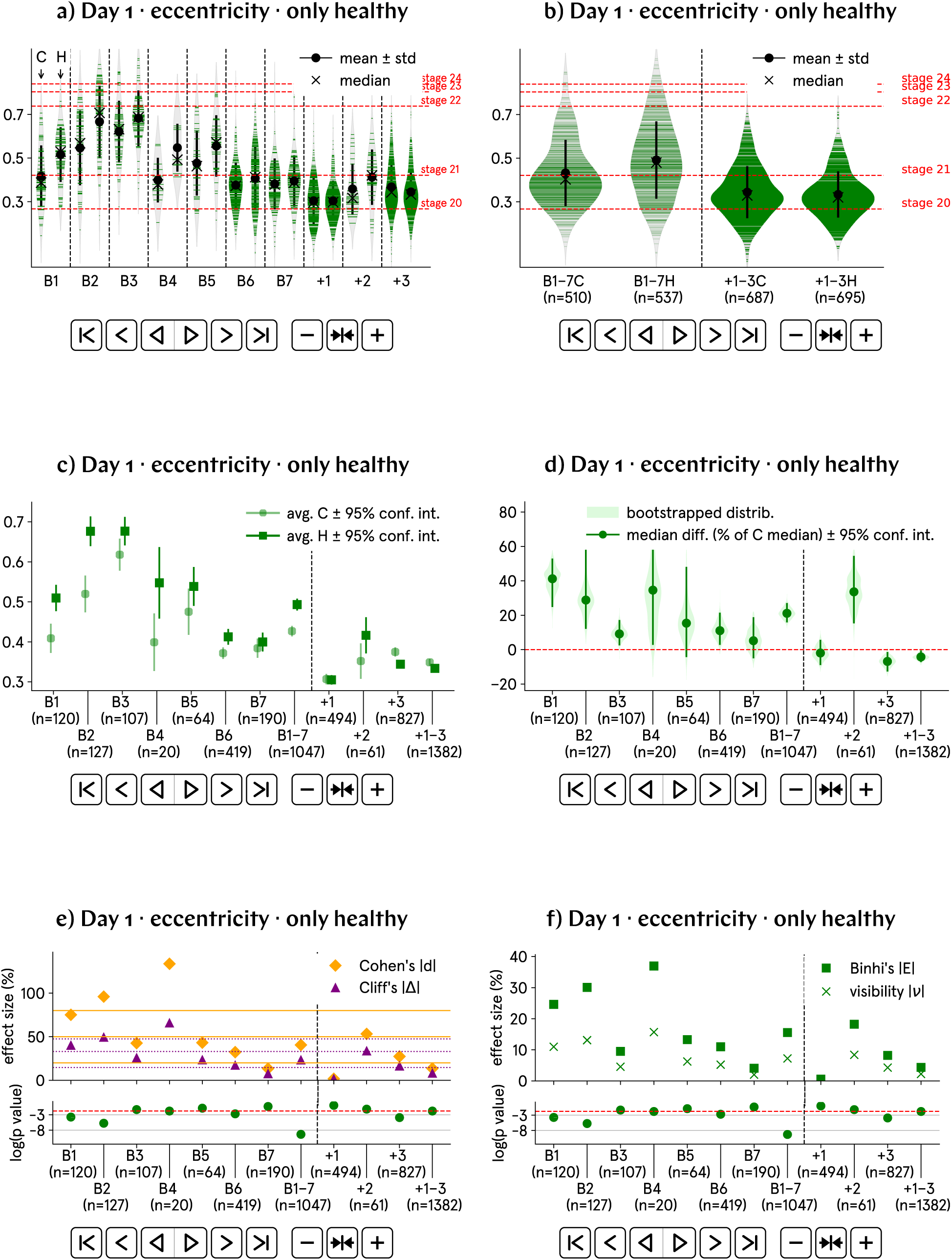
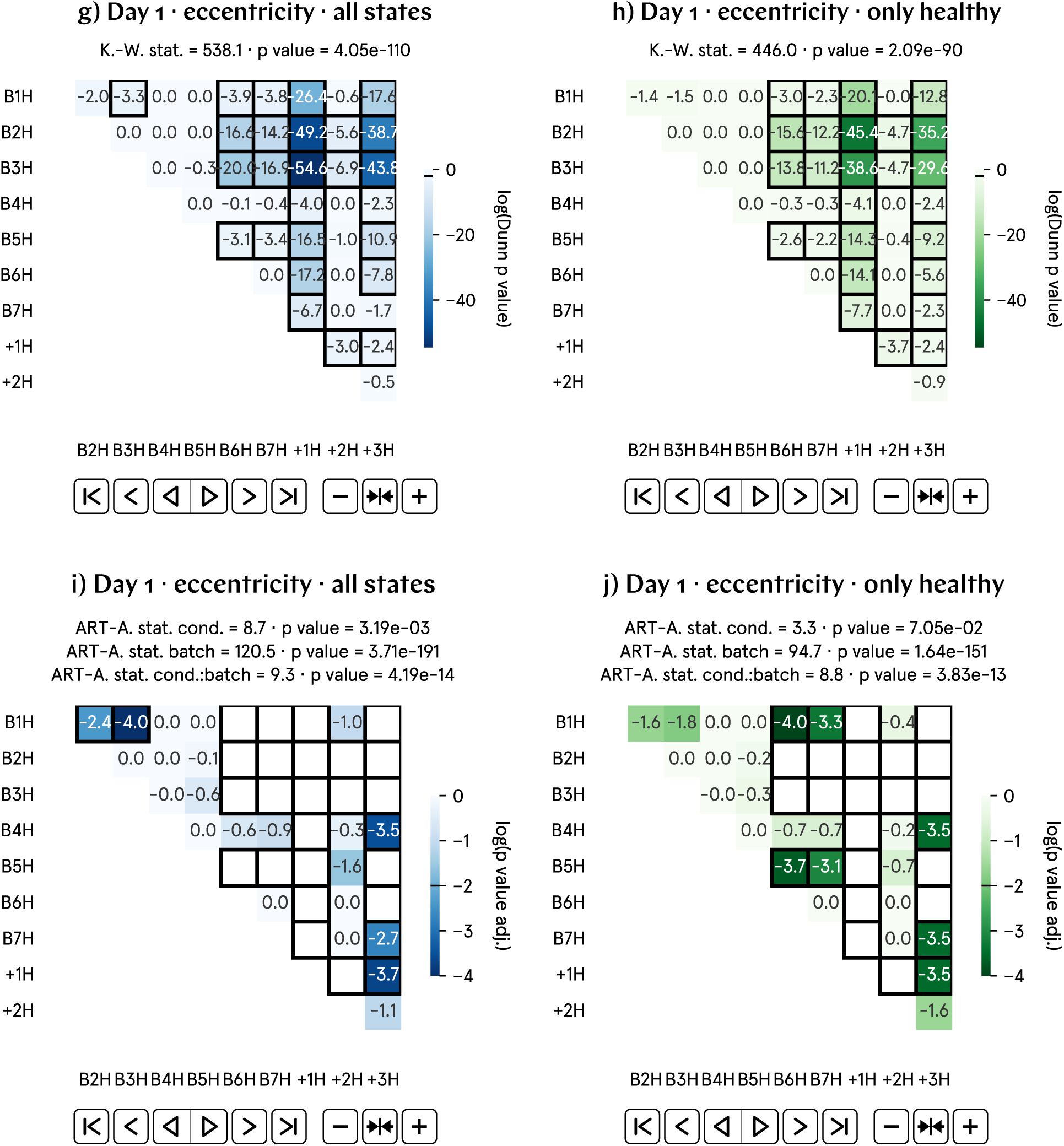
At day 1 p.f., the eccentricity of hypomagnetic embryos is approximately 20>% higher than that of the control population. a) and b) Non-normalized eccentricity by batch and aggregate data; B1–7 represent experimental runs, and +1–3 correspond to positive control runs, in which an artificial magnetic field inside the hypomagnetic chamber simulates Earth’s. c) and d) Analysis of the population averages and medians further support the observed 20% higher eccentricity. e) and f) Effect size metrics, namely Cohen’s *d* and Cliff’s Δ, and Binhi’s *E* and the visibility *ν*, confirm a medium to large effect, consistently reflecting the 20% level difference. g) through j) Given the substantial variation in eccentricity across batches, these plots may not provide an accurate representation of our data. Finally, we propose to automate embryo stage classification according to eccentricity, see predicted stages in a) and b).

### SI20. Day 1 elongation (norm.) results

**Fig. SI11:**
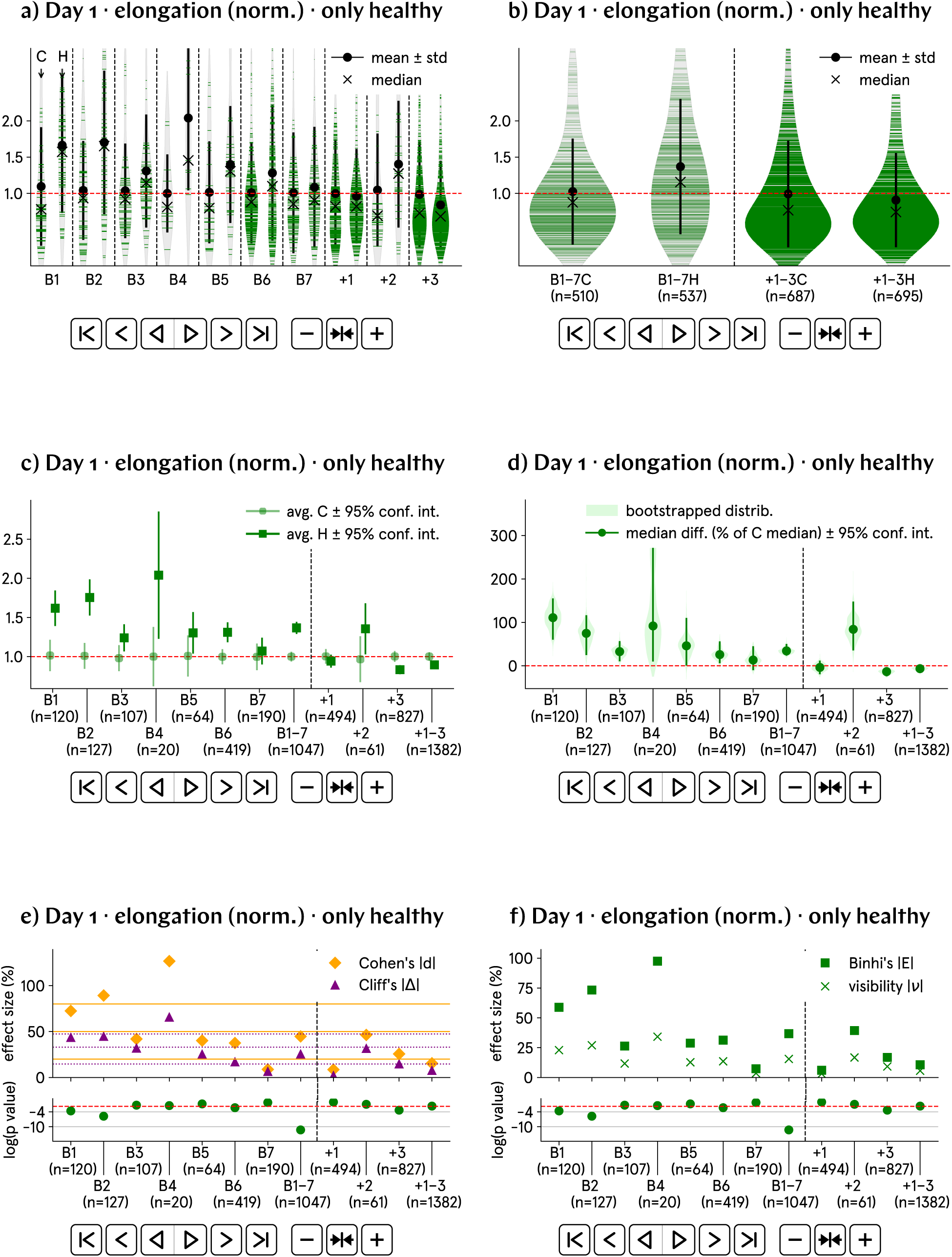
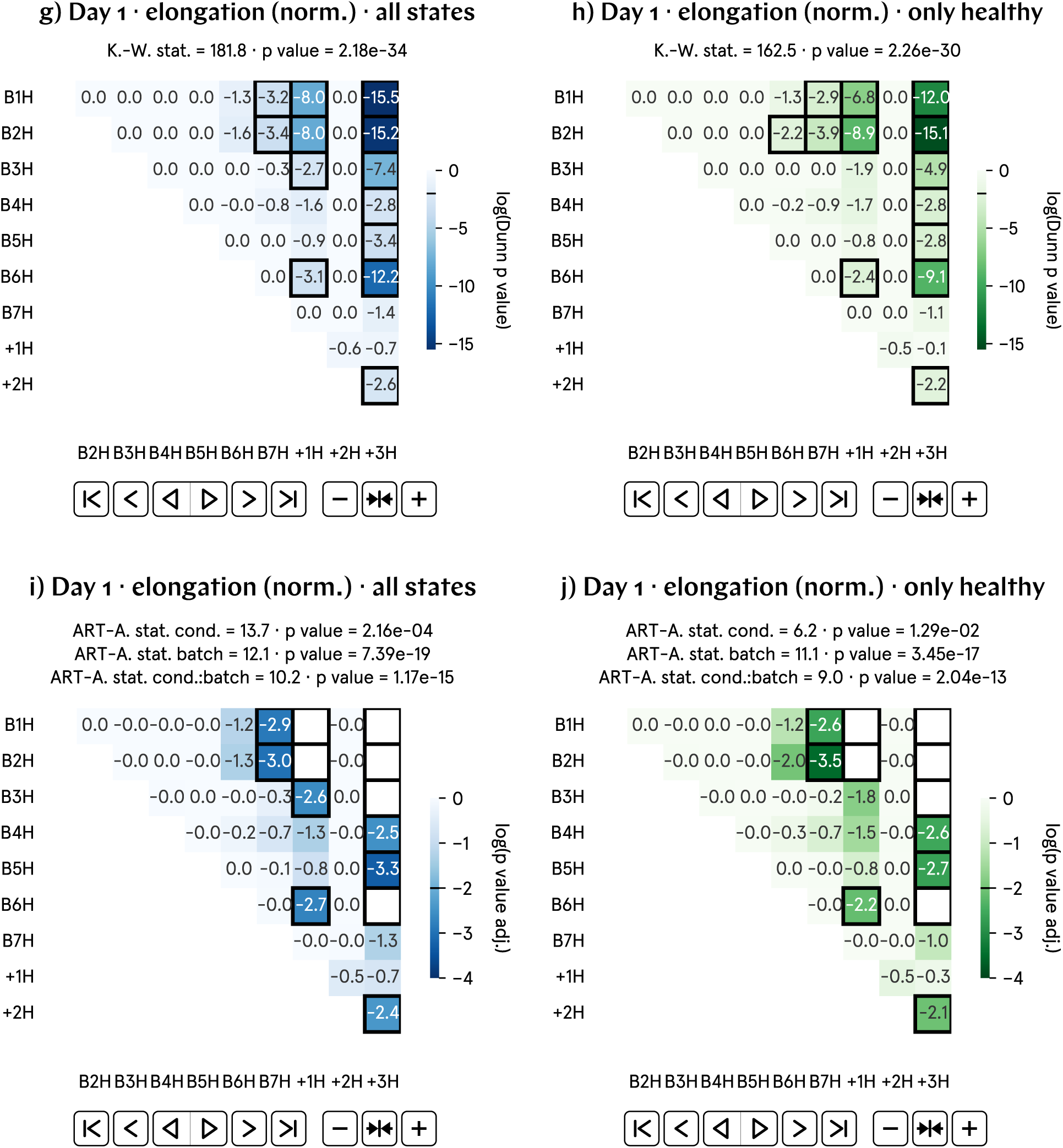
At day 1 p.f., the elongation of hypomagnetic embryos is approximately 45% higher than that of the control population. a) and b) Normalized elongation by batch and aggregate data; B1–7 represent experimental runs, and +1–3 correspond to positive control runs, in which an artificial magnetic field inside the hypomagnetic chamber simulates Earth’s. c) and d) Analysis of the population averages and medians further support the observed 45% higher elongation. e) and f) Effect size metrics, namely Cohen’s *d* and Cliff’s Δ, and Binhi’s *E* and the visibility *ν*, confirm a medium to large effect, consistently reflecting the 45% level difference. g) through j) Most statistically significant differences are found in the positive control tadpole populations raised in the hypomagnetic chamber, as expected.

### SI21. Day 1 elongation results

**Fig. SI12:**
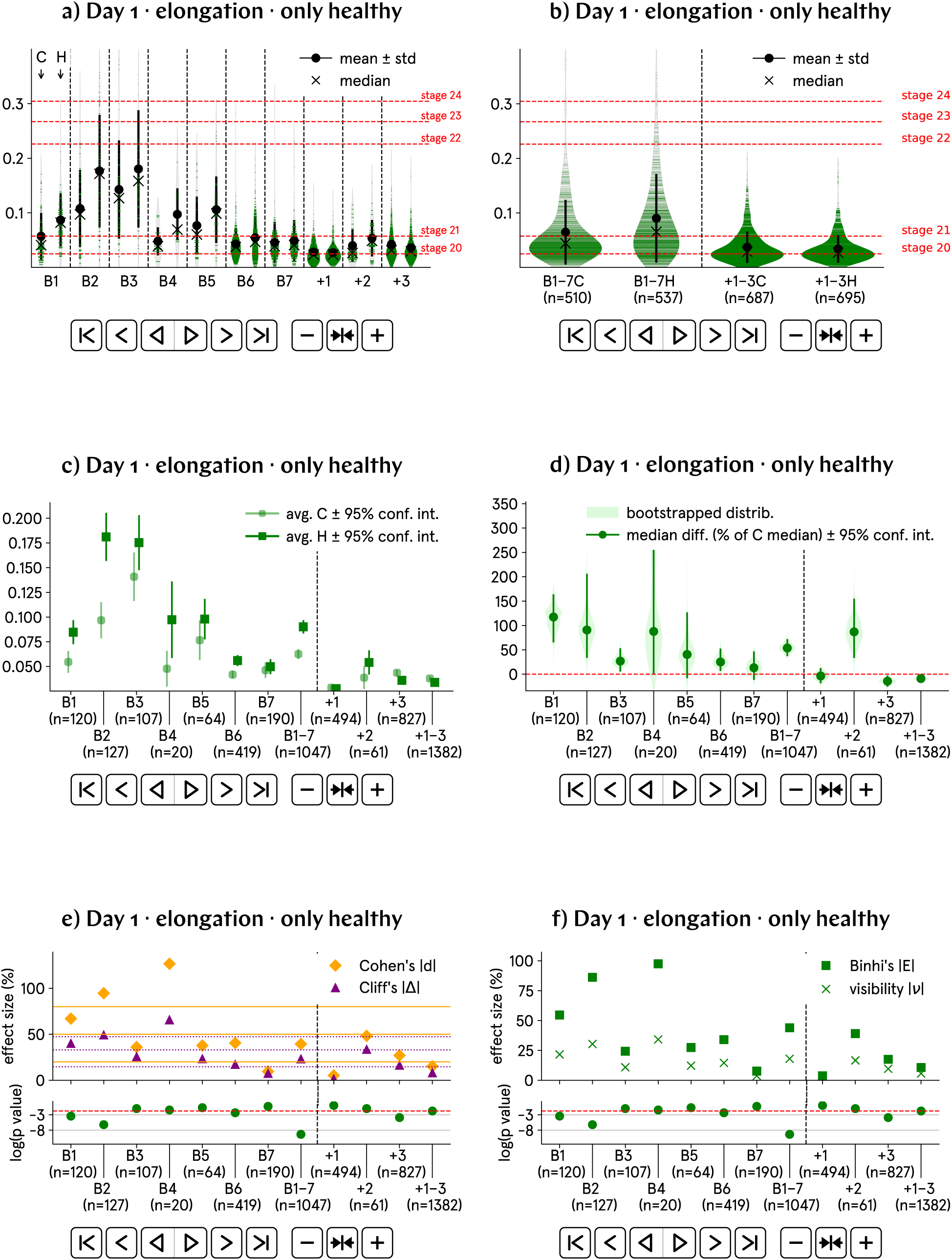
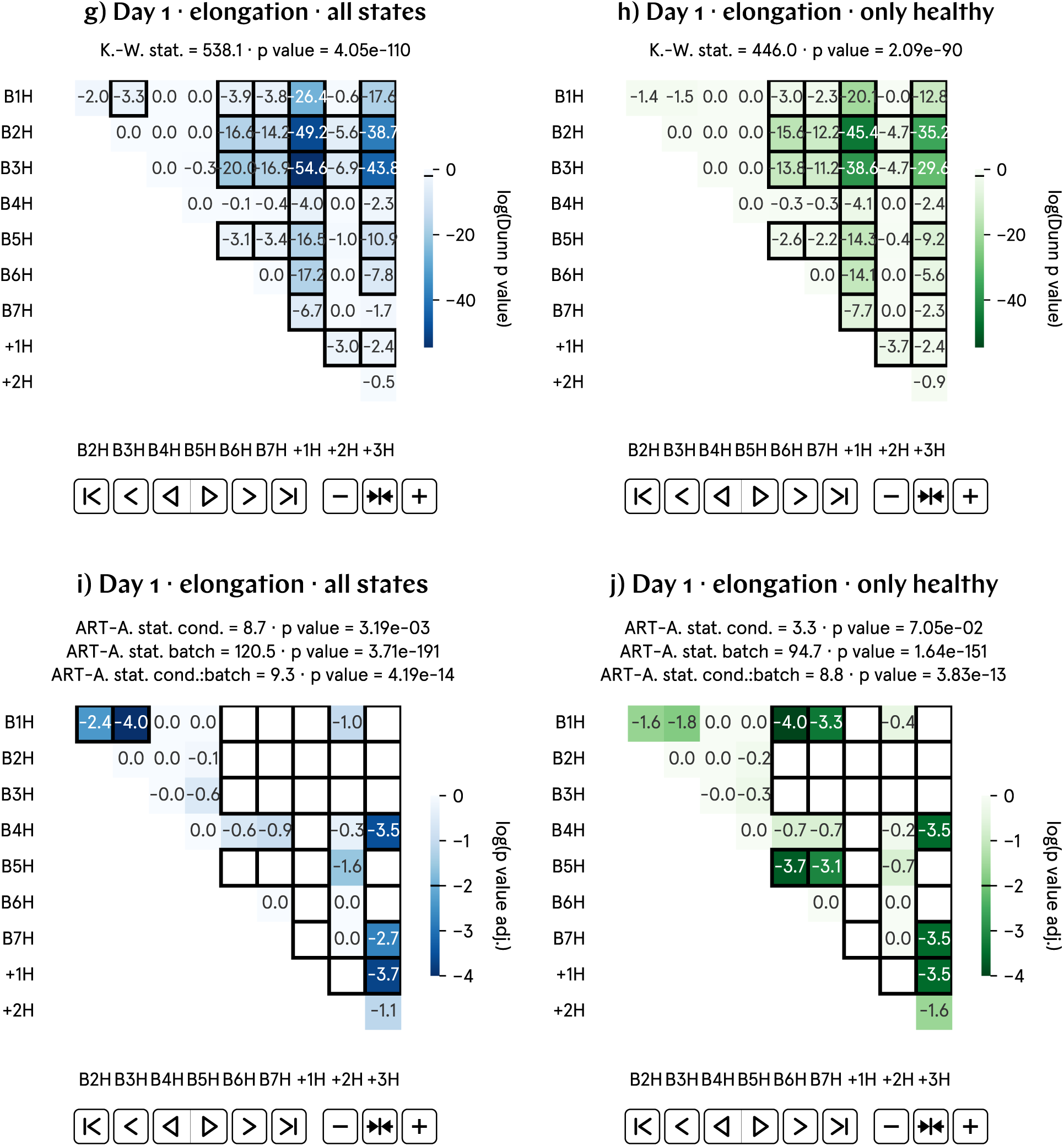
At day 1 p.f., the elongation of hypomagnetic embryos is approximately 45% higher than that of the control population. a) and b) Non-normalized elongation by batch and aggregate data; B1–7 represent experimental runs, and +1–3 correspond to positive control runs, in which an artificial magnetic field inside the hypomagnetic chamber simulates Earth’s. c) and d) Analysis of the population averages and medians further support the observed 45% higher elongation. e) and f) Effect size metrics, namely Cohen’s *d* and Cliff’s Δ, and Binhi’s *E* and the visibility *ν*, confirm a medium to large effect, consistently reflecting the 45% level difference. g) through j) Given the substantial variation in elongation across batches, these plots may not provide an accurate representation of our data. Finally, we propose to automate embryo stage classification according to elongation, see predicted stages in a) and b).

### SI22. Day 1 roundness (norm.) results

**Fig. SI13:**
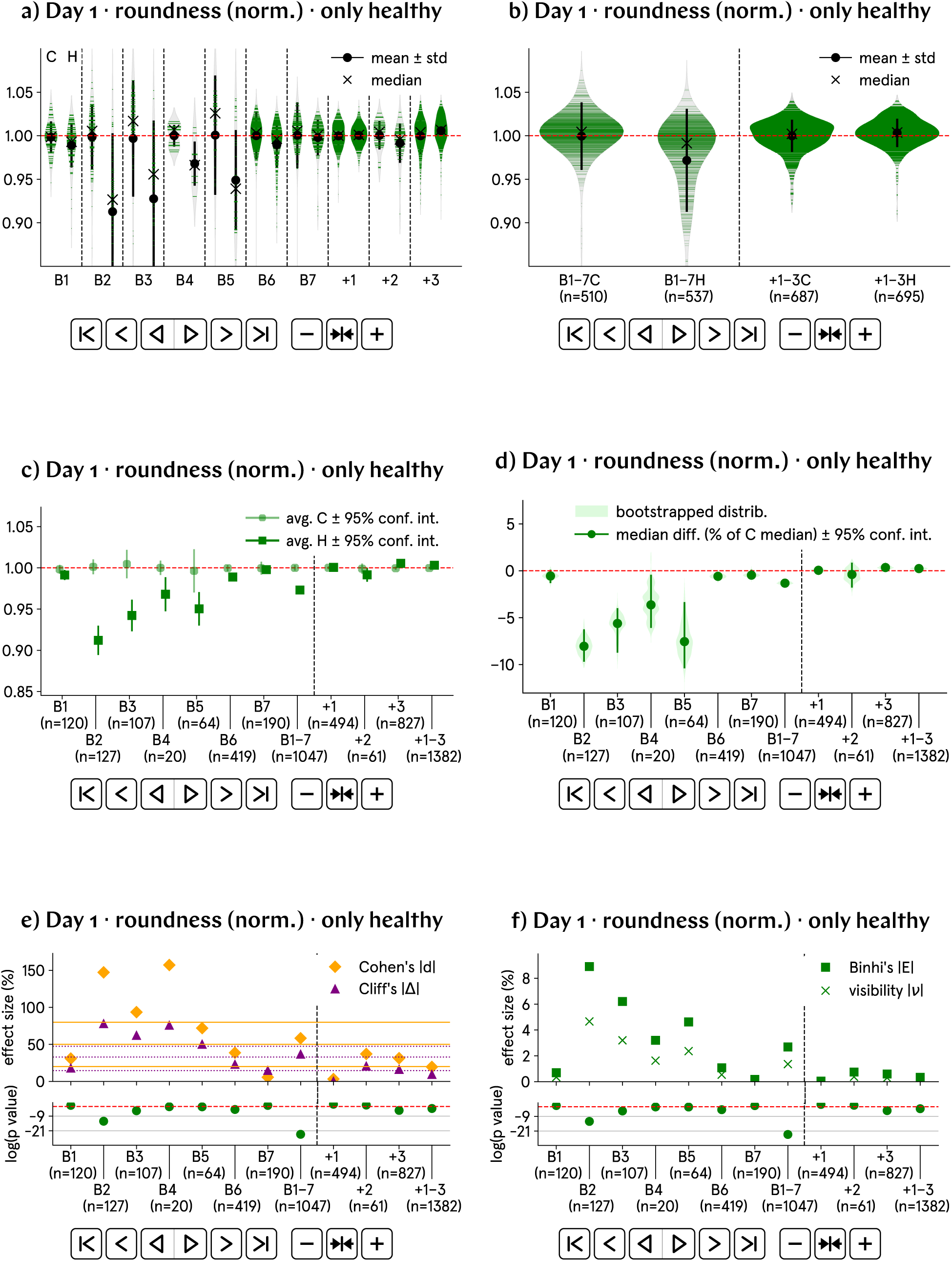
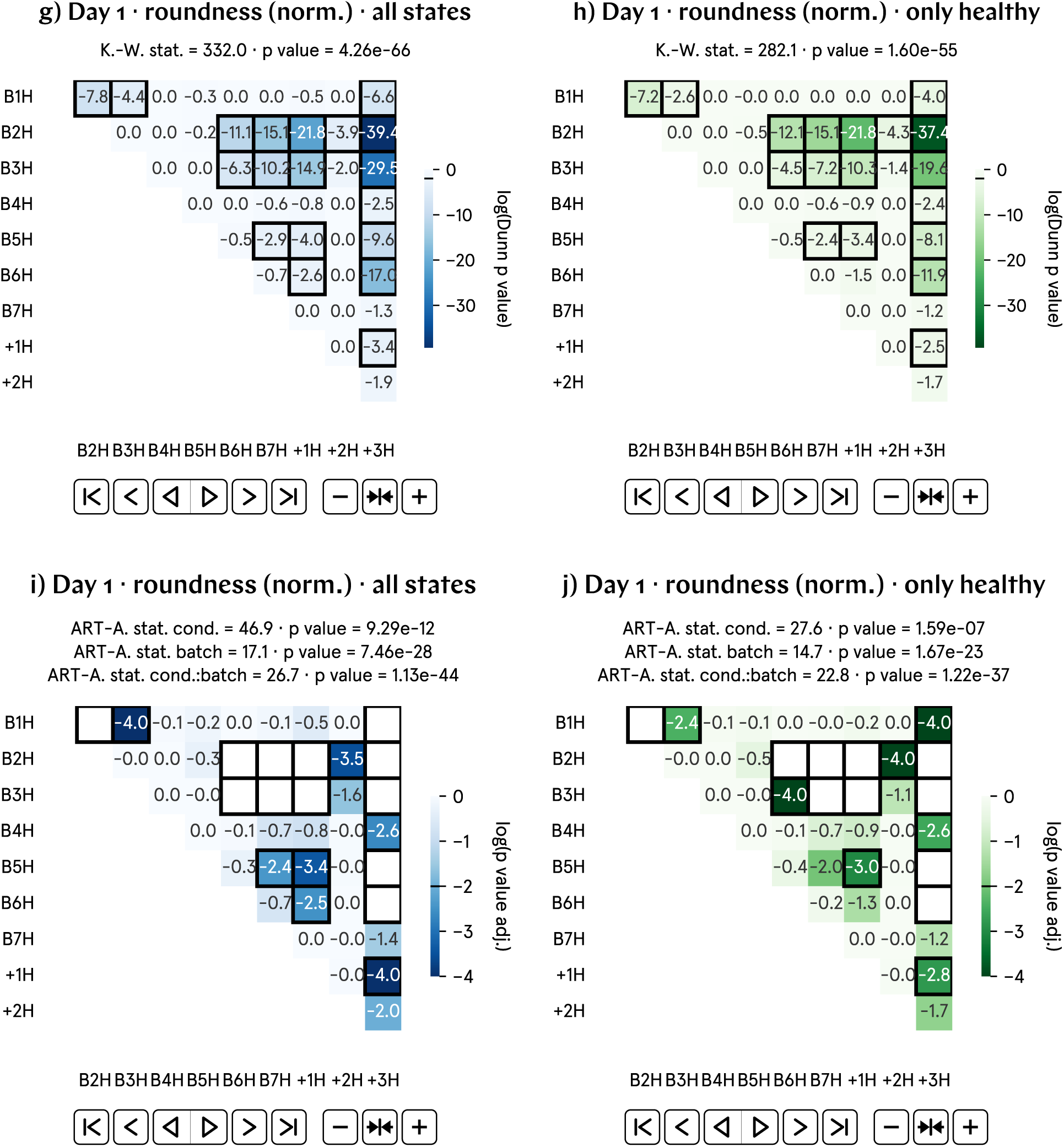
At day 1 p.f., the roundness of hypomagnetic embryos is approximately 3% lower than that of the control population. a) and b) Normalized roundness by batch and aggregate data; B1–7 represent experimental runs, and +1–3 correspond to positive control runs, in which an artificial magnetic field inside the hypomagnetic chamber simulates Earth’s. c) and d) Analysis of the population averages and medians further support the observed 3% lower roundness. e) and f) Effect size metrics, namely Cohen’s *d* and Cliff’s Δ, and Binhi’s *E* and the visibility *ν*, confirm a medium to large effect, consistently reflecting the 3% level difference. g) through j) There is significant variability in roundness across batches, even in the normalized data.

### SI23. Day 1 roundness results

**Fig. SI14:**
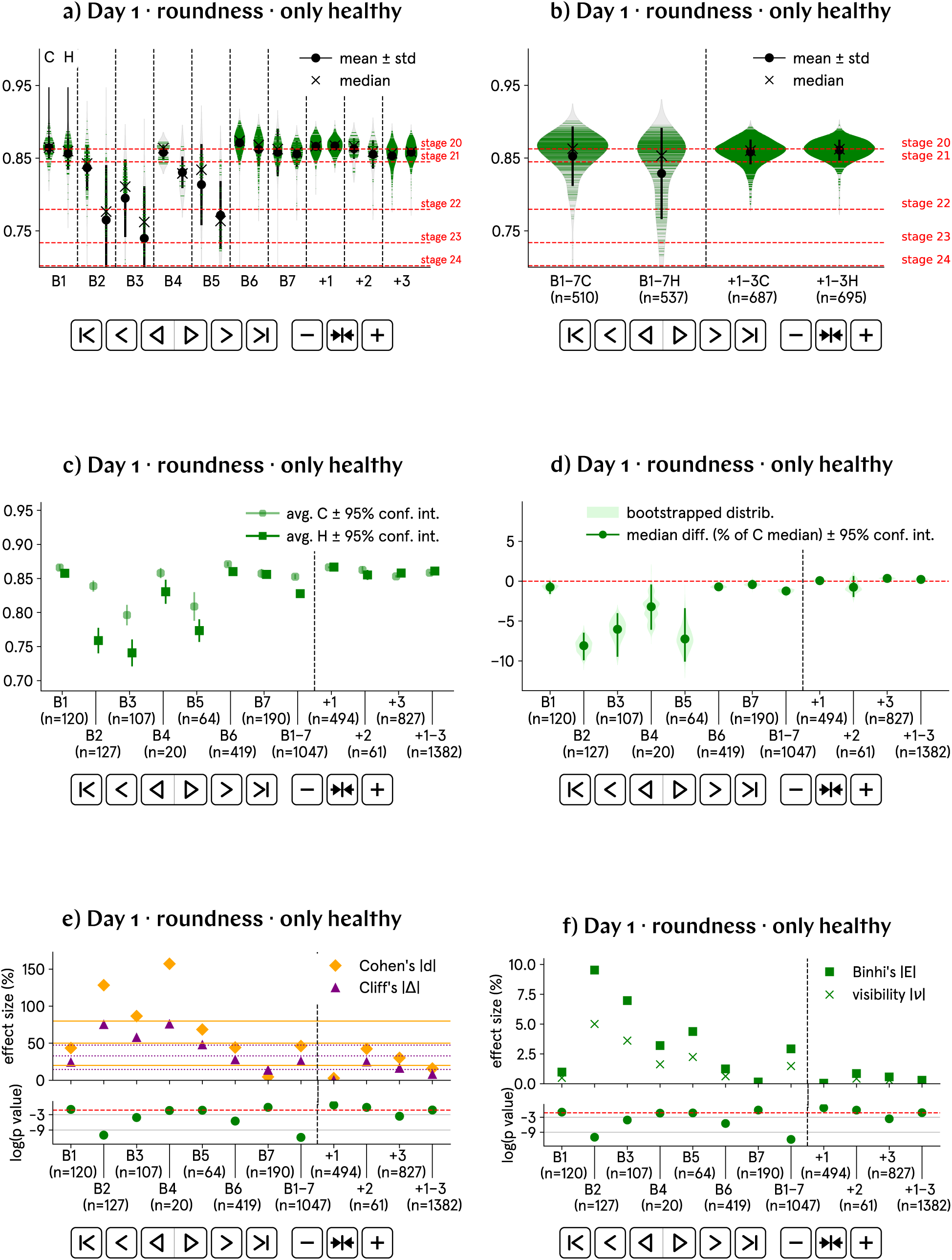
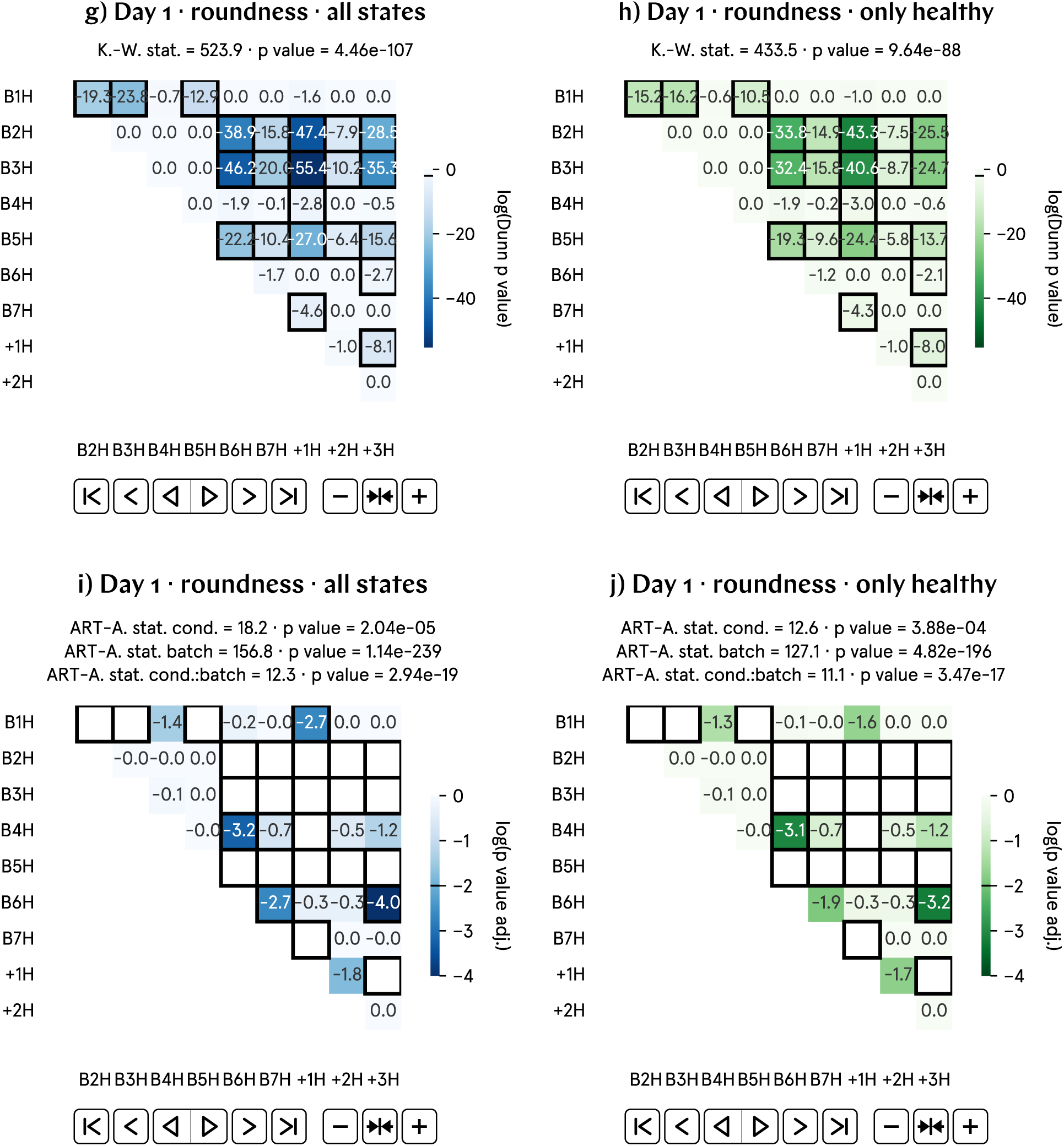
At day 1 p.f., the roundness of hypomagnetic embryos is approximately 3% lower than that of the control population. a) and b) Non-normalized roundness by batch and aggregate data; B1–7 represent experimental runs, and +1–3 correspond to positive control runs, in which an artificial magnetic field inside the hypomagnetic chamber simulates Earth’s. c) and d) Analysis of the population averages and medians further support the observed 3% lower roundness. e) and f) Effect size metrics, namely Cohen’s *d* and Cliff’s Δ, and Binhi’s *E* and the visibility *ν*, confirm a medium to large effect, consistently reflecting the 3% level difference. g) through j) Given the substantial variation in roundness across batches, these plots may not provide an accurate representation of our data. Finally, we propose to automate embryo stage classification according to roundness, see predicted stages in a) and b).

### SI24. Day 1 solidity (norm.) results

**Fig. SI5:**
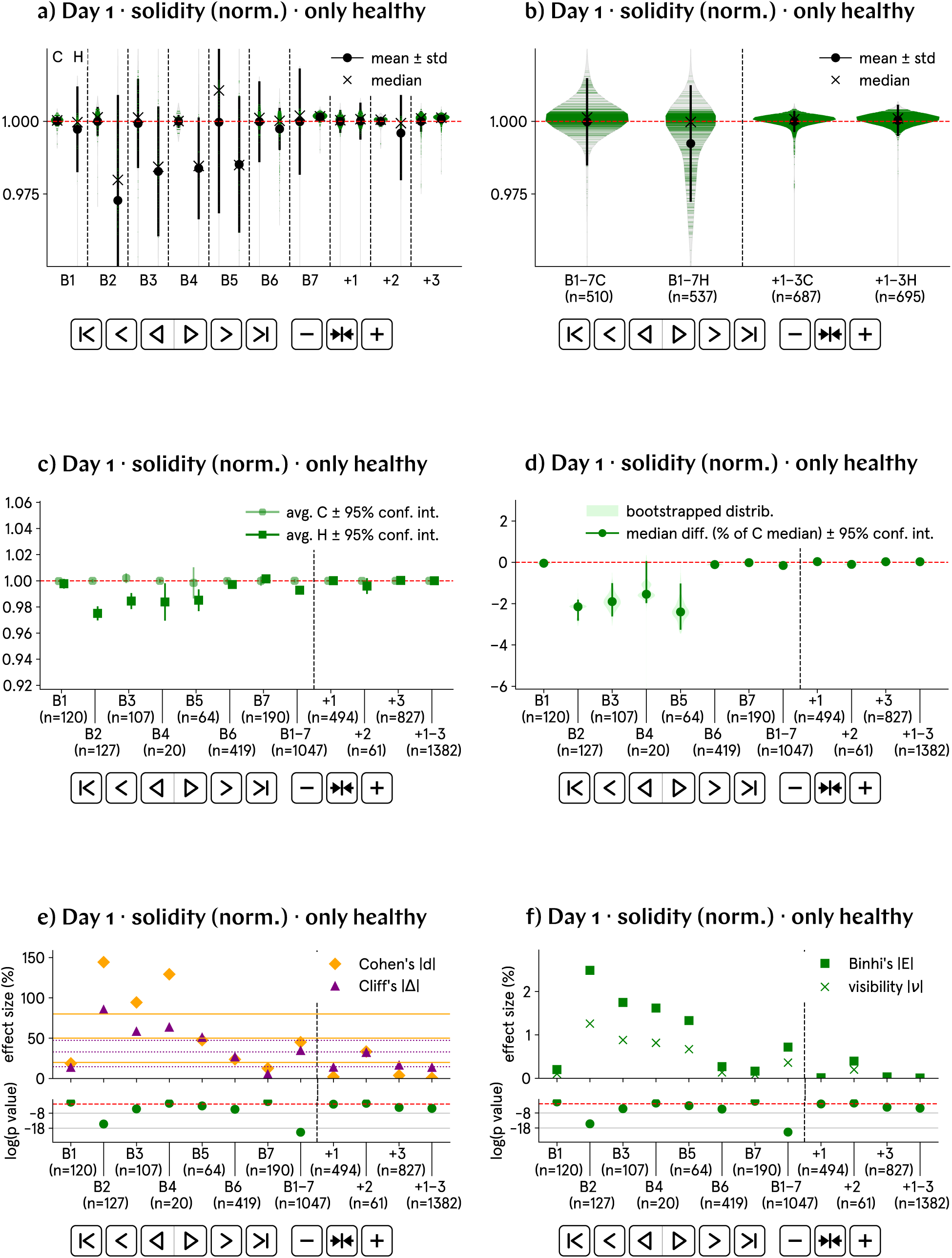
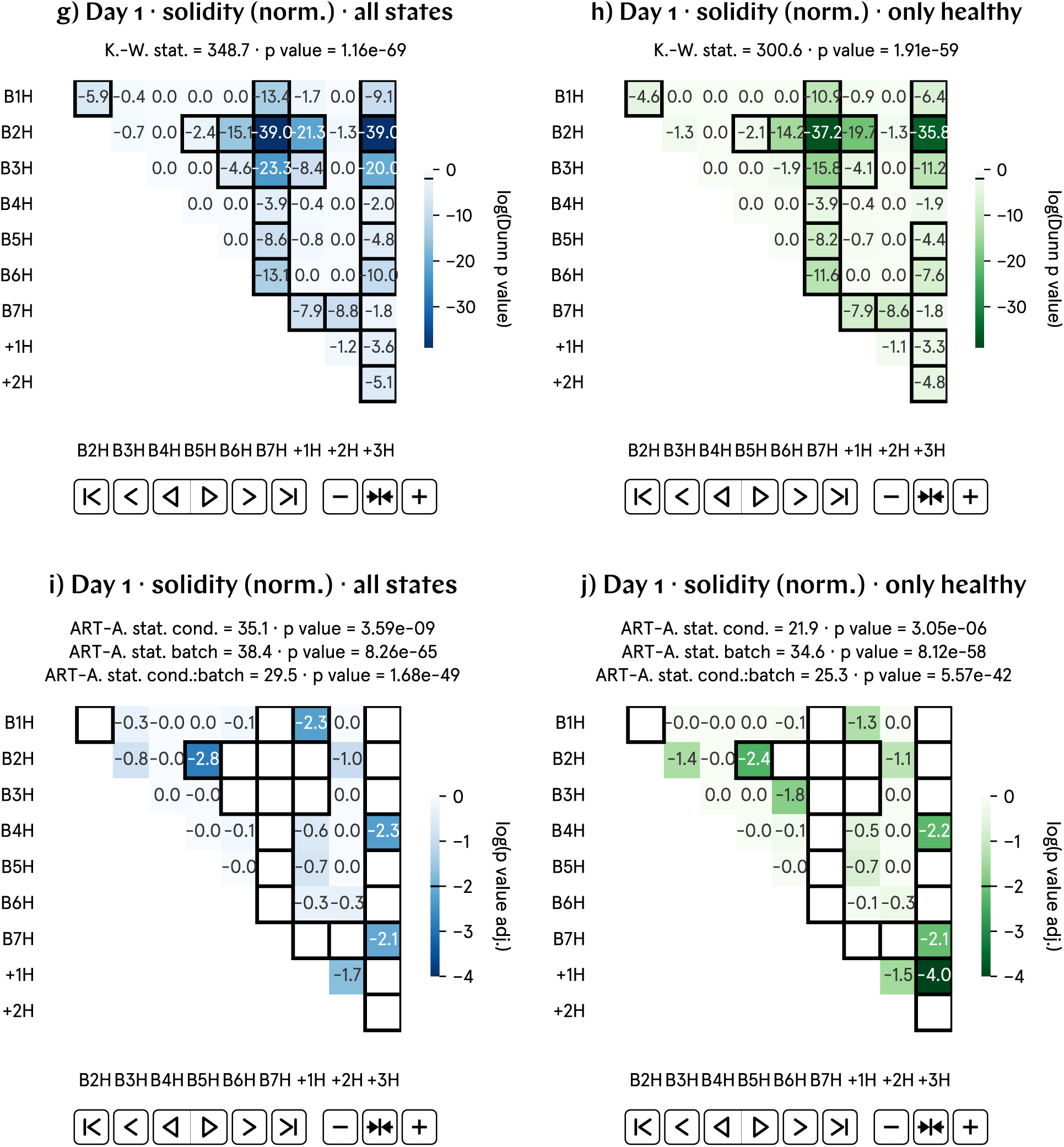
Residual field mapping and calibration. a) The magnetic field was mapped over a volume of 16’ 16’ 16’ and the residual field was measured in each of the points represented. b) The mapped field shows the field uniformity and can be inspected in the 3D interactive HTML plot available in the GitHub repository (see text for details). The photo in a) is taken with the same perspective as the plot in b).

### SI25. Day 1 solidity results

**Fig. SI16:**
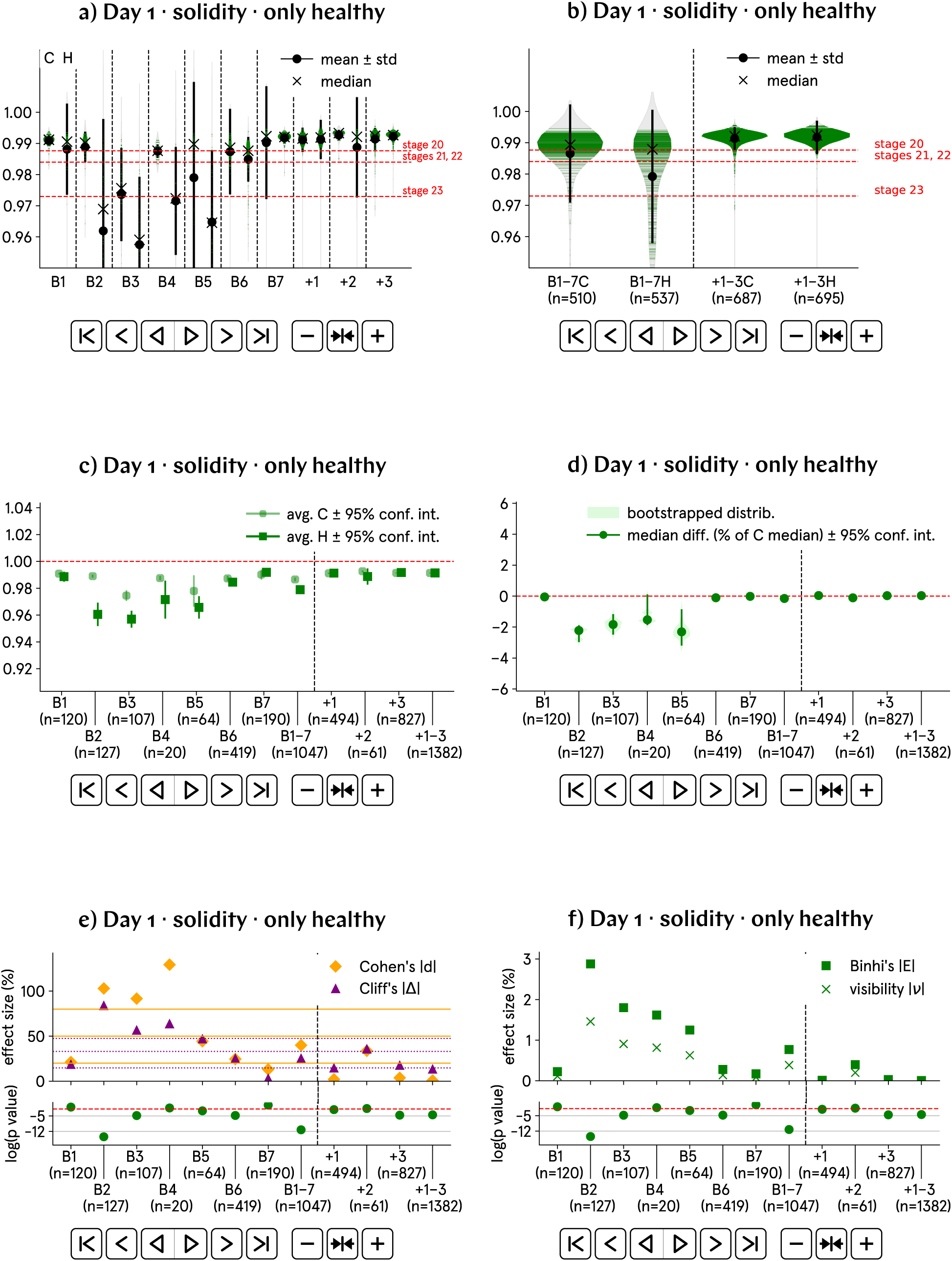
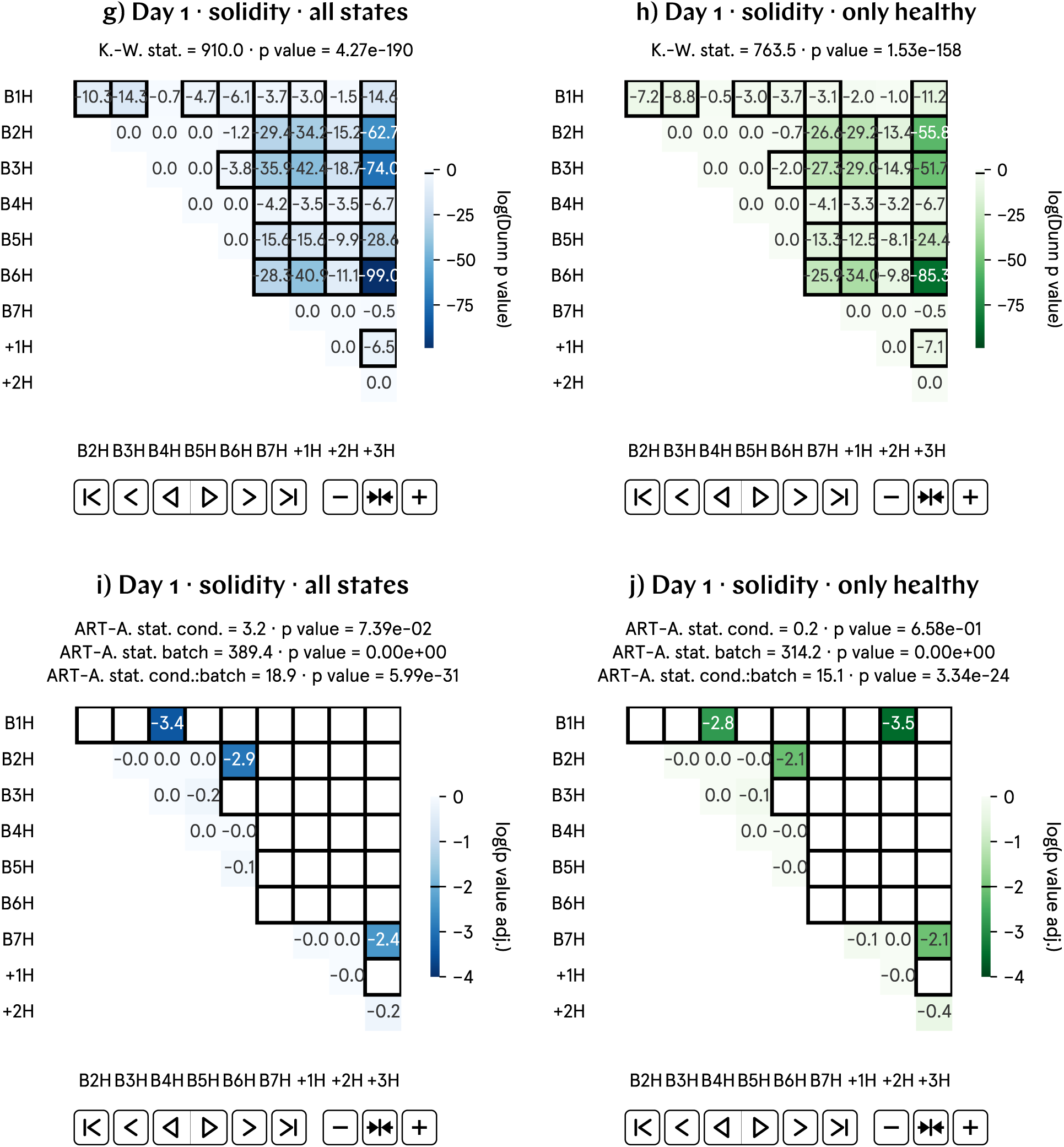
At day 1 p.f., the solidity of hypomagnetic embryos is approximately 1% lower than that of the control population. a) and b) Non-normalized solidity by batch and aggregate data; B1–7 represent experimental runs, and +1–3 correspond to positive control runs, in which an artificial magnetic field inside the hypomagnetic chamber simulates Earth’s. c) and d) Analysis of the population averages and medians further support the observed 1% lower solidity. e) and f) Effect size metrics, namely Cohen’s *d* and Cliff’s Δ, and Binhi’s *E* and the visibility *ν*, confirm a medium to large effect, consistently reflecting the 1% level difference. g) through j) Given the substantial variation in solidity across batches, these plots may not provide an accurate representation of our data. Finally, we propose to automate embryo stage classification according to solidity, see predicted stages in a) and b)

**Fig. SI17:**
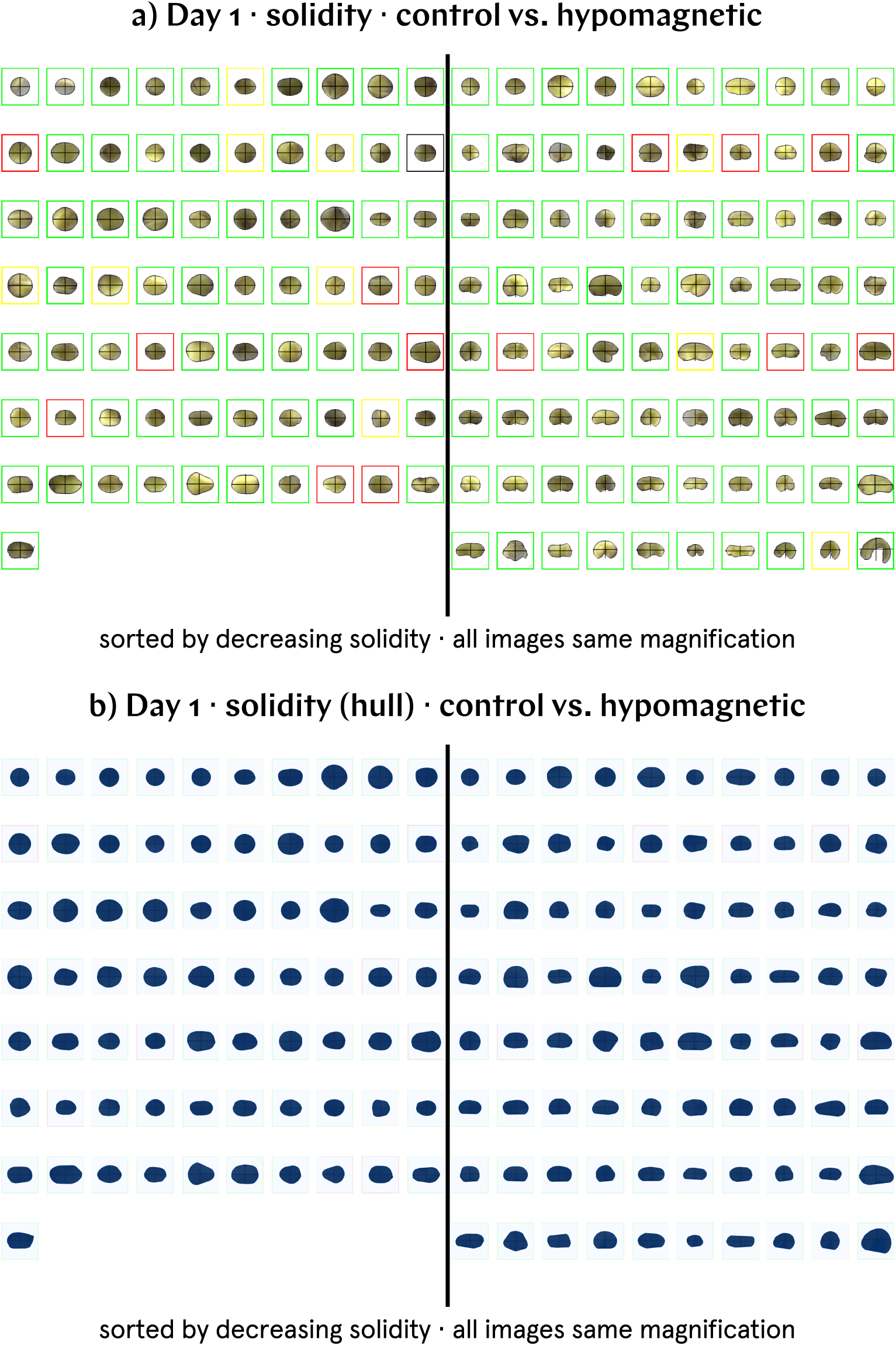
Batch B2 day 1 images ordered by decreasing solidity. a) Decreasing solidity is strongly correlated with advanced developmental stages, as the embryos transition in shape from egg-like to bean-like, and eventually to a more worm-like form. b) Solidity is defined by the area of the shape divided by the area of its convex hull; bean-like shapes with indentations have lower solidities than circles or ellipses. For illustration, we provide here the convex hull images of the tadpoles depicted in a).

### SI26. Day 2 color measures: L*^*^*a*^*^*b*^*^* yellowness, RGB yellowness, Yellowness Index, and HSV yellowness

From day 2 p.f. onwards, we observed a slight difference in color between the tadpoles in the different conditions: Hypomagnetic condition tadpoles seemed to be paler in yellow color. To quantify this qualitative observation for day 2 p.f. images, we employ four quantitative measures of yellowness; each measure is calculated after the following binarization procedure.

Day 2 RGB images from each separate batch are binarized using a global threshold value based on their combined histogram, ensuring consistency across all images in the same batch. To achieve this, a combined histogram of pixel intensities is created from all the images in a batch. The mean of the non-zero values in the combined histogram is calculated to approximate the average pixel intensity across the batch. If the histogram shows multiple peaks, the midpoint between the peaks is chosen as the global threshold. Otherwise, the mean of the non-zero values is used. After binarization, the white pixels are considered a better representation of the true tadpole color, as the darker pigmentation caused by melanin is more likely to be binarized as black. Color measures are then applied only to the white pixels in the binary mask.

The first of such yellowness measures, the one used in the main text, is the L*a*b* color space yellowness. L*a*b* [20] is a color space designed to be more perceptually uniform than the traditional RGB or CMYK color spaces, meaning that small changes in color correspond to similar changes in perceived color differences for humans. L*a*b* is widely used in image processing and computer vision because it separates color information from luminance and offers a more perceptually accurate representation of color. The pixels assigned to white in the binary mask are converted from RGB to L*a*b* color space; the b* channel (ranging from -128 to +127) represents the blue-yellow axis, with positive values corresponding to yellow. Our normalized (non-normalized) analysis for L*a*b* yellowness in day 2 images is found in Supplementary Fig. SI19 (Supplementary Fig. SI20).

The second color measure is the RGB yellowness. For each RGB pixel, we compute:

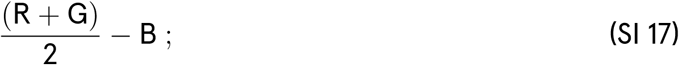

in other words, we average the red and green channels (which together contribute to yellow, since yellow in the RGB model is formed by mixing red and green) and subtract the blue channel (since blue is complementary to yellow; the more blue there is, the less yellow a pixel appears). This measure, ranging from *±*255, helps differentiate how yellow a pixel is relative to its blue component. The mean value of this RGB yellowness measure is calculated over the entire image. Positive values indicate more yellow (higher red and green values compared to blue), and negative values indicate more blue (higher blue values relative to red and green). Our normalized (non-normalized) analysis for RGB yellowness in day 2 images is found in Supplementary Fig. SI21 (Supplementary Fig. SI22).

The third color measure we employ is the Yellowness Index [59]. This measure is used to quantify the degree of yellowness in a material or an image, often in comparison to a reference white point, typically under a specific light source such as D65 (daylight). The formula below is specific for comparison with the D65 illuminant and involves converting the RGB values to the XYZ color space, which better reflects how humans perceive color. After conversion, we compute for each pixel:

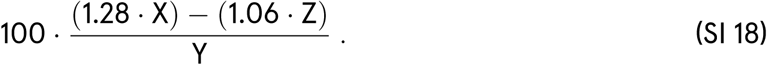

X, Y, and Z are the color values in the XYZ color space, which is a perceptually uniform color space designed to separate luminance (Y) from chromatic information (X, representing red-green colors, and Z, representing blue colors). 1.28 and 1.06 are constants to emphasize the balance between the X and Z components. The formula is structured to compare the values on the blue-yellow spectrum, where positive values correspond to a yellowish hue. This metric does not have strict predefined bounds; however, typical values are often found to be between 0 and 100 for yellowish materials. After the Yellowness Index is computed for each pixel, the image average is taken to describe the yellowness of the entire image under the D65 illuminant. Our normalized (non-normalized) analysis for the Yellowness Index in day 2 images is found in Supplementary Fig. SI23 (Supplementary Fig. SI24).

Finally, the HSV yellowness measure is calculated by first converting the RGB image to the HSV color space [60]. The HSV color space separates color information into three components, namely: the hue angle, H (representing the type of color; in this case, we are interested in yellow, which falls between 45*◦* and 75*◦*); the saturation S (measuring the intensity or purity of the color, measured in percent); and the value V (representing the brightness of the color, ranging from 0 to 1). Once the image is in the HSV color space, we filter for pixels that fall within the yellow hue range; for the pixels that pass this filter, we compute the below defined HSV yellowness measure:

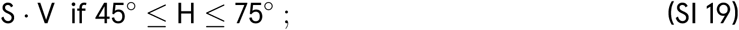

in other words, the HSV yellowness is the product of the saturation (how intense the yellow is) and value (how bright the yellow is). This means the yellowness is higher if the yellow is both bright and intense. This product is averaged over all pixels that fall within the yellow hue range to get the final yellowness value of the image. If no pixels fall within this hue range, the HSV yellowness is set to zero. The bounds for the calculated HSV yellowness are between 0 (no yellowness or desaturated/dark pixels) and 1 (maximum yellowness with fully saturated and bright yellow pixels). Our normalized (non-normalized) analysis for HSV yellowness in day 2 images is found in Supplementary Fig. SI25 (Supplementary Fig. SI26).

Similarly to day 1 images, the high variability of these measures across batches makes it more straightforward to perform comparisons in the normalized version of our plots.

A sample code output for day 2 images (of batch B4, in this example) is found in Supplementary Fig. SI18; only the programatically determined white pixels of the binary mask, shown, are taken into account in the calculation of yellowness.

**Fig. SI18:**
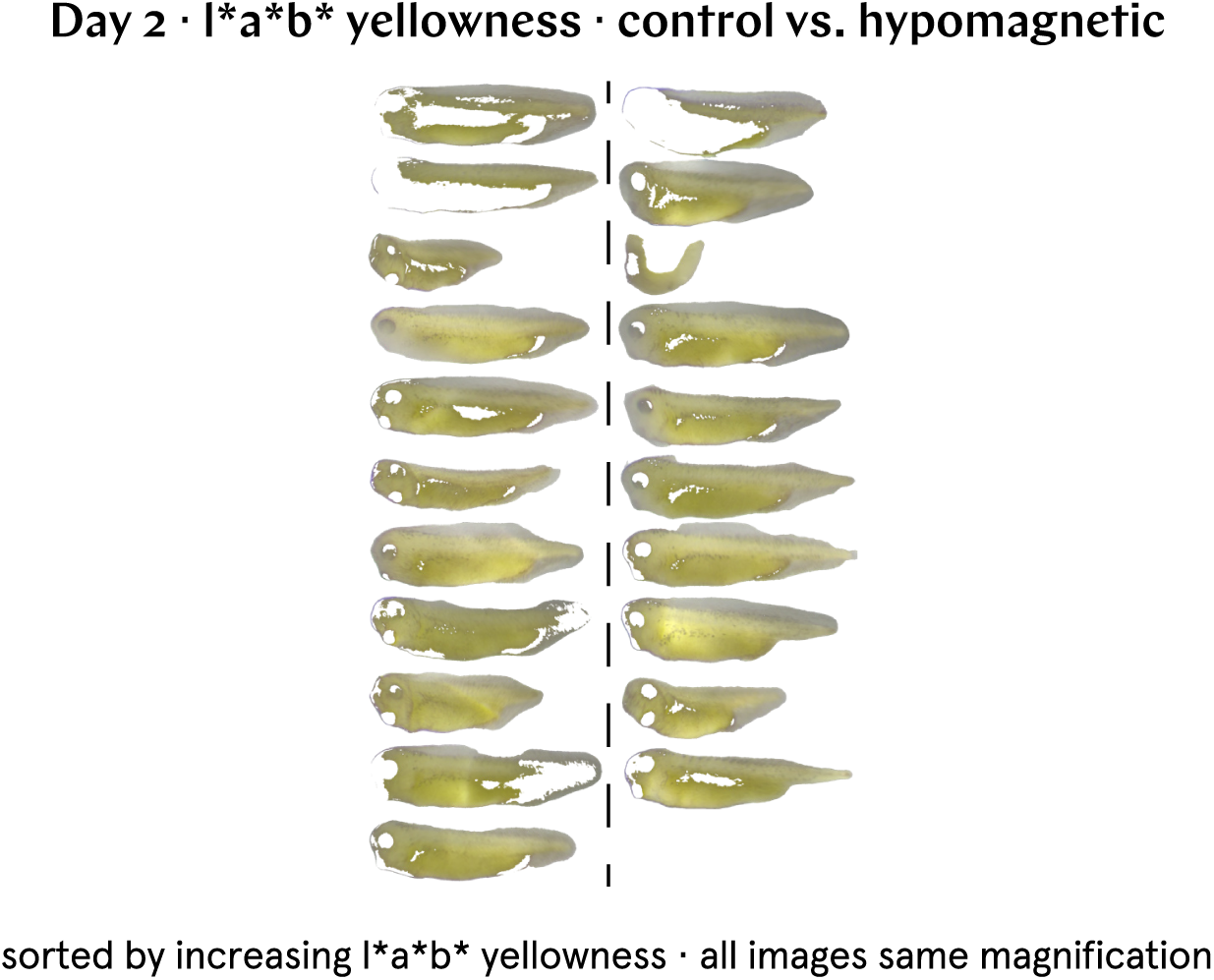
Sample code output for day 2 images. Day 2 images are binarized, and the light pixels are analyzed to assess various measures of yellow pigmentation. Here, batch B4 day 2 images are sorted by increasing L*a*b* yellowness.

### SI27. Day 2 L*^*^*a*^*^*b*^*^* yellowness (norm.) results

**Fig. SI19:**
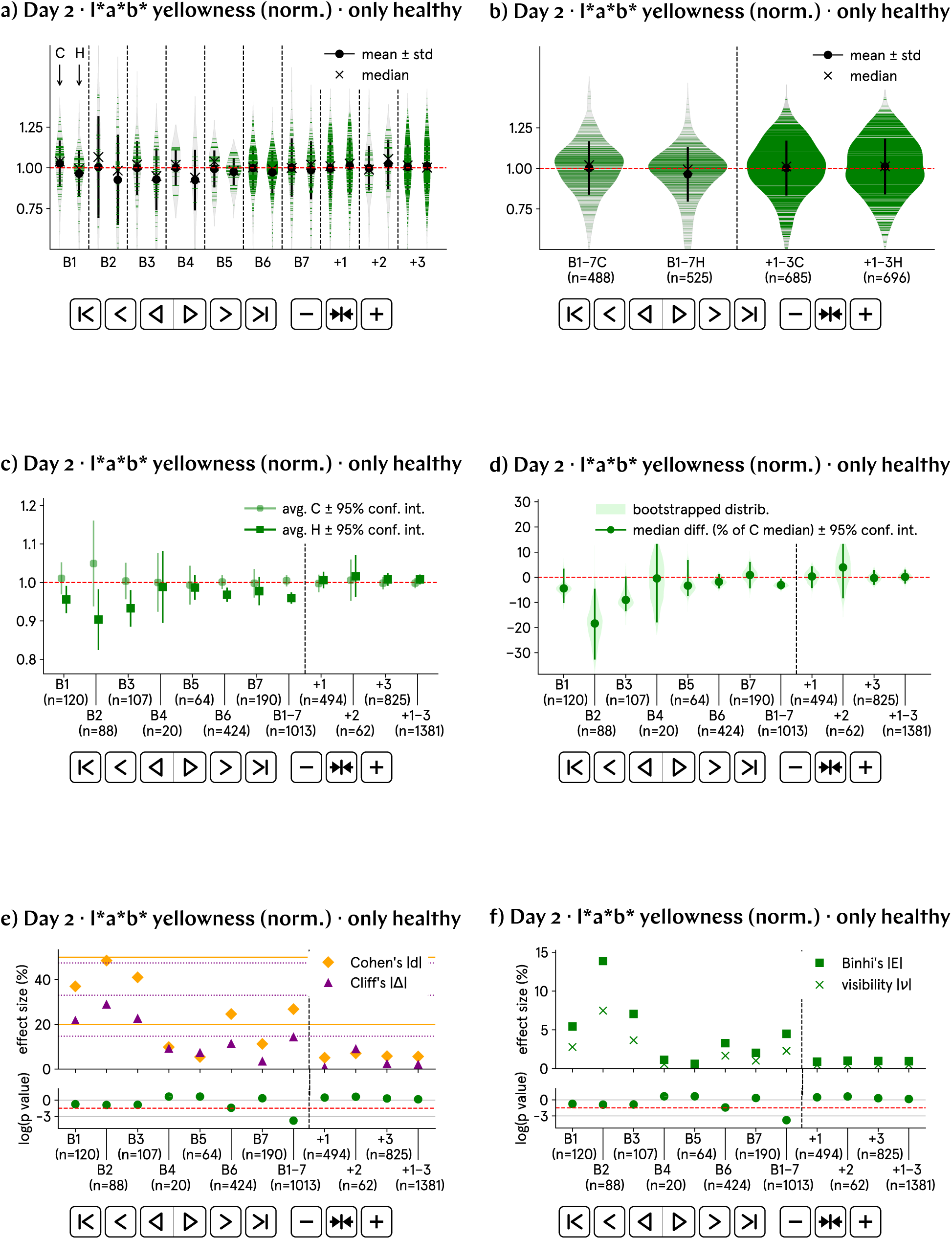
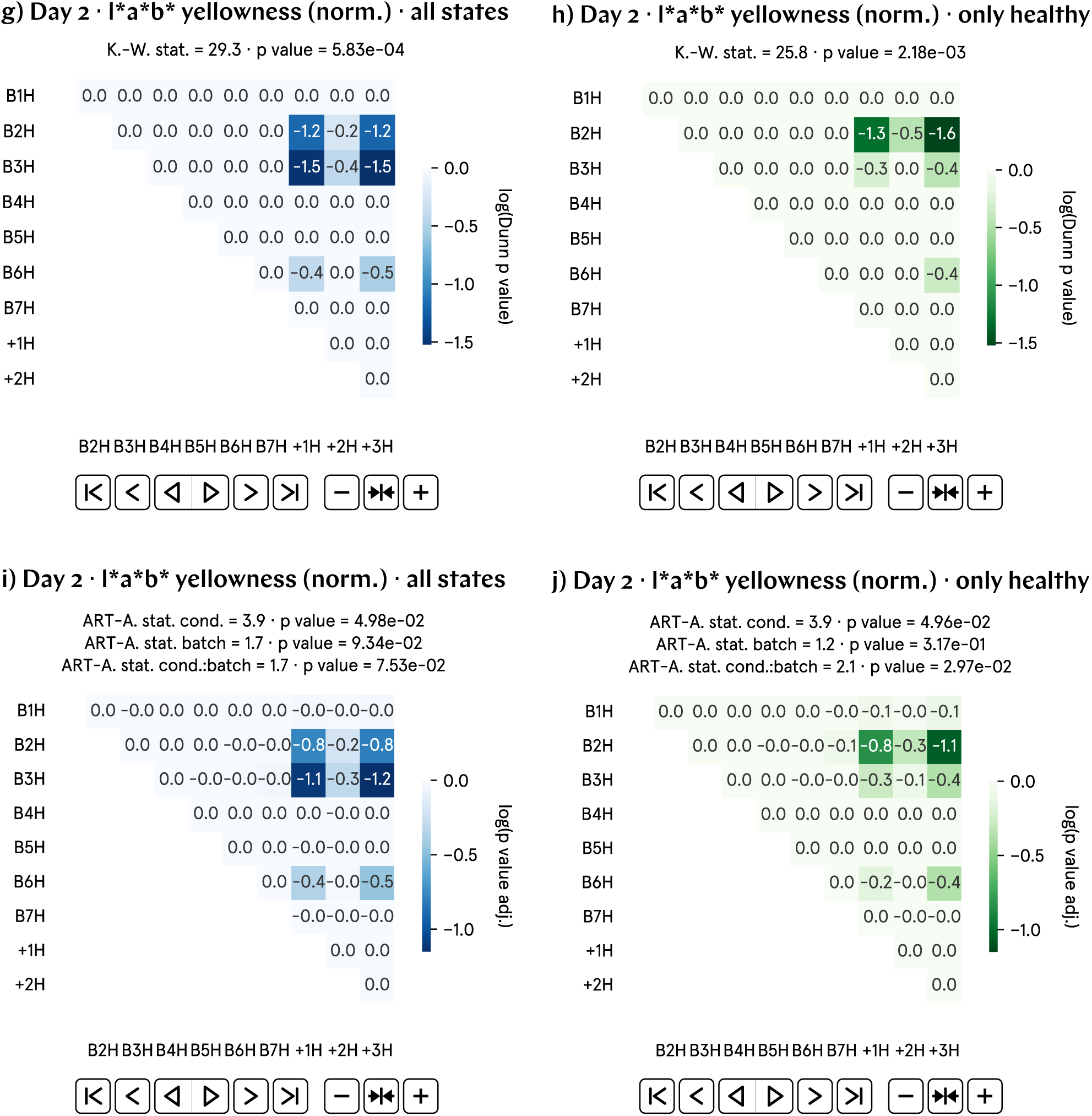
At day 2 p.f., the L*a*b* yellowness of hypomagnetic embryos is approximately 5% lower than that of the control population. a) and b) Normalized L*a*b* yellowness by batch and aggregate data; B1–7 represent experimental runs, and +1–3 correspond to positive control runs, in which an artificial magnetic field inside the hypomagnetic chamber simulates Earth’s. c) and d) Analysis of the population averages and medians further support the observed 5% lower L*a*b* yellowness. e) and f) Effect size metrics, namely Cohen’s *d* and Cliff’s Δ, and Binhi’s *E* and the visibility *ν*, confirm a small to medium effect, consistently reflecting the 5% level difference. g) through j) Most statistically significant differences are found in the positive control tadpole populations raised in the hypomagnetic chamber, as expected.

### SI28. Day 2 L*^*^*a*^*^*b*^*^* yellowness results

**Fig. SI20:**
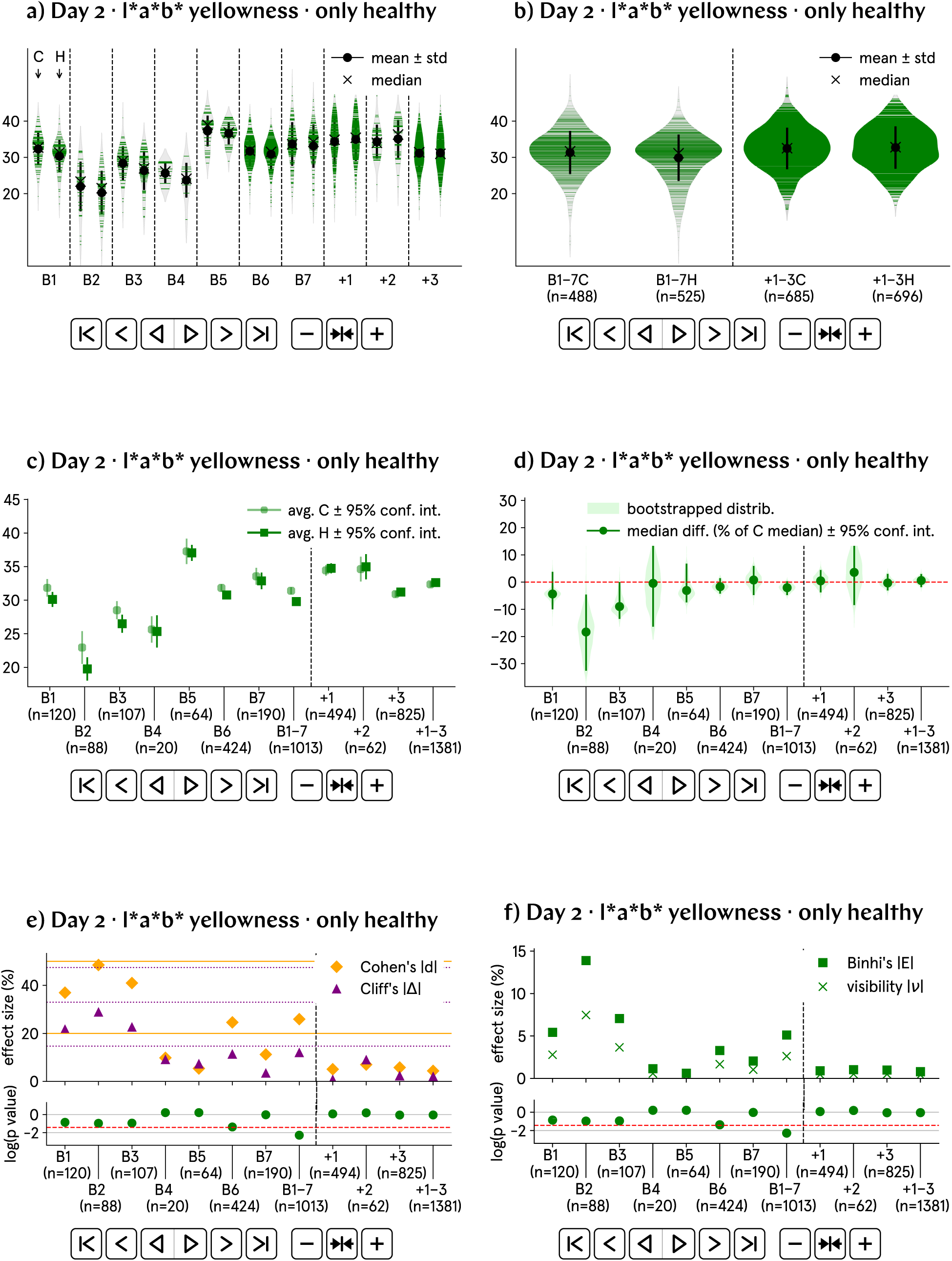
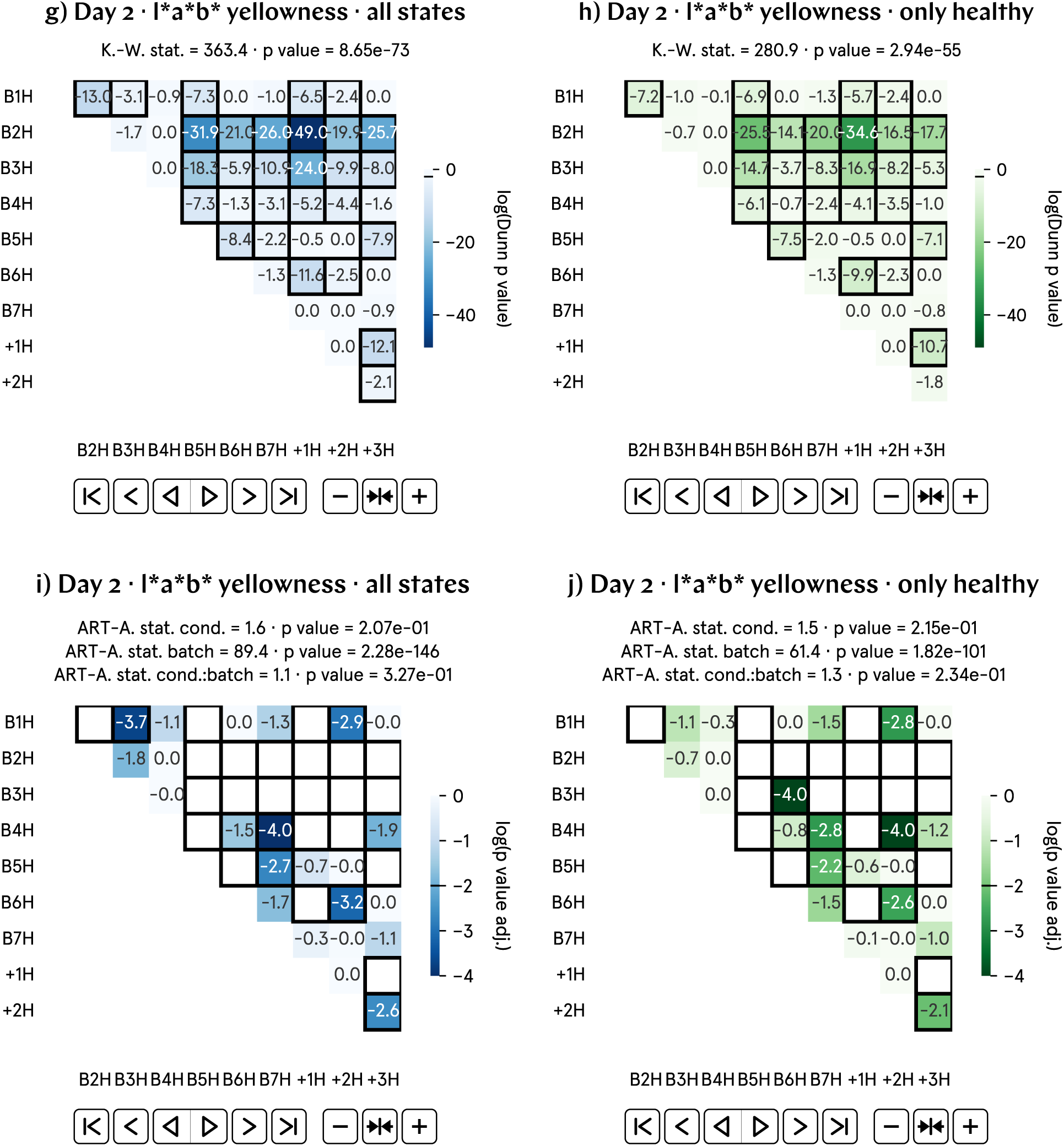
At day 2 p.f., the L*a*b* yellowness of hypomagnetic embryos is approximately 5% lower than that of the control population. a) and b) Non-normalized L*a*b* yellowness by batch and aggregate data; B1–7 represent experimental runs, and +1–3 correspond to positive control runs, in which an artificial magnetic field inside the hypomagnetic chamber simulates Earth’s. c) and d) Analysis of the population averages and medians further support the observed 5% lower L*a*b* yellowness. e) and f) Effect size metrics, namely Cohen’s *d* and Cliff’s Δ, and Binhi’s *E* and the visibility *ν*, confirm a small to medium effect, consistently reflecting the 5% level difference. g) through j) Given the substantial variation in L*a*b* yellowness across batches, these plots may not provide an accurate representation of our data.

### SI29. Day 2 RGB yellowness (norm.) results

**Fig. SI21:**
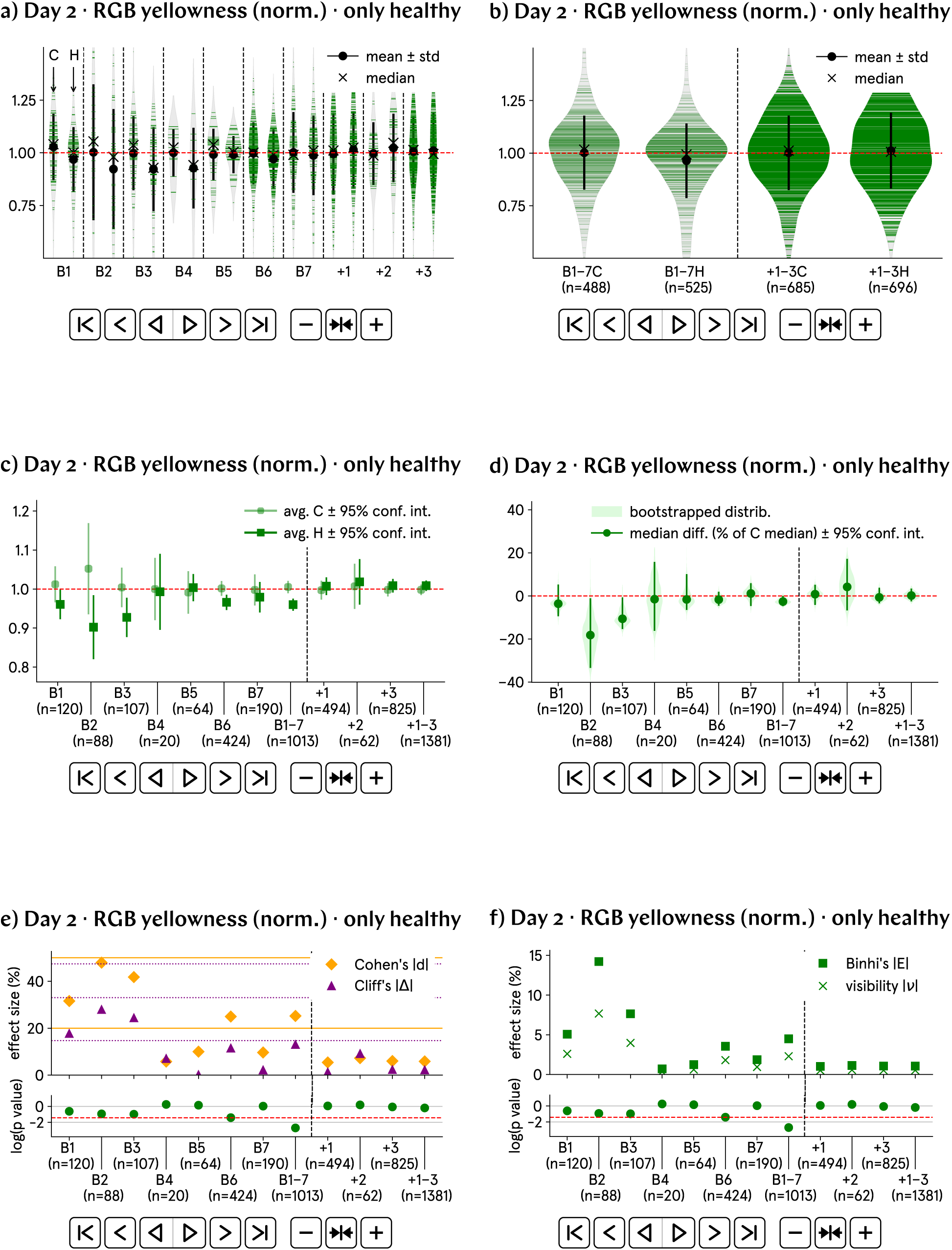
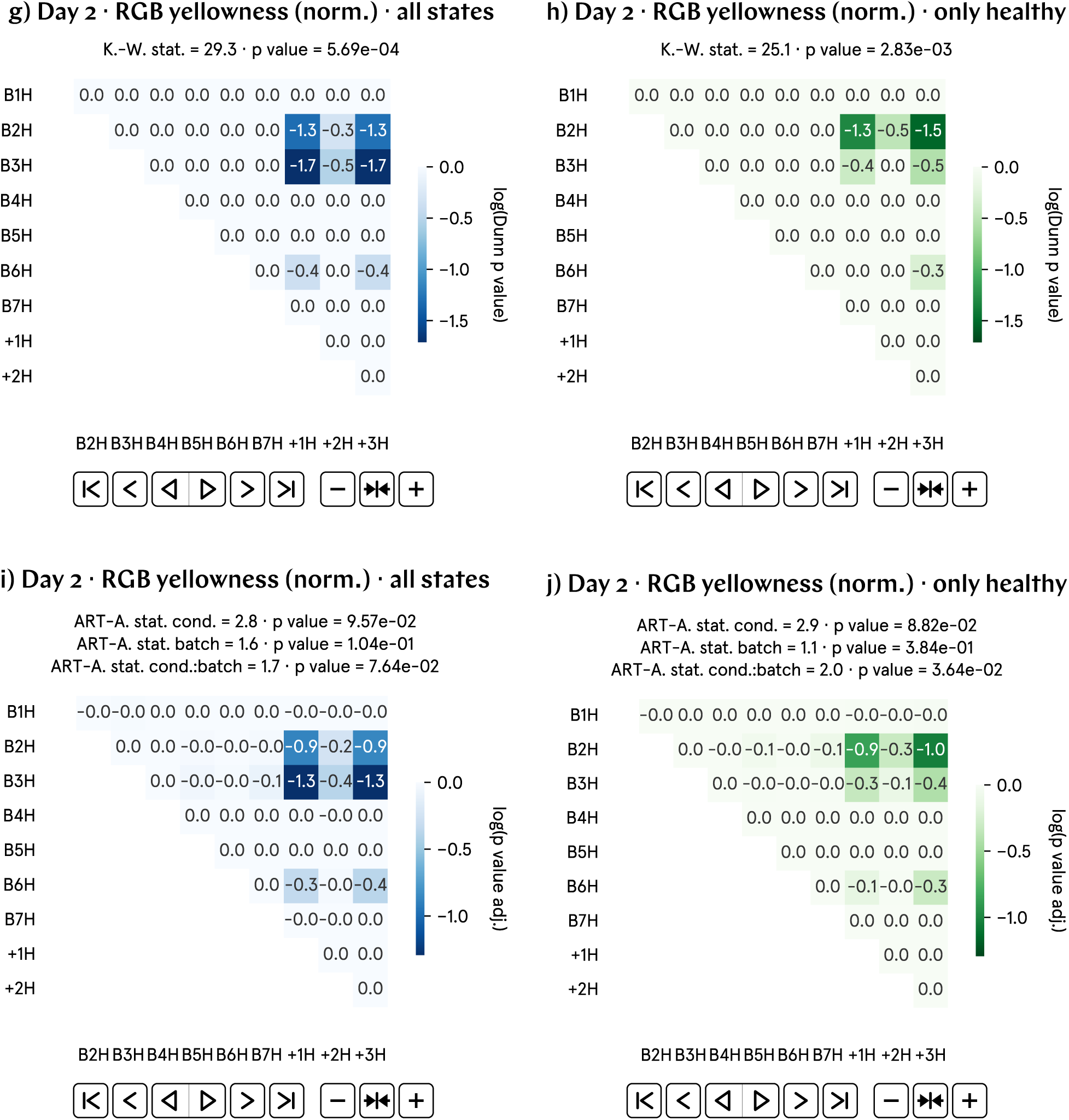
At day 2 p.f., the L*a*b* yellowness of hypomagnetic embryos is approximately 5% lower than that of the control population. a) and b) Non-normalized L*a*b* yellowness by batch and aggregate data; B1–7 represent experimental runs, and +1–3 correspond to positive control runs, in which an artificial magnetic field inside the hypomagnetic chamber simulates Earth’s. c) and d) Analysis of the population averages and medians further support the observed 5% lower L*a*b* yellowness. e) and f) Effect size metrics, namely Cohen’s *d* and Cliff’s Δ, and Binhi’s *E* and the visibility *ν*, confirm a small to medium effect, consistently reflecting the 5% level difference. g) through j) Given the substantial variation in L*a*b* yellowness across batches, these plots may not provide an accurate representation of our data.

### SI30. Day 2 RGB yellowness results

**Fig. SI22:**
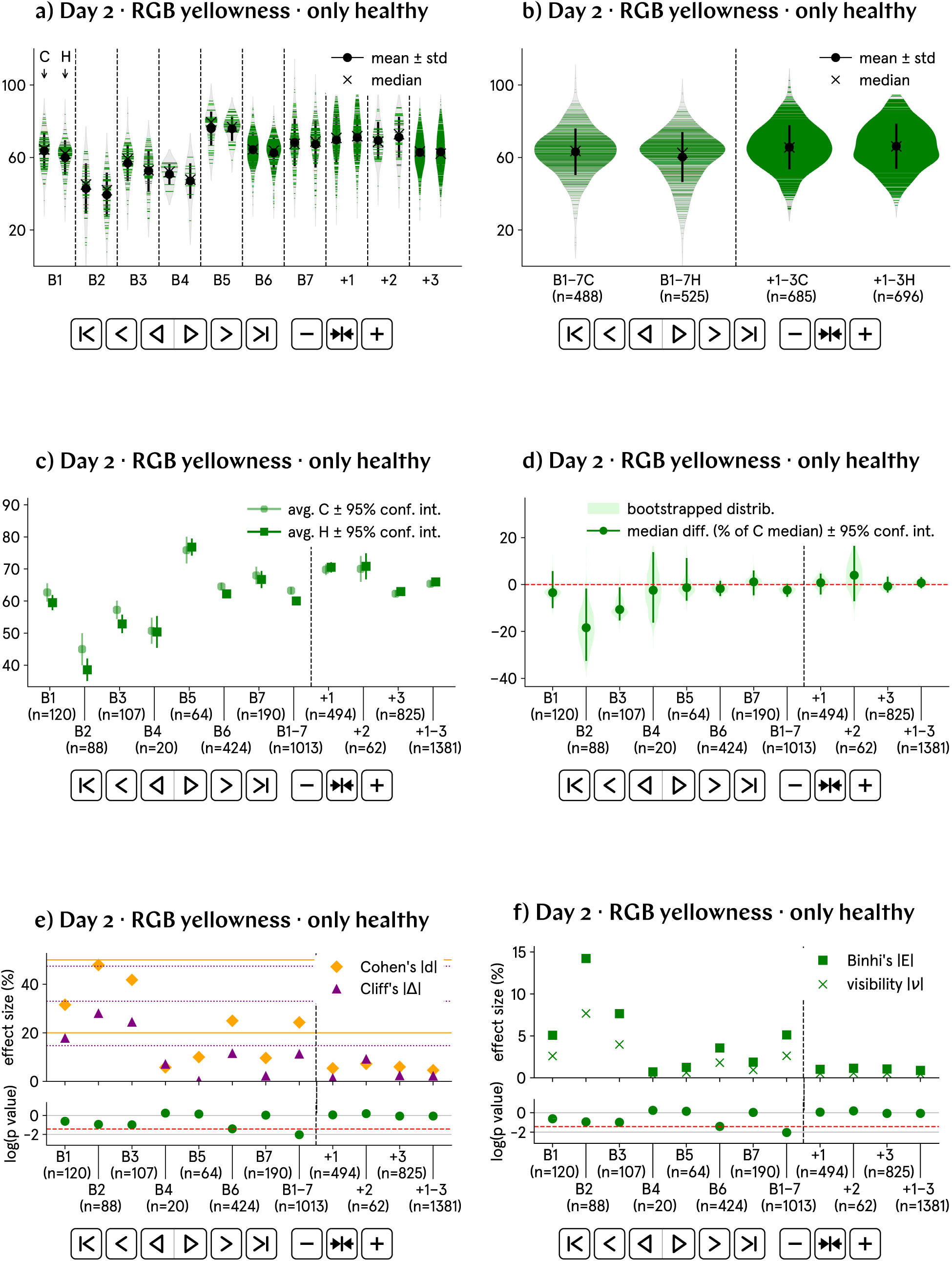
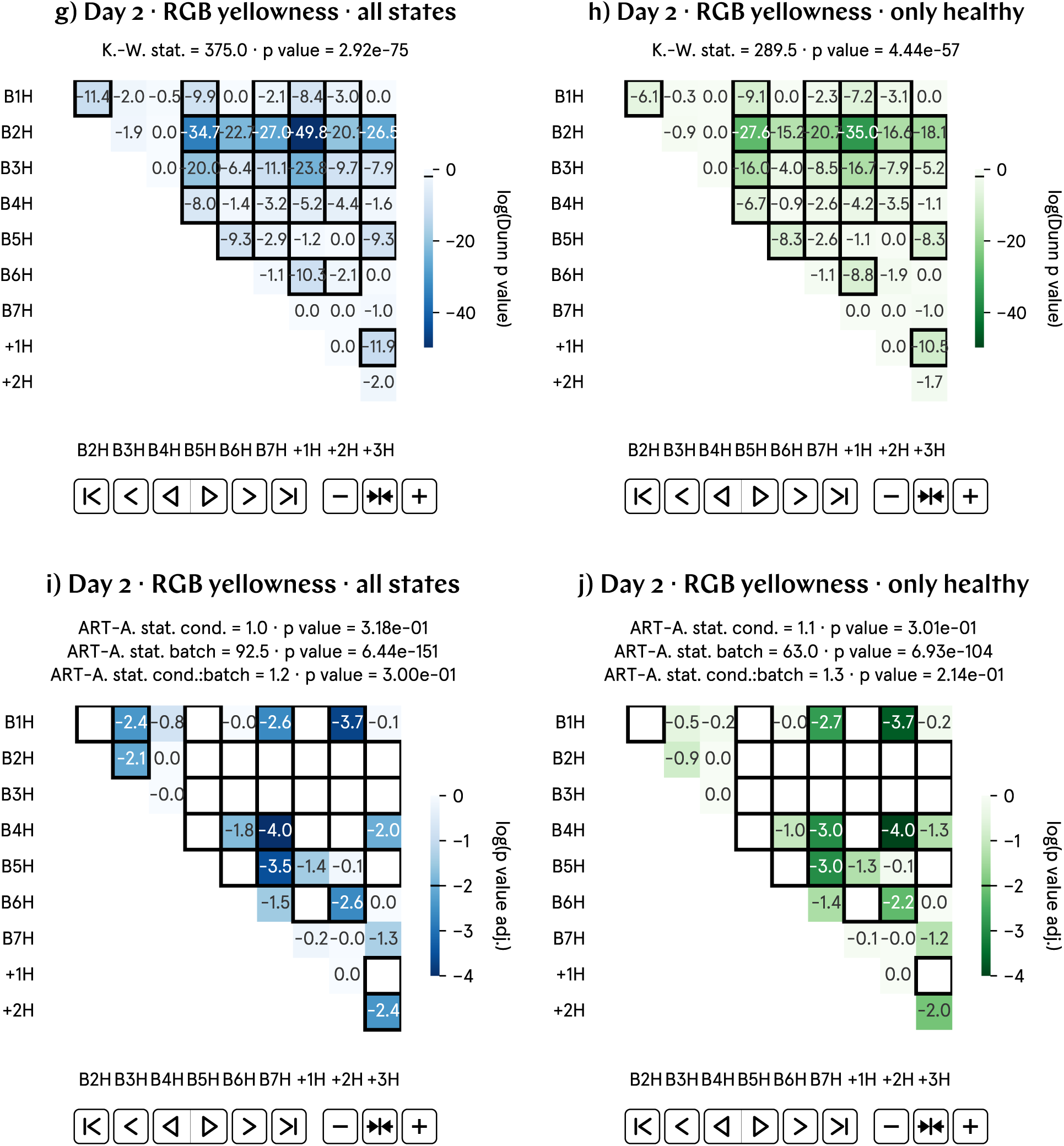
At day 2 p.f., the RGB yellowness of hypomagnetic embryos is approximately 5% lower than that of the control population. a) and b) Non-normalized RGB yellowness by batch and aggregate data; B1–7 represent experimental runs, and +1–3 correspond to positive control runs, in which an artificial magnetic field inside the hypomagnetic chamber simulates Earth’s. c) and d) Analysis of the population averages and medians further support the observed 5% lower RGB yellowness. e) and f) Effect size metrics, namely Cohen’s *d* and Cliff’s Δ, and Binhi’s *E* and the visibility *ν*, confirm a small to medium effect, consistently reflecting the 5% level difference. g) through j) Given the substantial variation in RGB yellowness across batches, these plots may not provide an accurate representation of our data.

### SI31. Day 2 Yellowness Index (norm.) results

**Fig. SI23:**
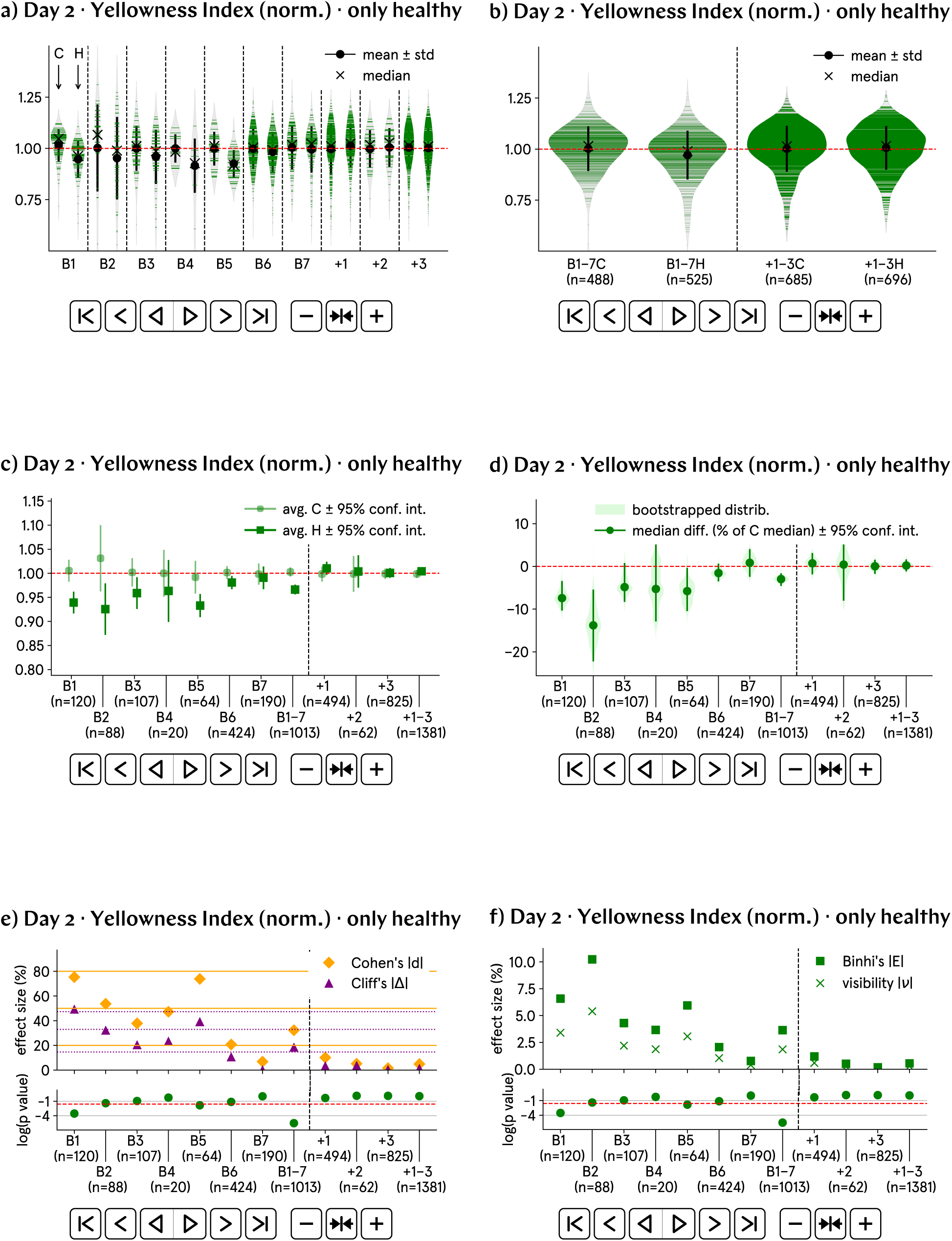
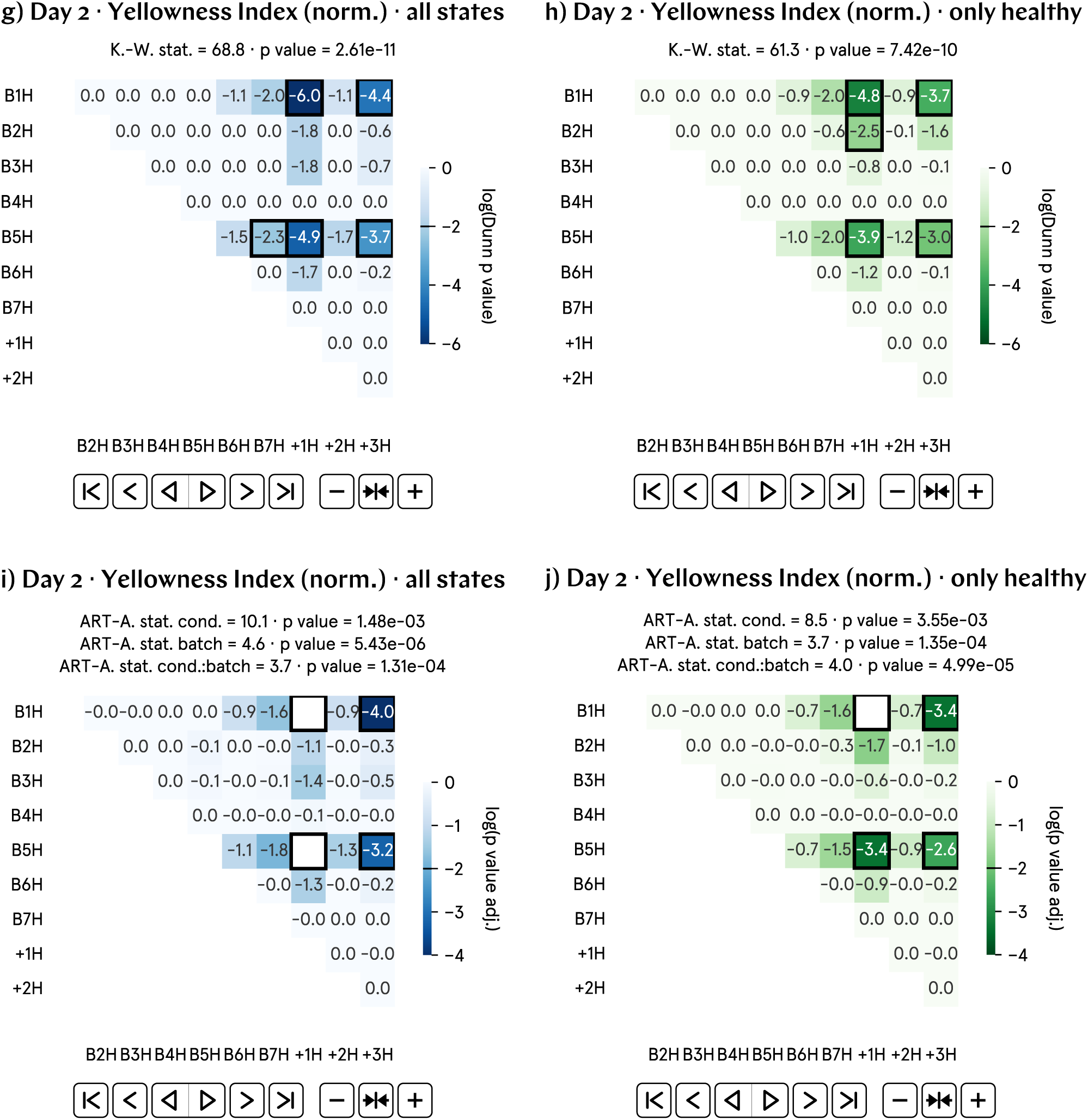
At day 2 p.f., the Yellowness Index of hypomagnetic embryos is approximately 5% lower than that of the control population. a) and b) Normalized Yellowness Index by batch and aggregate data; B1–7 represent experimental runs, and +1–3 correspond to positive control runs, in which an artificial magnetic field inside the hypomagnetic chamber simulates Earth’s. c) and d) Analysis of the population averages and medians further support the observed 5% lower Yellowness Index. e) and f) Effect size metrics, namely Cohen’s *d* and Cliff’s Δ, and Binhi’s *E* and the visibility *ν*, confirm a small to medium effect, consistently reflecting the 5% level difference. g) through j) Some statistically significant differences are found in the positive control tadpole populations raised in the hypomagnetic chamber, as expected. Surprisingly, batches B1 and B5 are also anomalous.

### SI32. Day 2 Yellowness Index results

**Fig. SI24:**
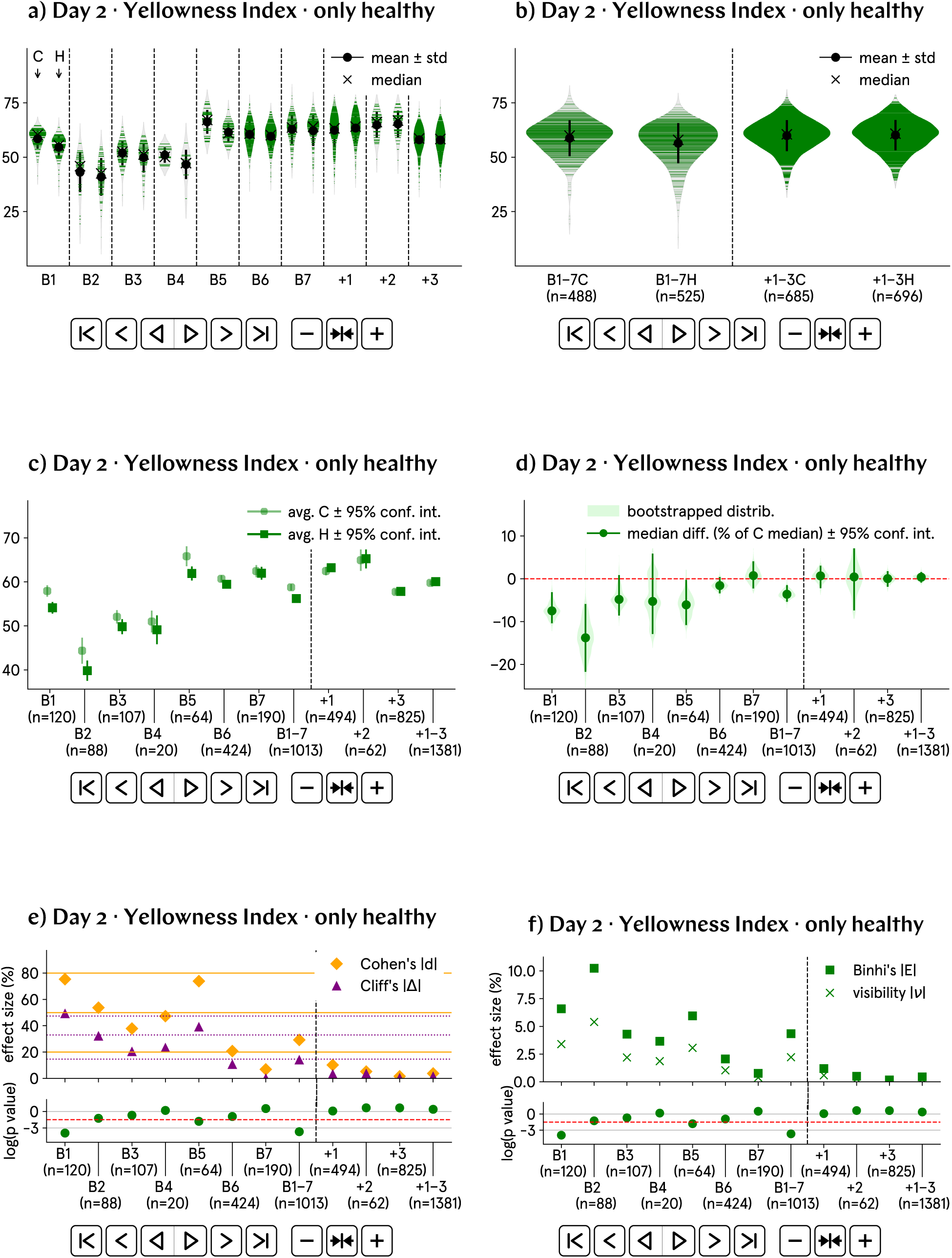
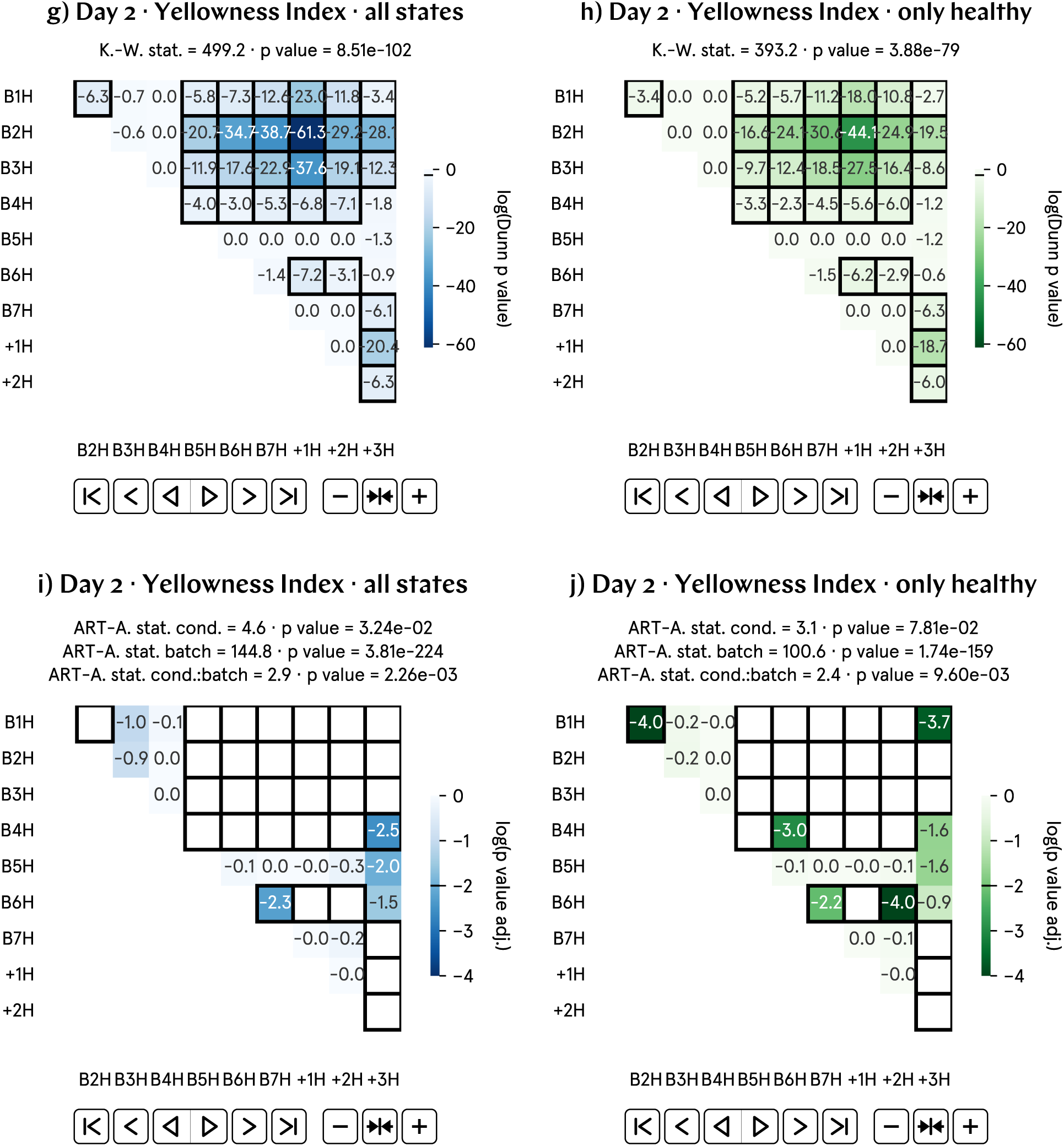
At day 2 p.f., the Yellowness Index of hypomagnetic embryos is approximately 5% lower than that of the control population. a) and b) Non-normalized Yellowness Index by batch and aggregate data; B1–7 represent experimental runs, and +1–3 correspond to positive control runs, in which an artificial magnetic field inside the hypomagnetic chamber simulates Earth’s. c) and d) Analysis of the population averages and medians further support the observed 5% lower Yellowness Index. e) and f) Effect size metrics, namely Cohen’s *d* and Cliff’s Δ, and Binhi’s *E* and the visibility *ν*, confirm a small to medium effect, consistently reflecting the 5% level difference. g) through j) Given the substantial variation in Yellowness Index across batches, these plots may not provide an accurate representation of our data.

### SI33. Day 2 HSV yellowness (norm.) results

**Fig. SI25:**
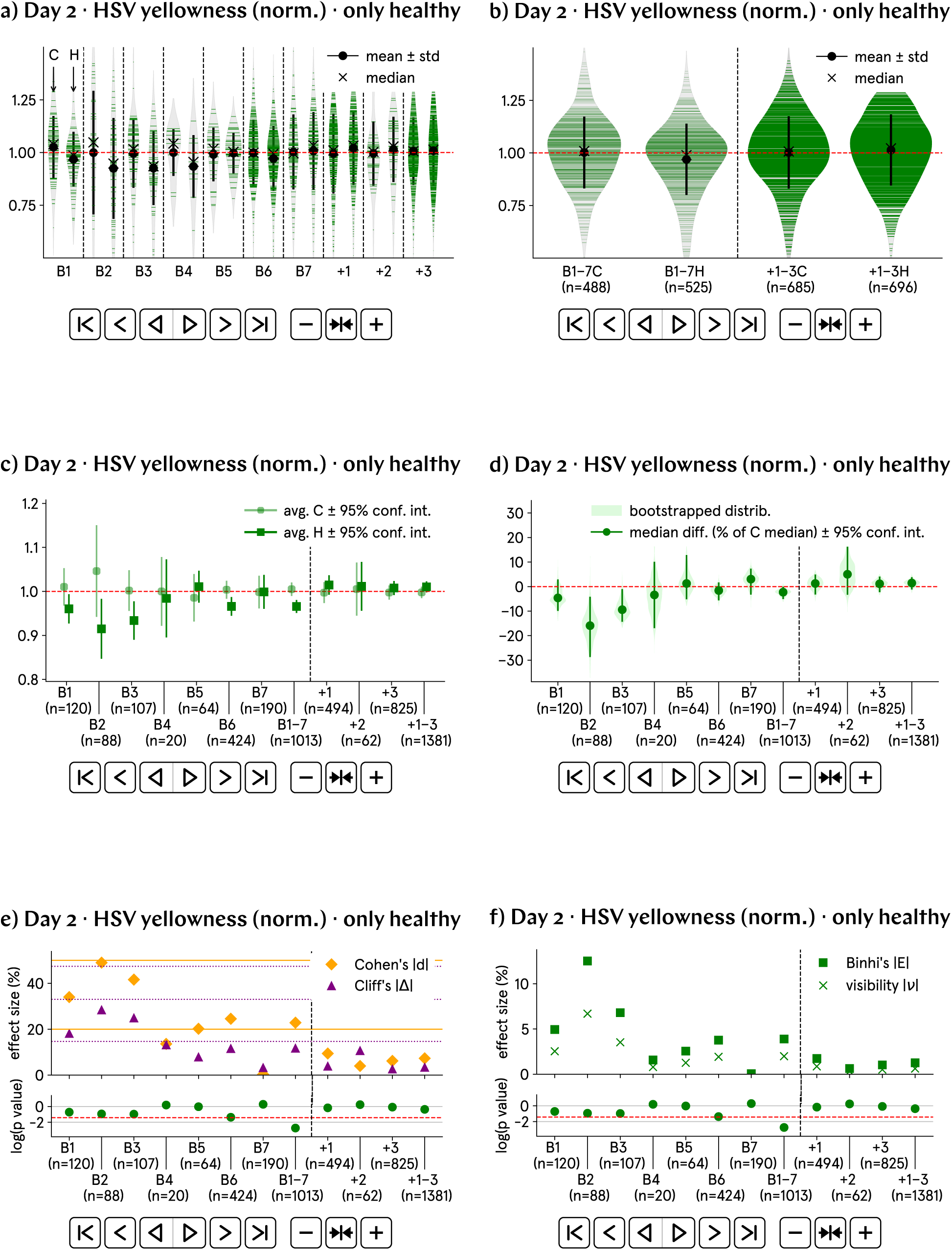
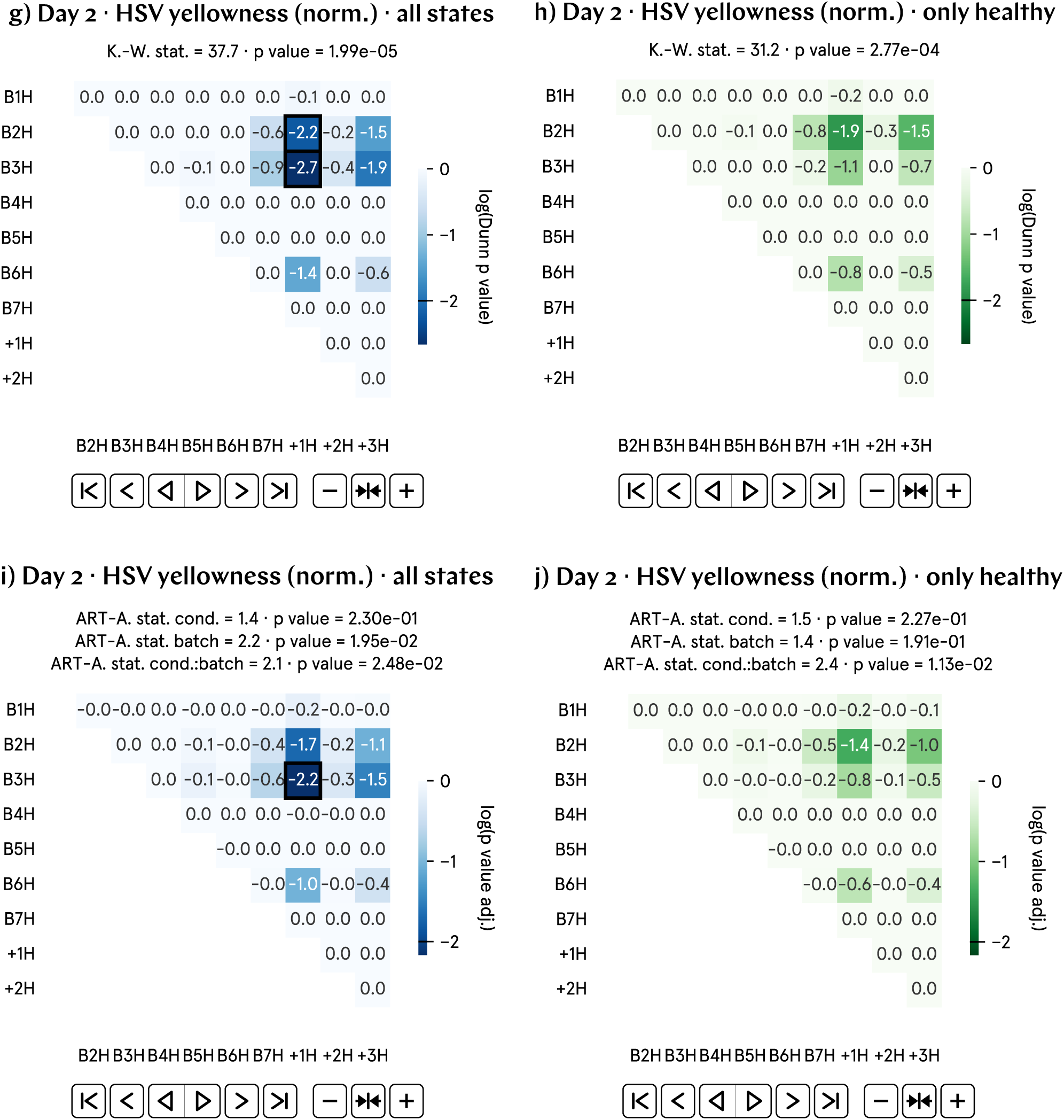
At day 2 p.f., the HSV yellowness of hypomagnetic embryos is approximately 5% lower than that of the control population. a) and b) Normalized HSV yellowness by batch and aggregate data; B1–7 represent experimental runs, and +1–3 correspond to positive control runs, in which an artificial magnetic field inside the hypomagnetic chamber simulates Earth’s. c) and d) Analysis of the population averages and medians further support the observed 5% lower HSV yellowness. e) and f) Effect size metrics, namely Cohen’s *d* and Cliff’s Δ, and Binhi’s *E* and the visibility *ν*, confirm a small to medium effect, consistently reflecting the 5% level difference. g) through j) Most statistically significant differences are found in the positive control tadpole populations raised in the hypomagnetic chamber, as expected. Surprisingly, batches B2 and B3 are also anomalous.

### SI34. Day 2 HSV yellowness results

**Fig. SI26:**
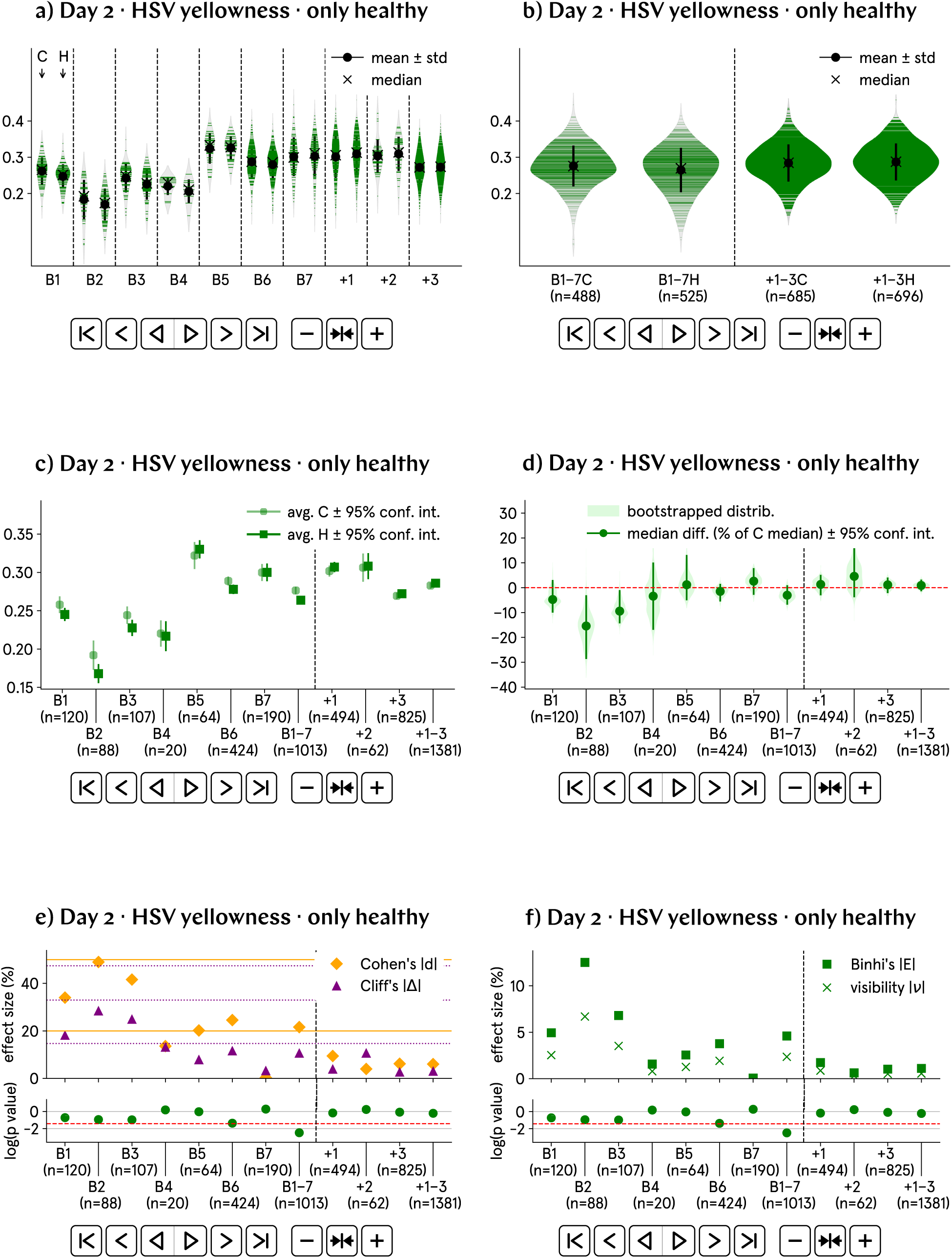
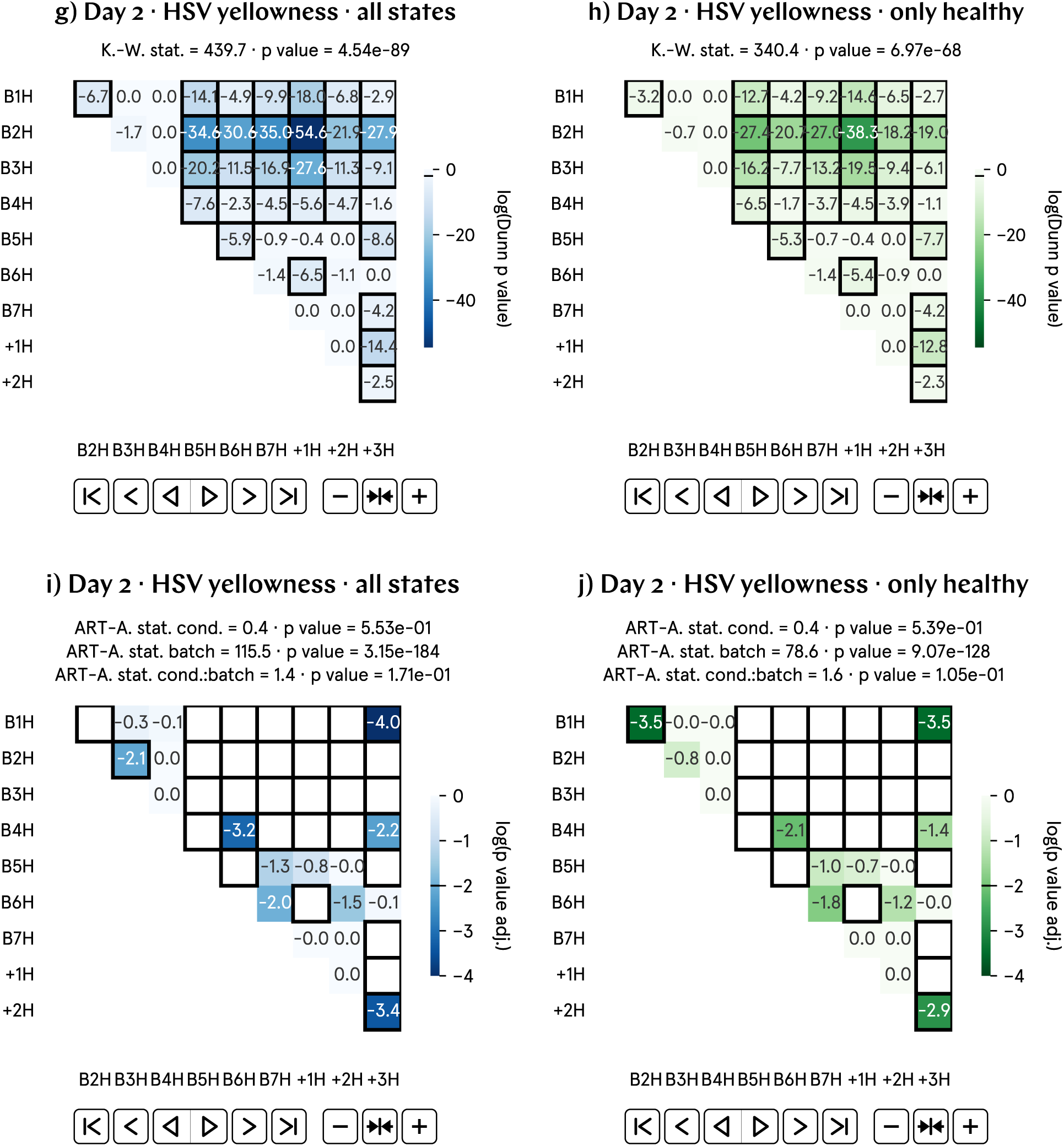
At day 2 p.f., the HSV yellowness of hypomagnetic embryos is approximately 5% lower than that of the control population. a) and b) Non-normalized HSV yellowness by batch and aggregate data; B1–7 represent experimental runs, and +1–3 correspond to positive control runs, in which an artificial magnetic field inside the hypomagnetic chamber simulates Earth’s. c) and d) Analysis of the population averages and medians further support the observed 5% lower HSV yellowness. e) and f) Effect size metrics, namely Cohen’s *d* and Cliff’s Δ, and Binhi’s *E* and the visibility *ν*, confirm a small to medium effect, consistently reflecting the 5% level difference. g) through j) Given the substantial variation in HSV yellowness across batches, these plots may not provide an accurate representation of our data.

### SI35. Day 3 morphological measures: major axis, perimeter, area, total curvature, solidity, elongation and roundness

For day 3 images, we continue to quantify the accelerated development of the tadpoles under hypomagnetic conditions.

The simplest and clearest measure is the length of the tadpoles’ major axis. Using python’s skimage.measure package, the major axis is calculated by fitting an ellipse to the largest region of the image and measuring its longest diameter. This axis represents the longest straight line through the region and is typically aligned with its orientation. Specifically, the major axis is the longest diameter of an ellipse that shares the same normalized second central moments as the region.

The second and third measures showing a differential development for hypomagnetic condition tadpoles is their perimeter and their area, also calculated with the python package skimage.measure.

The fourth measure used to quantify Day 3 images is the total image curvature. Using skimage.measure, the tadpole contour is extracted, and curvature is approximated through numerical differentiation. First, we compute the first derivative of the contour points, representing consecutive changes, followed by the second derivative, which captures the rate of change. The Euclidean norm of the second derivatives gives the curvature at each point. We report the absolute sum of these curvatures, reflecting the overall bending or complexity of the contour, with higher values indicating more pronounced or frequent bending. The use of the absolute values ensures that both concave and convex bends contribute positively to the total.

Day 3 images across different batches have similar, though not identical, magnifications. As a result, to ensure comparability across batches, we only present the normalized results for the major axis length (Supplementary Fig. SI28), perimeter (Supplementary Fig. SI29), area (Supplementary Fig. SI30), and total curvature (Supplementary Fig. SI31).

The fifth, sixth and seventh measures have been used for day 1 images and were discussed in Supplementary Section SI17. They are: solidity, elongation and roundness. Roundness on day 3 is particularly significant as it highlights the relationship between tadpole area and perimeter. With an approximate 7.5% increase in perimeter and a 5% increase in area, the roundness is expected to decrease by roughly 9%, aligning closely with our numerically calculated values. Our analysis for these measures, in normalized form (not normalized form, with stage predictions), are found respectively in Supplementary Fig. SI32 (Supplementary Fig. SI33), Supplementary Fig. SI34 (Supplementary Fig. SI35) and Supplementary Fig. SI36 (Supplementary Fig. SI37).

A sample code output for day 3 images (of batch B4, in this example) is found in Supplementary Fig. SI27, with each tadpole being annotated with its programmatically determined major and minor axes.

**Fig. SI27:**
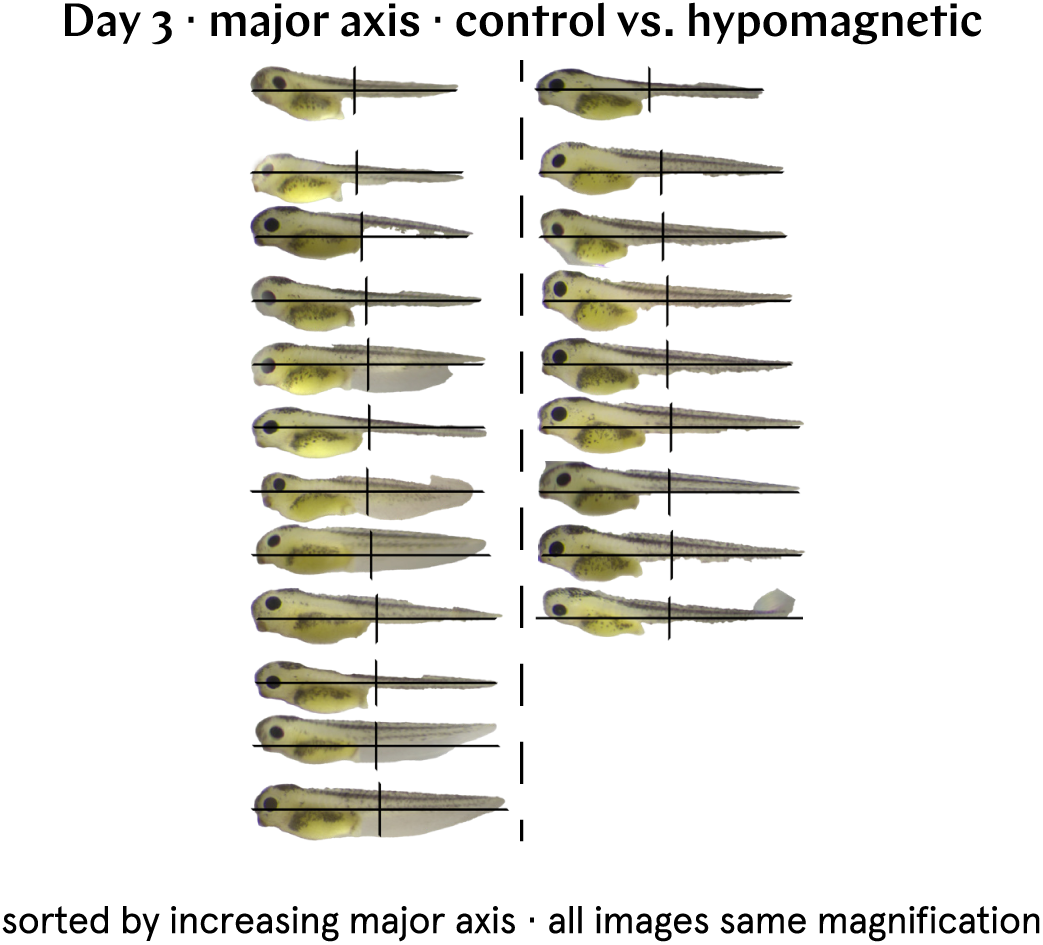
Sample code output for day 3 images. Day 3 images are segmented and subsequently analyzed for major axis length and other morphological properties. Here, day 3 images from batch B4 are arranged in order of increasing major axis length, with each tadpole displaying overlays of both major and minor axes.

### SI36. Day 3 major axis (norm.) results

**Fig. SI28:**
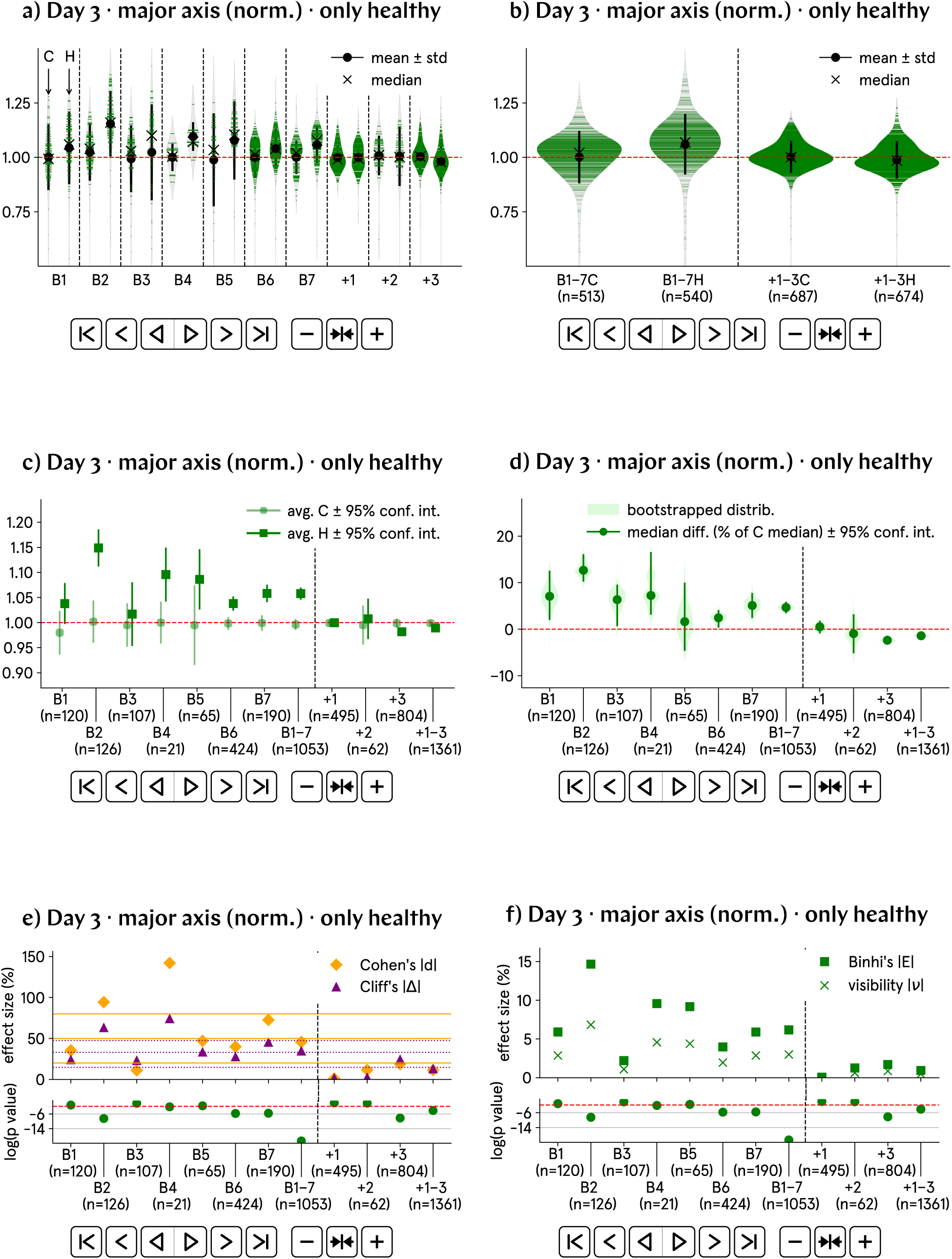
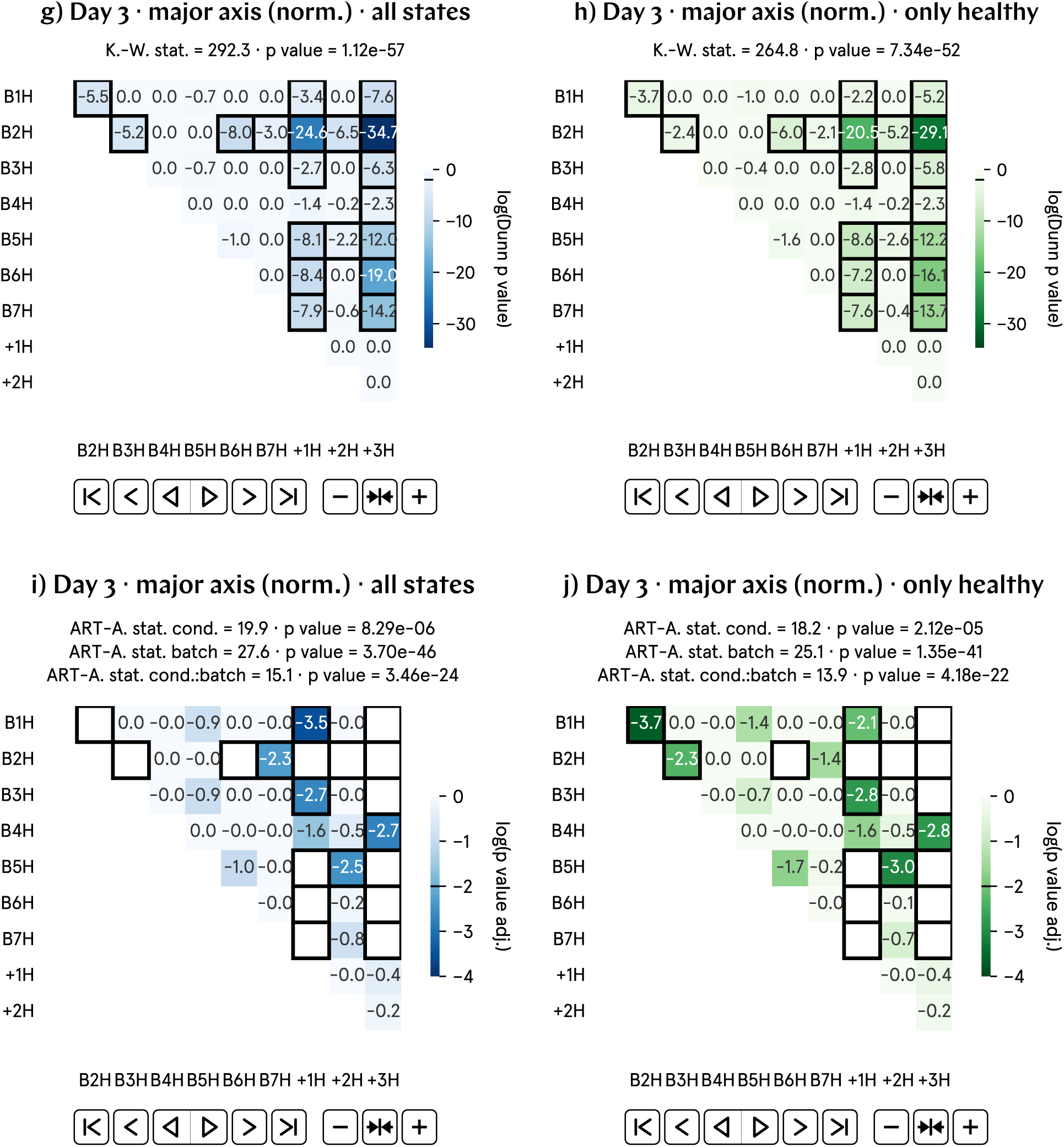
At day 3 p.f., the major axis of hypomagnetic embryos is approximately 7.5% higher than that of the control population. a) and b) Normalized major axis by batch and aggregate data; B1–7 represent experimental runs, and +1–3 correspond to positive control runs, in which an artificial magnetic field inside the hypomagnetic chamber simulates Earth’s. c) and d) Analysis of the population averages and medians further support the observed 7.5% higher major axis. e) and f) Effect size metrics, namely Cohen’s *d* and Cliff’s Δ, and Binhi’s *E* and the visibility *ν*, confirm a medium effect, consistently reflecting the 7.5% level difference. g) through j) Most statistically significant differences are found in the positive control tadpole populations raised in the hypomagnetic chamber, as expected. Batch B2 is also anomalous.

### SI37. Day 3 perimeter (norm.) results

**Fig. SI29:**
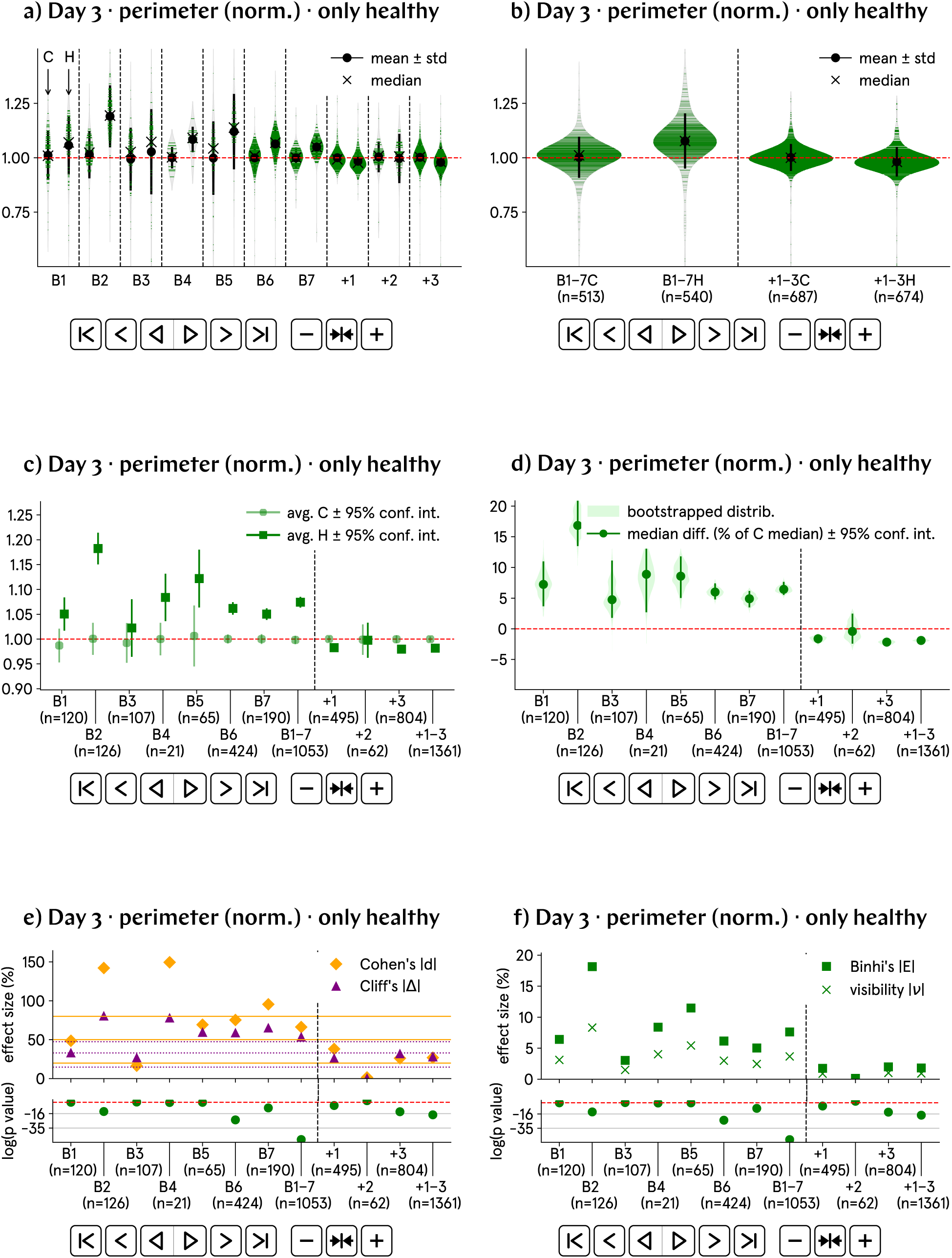
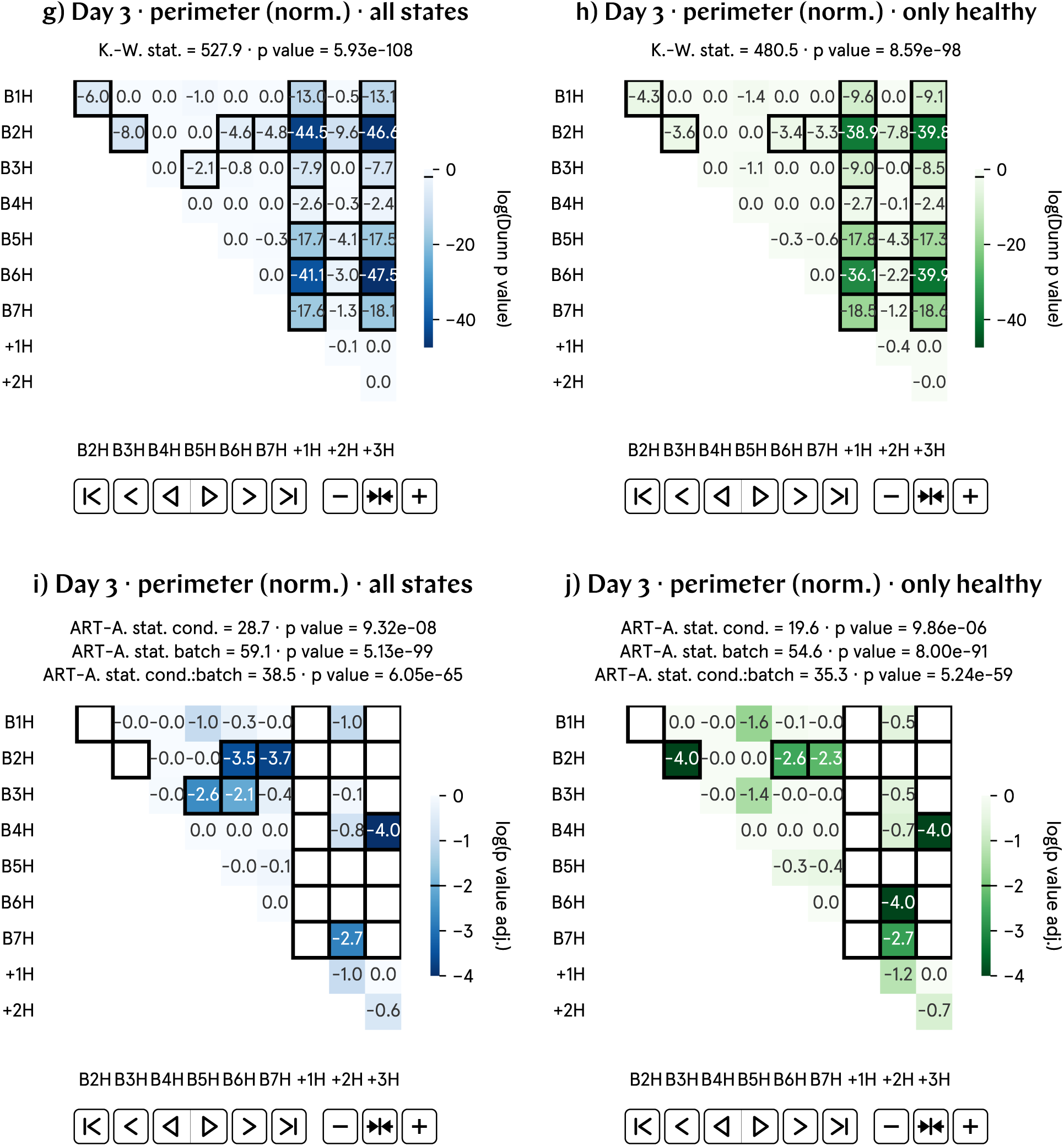
At day 3 p.f., the perimeter of hypomagnetic embryos is approximately 7.5% higher than that of the control population. a) and b) Normalized perimeter by batch and aggregate data; B1–7 represent experimental runs, and +1–3 correspond to positive control runs, in which an artificial magnetic field inside the hypomagnetic chamber simulates Earth’s. c) and d) Analysis of the population averages and medians further support the observed 7.5% higher perimeter. e) and f) Effect size metrics, namely Cohen’s *d* and Cliff’s Δ, and Binhi’s *E* and the visibility *ν*, confirm a medium effect, consistently reflecting the 7.5% level difference. g) through j) Most statistically significant differences are found in the positive control tadpole populations raised in the hypomagnetic chamber, as expected. Batch B2 is also anomalous.

### SI38. Day 3 area (norm.) results

**Fig. SI30:**
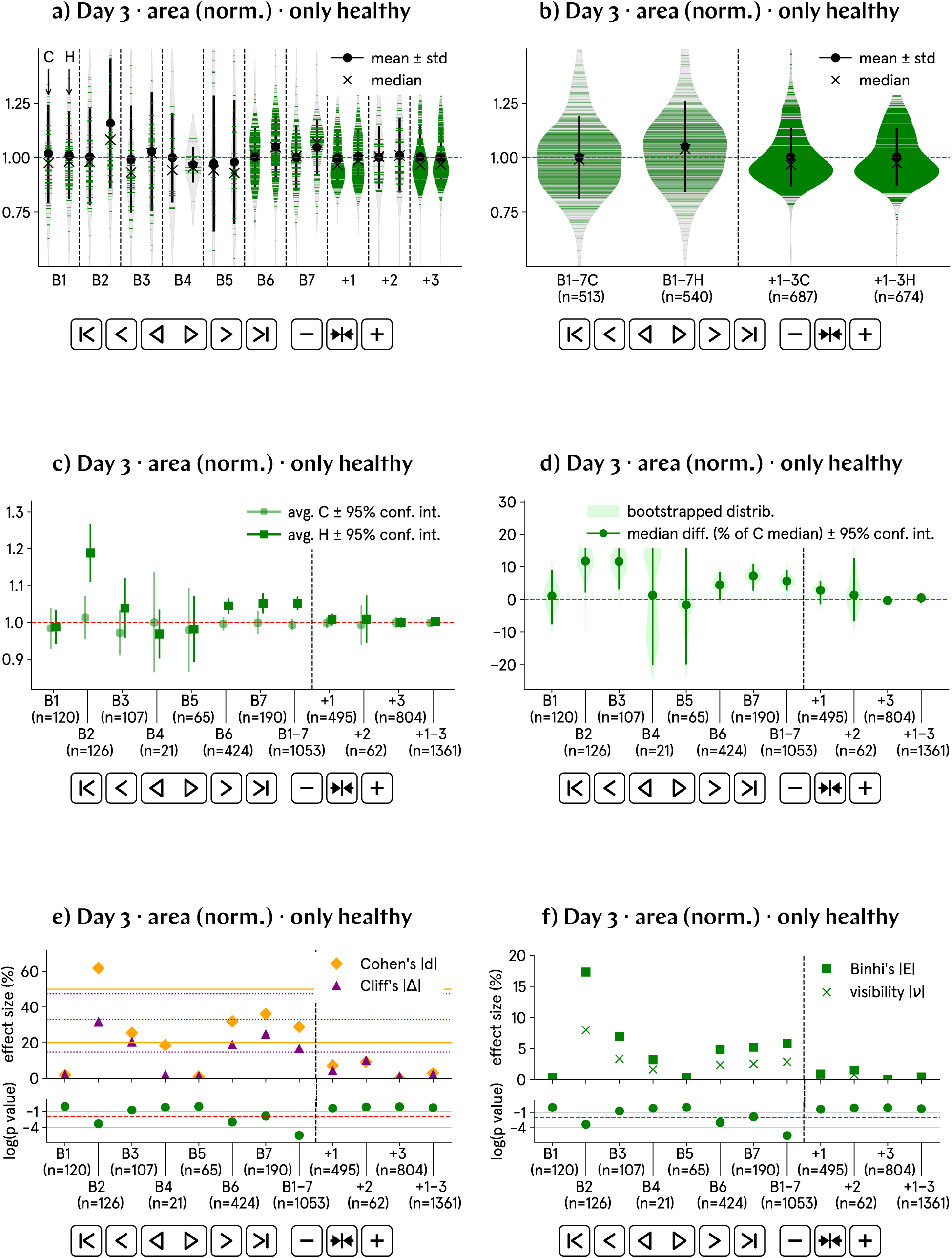
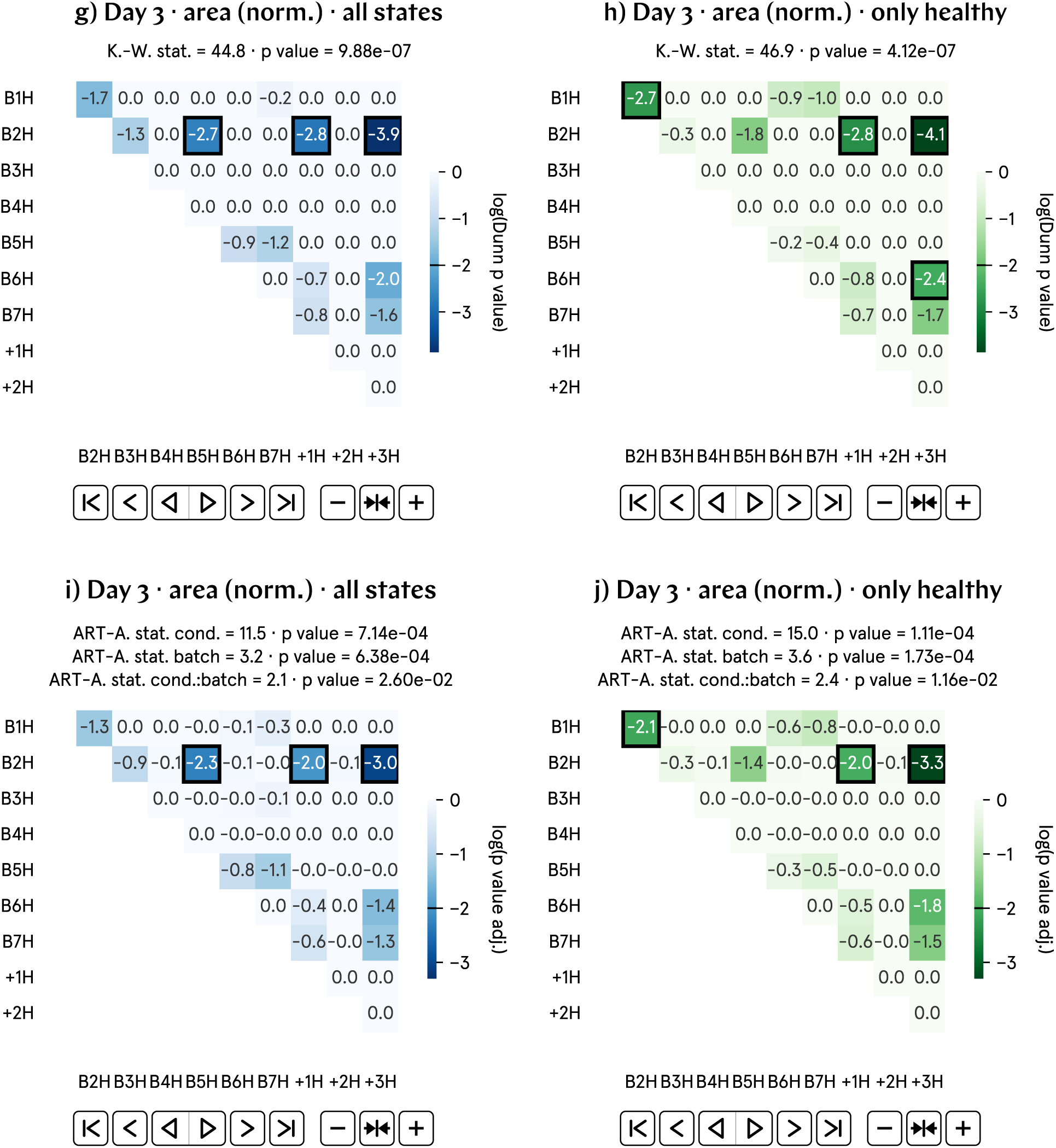
At day 3 p.f., the area of hypomagnetic embryos is approximately 5% higher than that of the control population. a) and b) Normalized area by batch and aggregate data; B1–7 represent experimental runs, and +1–3 correspond to positive control runs, in which an artificial magnetic field inside the hypomagnetic chamber simulates Earth’s. c) and d) Analysis of the population averages and medians further support the observed 5% higher area. e) and f) Effect size metrics, namely Cohen’s *d* and Cliff’s Δ, and Binhi’s *E* and the visibility *ν*, confirm a medium effect, consistently reflecting the 5% level difference. g) through j) Most statistically significant differences are found in the positive control tadpole populations raised in the hypomagnetic chamber, as expected. Batch B2 is also anomalous.

### SI39. Day 3 curvature (norm.) results

**Fig. SI31:**
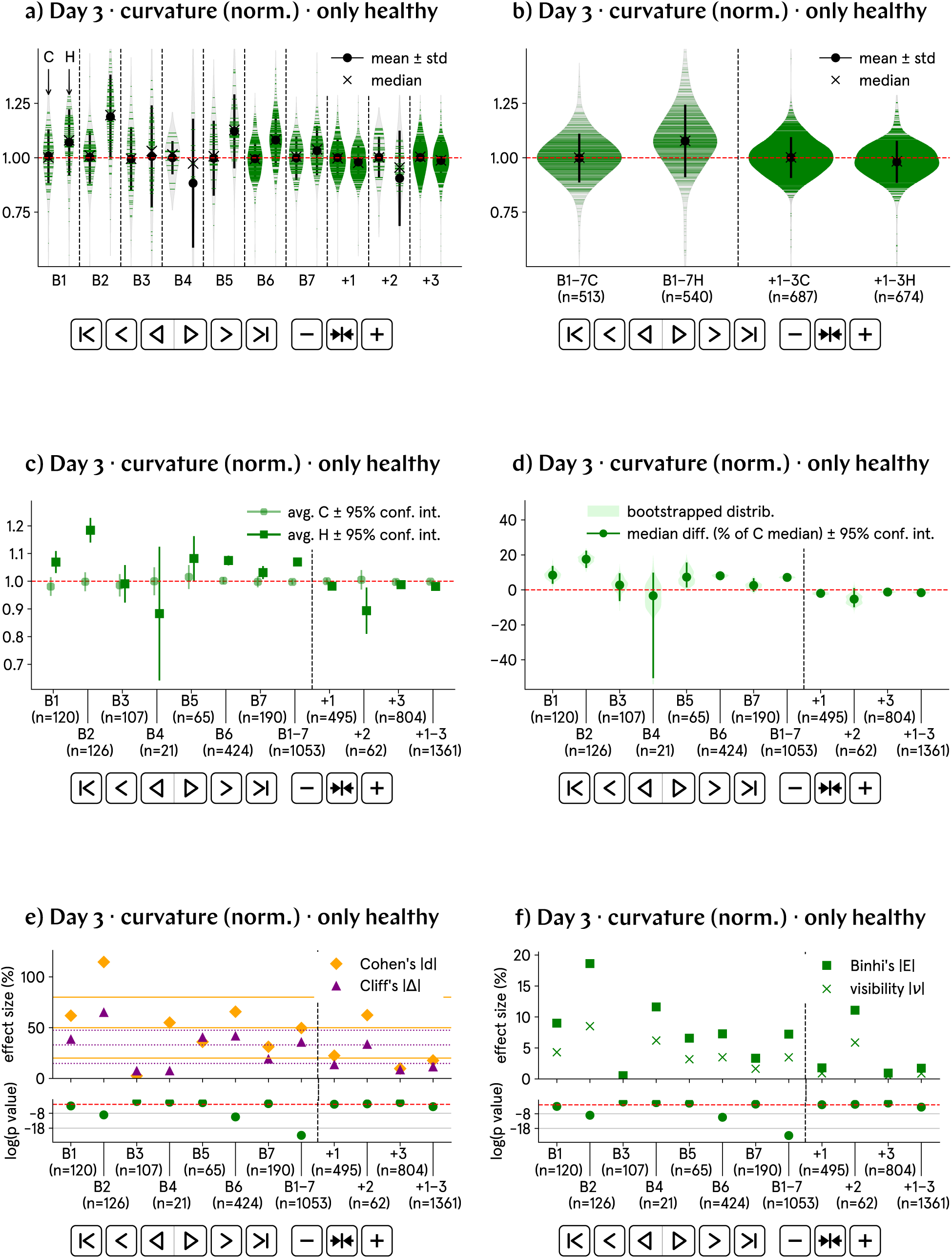
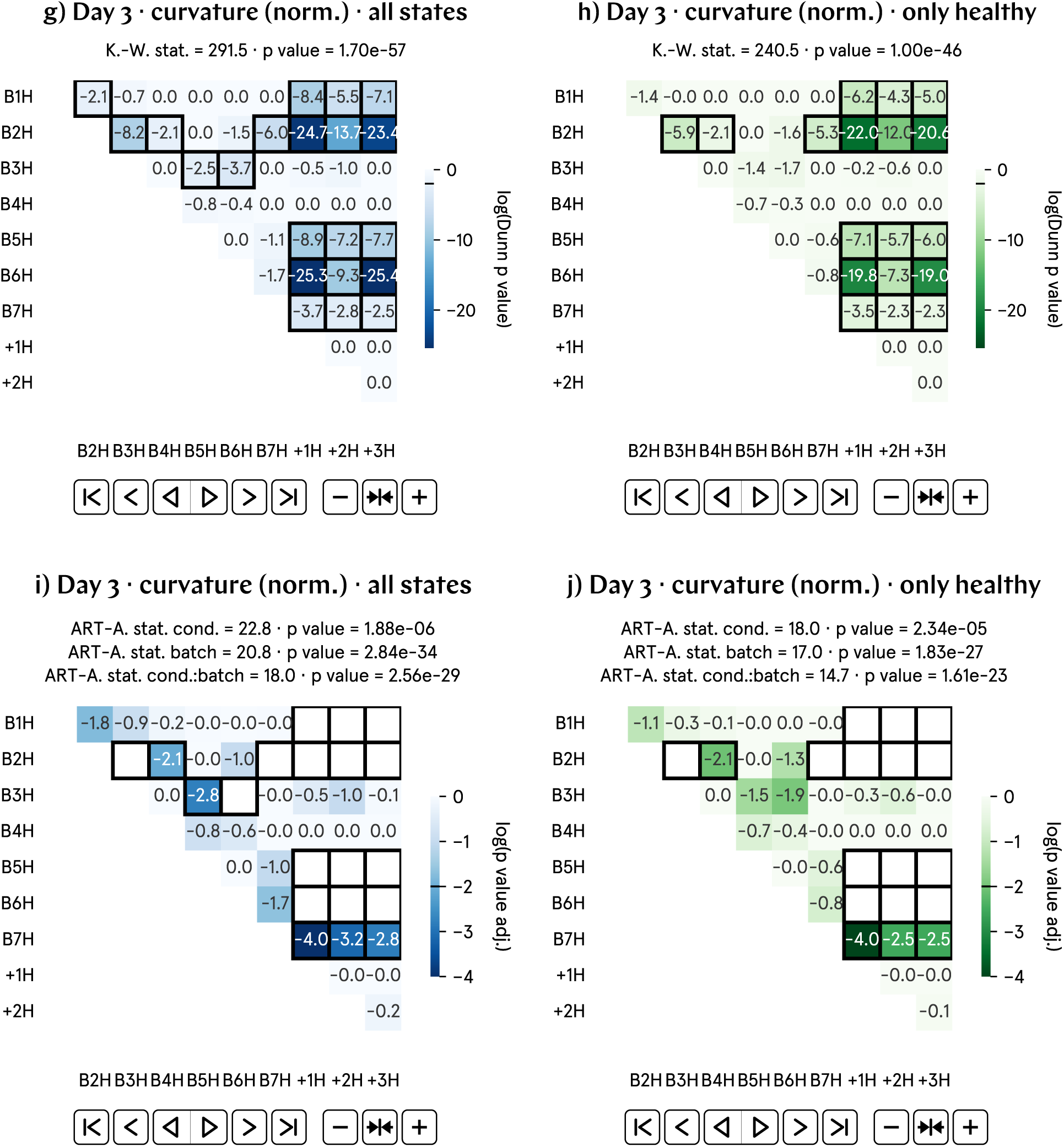
At day 3 p.f., the total curvature of hypomagnetic embryos is approximately 7.5% higher than that of the control population. a) and b) Normalized total curvature by batch and aggregate data; B1–7 represent experimental runs, and +1–3 correspond to positive control runs, in which an artificial magnetic field inside the hypomagnetic chamber simulates Earth’s. c) and d) Analysis of the population averages and medians further support the observed 7.5% higher total curvature. e) and f) Effect size metrics, namely Cohen’s *d* and Cliff’s Δ, and Binhi’s *E* and the visibility *ν*, confirm a medium effect, consistently reflecting the 7.5% level difference. g) through j) Most statistically significant differences are found in the positive control tadpole populations raised in the hypomagnetic chamber, as expected. Batch B2 is also anomalous.

### SI40. Day 3 solidity (norm.) results

**Fig. SI32:**
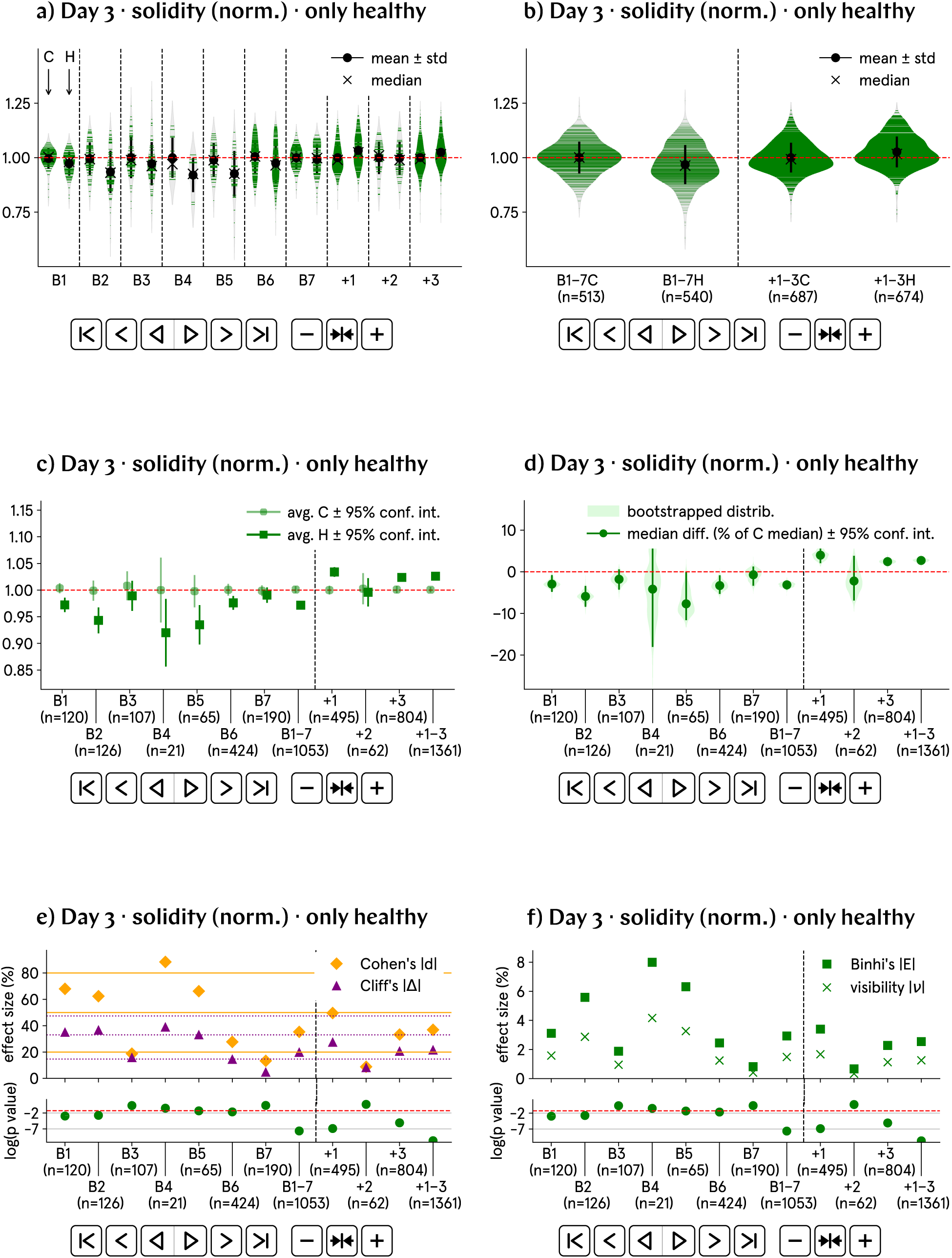
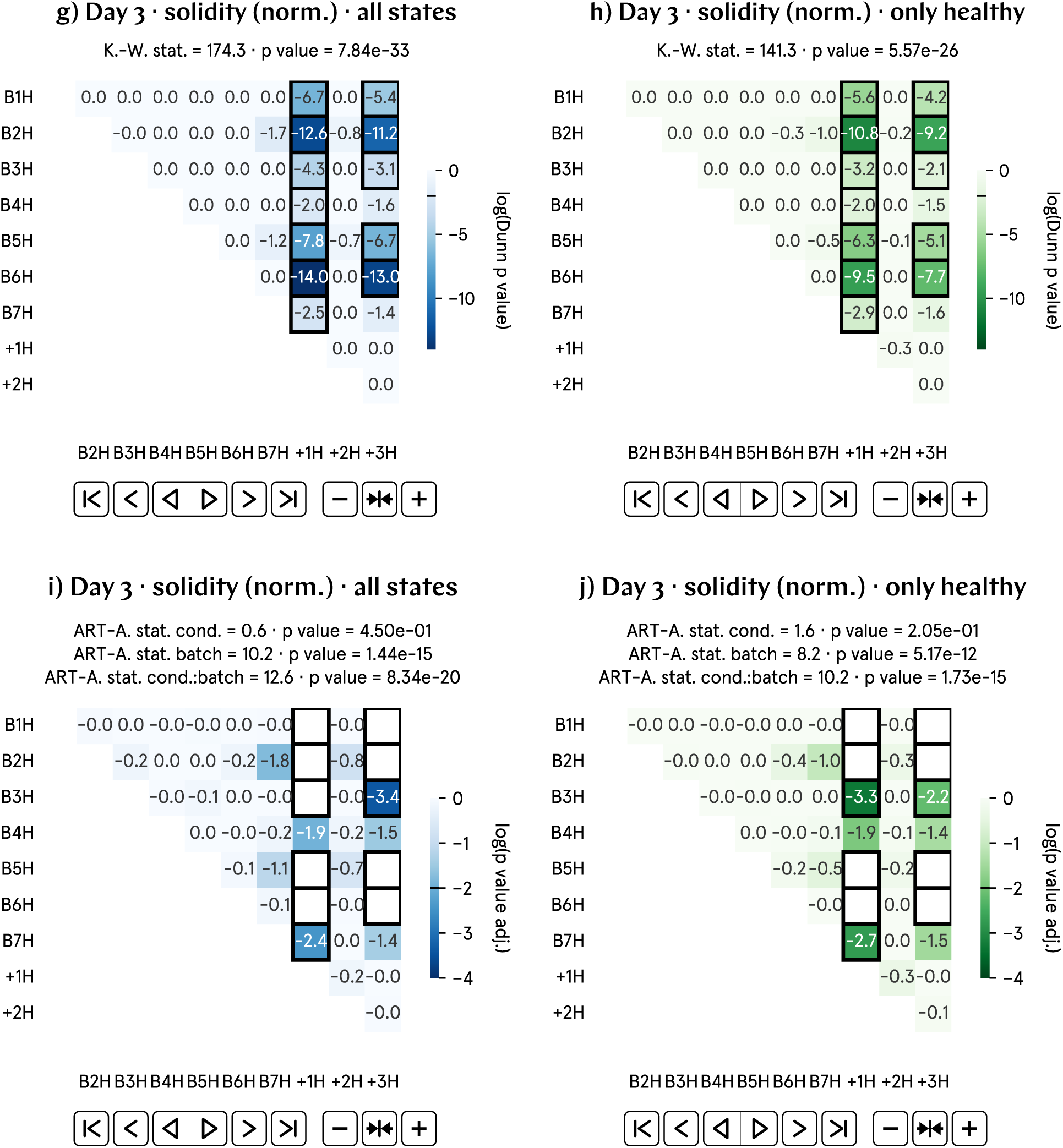
At day 3 p.f., the solidity of hypomagnetic embryos is approximately 3% lower than that of the control population. a) and b) Normalized solidity by batch and aggregate data; B1–7 represent experimental runs, and +1–3 correspond to positive control runs, in which an artificial magnetic field inside the hypomagnetic chamber simulates Earth’s. c) and d) Analysis of the population averages and medians further support the observed 3% lower solidity. e) and f) Effect size metrics, namely Cohen’s *d* and Cliff’s Δ, and Binhi’s *E* and the visibility *ν*, confirm a medium effect, consistently reflecting the 3% level difference. g) through j) Most statistically significant differences are found in the positive control tadpole populations raised in the hypomagnetic chamber, as expected.

### SI41. Day 3 solidity results

**Fig. SI33:**
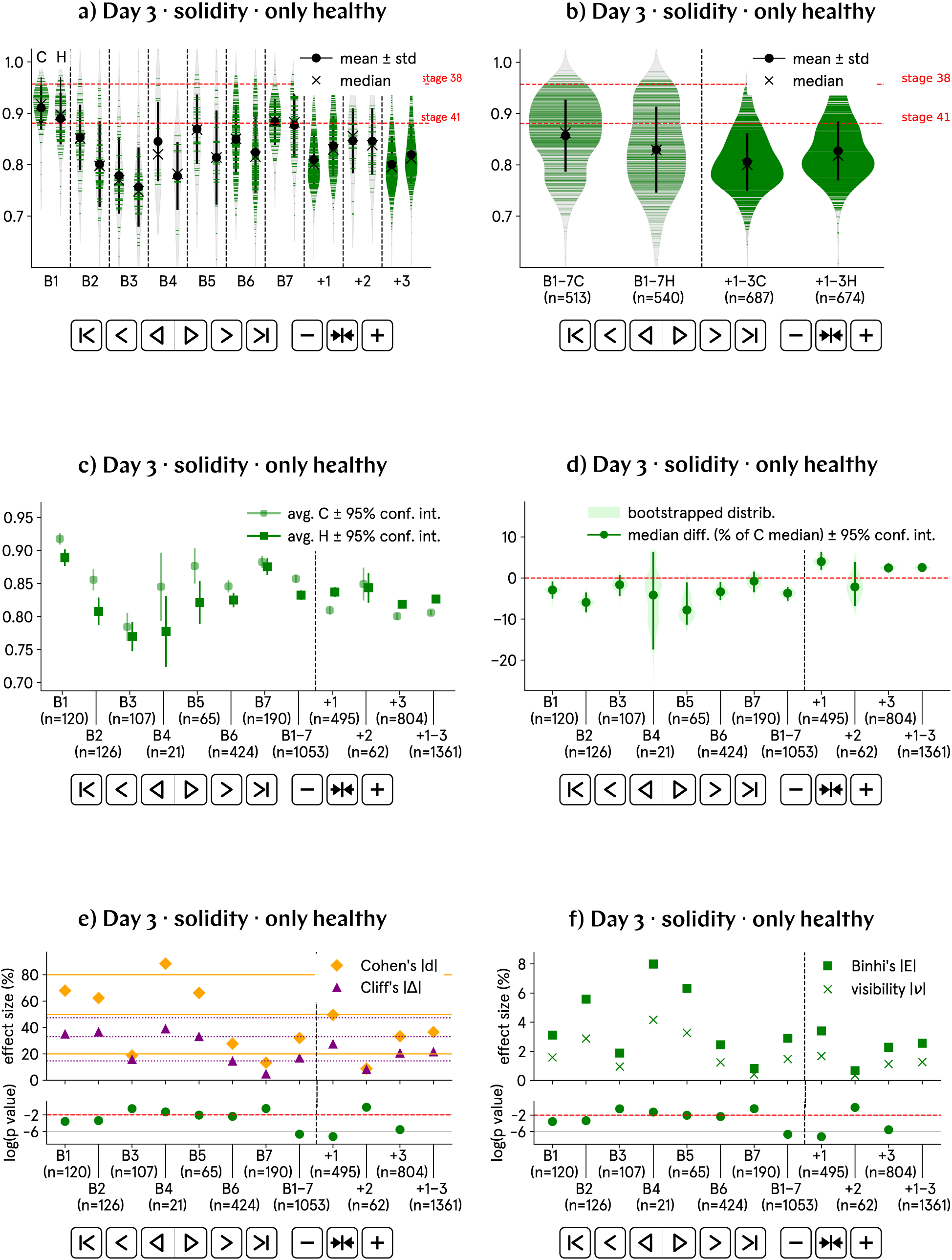
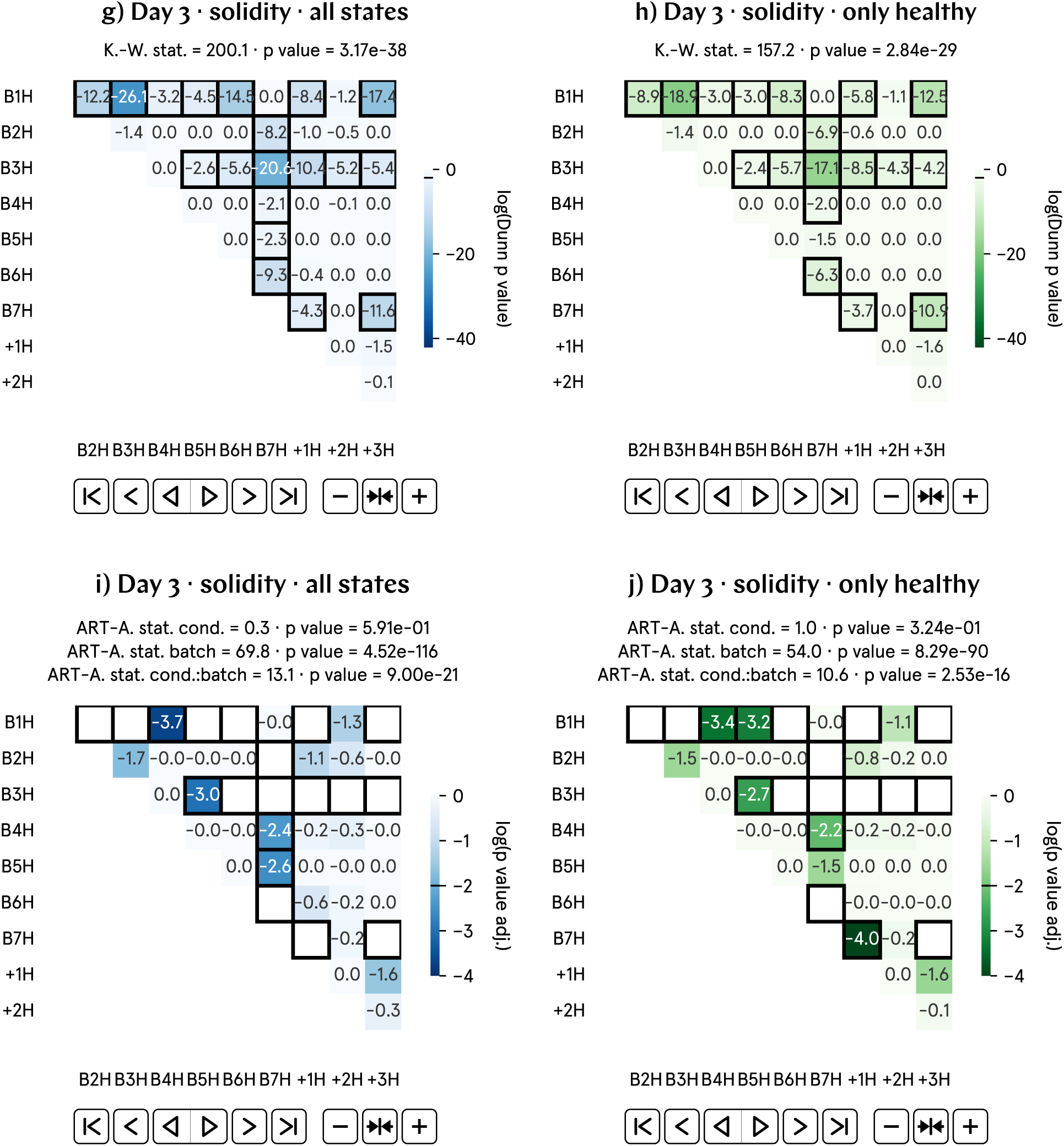
At day 3 p.f., the solidity of hypomagnetic embryos is approximately 3% lower than that of the control population. a) and b) Non-normalized solidity by batch and aggregate data; B1–7 represent experimental runs, and +1–3 correspond to positive control runs, in which an artificial magnetic field inside the hypomagnetic chamber simulates Earth’s. c) and d) Analysis of the population averages and medians further support the observed 3% lower solidity. e) and f) Effect size metrics, namely Cohen’s *d* and Cliff’s Δ, and Binhi’s *E* and the visibility *ν*, confirm a medium effect, consistently reflecting the 3% level difference. g) through j) Given the substantial variation in solidity across batches, these plots may not provide an accurate representation of our data. However, these plots do reveal that batches B1, B3 and B7 are anomalous according to this metric.

### SI42. Day 3 elongation (norm.) results

**Fig. SI34:**
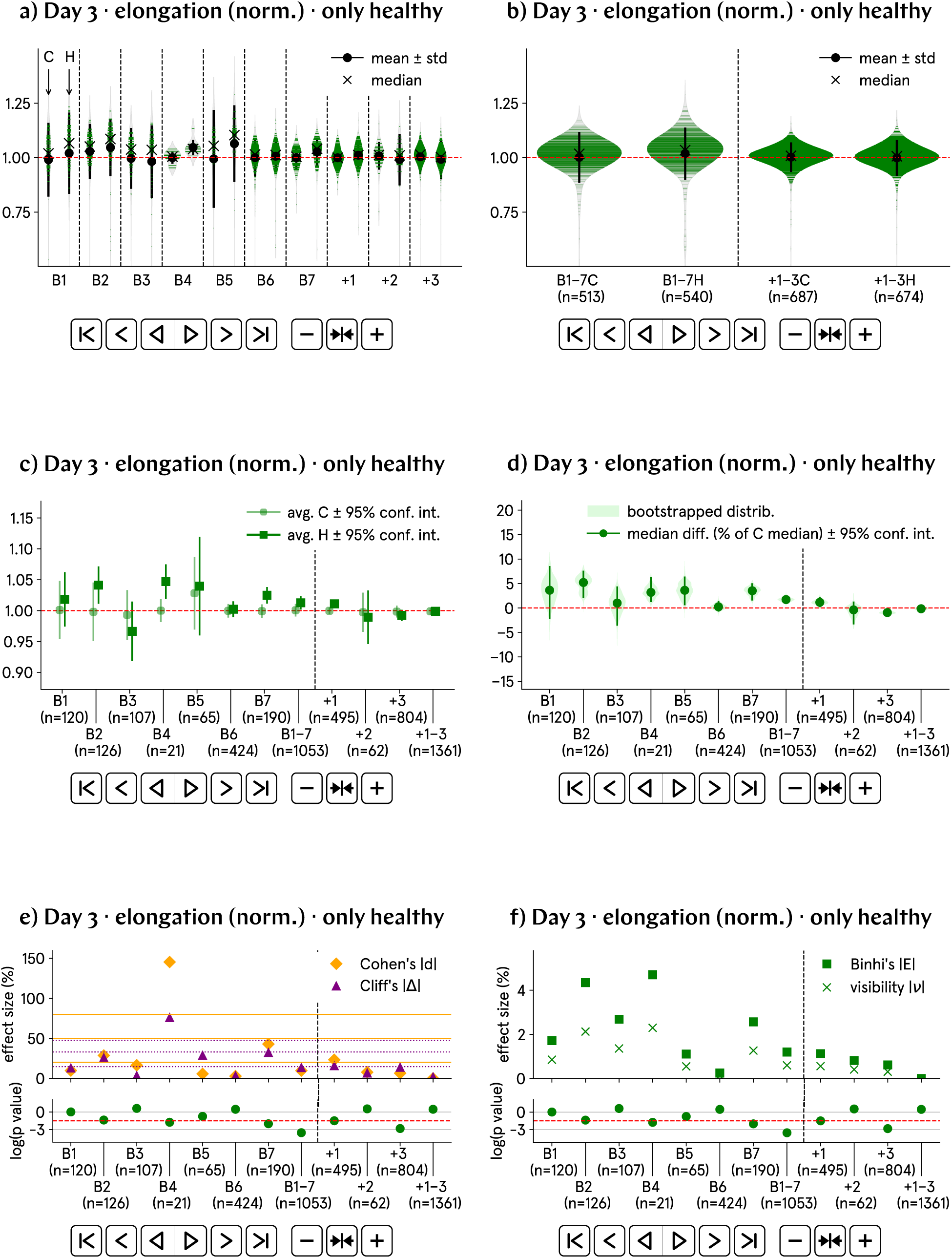
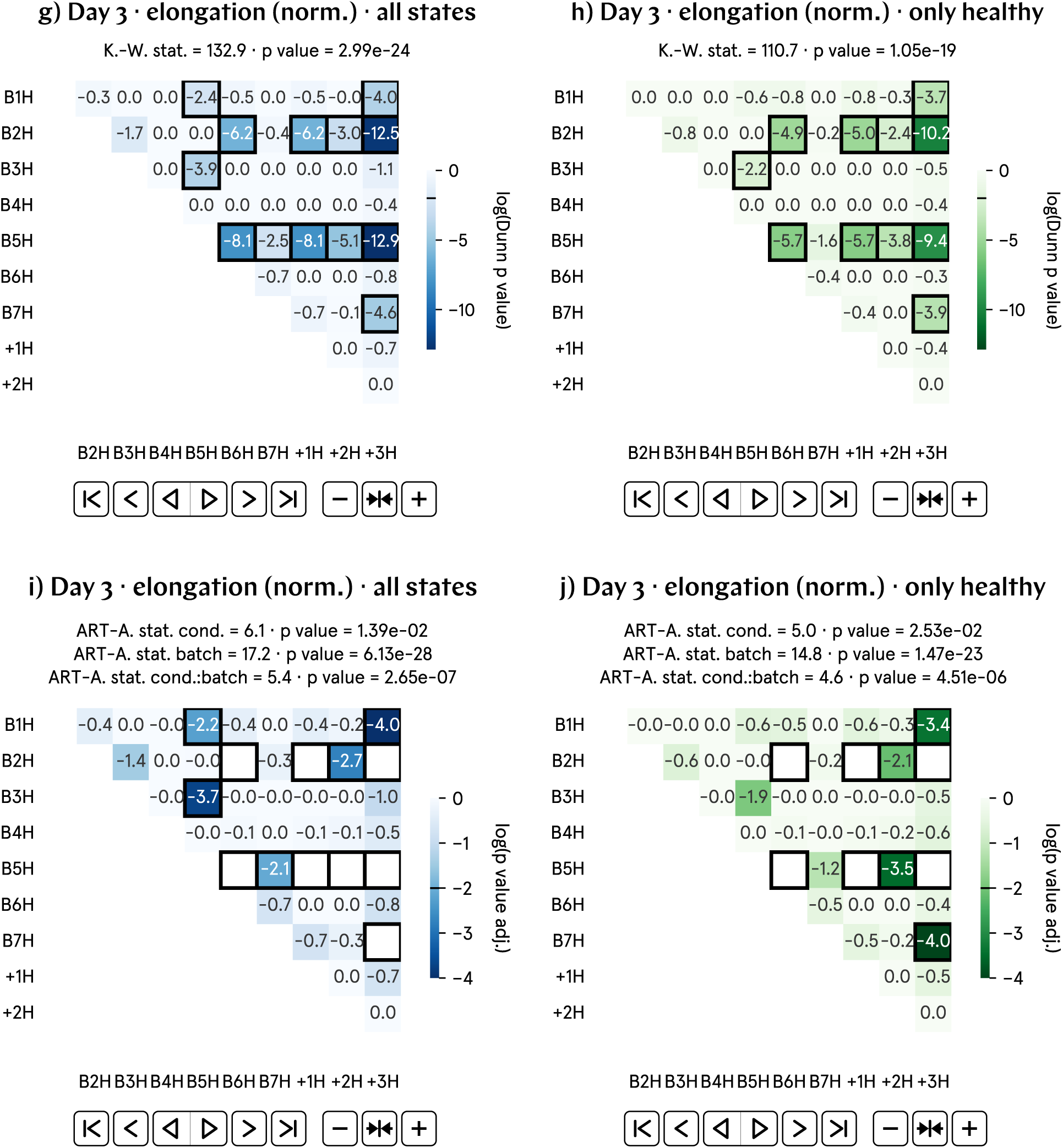
At day 3 p.f., the elongation of hypomagnetic embryos is approximately 1% higher than that of the control population. **a)** and **b)** Normalized elongation by batch and aggregate data; B1-7 represent experimental runs, and +1-3 correspond to positive control runs, in which an artificial magnetic field inside the hypomagnetic chamber simulates Earth’s. **c)** and **d)** Analysis of the population averages and medians further support the observed 1% higher elongation. **e)** and **f)** Effect size metrics, namely Cohen’s d and Cliff’s Δ, and Binhi’s *E* and the visibility *ν*, confirm a medium effect, consistently reflecting the 1% level difference. **g)** through **j)** Some statistically significant differences are found in the positive control tadpole populations raised in the hypomagnetic chamber, as expected. Batches B2 and B5 are also anomalous.

### SI43. Day 3 elongation results

**Fig. SI35:**
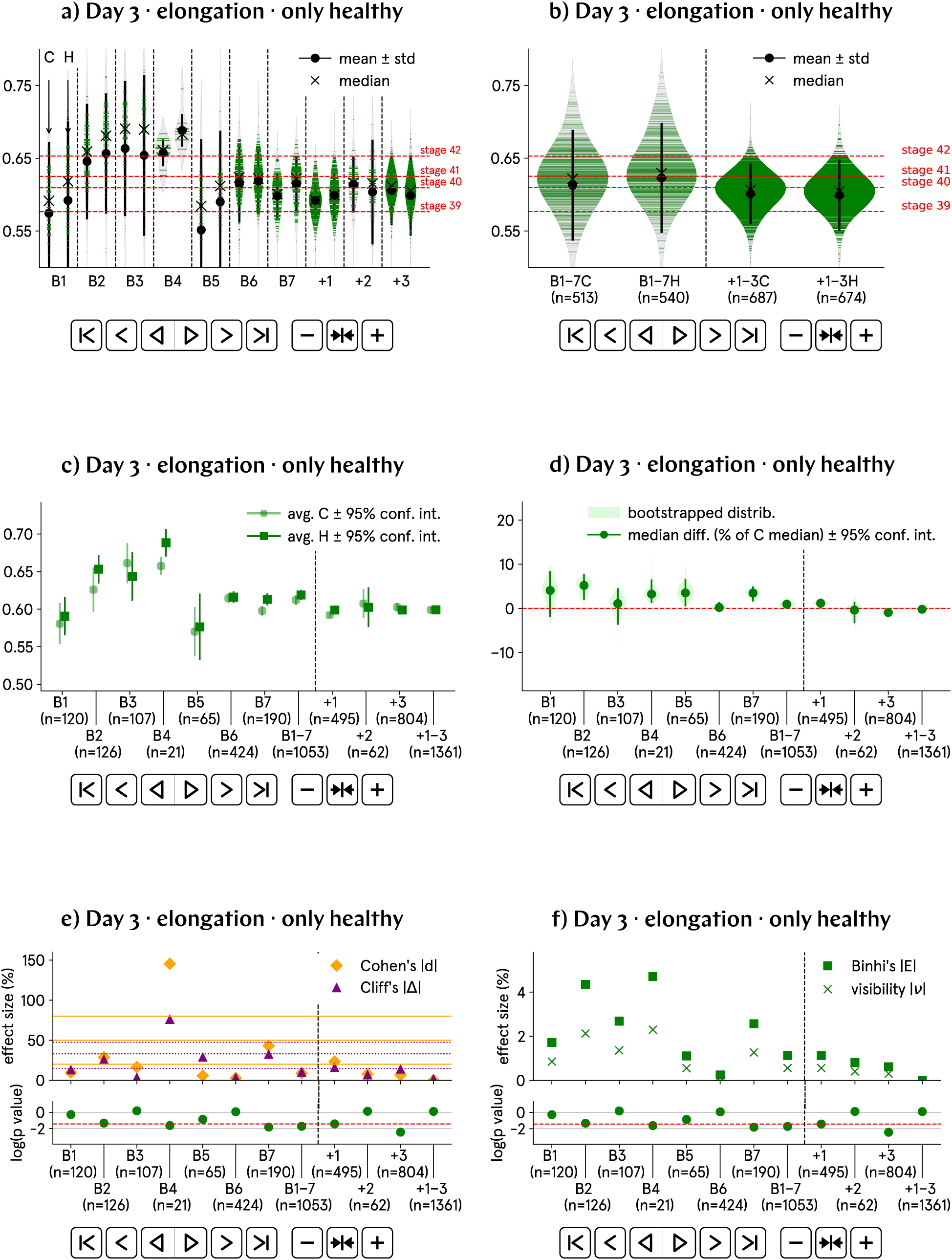
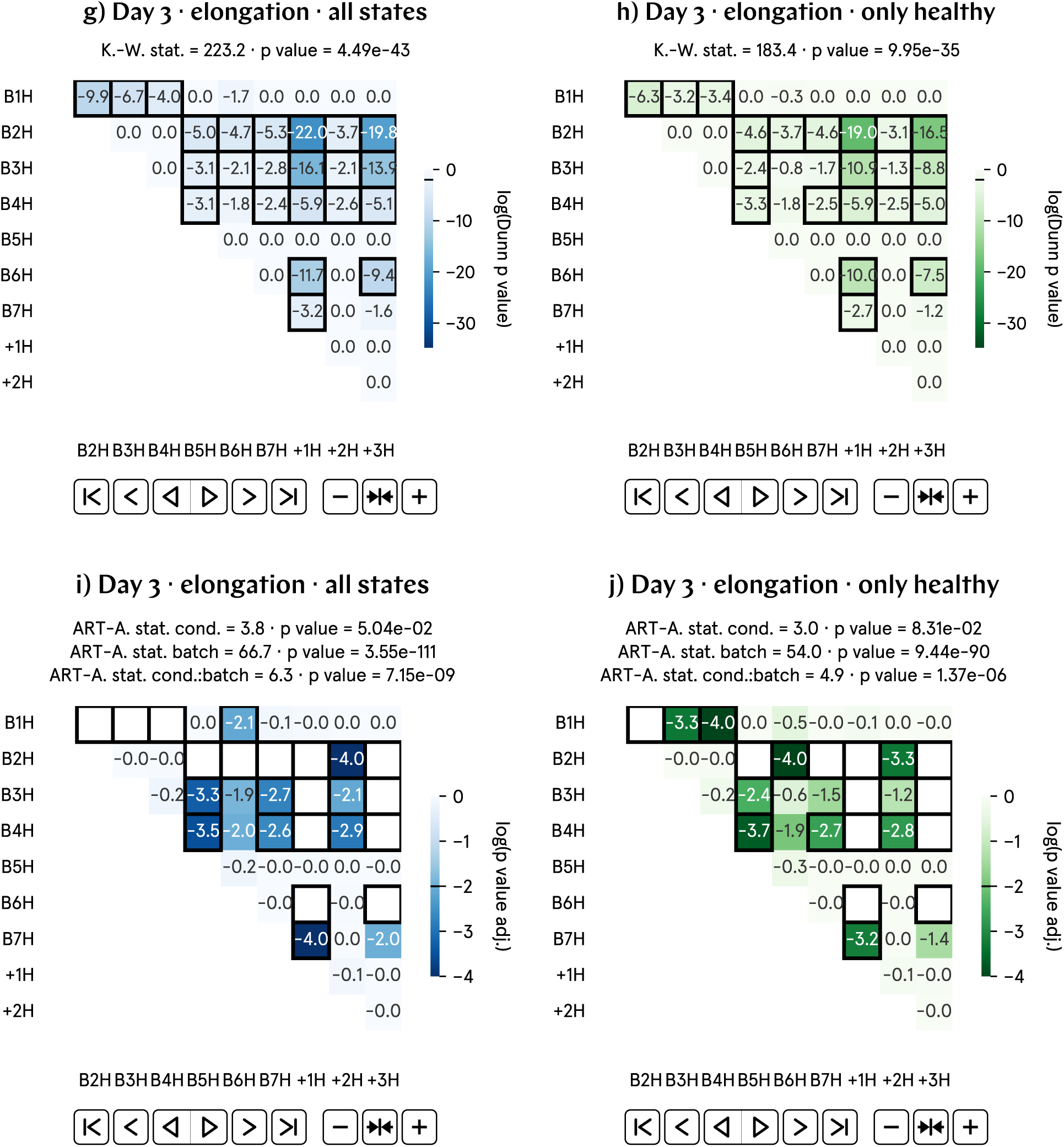
At day 3 p.f., the elongation of hypomagnetic embryos is approximately 1% higher than that of the control population. a) and b) Non-normalized elongation by batch and aggregate data; B1–7 represent experimental runs, and +1–3 correspond to positive control runs, in which an artificial magnetic field inside the hypomagnetic chamber simulates Earth’s. c) and d) Analysis of the population averages and medians further support the observed 1% higher elongation. e) and f) Effect size metrics, namely Cohen’s *d* and Cliff’s Δ, and Binhi’s *E* and the visibility *ν*, confirm a medium effect, consistently reflecting the 1% level difference. g) through j) Given the substantial variation in elongation across batches, these plots may not provide an accurate representation of our data.

### SI44. Day 3 roundness (norm.) results

**Fig. SI36:**
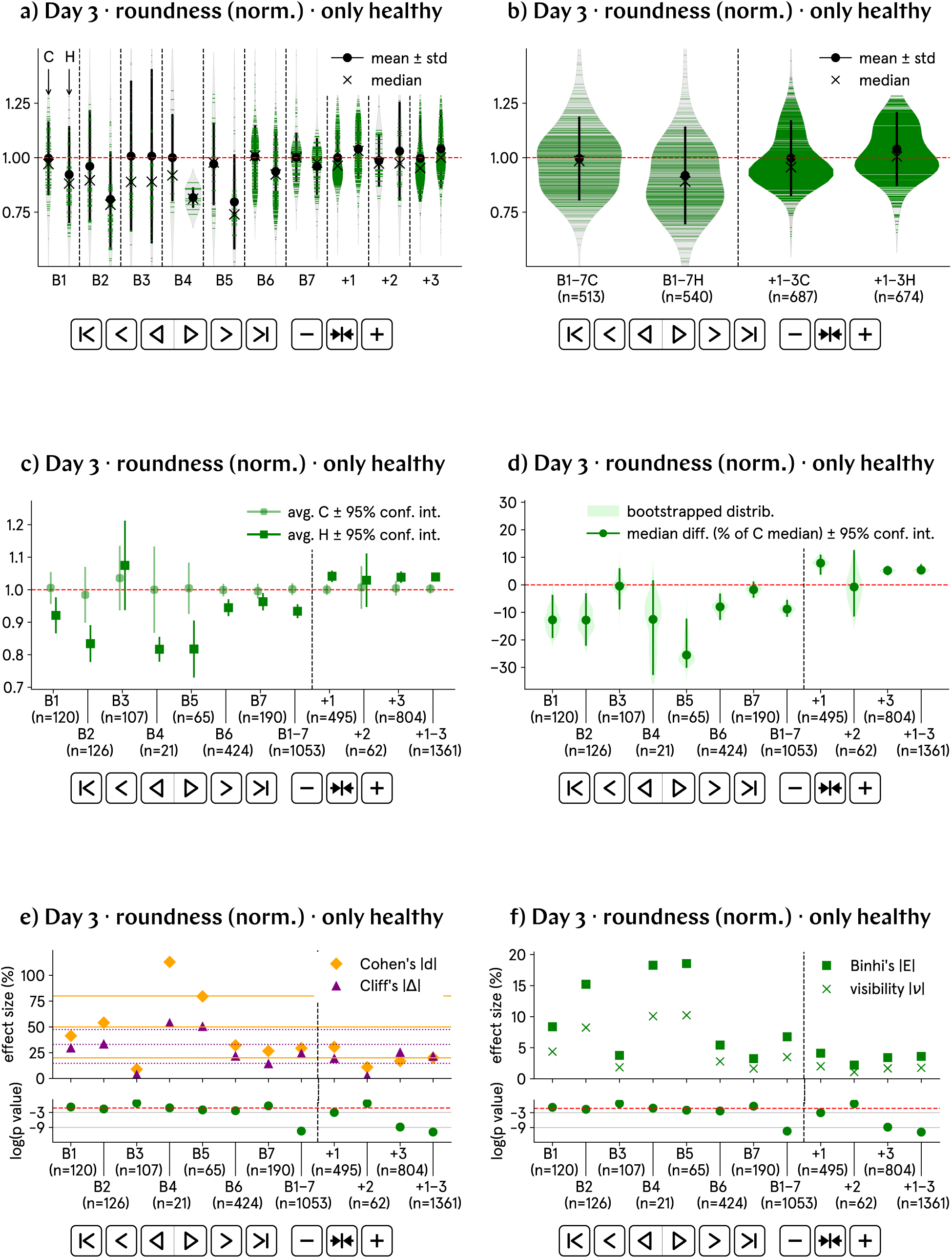
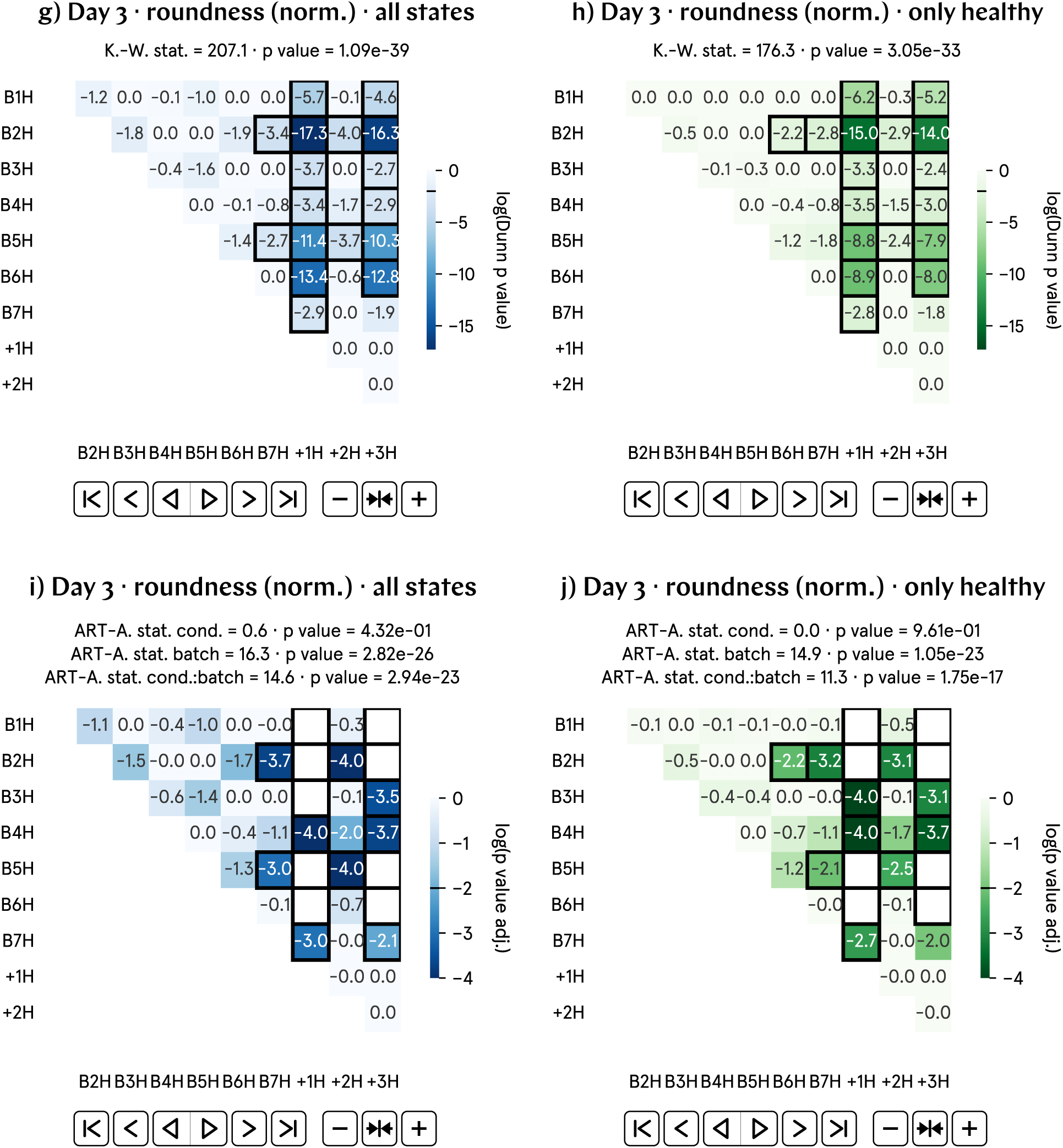
At day 3 p.f., the roundness of hypomagnetic embryos is approximately 9% lower than that of the control population. a) and b) Normalized roundness by batch and aggregate data; B1–7 represent experimental runs, and +1–3 correspond to positive control runs, in which an artificial magnetic field inside the hypomagnetic chamber simulates Earth’s. c) and d) Analysis of the population averages and medians further support the observed 9% lower roundness. e) and f) Effect size metrics, namely Cohen’s *d* and Cliff’s Δ, and Binhi’s *E* and the visibility *ν*, confirm a medium effect, consistently reflecting the 9% level difference. g) through j) Most statistically significant differences are found in the positive control tadpole populations raised in the hypomagnetic chamber, as expected. Batches B2 and B5 are also anomalous.

### SI45. Day 3 roundness results

**Fig. SI37:**
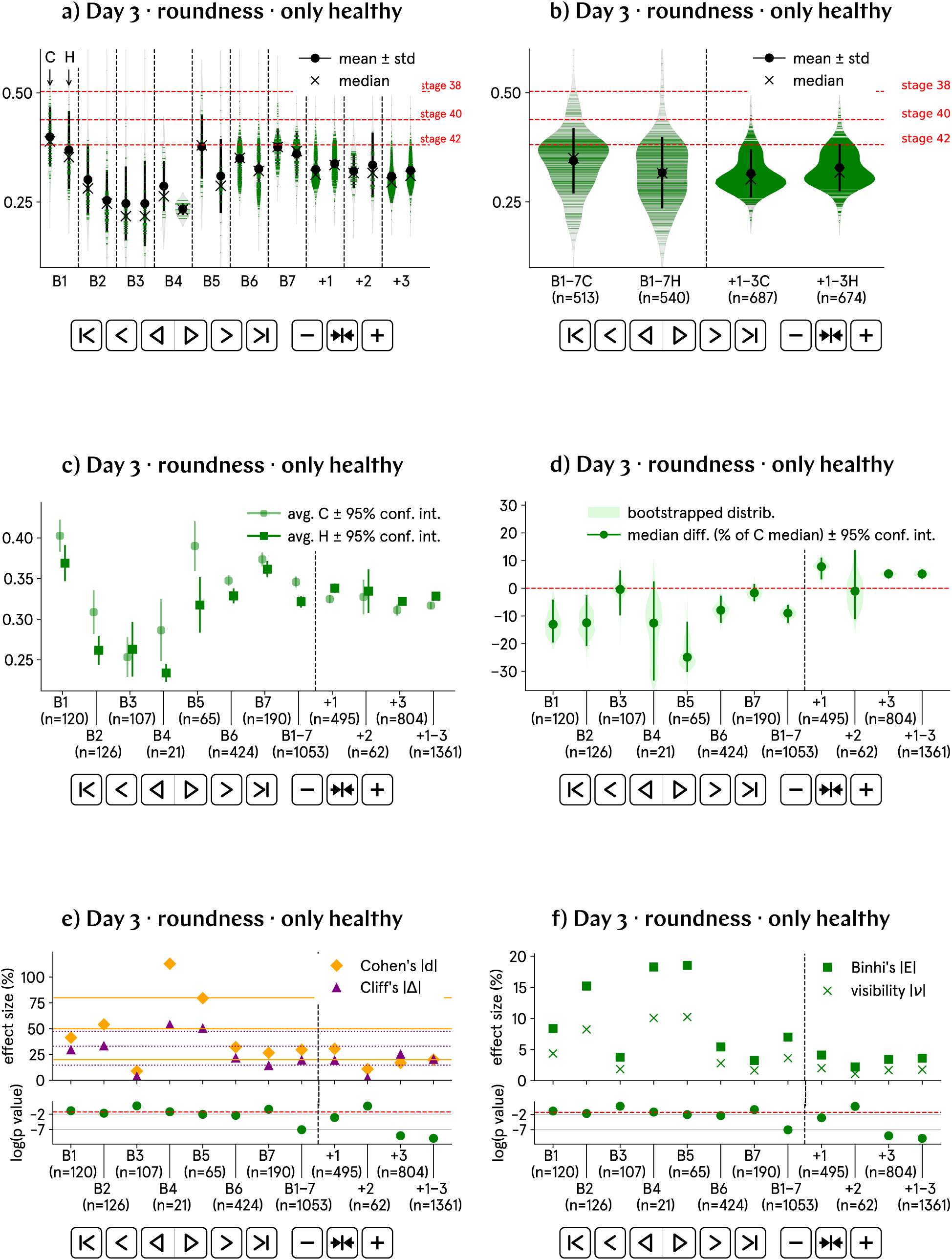
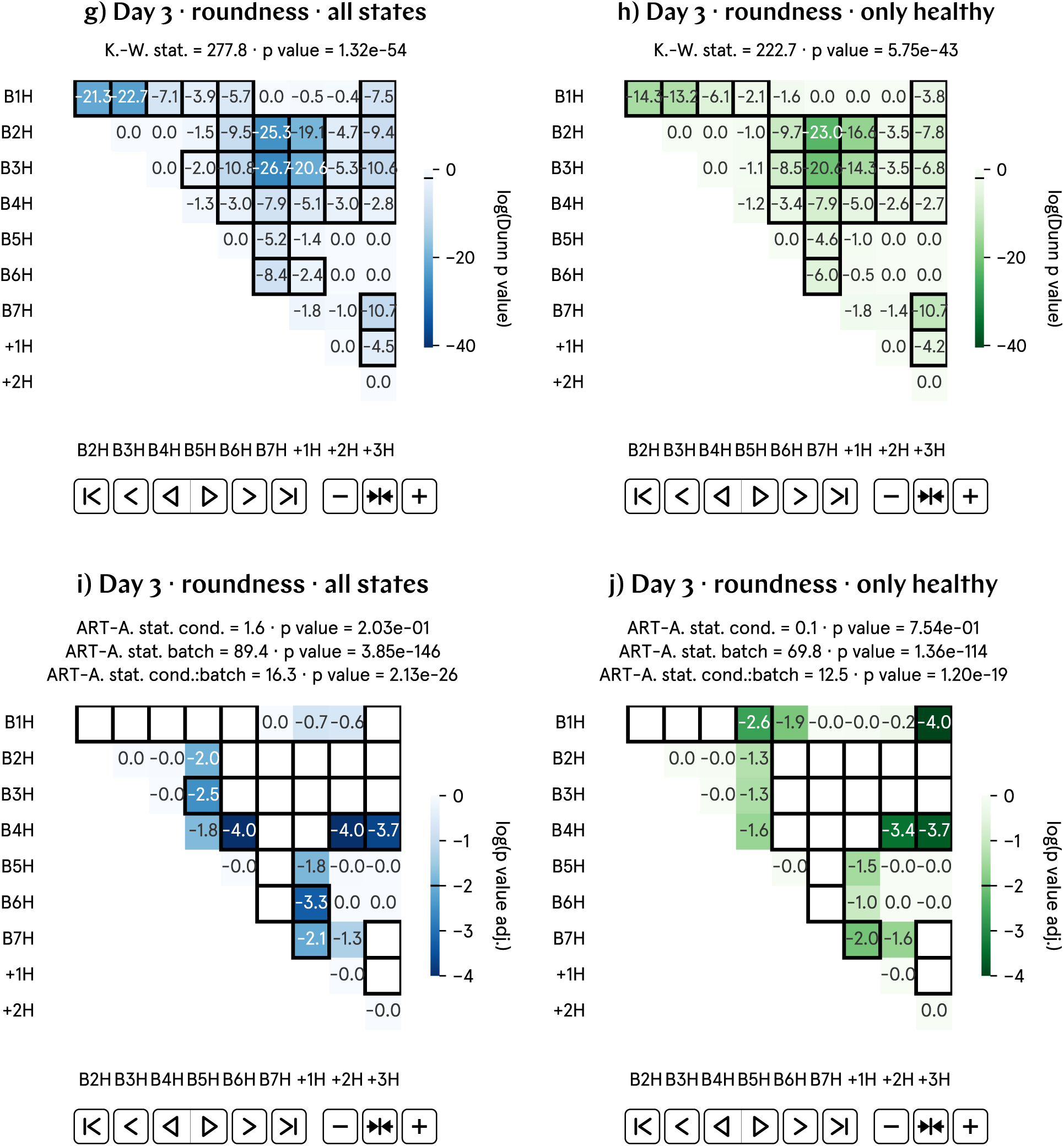
At day 3 p.f., the roundness of hypomagnetic embryos is approximately 9% lower than that of the control population. a) and b) Non-normalized roundness by batch and aggregate data; B1–7 represent experimental runs, and +1–3 correspond to positive control runs, in which an artificial magnetic field inside the hypomagnetic chamber simulates Earth’s. c) and d) Analysis of the population averages and medians further support the observed 9% lower roundness. e) and f) Effect size metrics, namely Cohen’s *d* and Cliff’s Δ, and Binhi’s *E* and the visibility *ν*, confirm a medium effect, consistently reflecting the 9% level difference. g) through j) Given the substantial variation in roundness across batches, these plots may not provide an accurate representation of our data.

### SI46. Day 2 to 3 measures: major axis, total curvature, convexity and solidity

We also quantify the extent to which the progression of morphological parameters between days 2 and 3 is different across control and hypomagnetic populations. For each particular tadpole, we thus track changes in major axis, curvature, convexity and solidity.

From the above morphological parameters, the only that remains to be defined is the convexity. Convexity quantifies how close a shape is to being convex. It compares the perimeter of the shape to the perimeter of its convex hull, which is the smallest convex shape that can fully enclose the shape. Convexity is defined as:

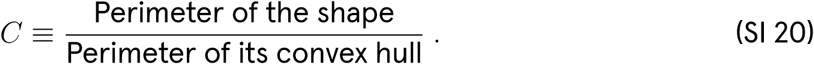

Convexity values range from 1 (for perfectly convex shapes) to values less than 1 for shapes with concavities.

Our analysis of the progression of morphological parameters employs four distinct comparison measures (explained in Supplementary Section SI15). All measures consistently support the conclusion that the control and hypomagnetic tadpole populations evolve differently from day 2 to day 3 in a statistically significant manner. The results for the major axis are presented in Supplementary Figs. SI38 through SI41; for total curvature in Supplementary Figs. SI42 through SI45; for convexity in Supplementary Figs. SI46 through SI49; and for solidity in Supplementary Figs. SI50 through SI53.

The data are normalized to the control group average, per batch. This normalization is especially important for properties like major axis and curvature, where slight magnification changes occur between image acquisitions, ensuring a fair comparison. Although convexity and solidity inherently do not require normalization, they are normalized to facilitate consistent comparisons across batches and across the four different effect change measures.

We are unable to confidently explain why the control group during the positive control runs (+1–3) exhibit an overall larger Z-score difference in convexity and solidity when compared to the control group during the experimental runs (B1–7). This discrepancy may reflect limitations in these specific measures when capturing tadpole shape changes in our dataset. However, it is important to note that this trend is reversed in the experimental runs, further highlighting the distinct morphological progression between tadpoles exposed to the hypomagnetic versus the control magnetic field condition.

During bootstrapping, the normalization of the progression data resulted in excessively large sampled values for all four measures. Consequently, the bootstrapping plots of type d), from Supplementary Fig. SI38 to Supplementary Fig. SI53, were generated using non-normalized data as an exception. Despite this adjustment, unusually large sampled values persisted, likely due to small denominators in the calculations. Additionally, some non-normalized bootstrapping results contradict the normalized distributions; for example, in the convexity results of Supplementary Fig. SI46, bootstrapped plot d) shows a positive median difference, whereas b) suggests a decrease in convexity for day 3 under hypomagnetic conditions. This inconsistency raises concerns about the reliability of these non-normalized bootstrapped plots.

Finally, we highlight the highly anomalous plot b) of Supplementary Fig. SI44, which shows the aggregate violins for percentage change in curvature progression. Unlike the other effect-change measures, percentage change uses only the day 2 property value in the denominator. If this value is small, the resulting percentage change can become disproportionately large. This is likely responsible for the anomaly observed in the plot.

### SI47. Day 2 to 3 major axis progression (norm.) results, Z-score difference

**Fig. SI38:**
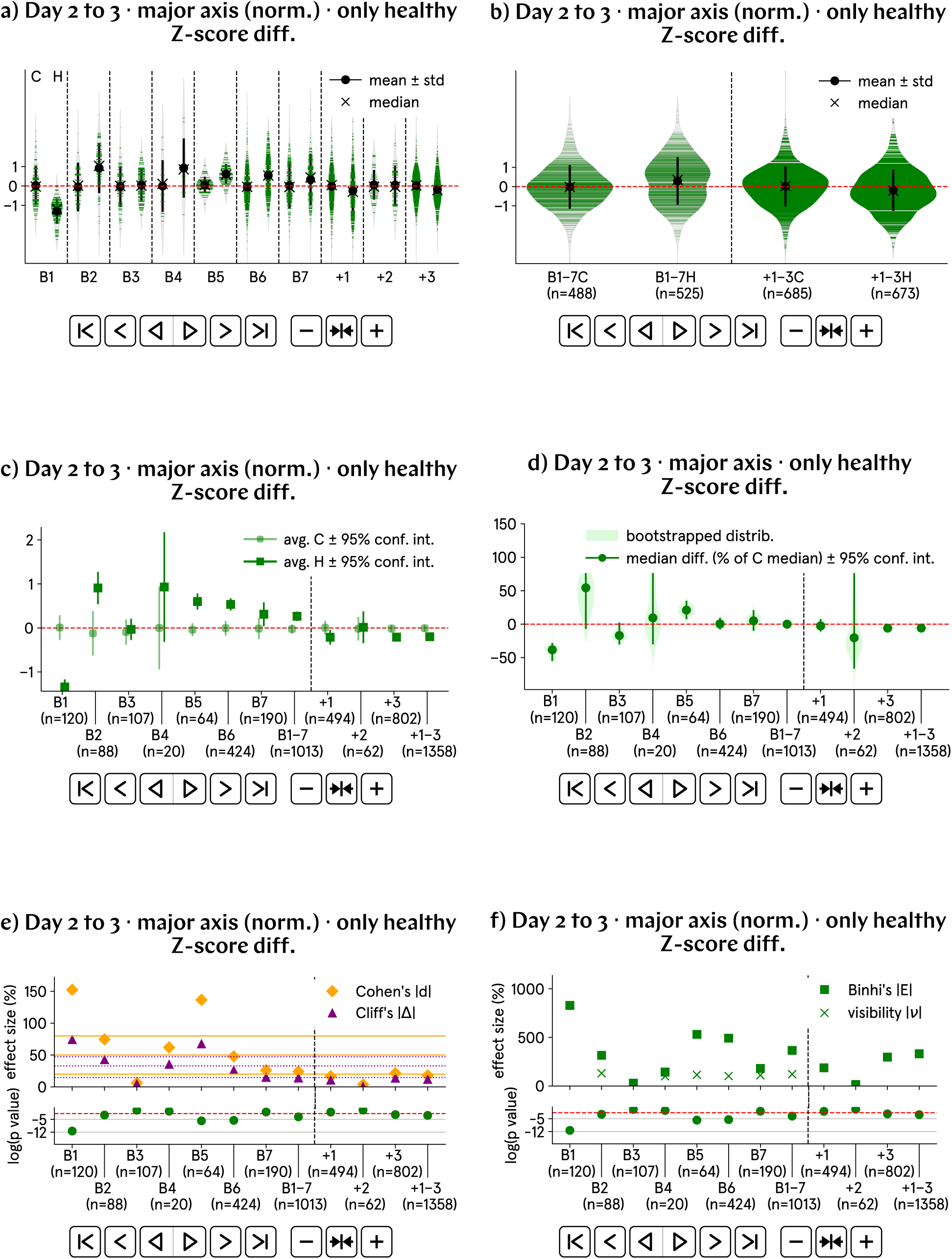
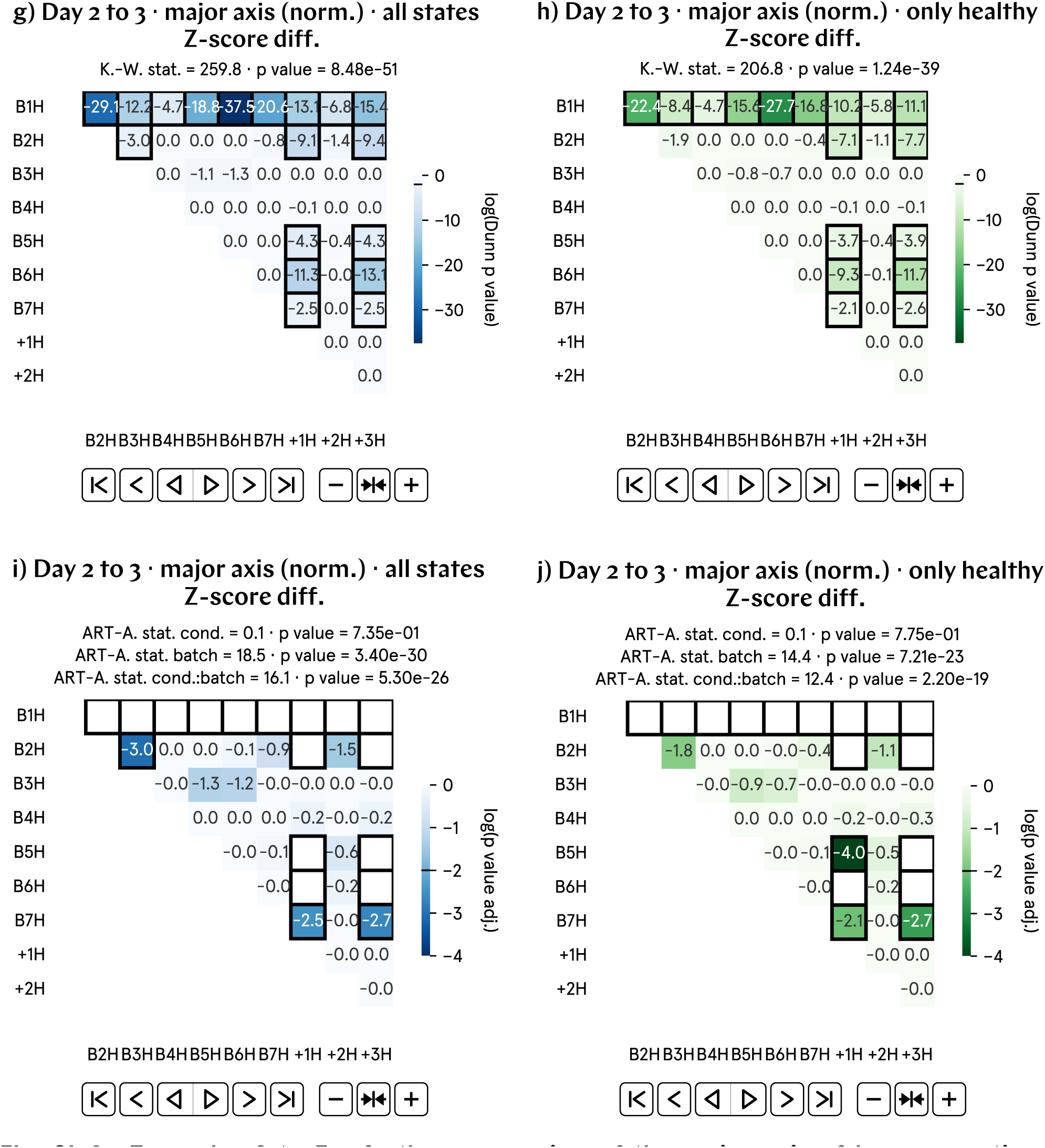
From day 2 to 3 p.f., the progression of the major axis of hypomagnetic embryos, as measured by the Z-score difference, is approximately 0.5 higher than that of the control population. a) and b) Normalized major axis Z-score difference by batch and aggregate data; B1–7 represent experimental runs, and +1–3 correspond to positive control runs, in which an artificial magnetic field inside the hypomagnetic chamber simulates Earth’s. c) and d) Analysis of the population averages and medians further support the observed 0.5 higher value. e) Cohen’s *d* and Cliff’s Δ confirm a medium effect.

### SI48. Day 2 to 3 major axis progression (norm.) results, visibility

**Fig. SI39:**
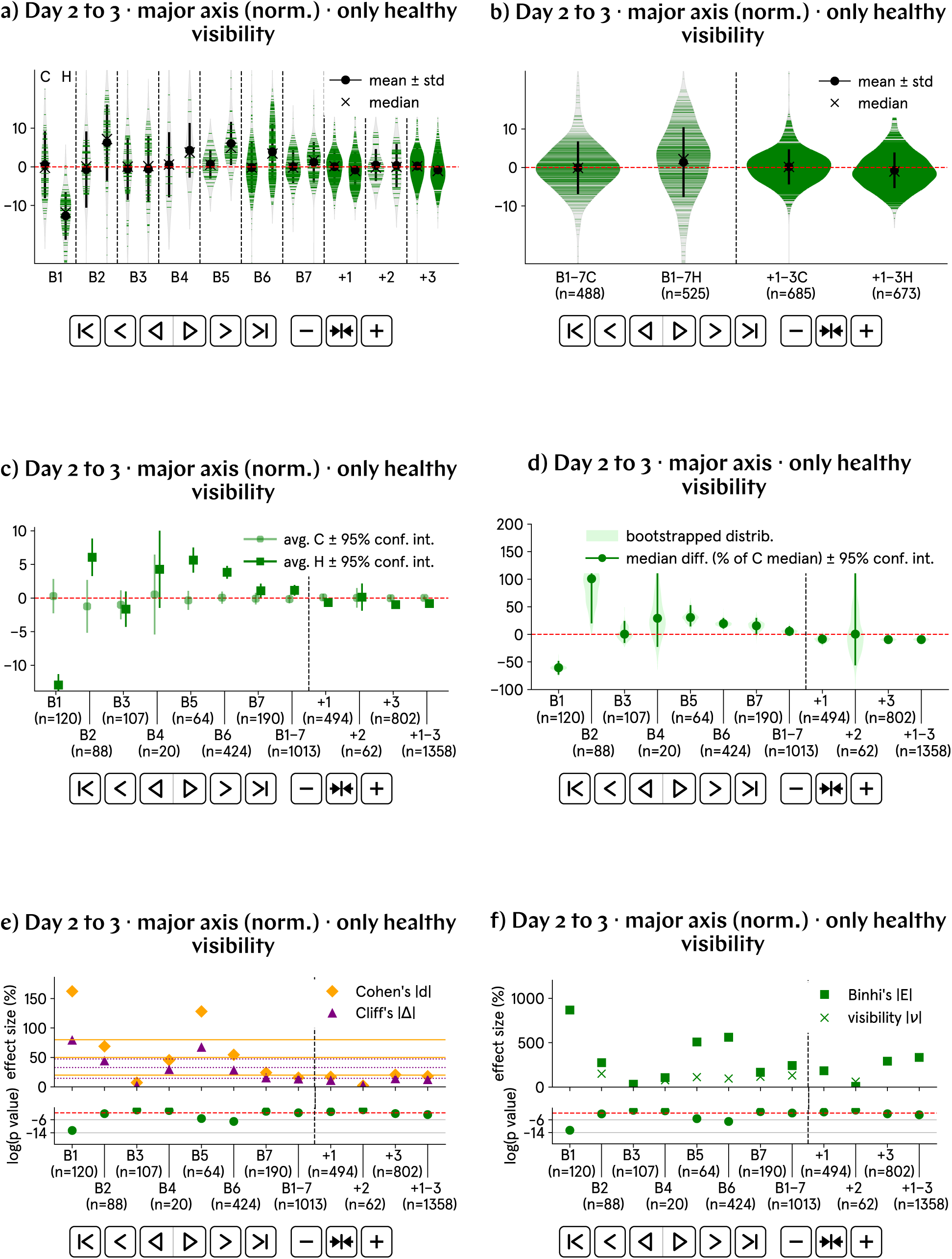
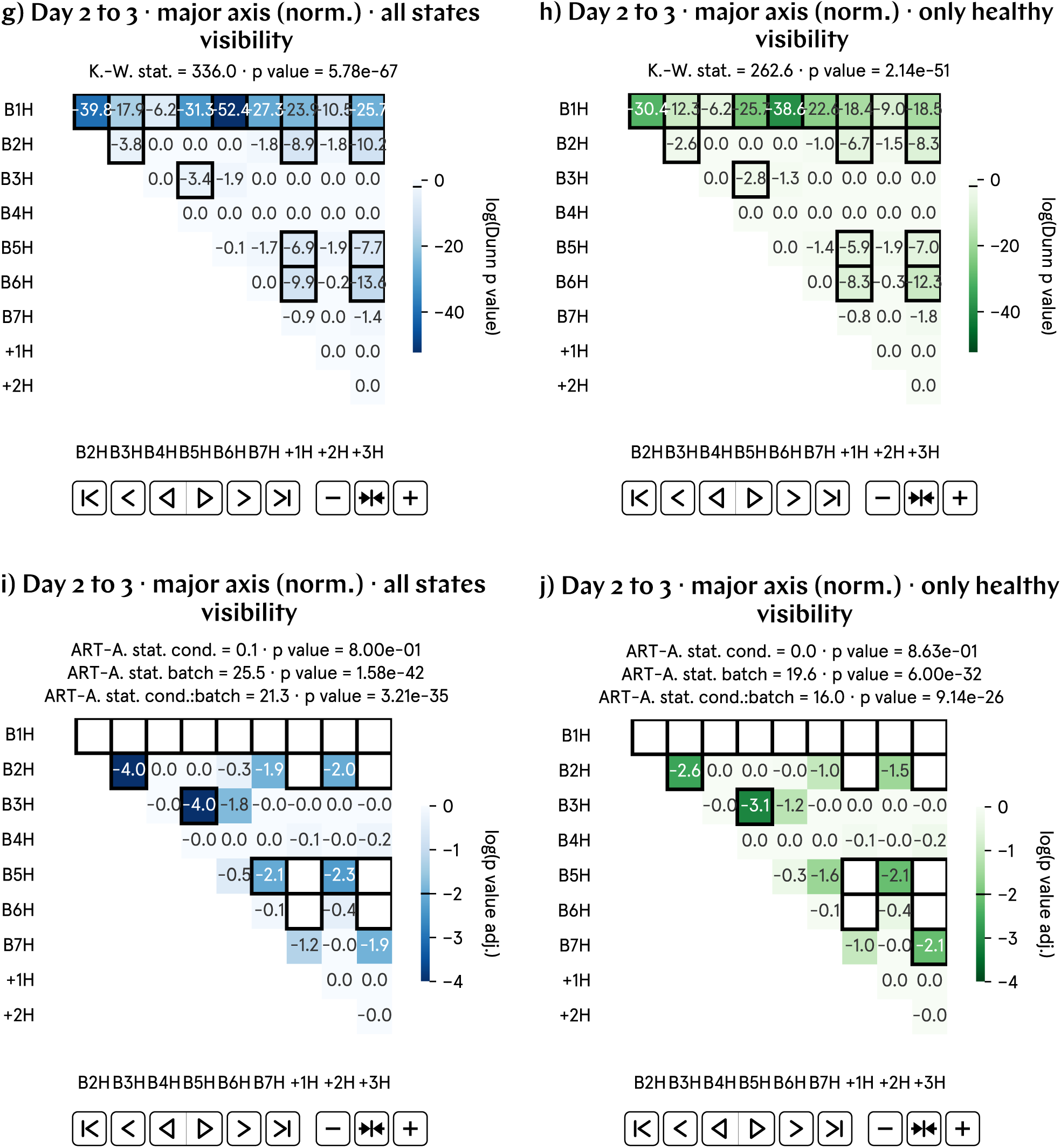
From day 2 to 3 p.f., the progression of the major axis of hypomagnetic embryos, as measured by the visibility, is approximately 3% higher than that of the control population. a) and b) Normalized major axis visibility by batch and aggregate data; B1–7 represent experimental runs, and +1–3 correspond to positive control runs, in which an artificial magnetic field inside the hypomagnetic chamber simulates Earth’s. c) and d) Analysis of the population averages and medians further support the observed 3% higher value. e) Cohen’s *d* and Cliff’s Δ confirm a medium effect. f) Binhi’s *E* and the visibility *ν* overestimate the effect size. g) through j) Most statistically significant differences are found in the positive control tadpole populations raised in the hypomagnetic chamber, as expected. Batch B1 is also anomalous.

### SI49. Day 2 to 3 major axis progression (norm.) results, percentage change

**Fig. SI40:**
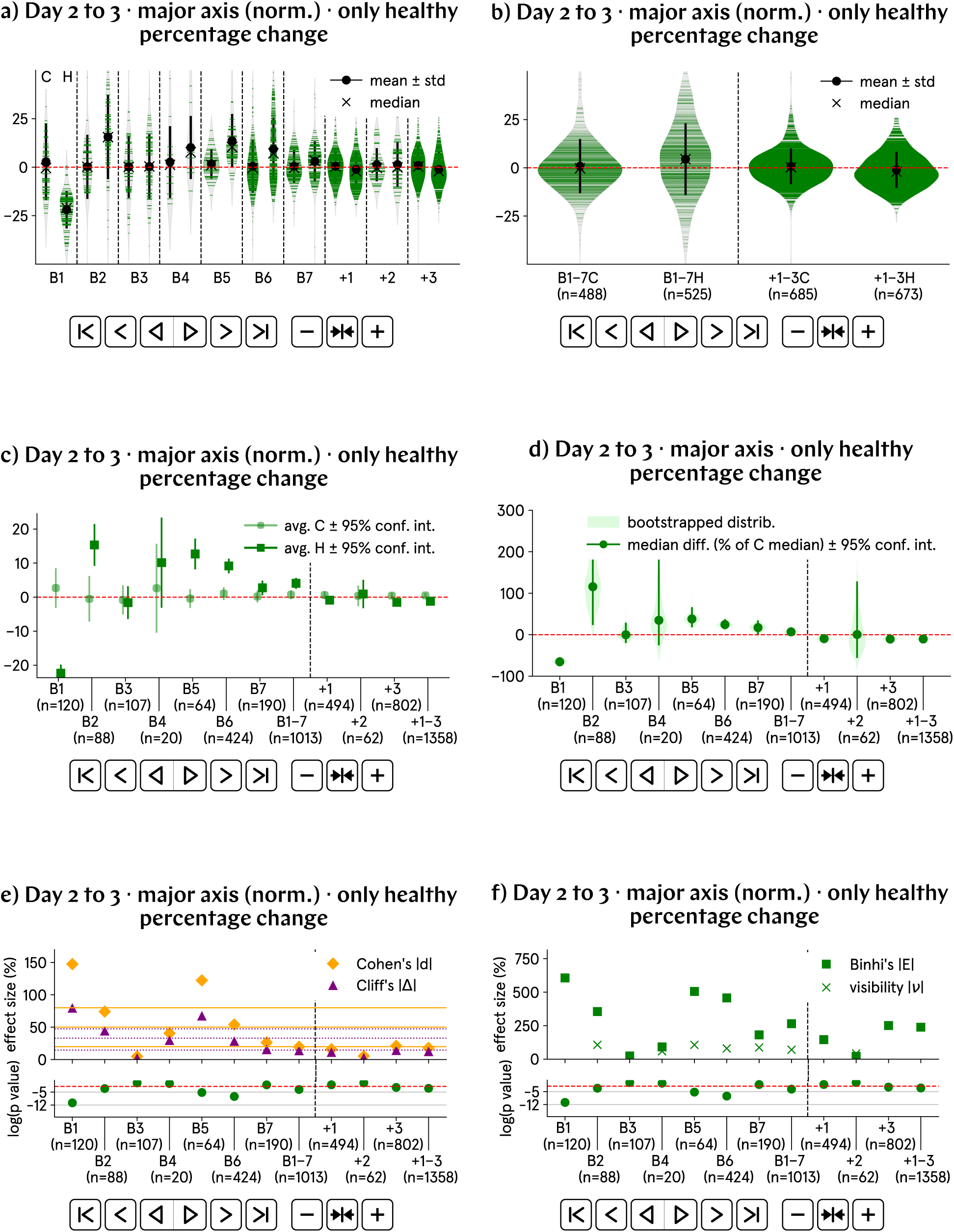
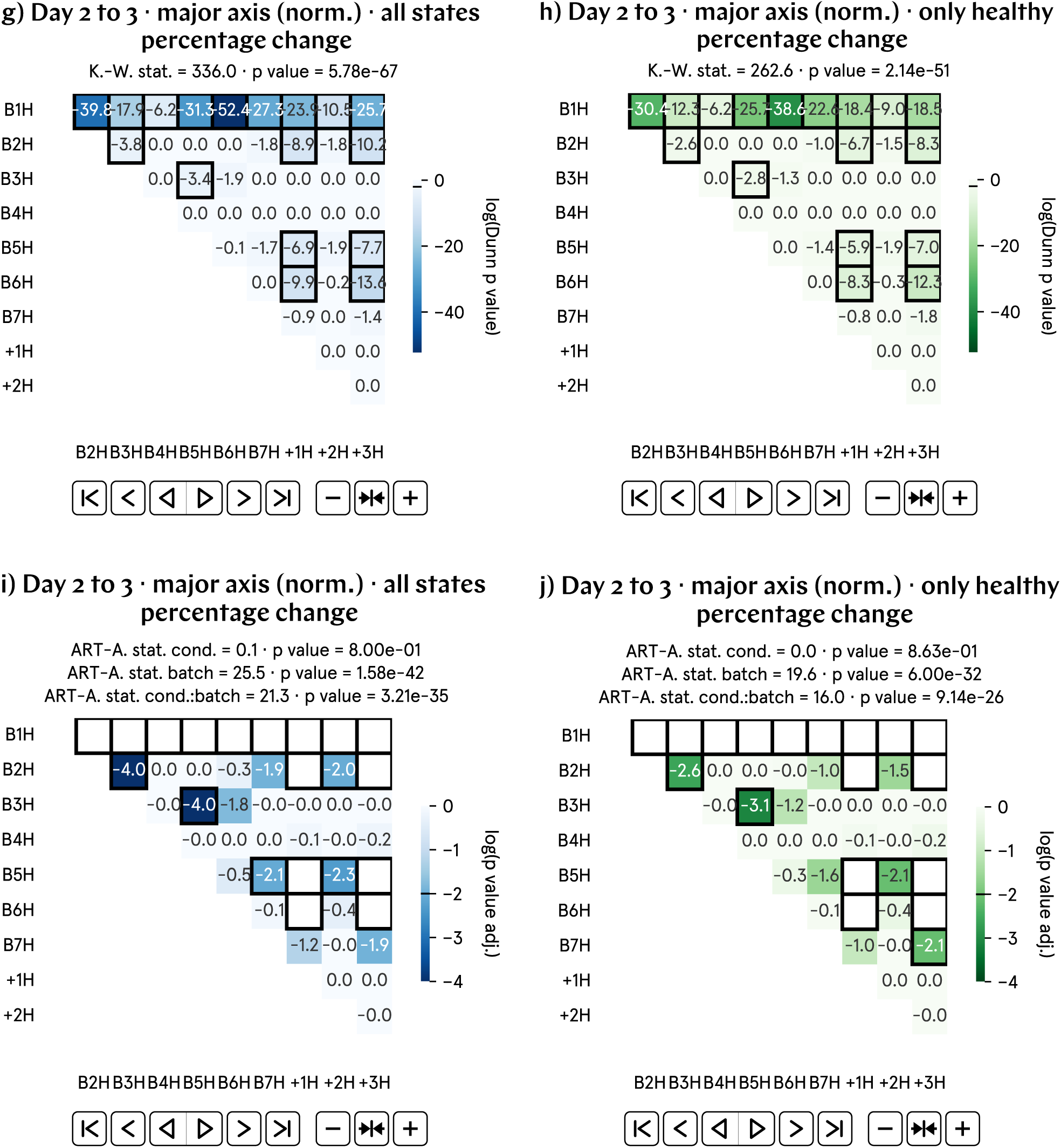
From day 2 to 3 p.f., the progression of the major axis of hypomagnetic embryos, as measured by the percentage change, is approximately 5% higher than that of the control population. a) and b) Normalized major axis percentage change by batch and aggregate data; B1–7 represent experimental runs, and +1–3 correspond to positive control runs, in which an artificial magnetic field inside the hypomagnetic chamber simulates Earth’s. c) and d) Analysis of the population averages and medians further support the observed 5% higher value. e) Cohen’s *d* and Cliff’s Δ confirm a medium effect. f) Binhi’s *E* and the visibility *ν* overestimate the effect size. g) through j) Most statistically significant differences are found in the positive control tadpole populations raised in the hypomagnetic chamber, as expected. Batch B1 is also anomalous.

### SI50. Day 2 to 3 major axis progression (norm.) results, relative difference

**Fig. SI41:**
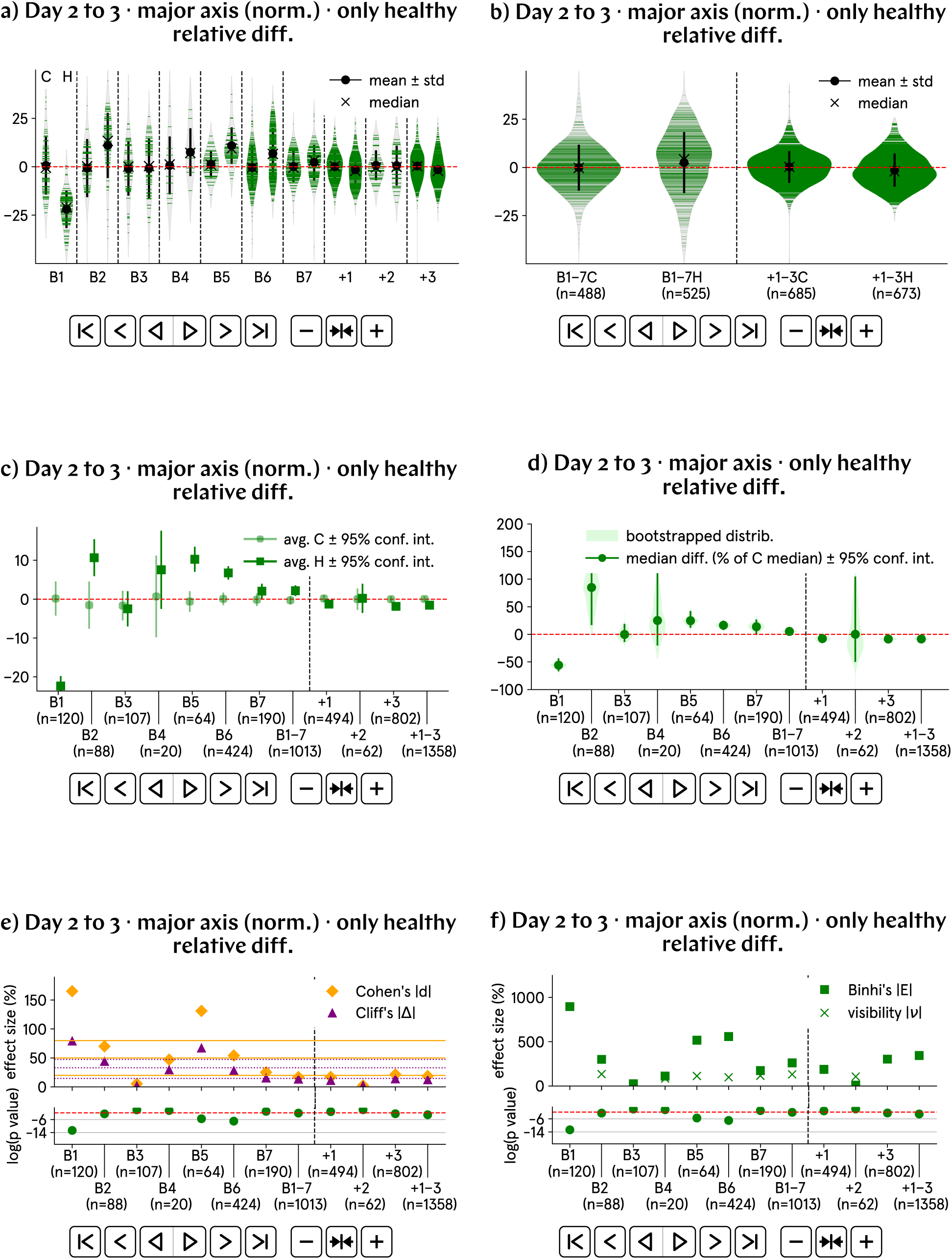
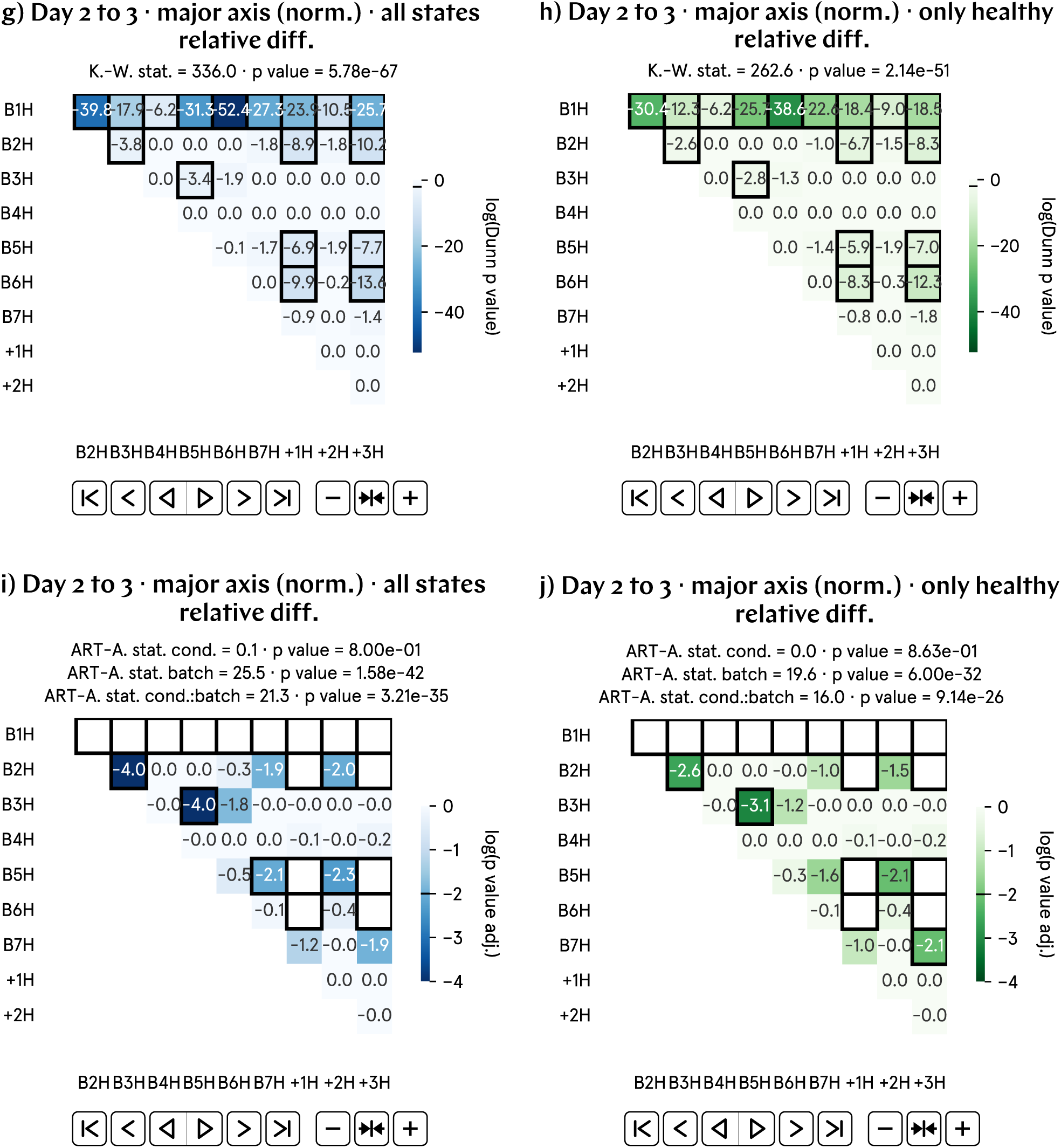
From day 2 to 3 p.f., the progression of the major axis of hypomagnetic embryos, as measured by the relative difference, is approximately 3% higher than that of the control population. a) and b) Normalized major axis relative difference by batch and aggregate data; B1–7 represent experimental runs, and +1–3 correspond to positive control runs, in which an artificial magnetic field inside the hypomagnetic chamber simulates Earth’s. c) and d) Analysis of the population averages and medians further support the observed 3% higher value. e) Cohen’s *d* and Cliff’s Δ confirm a medium effect. f) Binhi’s *E* and the visibility *ν* overestimate the effect size. g) through j) Most statistically significant differences are found in the positive control tadpole populations raised in the hypomagnetic chamber, as expected. Batch B1 is also anomalous.

### SI51. Day 2 to 3 curvature progression (norm.) results, Z-score difference

**Fig. SI42:**
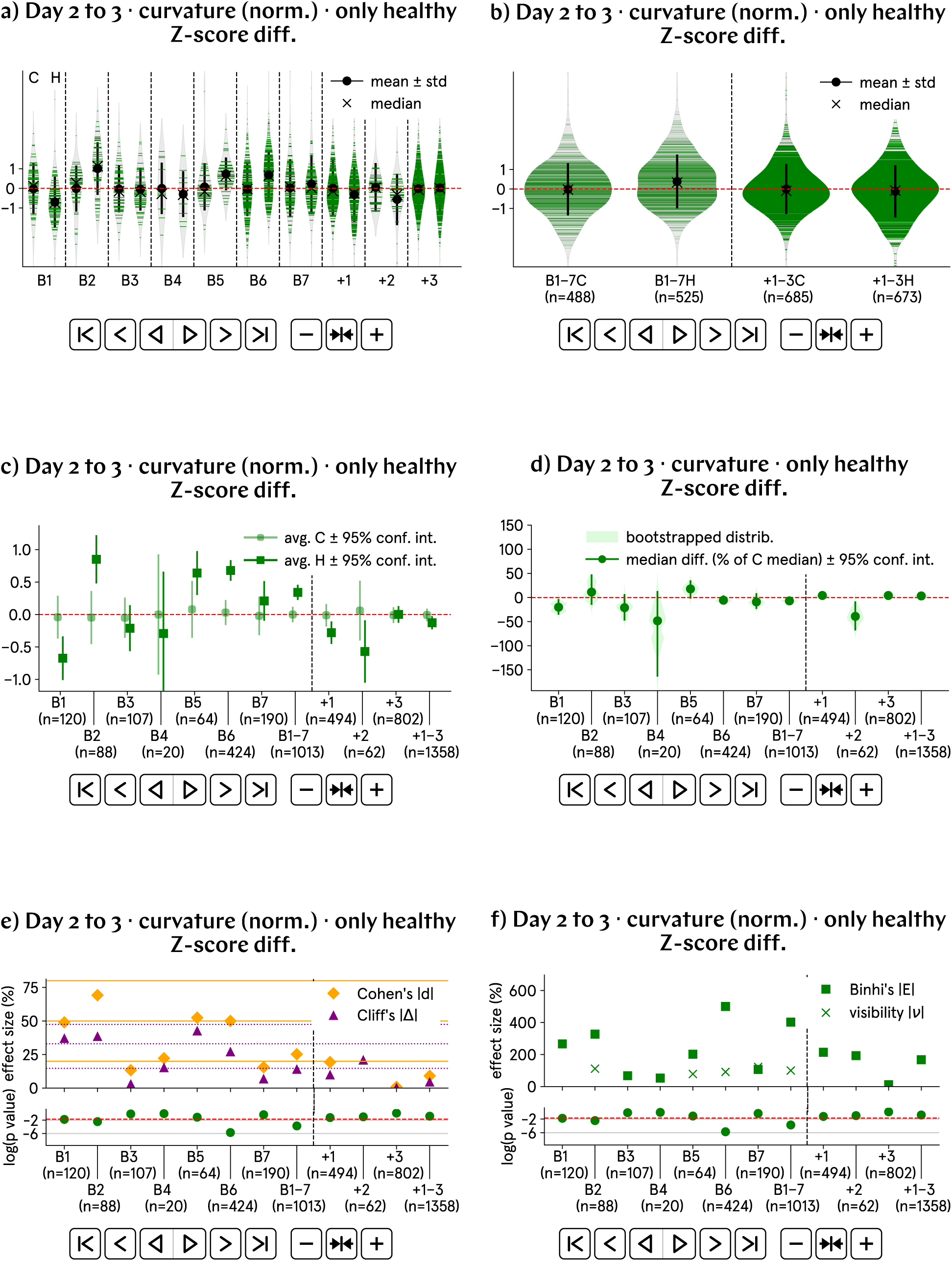
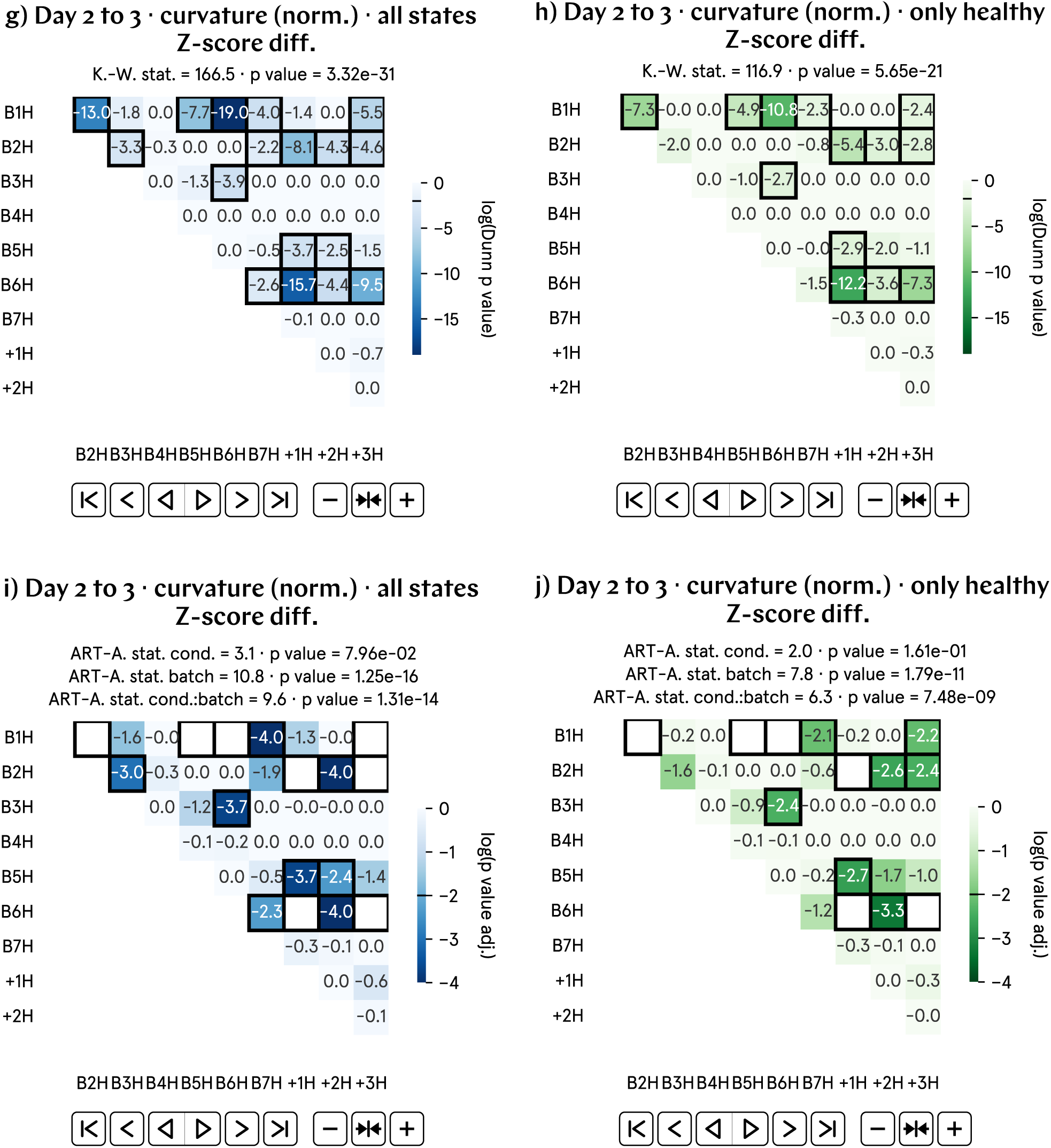
From day 2 to 3 p.f., the progression of the total curvature of hypomagnetic embryos, as measured by the Z-score difference, is approximately 0.4 higher than that of the control population. a) and b) Normalized total curvature Z-score difference by batch and aggregate data; B1–7 represent experimental runs, and +1–3 correspond to positive control runs, in which an artificial magnetic field inside the hypomagnetic chamber simulates Earth’s. c) and d) Analysis of the population averages and medians further support the observed 0.4 higher value. e) Cohen’s *d* and Cliff’s Δ confirm a medium effect. f) Binhi’s *E* and the visibility *ν* overestimate the effect size. g) through j) Most statistically significant differences are found in the positive control tadpole populations raised in the hypomagnetic chamber, as expected. Batches B1, B2 and B6 are also anomalous

### SI52. Day 2 to 3 curvature progression (norm.) results, visibility

**Fig. SI43:**
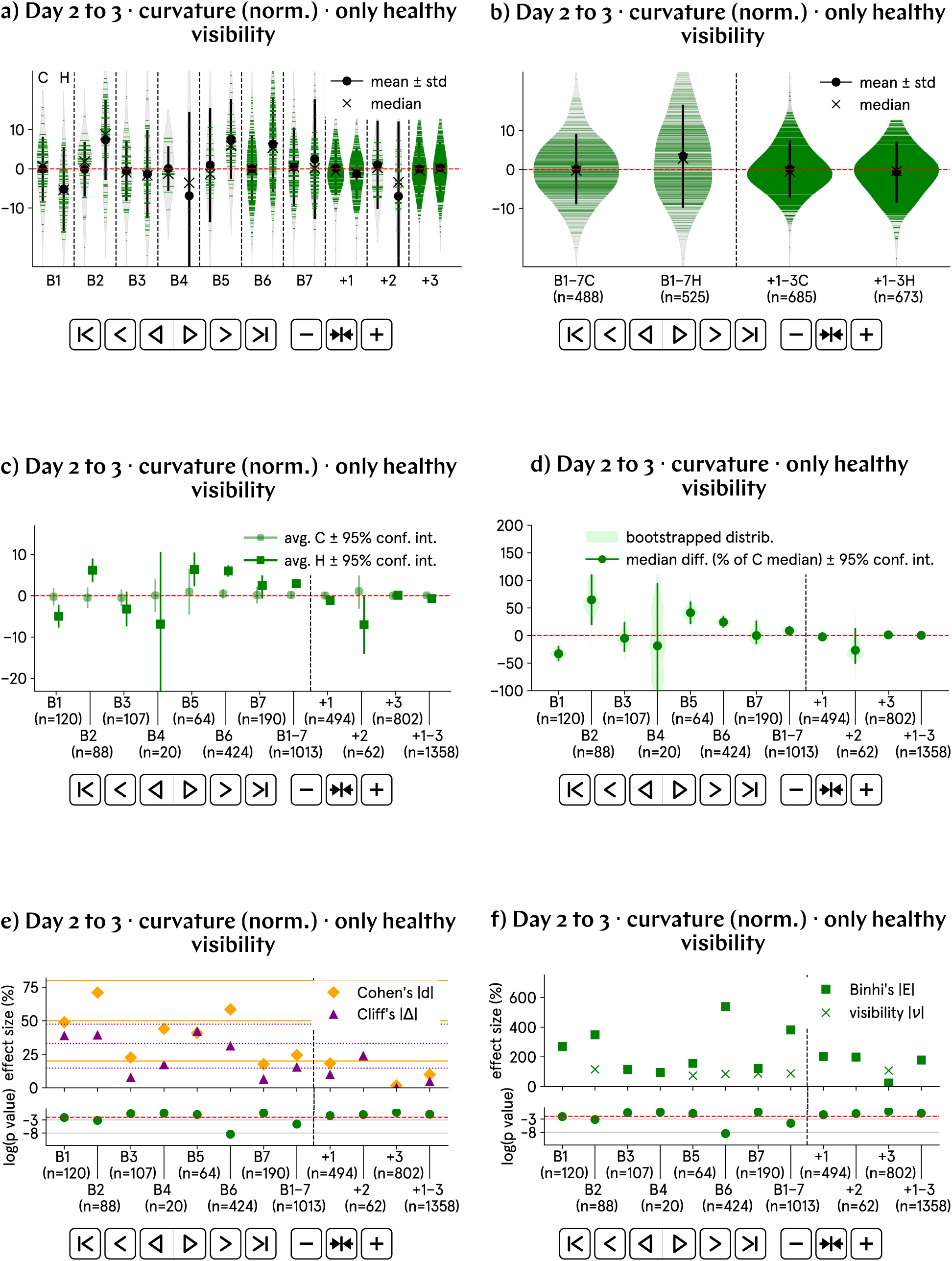
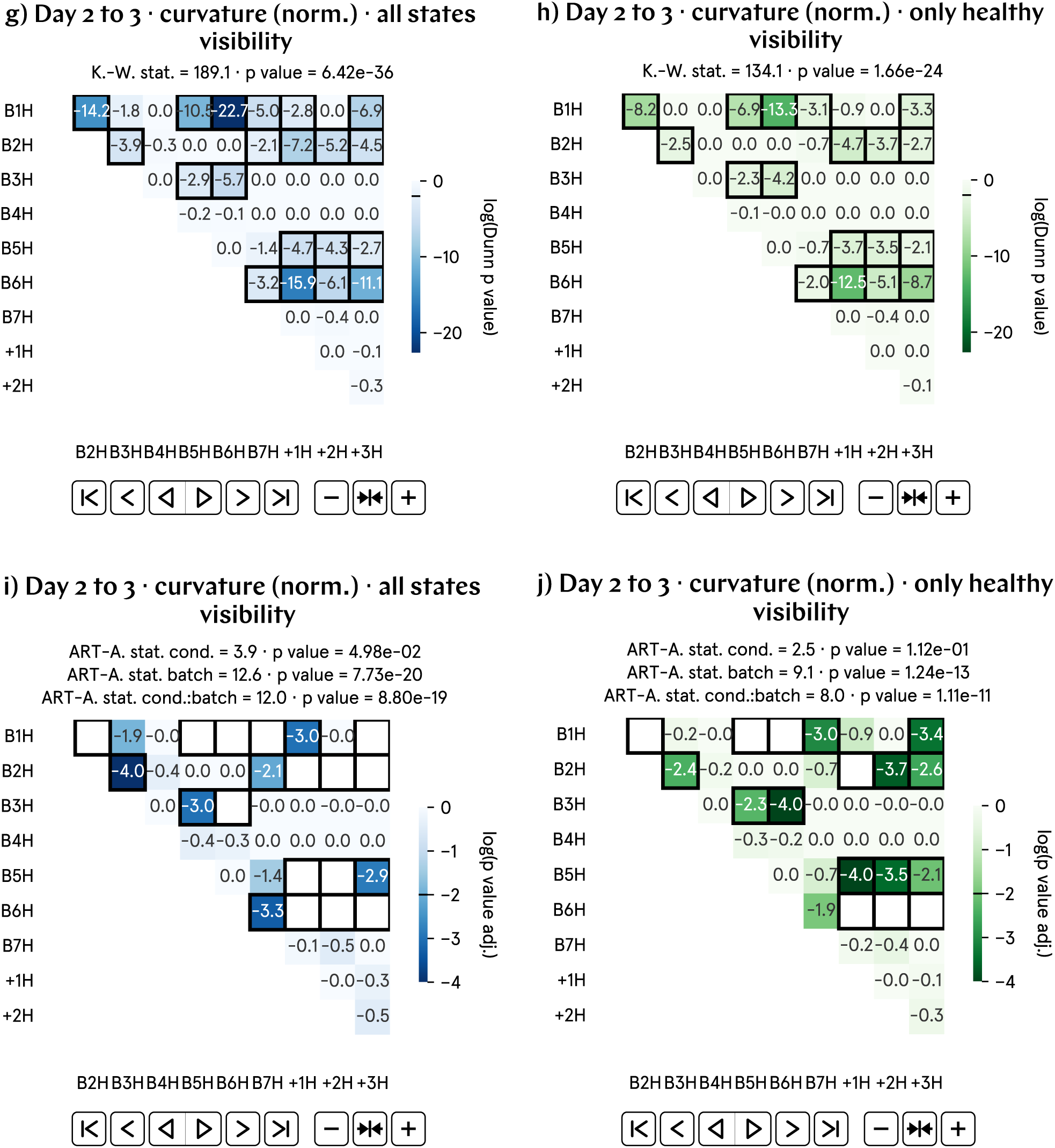
From day 2 to 3 p.f., the progression of the total curvature of hypomagnetic embryos, as measured by the visibility, is approximately 3% higher than that of the control population. a) and b) Normalized total curvature visibility by batch and aggregate data; B1–7 represent experimental runs, and +1–3 correspond to positive control runs, in which an artificial magnetic field inside the hypomagnetic chamber simulates Earth’s. c) and d) Analysis of the population averages and medians further support the observed 3% higher value. e) Cohen’s *d* and Cliff’s Δ confirm a medium effect. f) Binhi’s *E* and the visibility *ν* overestimate the effect size. g) through j) Most statistically significant differences are found in the positive control tadpole populations raised in the hypomagnetic chamber, as expected. Batches B1, B2 and B6 are also anomalous.

### SI53. Day 2 to 3 curvature progression (norm.) results, percentage change

**Fig. SI44:**
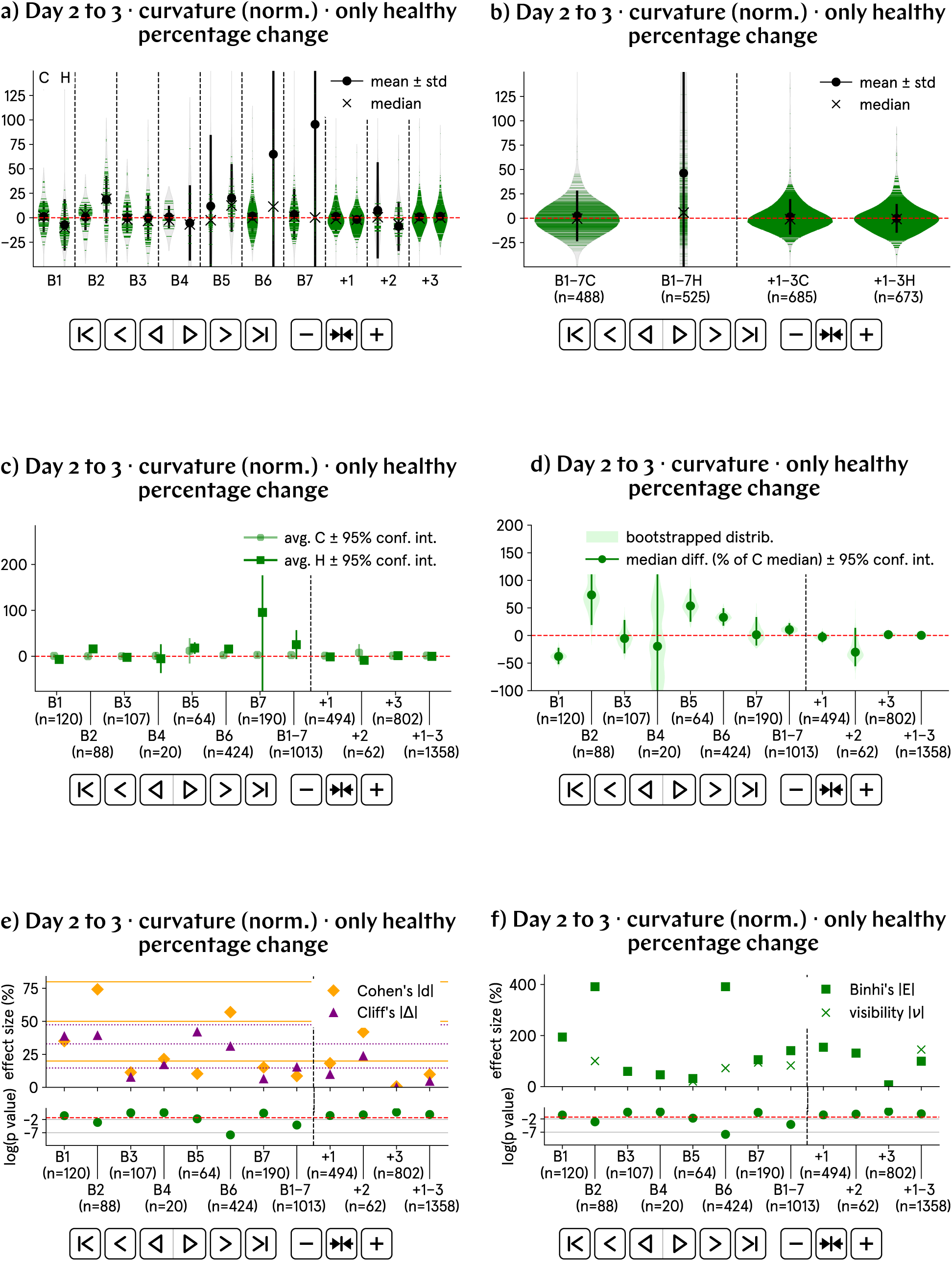
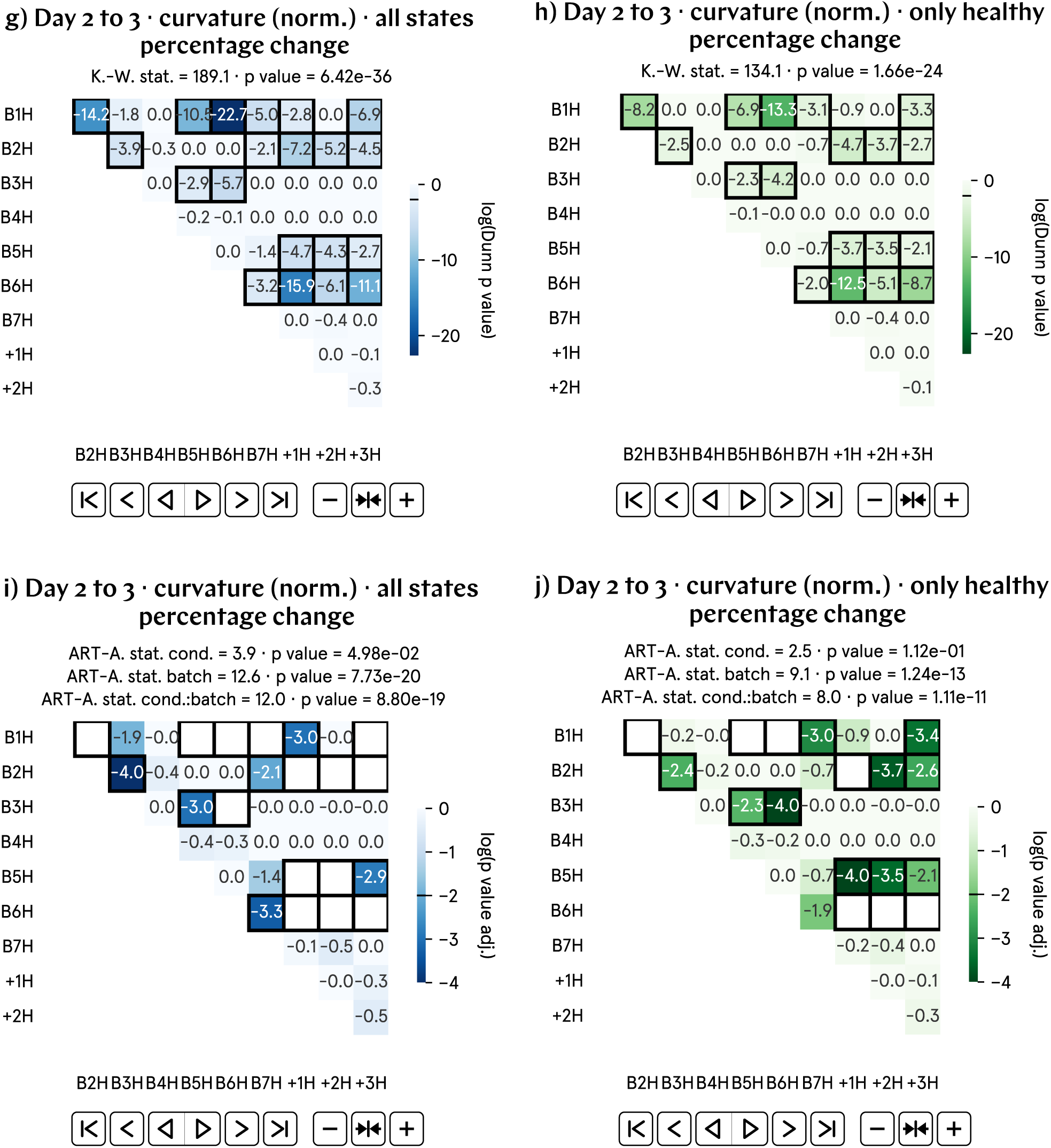
From day 2 to 3 p.f., the progression of the total curvature of hypomagnetic embryos, as measured by the percentage change, is approximately 25–50% higher than that of the control population–reflecting the limitations of this metric. a) and b) Normalized total curvature percentage change by batch and aggregate data. The highly anomalous violin in b) is a reflection of the limitations of this metric (due to calculations involving small denominators). c) and d) Analysis of the population averages and medians further support the observed 25% value. e) Cohen’s *d* and Cliff’s Δ confirm a medium effect. f) Binhi’s *E* and the visibility *ν* overestimate the effect size. g) through j) Most statistically significant differences are found in the positive control tadpole populations raised in the hypomagnetic chamber, as expected. Batches B1, B2 and B6 are also anomalous.

### SI54. Day 2 to 3 curvature progression (norm.) results, relative difference

**Fig. SI45:**
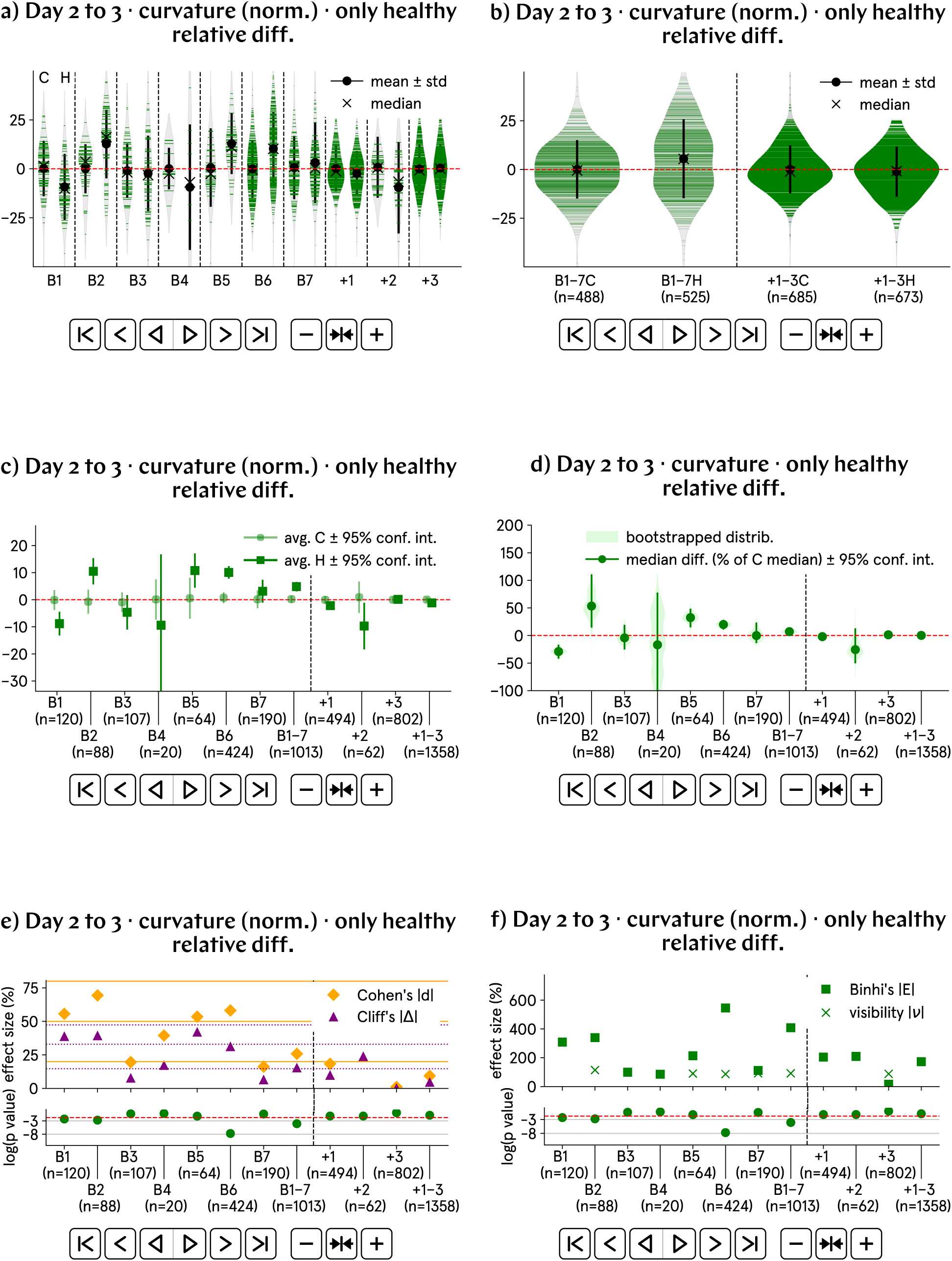
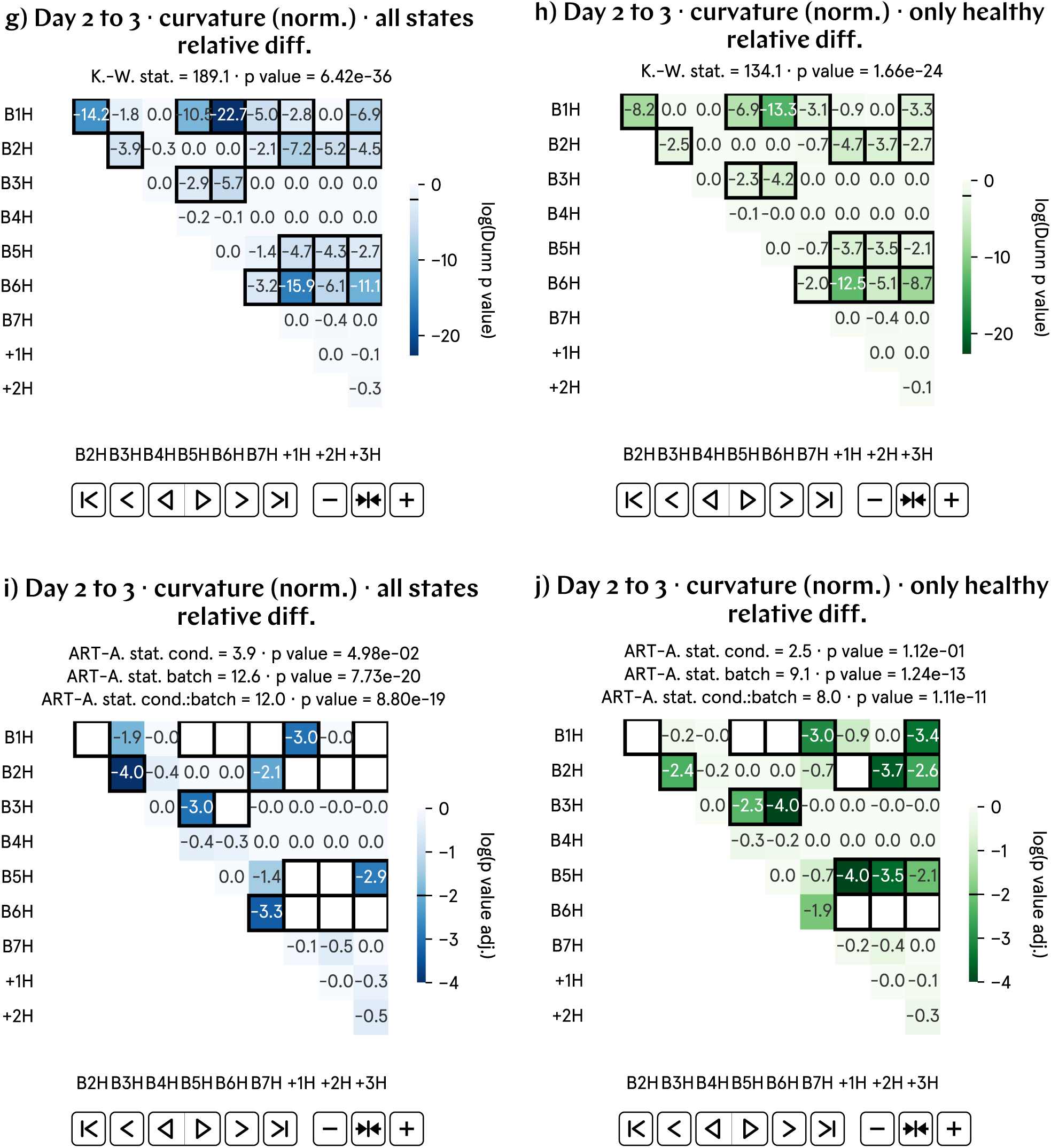
From day 2 to 3 p.f., the progression of the total curvature of hypomagnetic embryos, as measured by the relative difference, is approximately 5% higher than that of the control population. a) and b) Normalized total curvature relative difference by batch and aggregate data; B1–7 represent experimental runs, and +1–3 correspond to positive control runs, in which an artificial magnetic field inside the hypomagnetic chamber simulates Earth’s. c) and d) Analysis of the population averages and medians further support the observed 5% higher value. e) Cohen’s *d* and Cliff’s Δ confirm a medium effect. f) Binhi’s *E* and the visibility *ν* overestimate the effect size. g) through j) Most statistically significant differences are found in the positive control tadpole populations raised in the hypomagnetic chamber, as expected. Batches B1, B2 and B6 are also anomalous.

### SI55. Day 2 to 3 convexity progression (norm.) results, Z-score difference

**Fig. SI46:**
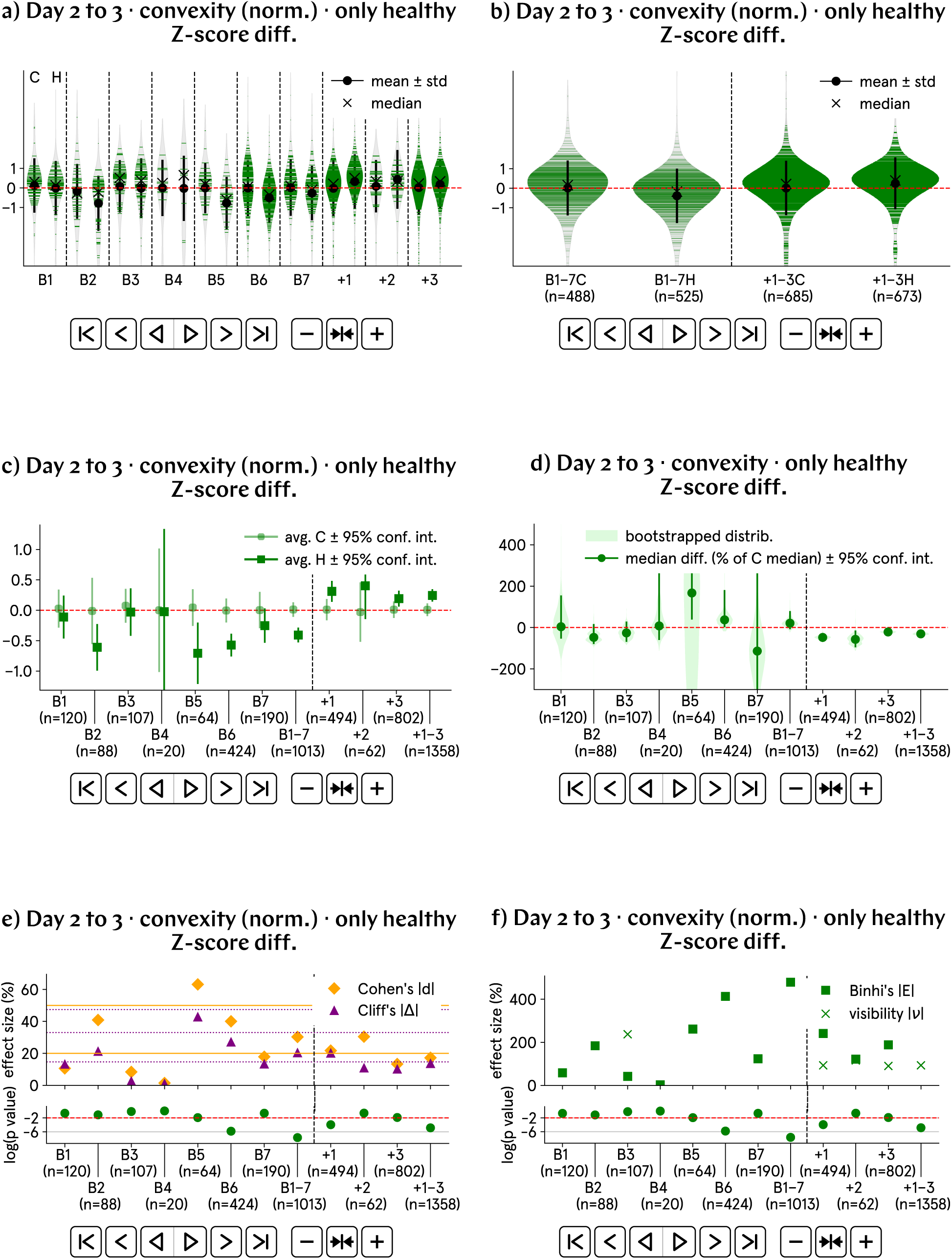
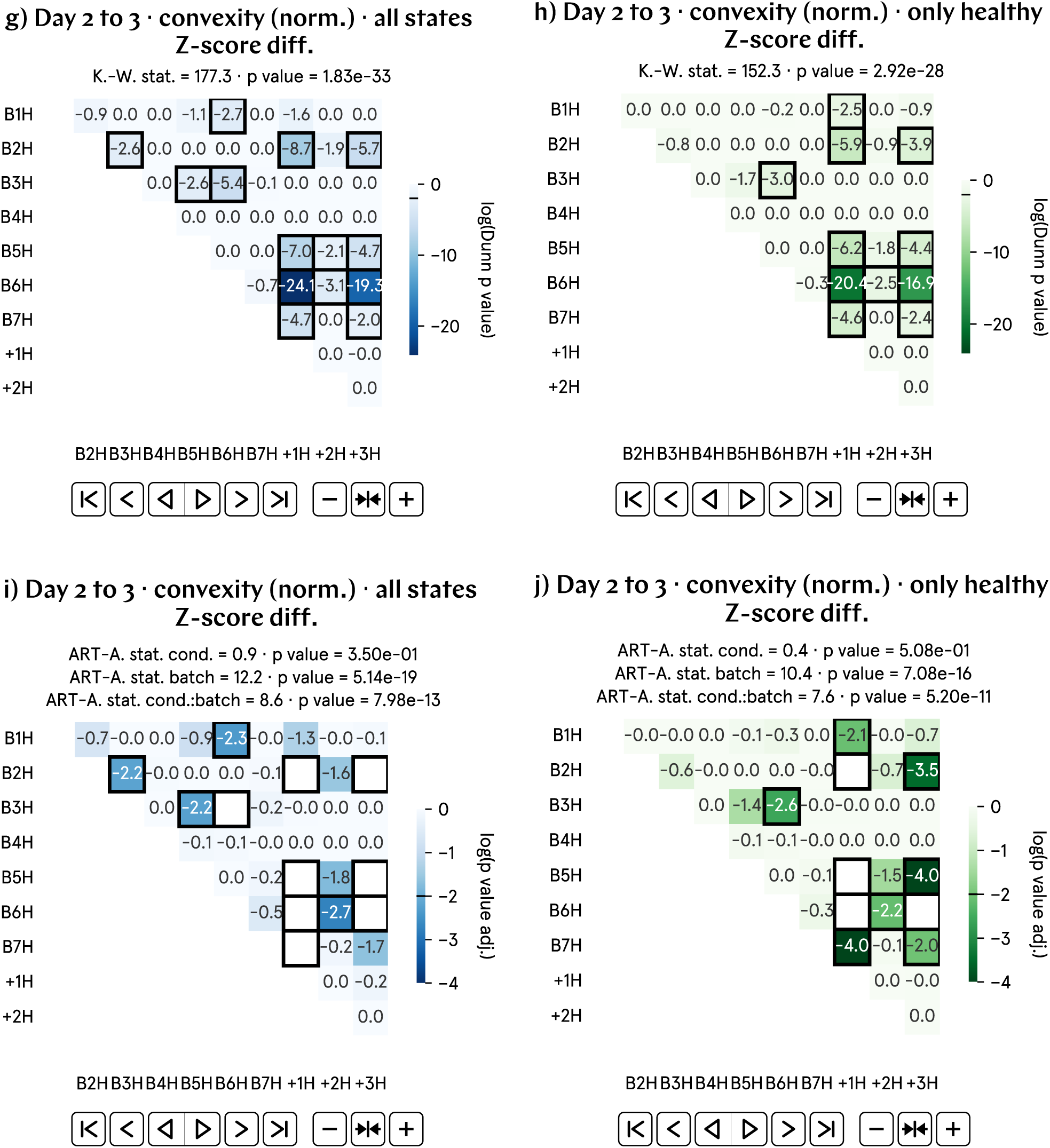
From day 2 to 3 p.f., the progression of the convexity of hypomagnetic embryos, as measured by the Z-score difference, is approximately 0.4 lower than that of the control population. a) and b) Normalized convexity Z-score difference by batch and aggregate data; B1–7 represent experimental runs, and +1–3 correspond to positive control runs, in which an artificial magnetic field inside the hypomagnetic chamber simulates Earth’s. c) and d) Analysis of the population averages and medians further support the observed 0.4 lower value. e) Cohen’s *d* and Cliff’s Δ confirm a medium effect. f) Binhi’s *E* and the visibility *ν* overestimate the effect size. g) through j) Most statistically significant differences are found in the positive control tadpole populations raised in the hypomagnetic chamber, as expected.

### SI56. Day 2 to 3 convexity progression (norm.) results, visibility

**Fig. SI47:**
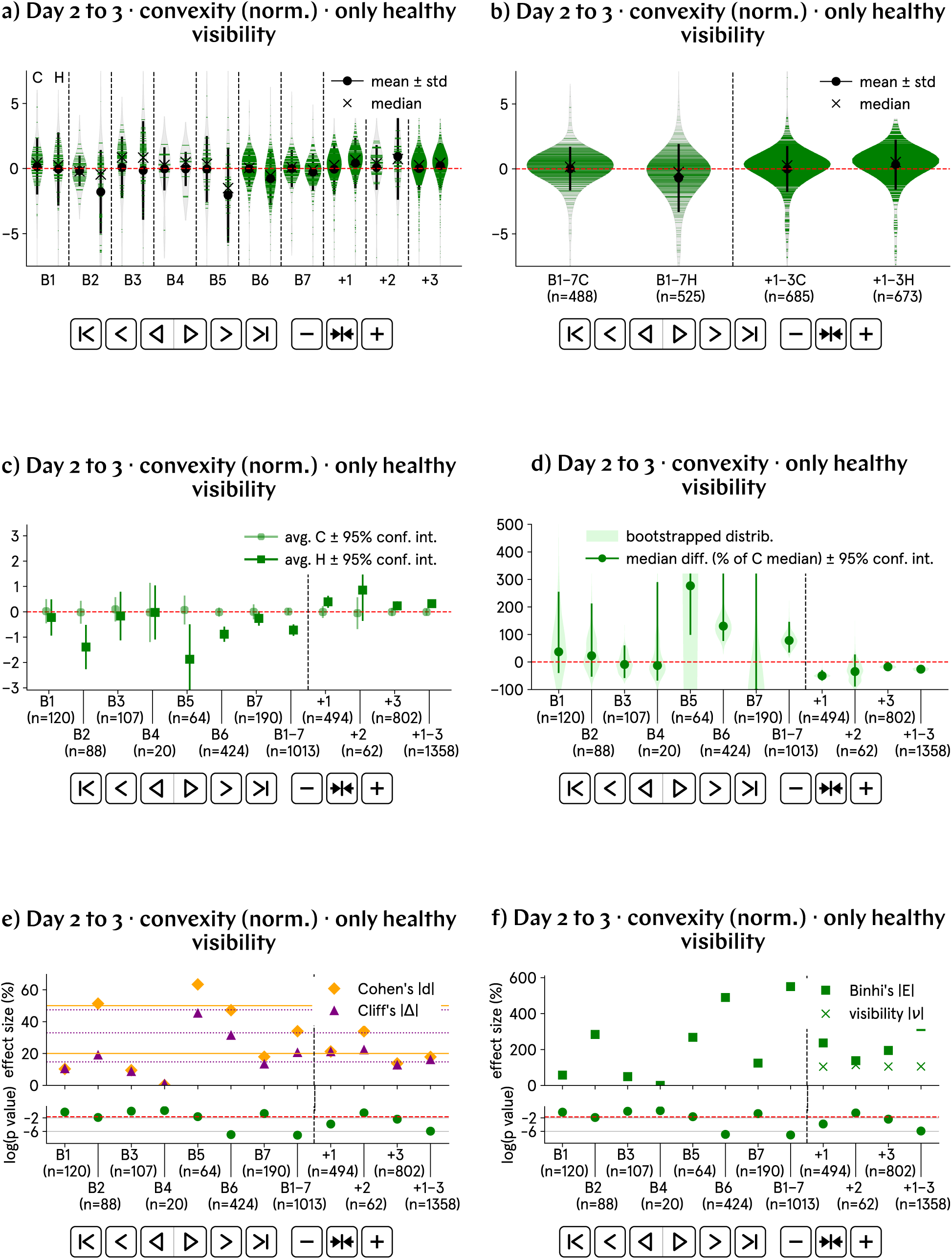
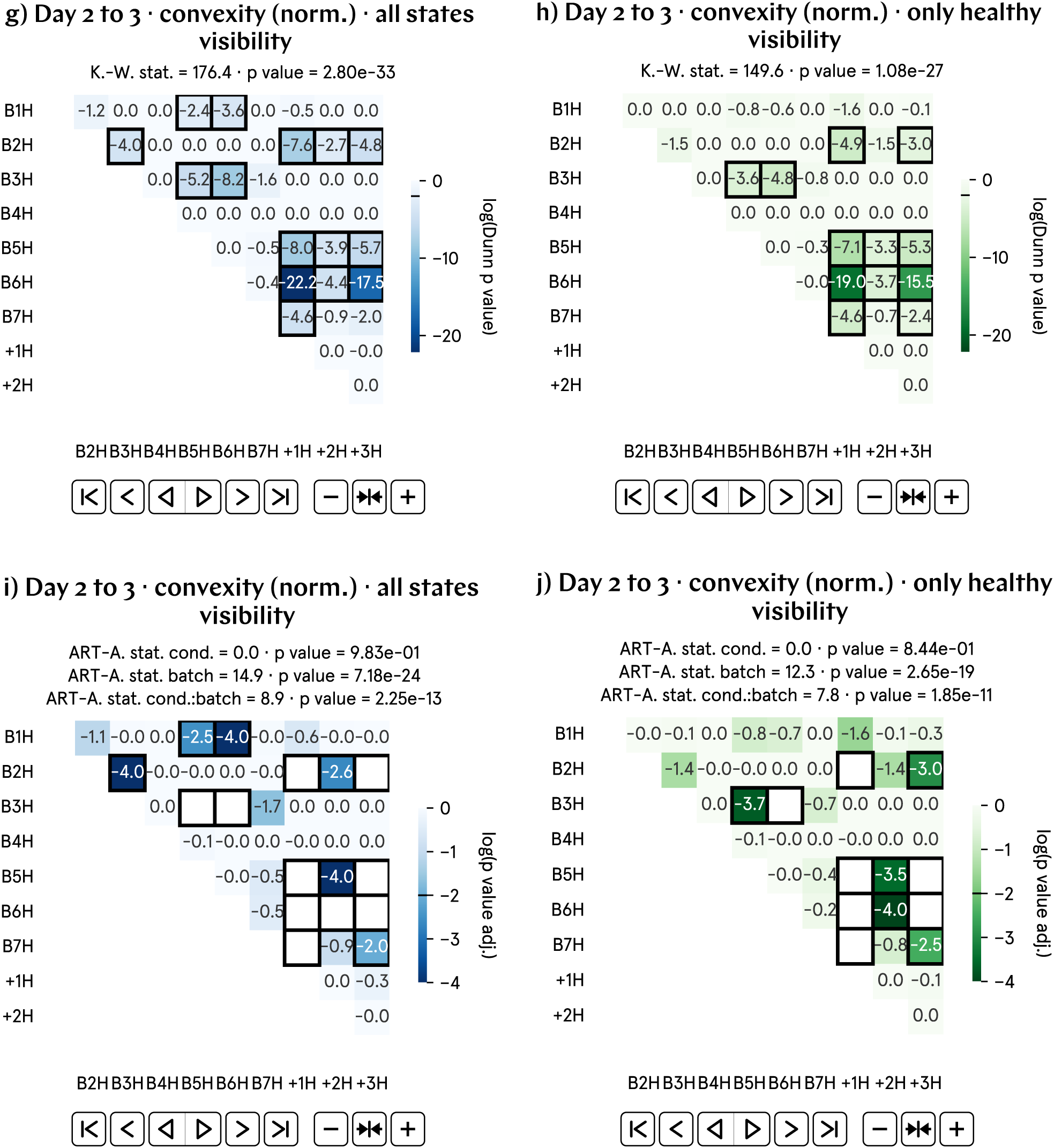
From day 2 to 3 p.f., the progression of the convexity of hypomagnetic embryos, as measured by the visibility, is approximately 1% lower than that of the control population. a) and b) Normalized convexity visibility by batch and aggregate data; B1–7 represent experimental runs, and +1–3 correspond to positive control runs, in which an artificial magnetic field inside the hypomagnetic chamber simulates Earth’s. c) and d) Analysis of the population averages and medians further support the observed 1% lower value. e) Cohen’s *d* and Cliff’s Δ confirm a medium effect. f) Binhi’s *E* and the visibility *ν* overestimate the effect size. g) through j) Most statistically significant differences are found in the positive control tadpole populations raised in the hypomagnetic chamber, as expected.

### SI57. Day 2 to 3 convexity progression (norm.) results, percentage change

**Fig. SI48:**
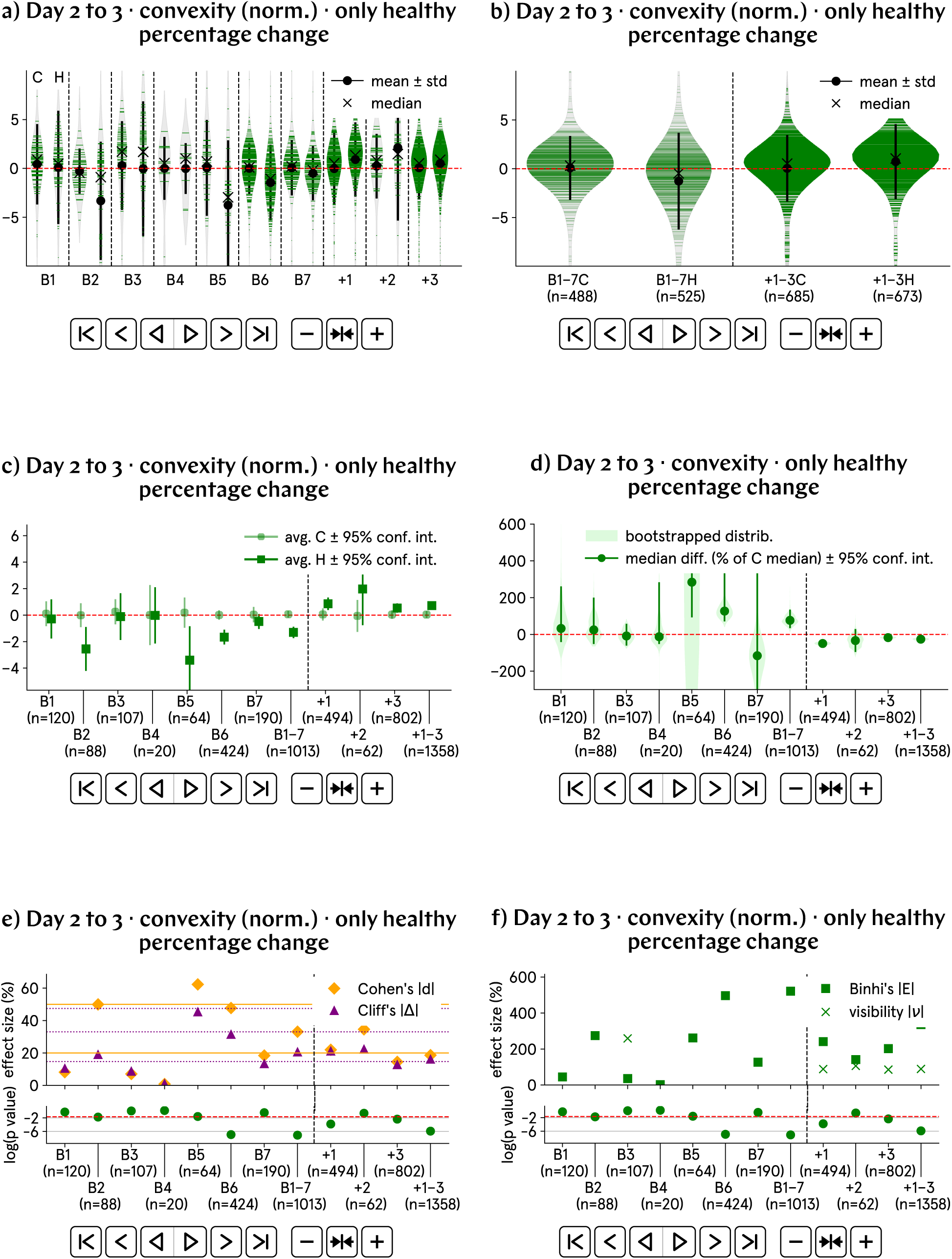
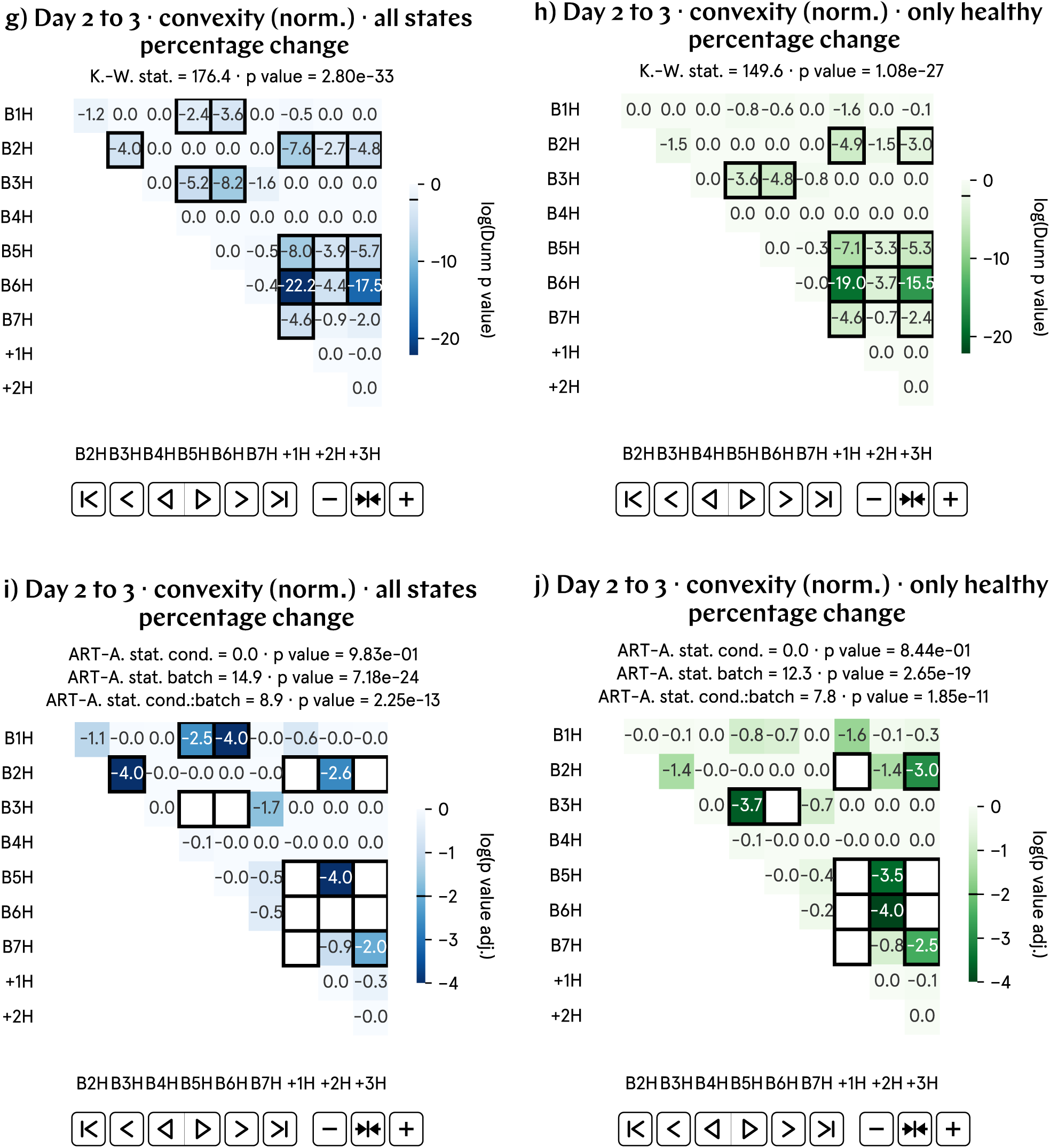
From day 2 to 3 p.f., the progression of the convexity of hypomagnetic embryos, as measured by the percentage change, is approximately 2% lower than that of the control population. a) and b) Normalized convexity percentage change by batch and aggregate data; B1–7 represent experimental runs, and +1–3 correspond to positive control runs, in which an artificial magnetic field inside the hypomagnetic chamber simulates Earth’s. c) and d) Analysis of the population averages and medians further support the observed 2% lower value. e) Cohen’s *d* and Cliff’s Δ confirm a medium effect. f) Binhi’s *E* and the visibility *ν* overestimate the effect size. g) through j) Most statistically significant differences are found in the positive control tadpole populations raised in the hypomagnetic chamber, as expected.

### SI58. Day 2 to 3 convexity progression (norm.) results, relative difference

**Fig. SI49:**
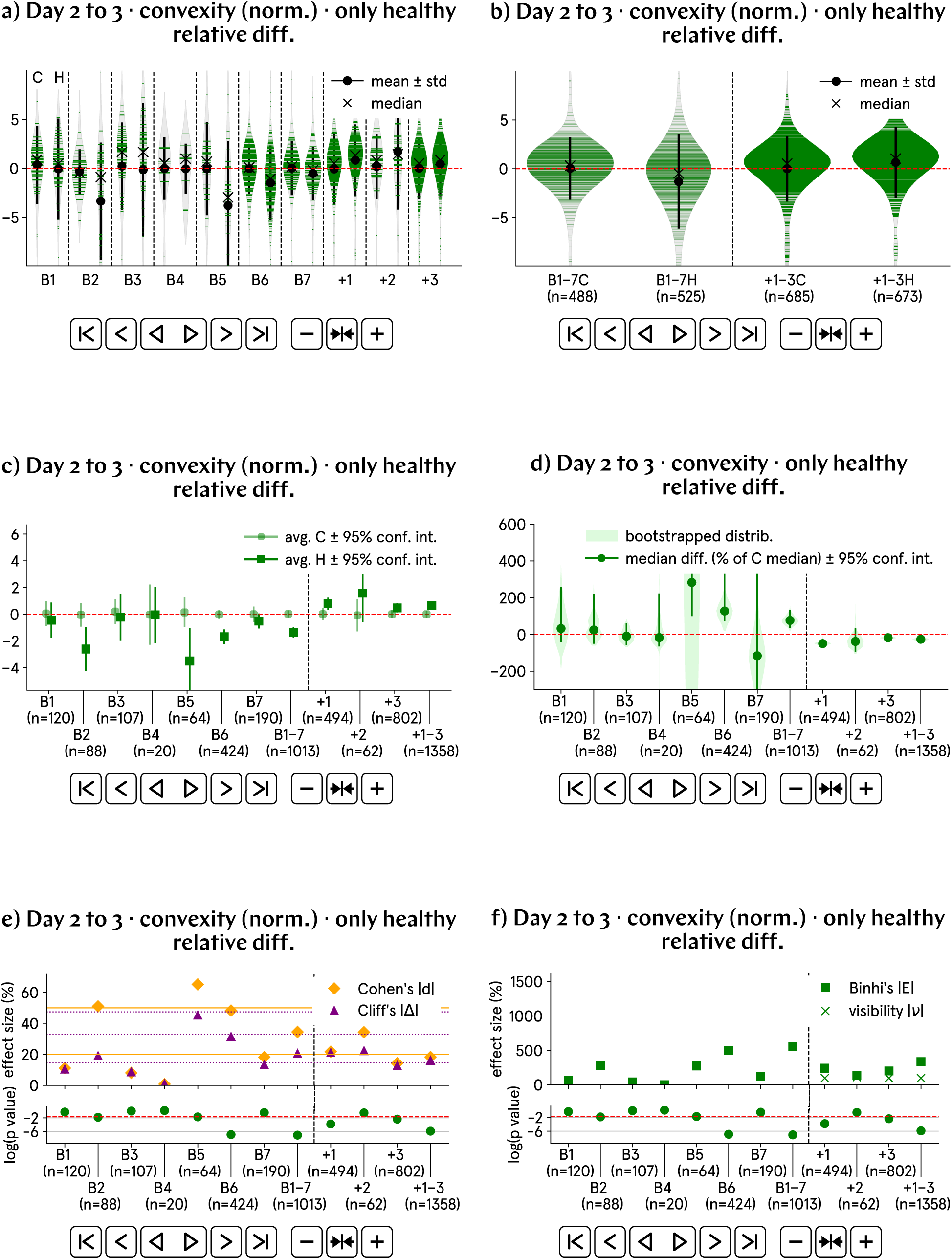
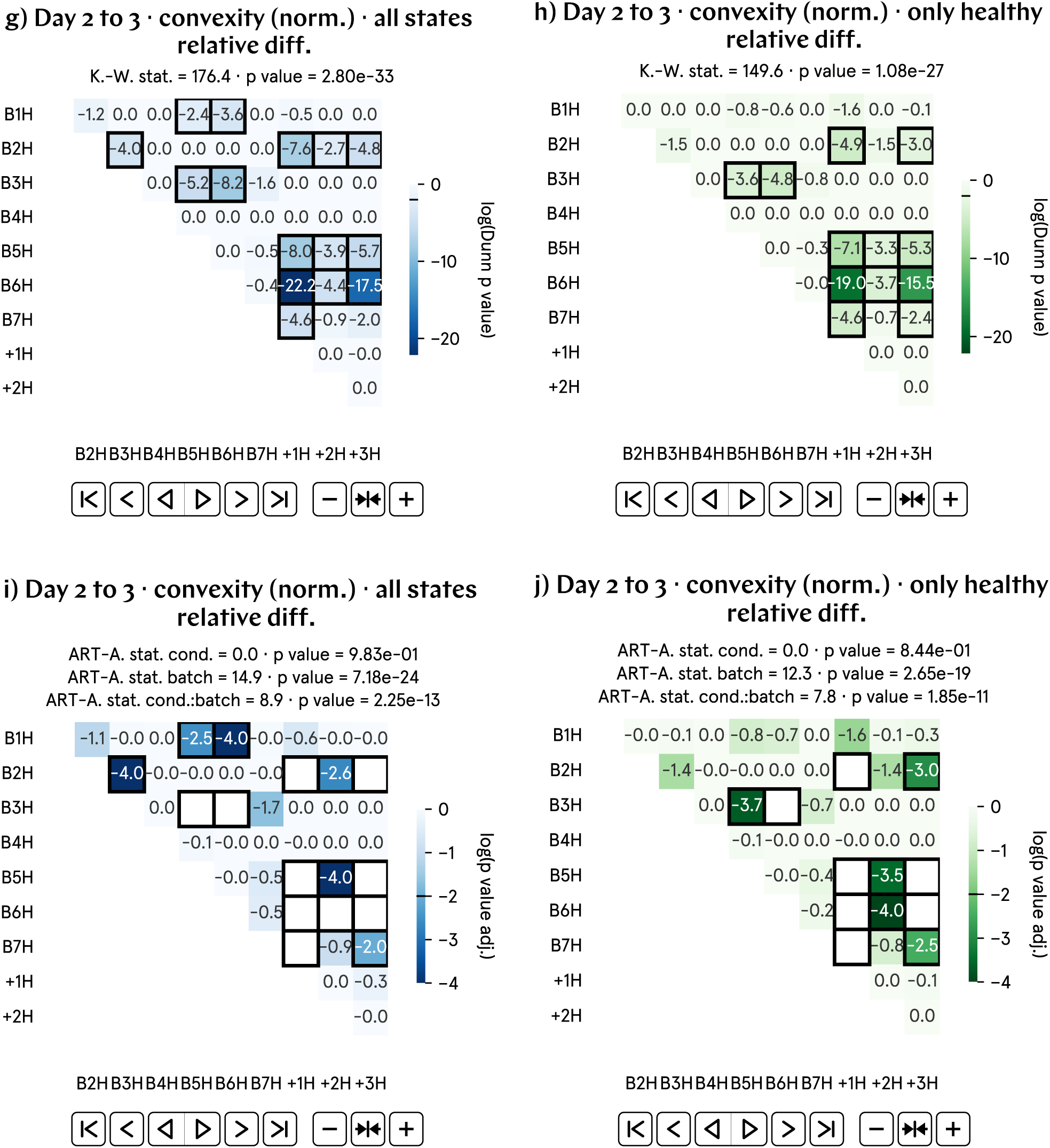
From day 2 to 3 p.f., the progression of the convexity of hypomagnetic embryos, as measured by the relative difference, is approximately 2% lower than that of the control population. a) and b) Normalized convexity relative difference by batch and aggregate data; B1–7 represent experimental runs, and +1–3 correspond to positive control runs, in which an artificial magnetic field inside the hypomagnetic chamber simulates Earth’s. c) and d) Analysis of the population averages and medians further support the observed 2% lower value. e) Cohen’s *d* and Cliff’s Δ confirm a medium effect. f) Binhi’s *E* and the visibility *ν* overestimate the effect size. g) through j) Most statistically significant differences are found in the positive control tadpole populations raised in the hypomagnetic chamber, as expected.

### SI59. Day 2 to 3 solidity progression (norm.) results, Z-score difference

**Fig. SI50:**
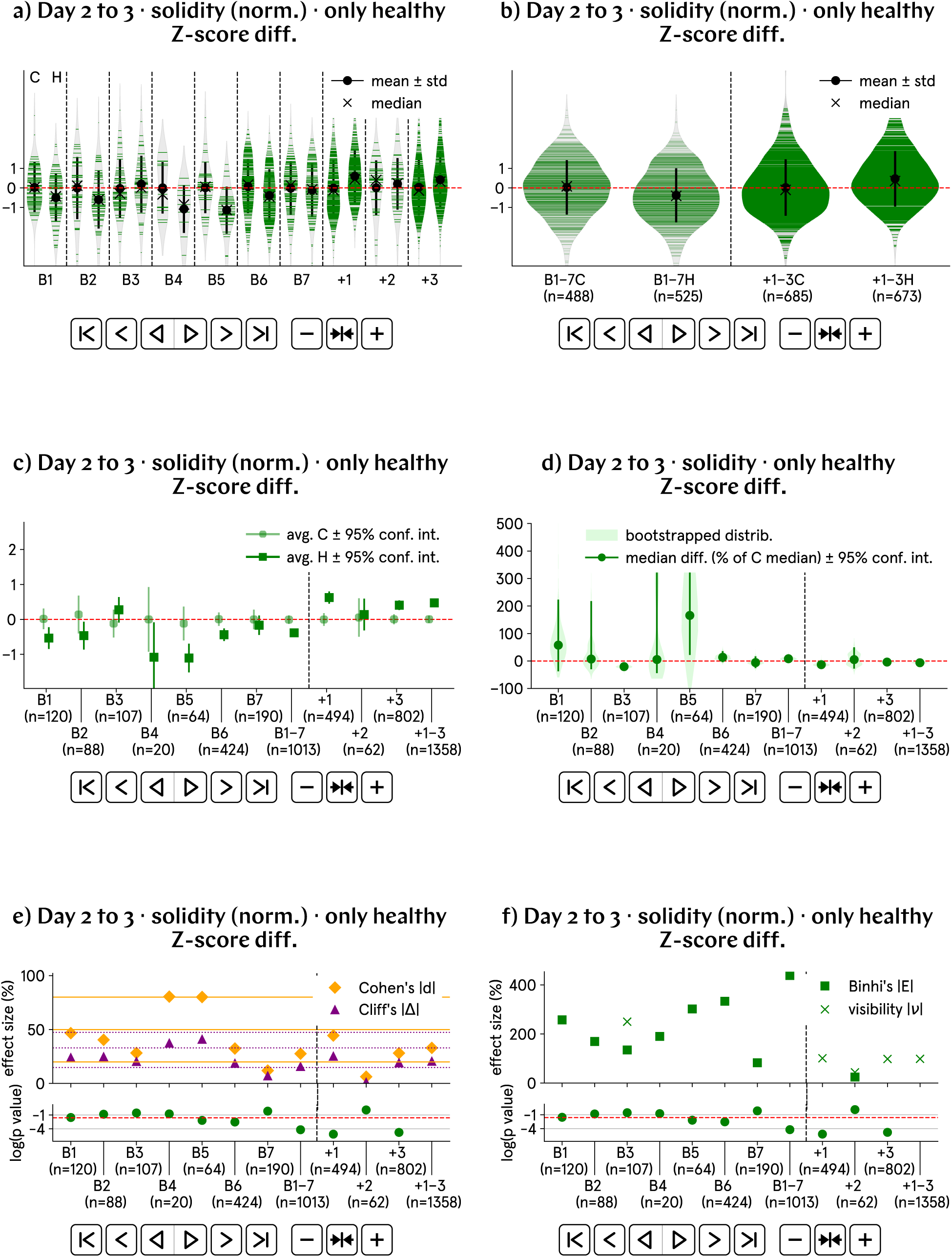
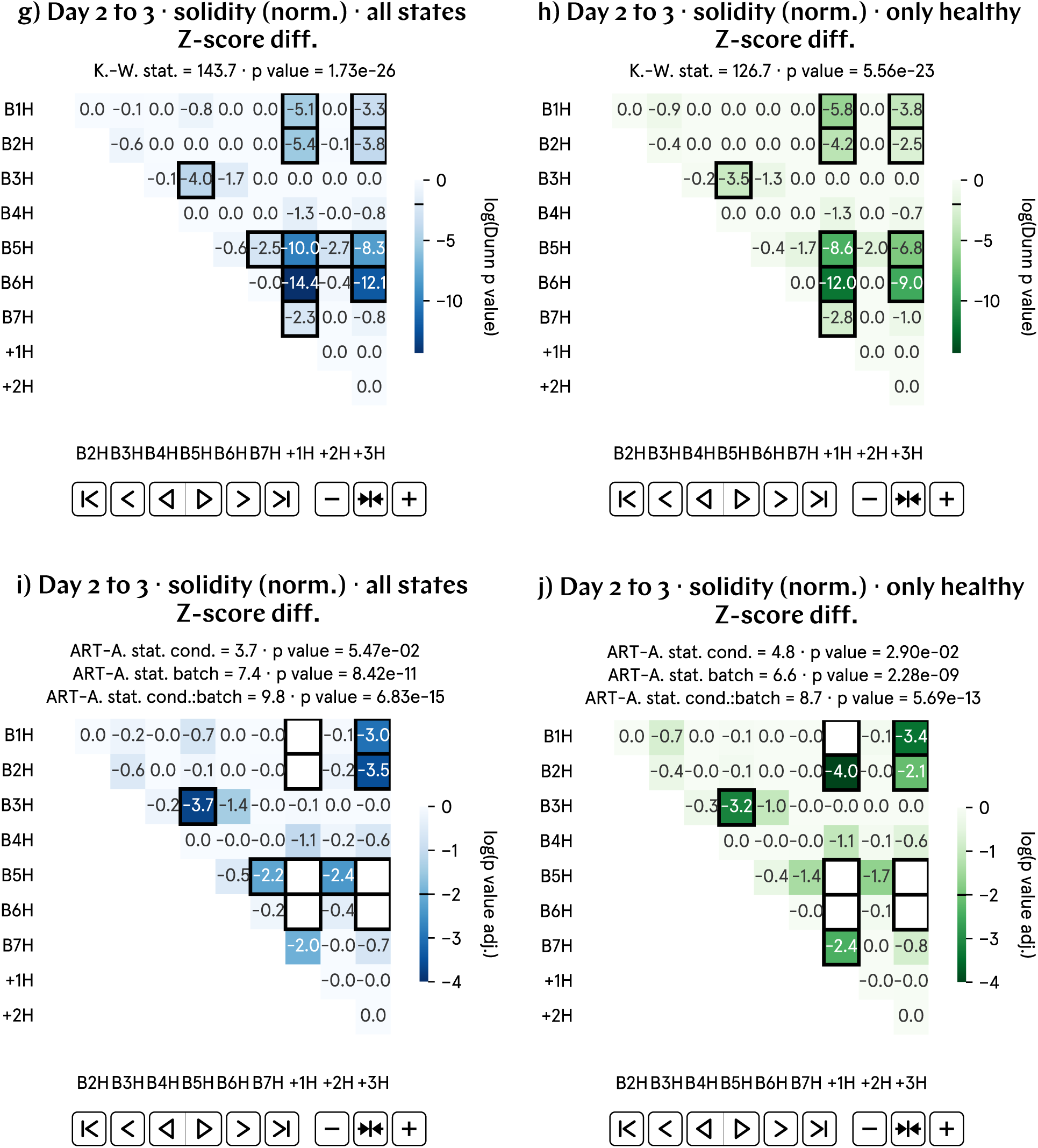
From day 2 to 3 p.f., the progression of the solidity of hypomagnetic embryos, as measured by the Z-score difference, is approximately 0.4 lower than that of the control population. a) and b) Normalized solidity Z-score difference by batch and aggregate data; B1–7 represent experimental runs, and +1–3 correspond to positive control runs, in which an artificial magnetic field inside the hypomagnetic chamber simulates Earth’s. c) and d) Analysis of the population averages and medians further support the observed 0.4 lower value. e) Cohen’s *d* and Cliff’s Δ confirm a medium effect. f) Binhi’s *E* and the visibility *ν* overestimate the effect size. g) through j) Most statistically significant differences are found in the positive control tadpole populations raised in the hypomagnetic chamber, as expected.

### SI60. Day 2 to 3 solidity progression (norm.) results, visibility

**Fig. SI51:**
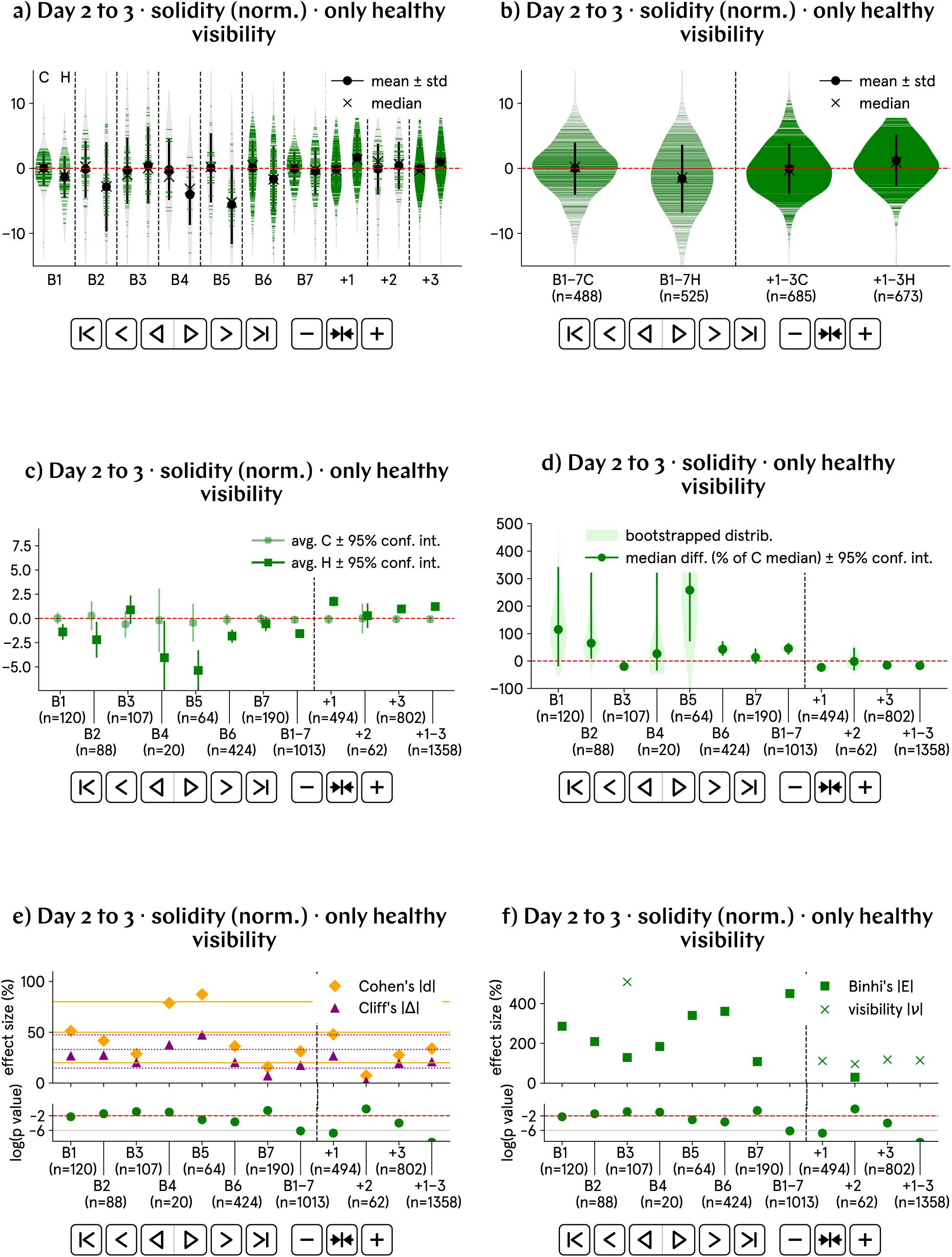
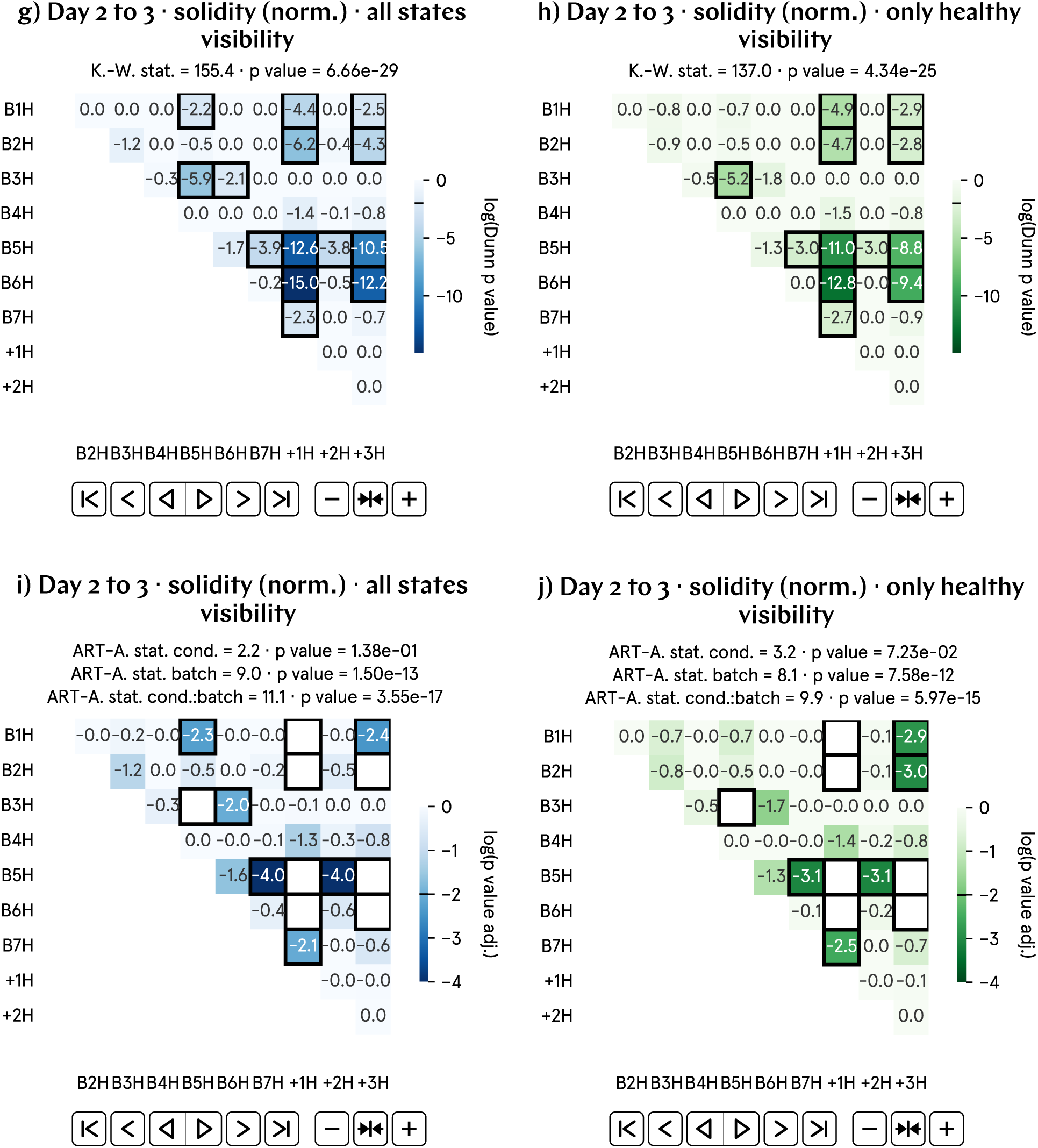
From day 2 to 3 p.f., the progression of the solidity of hypomagnetic embryos, as measured by the visibility, is approximately 2% lower than that of the control population. a) and b) Normalized solidity visibility by batch and aggregate data; B1–7 represent experimental runs, and +1–3 correspond to positive control runs, in which an artificial magnetic field inside the hypomagnetic chamber simulates Earth’s. c) and d) Analysis of the population averages and medians further support the observed 2% lower value. e) Cohen’s *d* and Cliff’s Δ confirm a medium effect. f) Binhi’s *E* and the visibility *ν* overestimate the effect size. g) through j) Most statistically significant differences are found in the positive control tadpole populations raised in the hypomagnetic chamber, as expected.

### SI61. Day 2 to 3 solidity progression (norm.) results, percentage change

**Fig. SI52:**
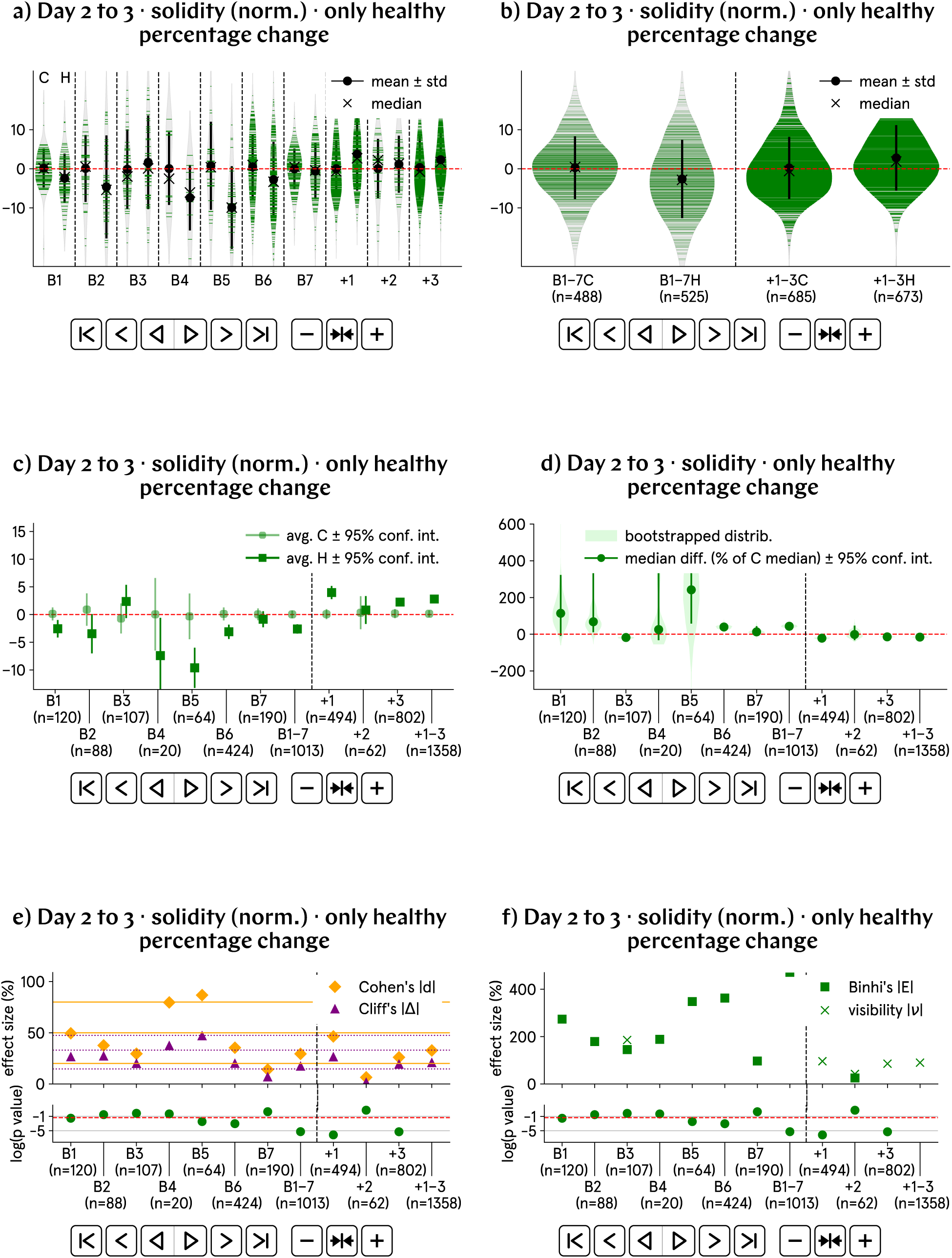
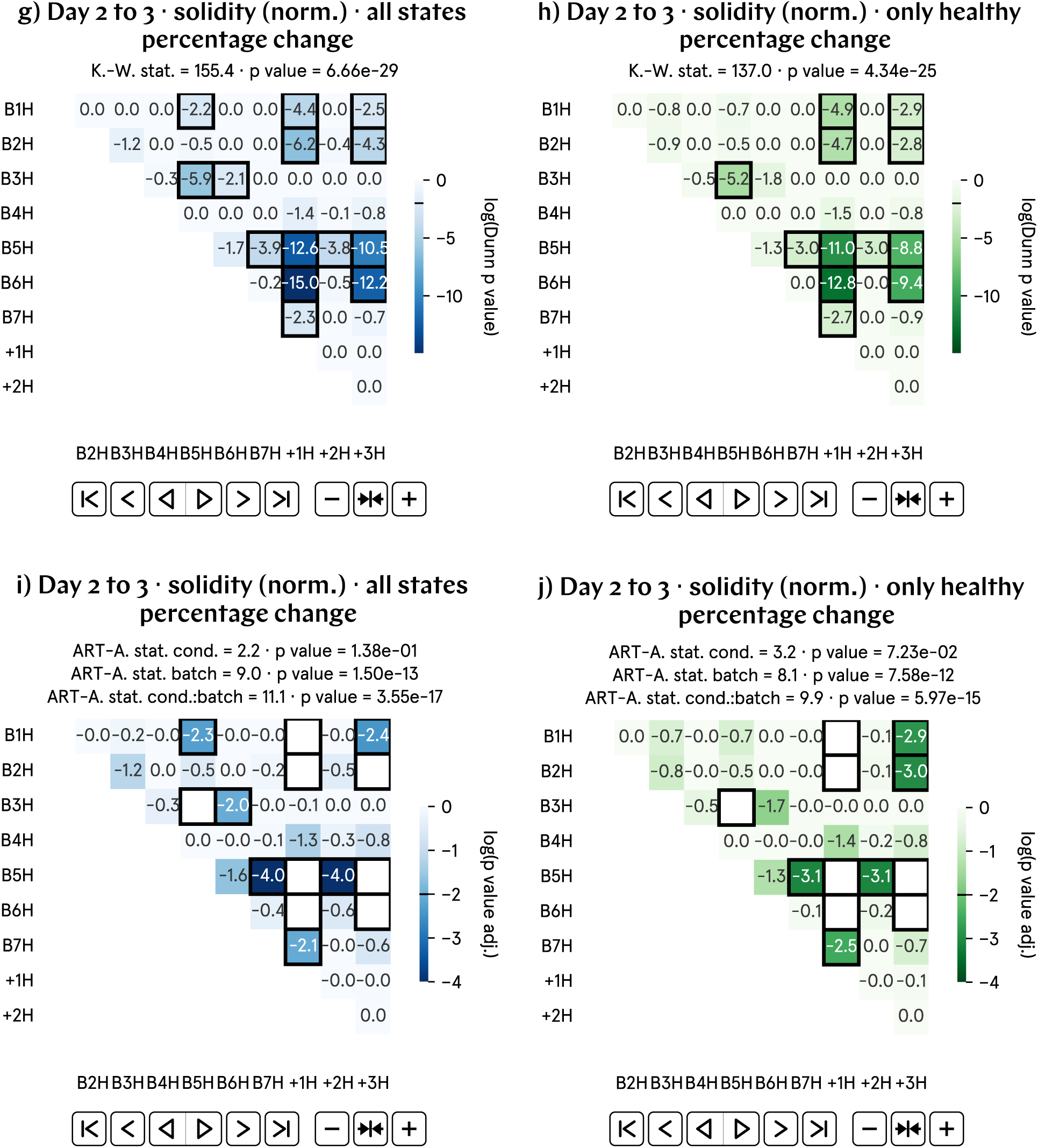
From day 2 to 3 p.f., the progression of the solidity of hypomagnetic embryos, as measured by the percentage change, is approximately 2% lower than that of the control population. a) and b) Normalized solidity percentage change by batch and aggregate data; B1–7 represent experimental runs, and +1–3 correspond to positive control runs, in which an artificial magnetic field inside the hypomagnetic chamber simulates Earth’s. c) and d) Analysis of the population averages and medians further support the observed 2% lower value. e) Cohen’s *d* and Cliff’s Δ confirm a medium effect. f) Binhi’s *E* and the visibility *ν* overestimate the effect size. g) through j) Most statistically significant differences are found in the positive control tadpole populations raised in the hypomagnetic chamber, as expected.

### SI62. Day 2 to 3 solidity progression (norm.) results, relative difference

**Fig. SI53:**
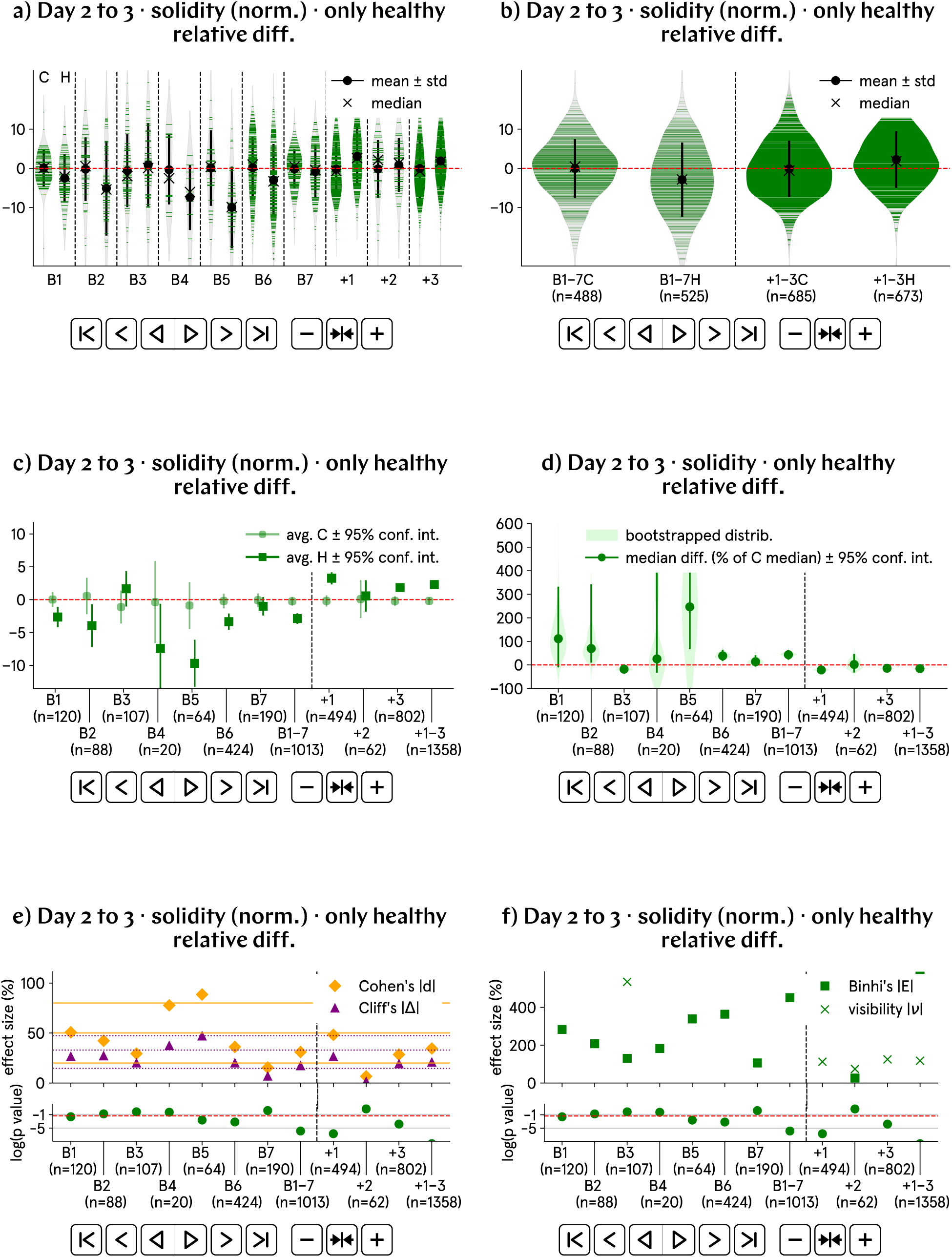
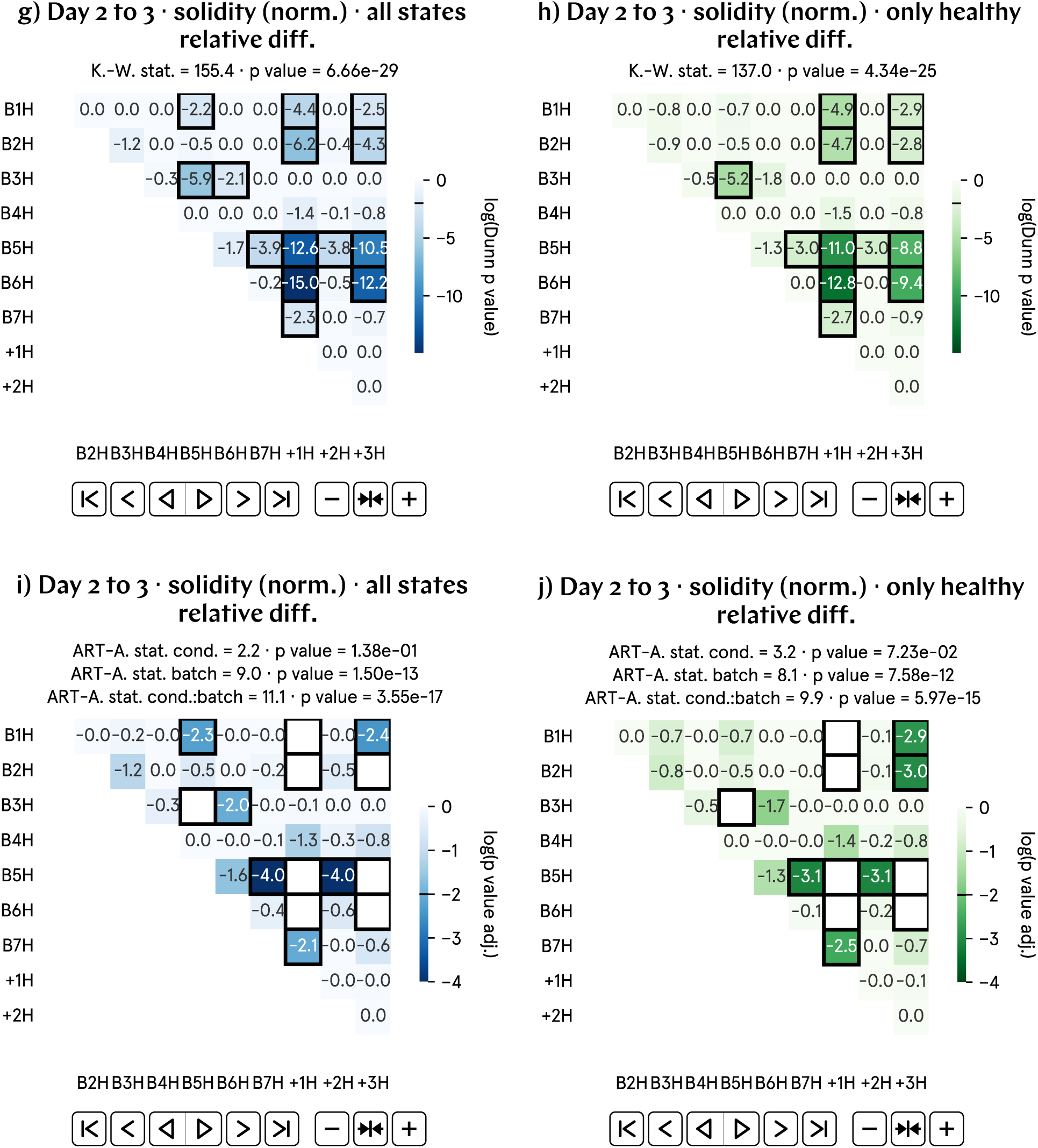
From day 2 to 3 p.f., the progression of the solidity of hypomagnetic embryos, as measured by the relative difference, is approximately 3% lower than that of the control population. a) and b) Normalized solidity relative difference by batch and aggregate data; B1–7 represent experimental runs, and +1–3 correspond to positive control runs, in which an artificial magnetic field inside the hypomagnetic chamber simulates Earth’s. c) and d) Analysis of the population averages and medians further support the observed 3% lower value. e) Cohen’s *d* and Cliff’s Δ confirm a medium effect. f) Binhi’s *E* and the visibility *ν* overestimate the effect size. g) through j) Most statistically significant differences are found in the positive control tadpole populations raised in the hypomagnetic chamber, as expected.

### SI63. Additional morphological measures

For completeness, we display here our results for the normalized perimeter in day 1 and 2 images (respectively, in Supplementary Figs. SI54 and SI57); both are overall slightly larger in hypomagnetic tadpoles. The trend persists in the day 3 images, where the previous analysis revealed a significantly larger perimeter for hypomagnetic tadpoles (see Supplementary Fig. SI29).

In day 2 images, the major axis of hypomagnetic tadpoles is only marginally longer (as depicted in Supplementary Fig. SI56), whereas in day 3 images, the major axis is significantly extended for hypomagnetic tadpoles (see Section 6). We hypothesize that the segmentation of day 2 images, with tadpoles with less pigmentation, is less robust and a worse predictor of true tadpole length. In addition, the imaging conditions on day 2 were less controlled, as the live tadpoles were not manipulated, potentially resulting in distortions or tadpoles being out of the focal plane. In contrast, on day 3, the fixed tadpoles were carefully positioned with tweezers to lie flat on their sides, providing more reliable measurements.

There was no statistical impact from the tadpoles’ magnetic field environment over the area for day 1 and 2 images; detailed analyses can be found in Supplementary Figs. SI55 and SI58. Segmentation for day 1 images is highly robust, while for day 2 images, it may be less reliable, potentially obscuring the true area measurements.

We quantitatively compare our findings to the only study we found reporting size variability in *Xenopus* embryos [14]. In their analysis of embryos between NF stages 10 and 15 [15], they demonstrate that natural size variations are compensated by a scaling mechanism in gene expression, where larger embryos exhibit proportionally larger gene expression domains, leading to proper tissue and organ development despite size differences. In their analysis of over 2,000 early-stage embryos, they report a maximum 38% difference in major axis length. As shown in the a) panel of Supplementary Fig. SI59, the day 1 p.f. images from our dataset, which align closely with the stages they studied, exhibit a very similar variability in major axis length for control embryos, calculated in our case as the difference between the averages of the largest and smallest 5%. Specifically, for the control embryos in the aggregated data B1–7, this variability closely matches the 38% reported (denoted with a red dashed line); in contrast, hypomagnetic embryos display significantly greater variability (approximately 20% higher). In the b) and c) panels of Supplementary Fig. SI59, we plot the major axis variability for day 2 and 3 p.f. images in our dataset. We consistently find a higher variability in the hypomagnetic tadpole population; this variability is much reduced in tadpoles during the positive control runs. The comparison with the data from [14] in b) and c) is an extrapolation, as the persistence of variability at similar magnitudes in later-stage embryos remains unquantified by any other study than the present one. For all points in these plots, full error propagation was done automatically using the python package unumpy.

### SI64. Day 1 perimeter (norm.) results

**Fig. SI54:**
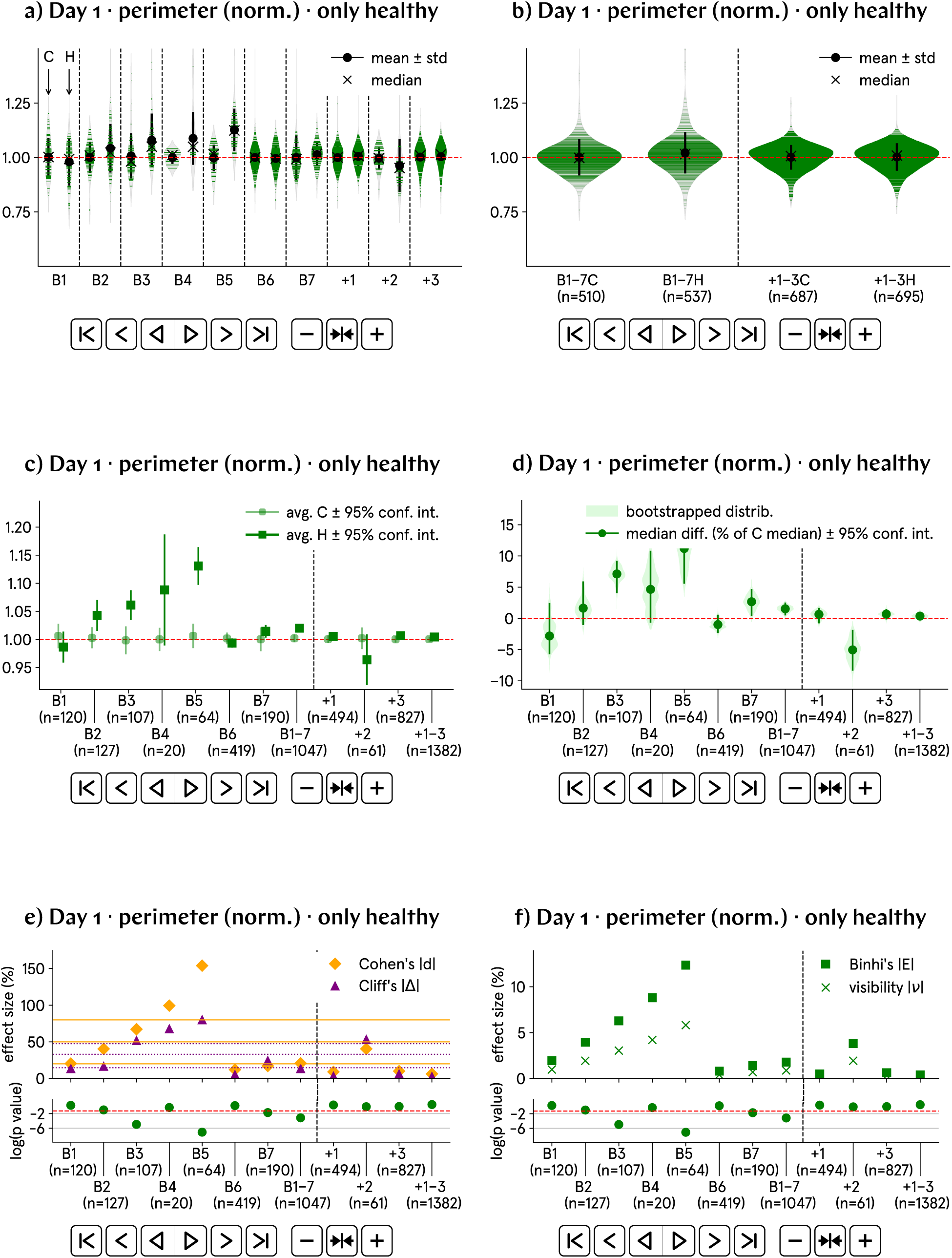
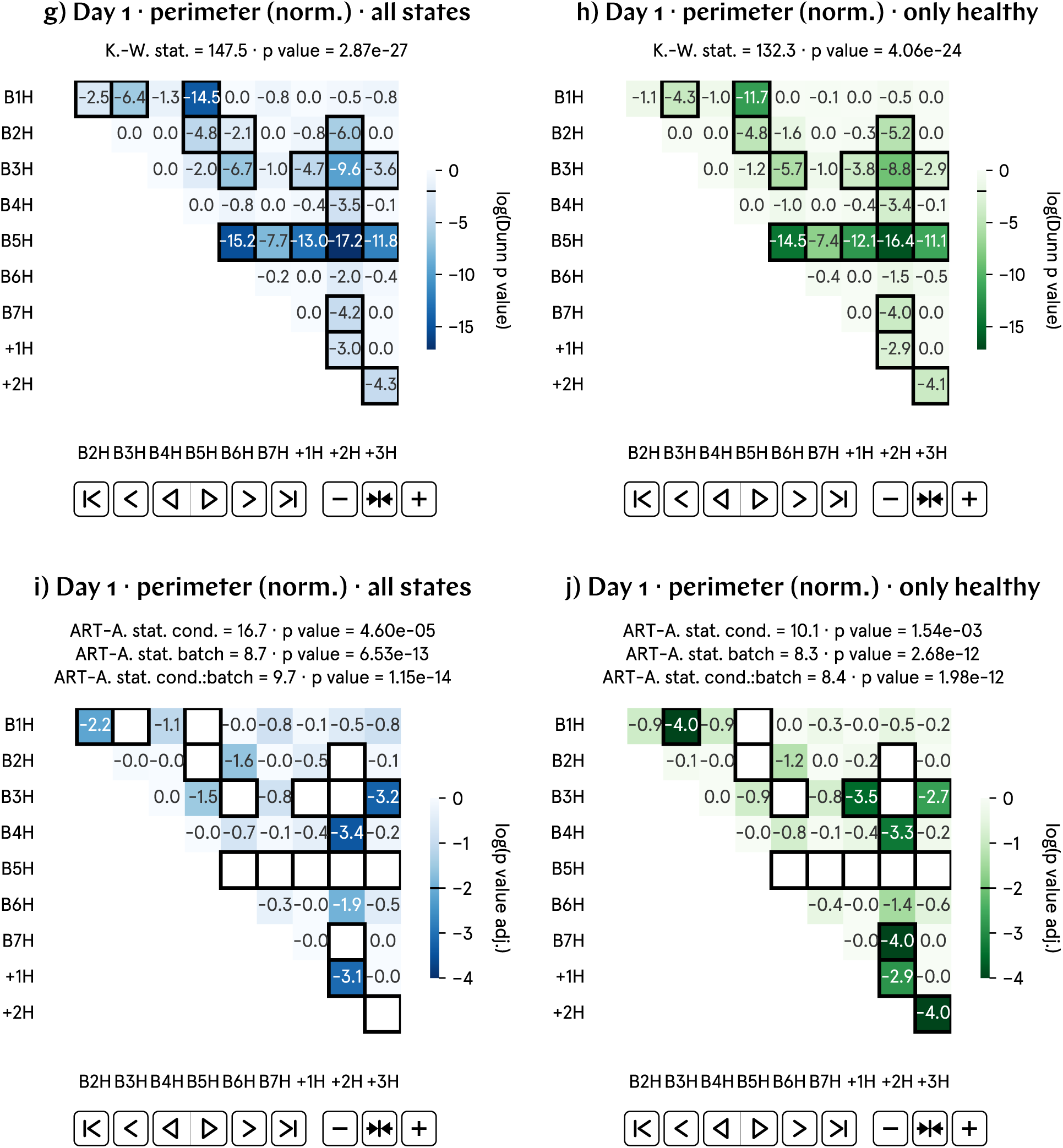
Average day 1 perimeters are only slightly larger in hypomagnetic tadpoles. This contributes to the observed reduced roundness in hypomagnetic tadpoles, as the areas remain mostly unaffected across populations.

### SI65. Day 1 area (norm.) results

**Fig. SI55:**
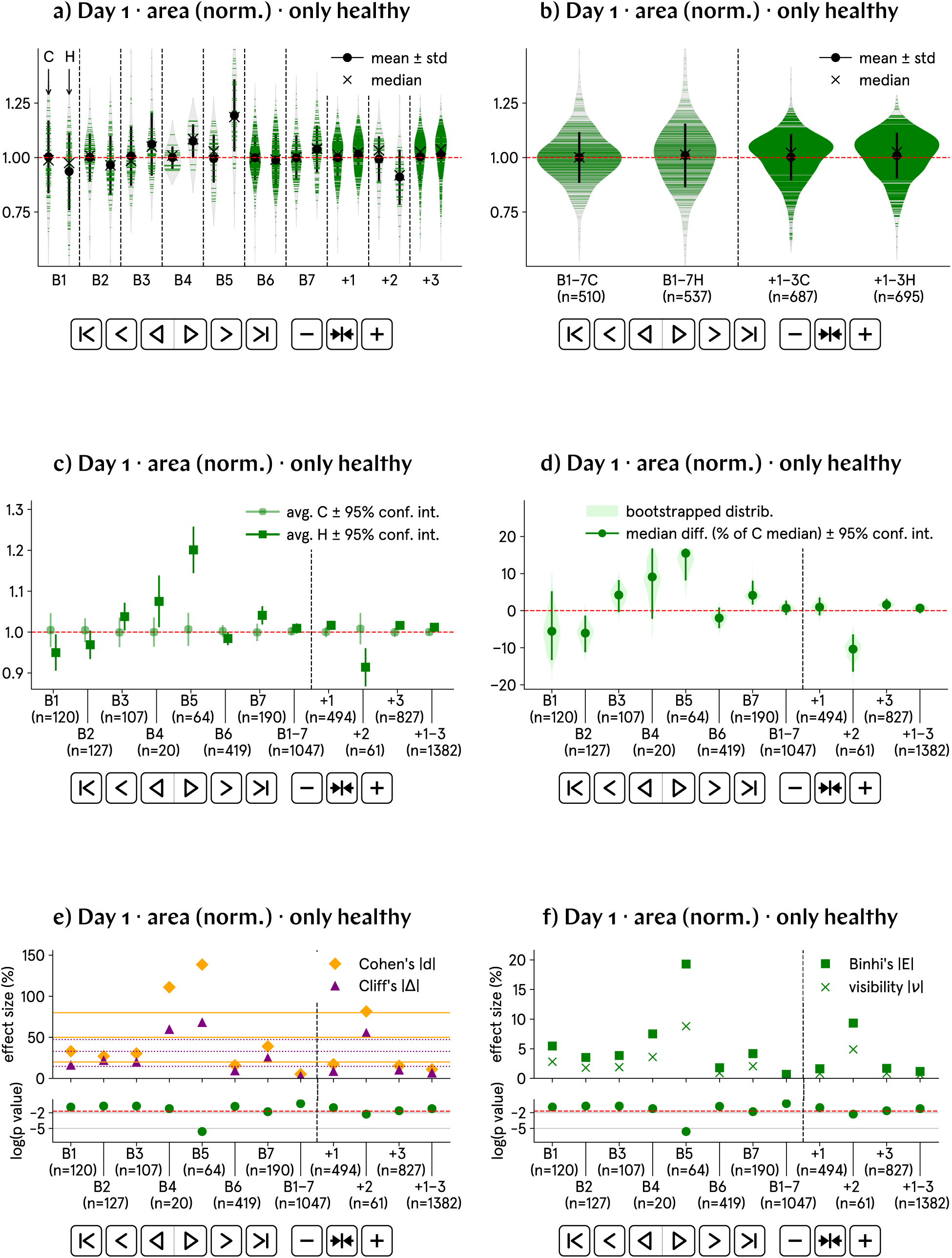
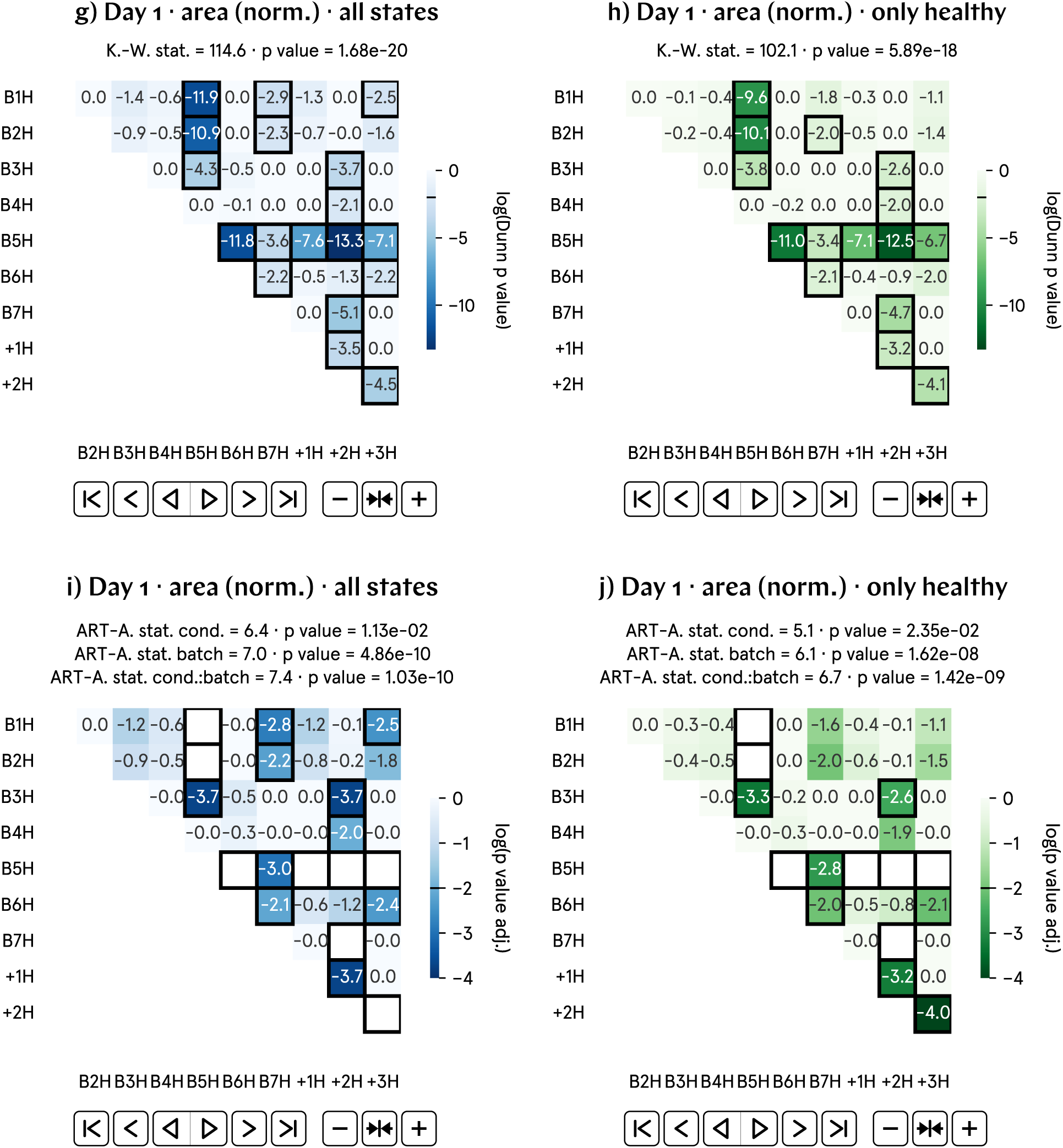
Average day 1 areas are not statistically changed as a function of magnetic field environment. However, the violin plots of b) do show a qualitatively different kernel density distribution for control and hypomagnetic tadpoles; this qualitative difference in distribution is mostly gone for the positive control runs.

### SI66. Day 2 major axis (norm.) results

**Fig. SI56:**
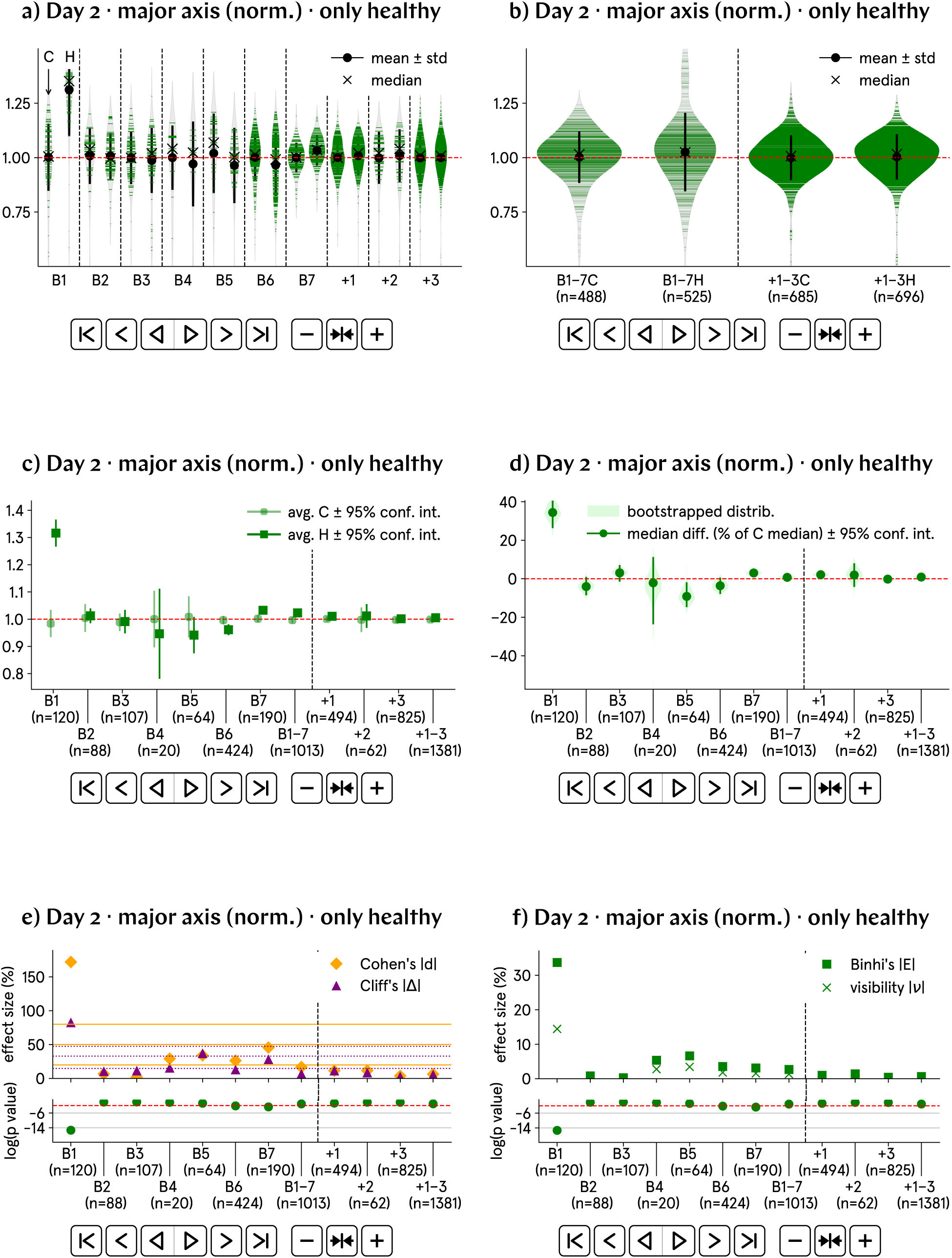
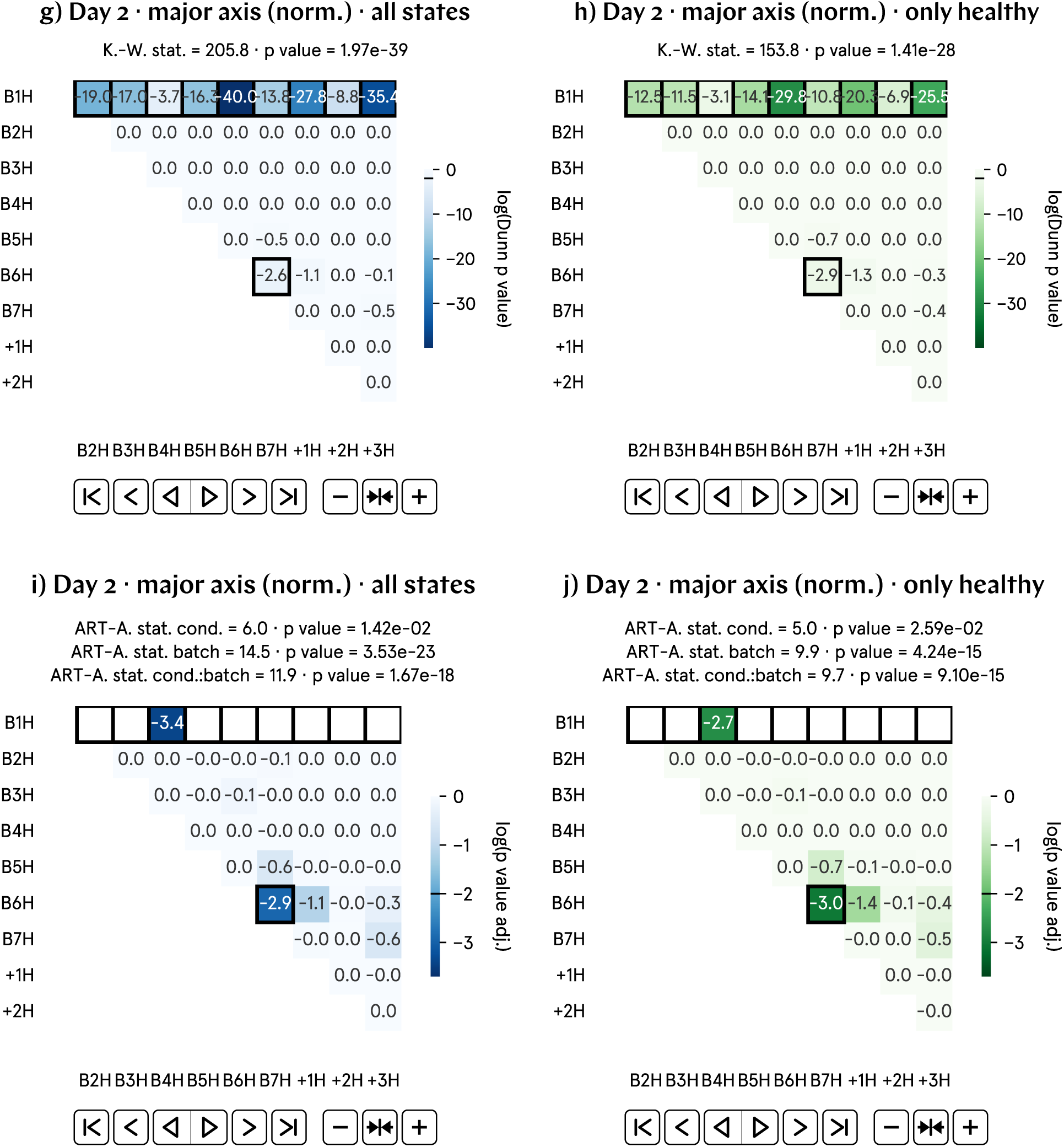
Average day 2 major axes are only slightly longer in hypomagnetic tadpoles. However, the violin plots in b) reveal a qualitatively distinct kernel density distribution between control and hypomagnetic tadpoles, with the hypomagnetic population exhibiting notably longer tails at the higher end of the size spectrum. This qualitative difference in distribution largely disappears in the positive control runs.

### SI67. Day 2 perimeter (norm.) results

**Fig. SI57:**
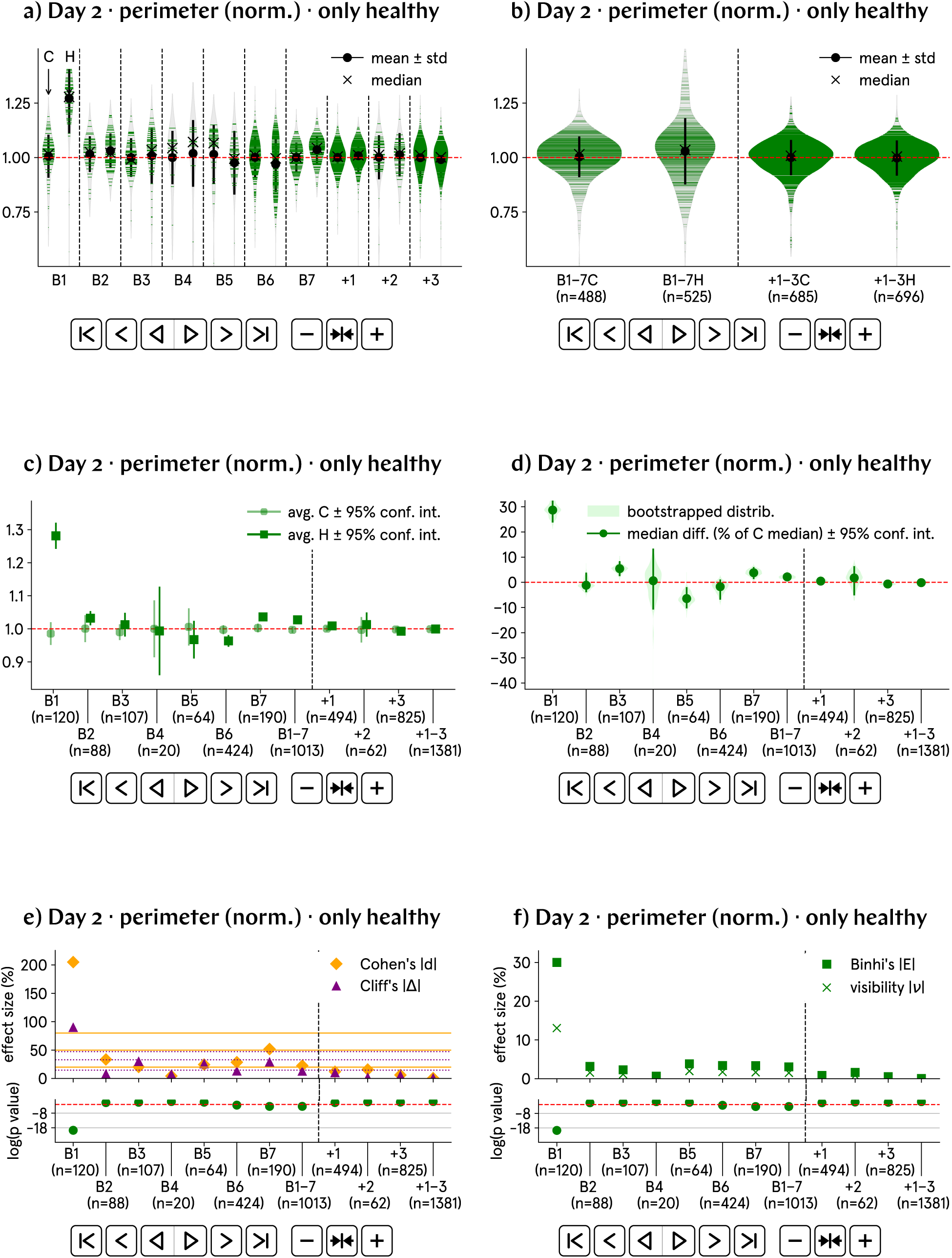
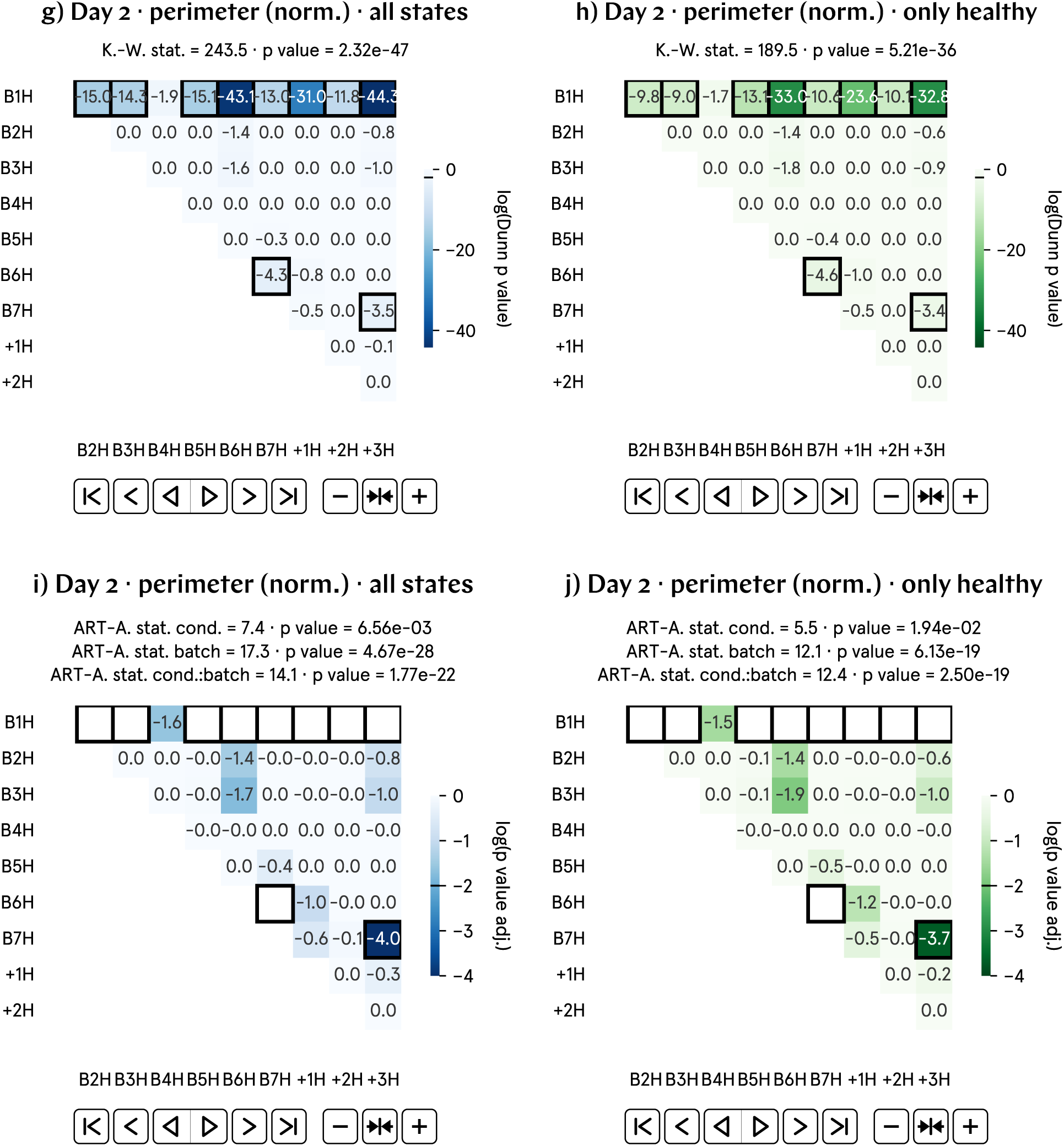
Average day 2 perimeters are only slightly larger in hypomagnetic tadpoles. However, the violin plots of b) do show a qualitatively different kernel density distribution for control and hypomagnetic tadpoles; this qualitative difference in distribution is gone for the positive control runs.

### SI68. Day 2 area (norm.) results

**Fig. SI58:**
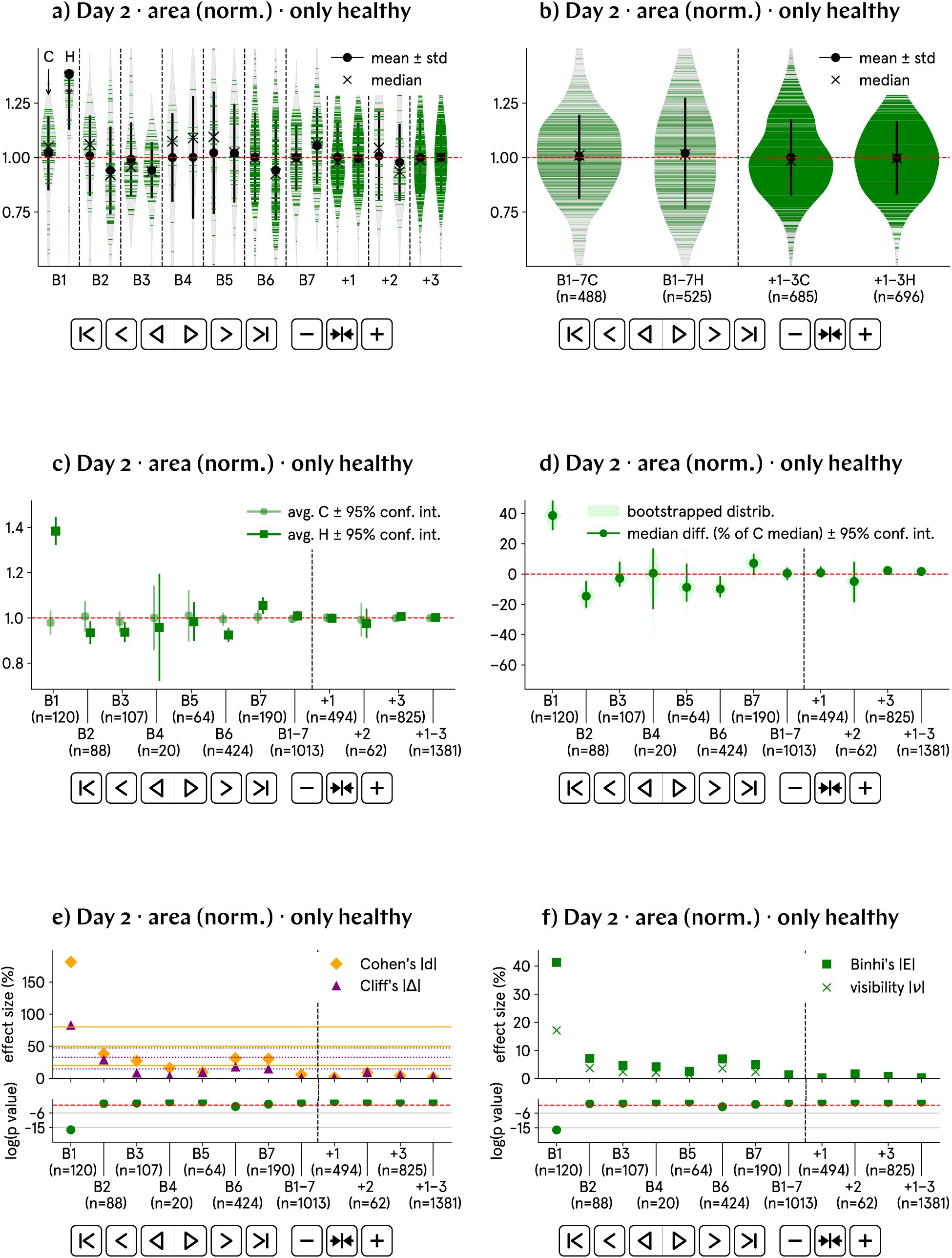
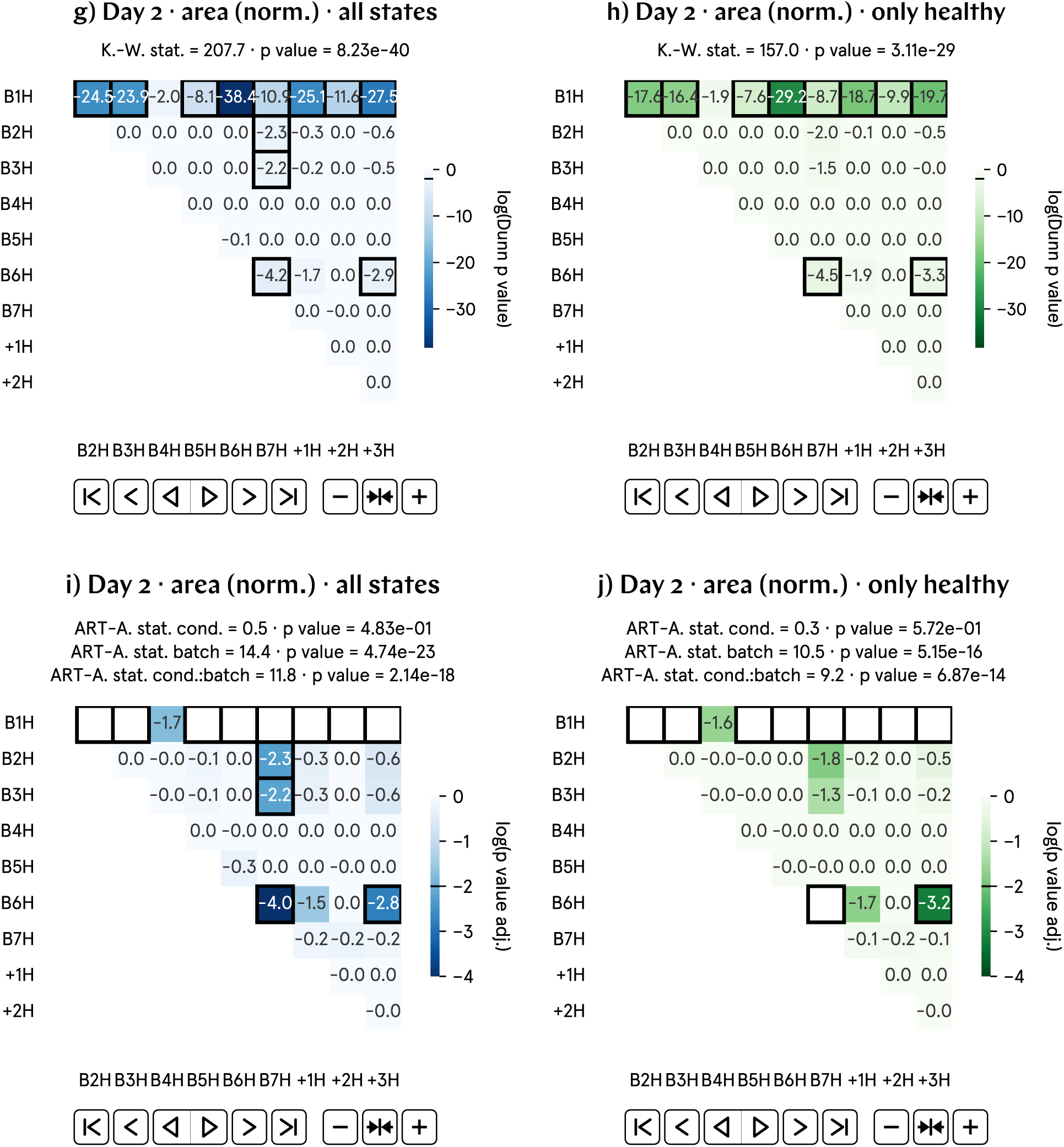
Average day 2 areas are not statistically changed as a function of magnetic field environment. Still, the violin plots of b) do show a qualitatively different kernel density distribution for control and hypomagnetic tadpoles; this qualitative difference in distribution is, however, still present for the positive control runs.

**Fig. SI59:**
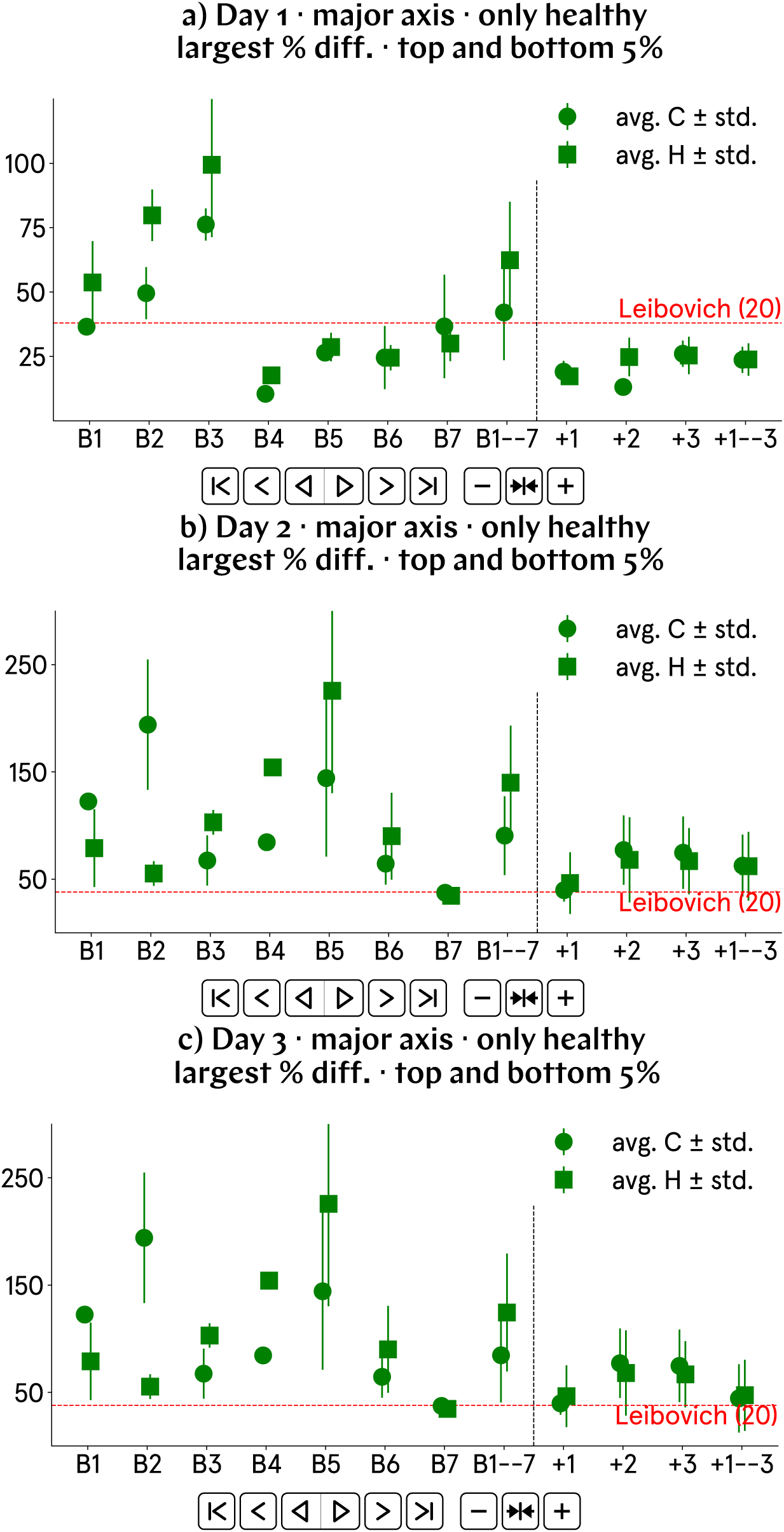
The variability in major axis length is always larger for hypomagnetic tadpoles. a) The variability in major axis length for control embryos in day 1 p.f. images closely aligns with the 38% maximum difference reported in [14] for similar stage embryos. In contrast, the hypomagnetic embryos exhibit approximately 20% greater variability. b) and c) The comparison with the data from [14] is an extrapolation; however, we consistently find a higher variability in the hypomagnetic tadpole population; this variability is considerably reduced for tadpoles during the positive control runs.

### SI69. Guide to GitHub repository

All our work is freely available in the GitHub repository for this work, found at github.com/Quantum-Biology-Institute/Weak-magnetic-field-effects-in-biology-are-measurable.

The repository contains 13 folders; their names and the Supplementary Section where their description can be found are as follows:

**Table SI4:** GitHub folders and the Supplementary Sections containing their descriptions.

| Folder name | Description in Supplementary Section |
| --- | --- |
| 0 Raw data | SI70 |
| 1 Utilities | SI73 |
| 2 Experimental overview | SI73 |
| 3 D1 quantification | SI73 |
| 4 D2 quantification | SI73 |
| 5 D3 quantification | SI73 |
| 6 DeltaD2D3 | SI73 |
| 7 D1 images | SI72 |
| 8 D2 images | SI72 |
| 9 D3 images | SI72 |
| 10 Posters | SI71 |
| 11 Calibration data | SI75 |
| 12 Latex code | N/A (L<sup>A</sup>T<sub>E</sub>X files used to build this document) |

### SI70. Guide to raw data

All raw data in.jpg format is available in the GitHub folder 0 Raw data. This folder contains 10 subfolders: Batches named B1–7 correspond to experimental runs, and batches named B8–10 to positive control runs. The folder names include the fertilization date (in format YYYY_MM_DD). Each batch folder contains three subfolders with raw images for days 1, 2, and 3 p.f., named for the days the images were taken (also in format YYYY_MM_DD); the images are organized in further subfolders by control or sample (meaning hypomagnetic) conditions and plate number.

Batch B1 includes images for up to 7 days p.f. Batch B5 also contains images for tadpoles raised outside our control incubator, on a lab bench (folders labeled ‘outside’). Batches B5 and B6 have additional images for tadpoles raised together in Petri dishes, not in individual wells (folders labeled ‘PD’).

### SI71. Guide to plate posters

For each of the 111 well plates used in this work, along with the Petri dishes also used in batches B5 and B6, we created posters illustrating the 3-day tadpole progression, available in the GitHub folder 10 Posters. Plate posters display either 24 or 48 unique tadpoles, depending on whether a 24- or 48-well plate was used in the experiment.

Each plate poster is named by: batch number (with batches named B1–7 corresponding to experimental runs, and batches named B8–10 to positive control runs); experimental condition, control (C) or hypomagnetic (H); and plate number. For example, poster B3CP1.pdf is for batch 3, control condition, first plate. For Batch B1, we also provide posters including images for up to 7 days p.f. (named ‘B1XPY_7days.pdf’, with X = *{*C, H*}* the condition and Y the plate number).

Within each poster, tadpole images are numbered by well number. The accompanying text for each tadpole is color-coded based on the human assessment of tadpole health (see details in Supplementary Section SI7): green (healthy), orange (one health condition), red (more than one health condition), and black (dead by day 1 p.f.). If a health condition is observed, it is noted in the accompanying text.

Missing images in the posters are replaced with the cute icon shown below. If the reason for the missing image is known, it is noted in the accompanying text.

### SI72. Guide to segmented images

Post-segmentation.png images that are used in the automated analysis are found in GitHub folders [7 D1 images](https://github.com/Quantum-Biology-Institute/Weak-magnetic-field-effects-in-biology-are-measurable/tree/main/7%20D1%20images), [8 D2 images](https://github.com/Quantum-Biology-Institute/Weak-magnetic-field-effects-in-biology-are-measurable/tree/main/8%20D2%20images) and [9 D3 images](https://github.com/Quantum-Biology-Institute/Weak-magnetic-field-effects-in-biology-are-measurable/tree/main/9%20D3%20images), for days 1, 2, and 3 p.f., respectively.

These images are separated by batches, conditions and plates. Batches B1 through B7 correspond to our experimental runs; batches B8 through B10 correspond to the positive control runs. Inside each batch folder, images are subdivided and uniquely labeled into control or hypomagnetic condition (with a C or an H, respectively); the day of the image (day 1, 2, or 3 p.f.); the plate number; and finally the well number within the plate. For example, the image B1CD1P3-08.png represents the tadpole image for batch B1, in the control condition, at day 1 p.f., located in plate 3 and well 8. Wells are numbered sequentially by row: in a 48-well plate, A1 to A8 correspond to wells that we number 1 to 8, B1 to B8 to wells 9 to 16, and so forth. The same numbering system applies to 24-well plates.

### SI73. Guide to python code

All the used python code is available in the GitHub repository for this work, found at github.com/Quantum-Biology-Institute/Weak-magnetic-field-effects-in-biology-are-measurable, in multiple folders.

We recommend that interested researchers download all GitHub folders to a local directory and then modify the base path in the python files (located at the first line of each script) to match the directory where the files were downloaded to.

Folder [1 Utilities](https://github.com/Quantum-Biology-Institute/Weak-magnetic-field-effects-in-biology-are-measurable/tree/main/1%20Utilities) contains the code for the automated image analysis; most auxiliary functions are found in the file utilities.py. Folder [2 Experimental overview](https://github.com/Quantum-Biology-Institute/Weak-magnetic-field-effects-in-biology-are-measurable/tree/main/2%20Experimental%20overview) contains the file Posters.py that is used to build the poster with 3-day images for each well-plate.

The major analysis workflow is as follows:

1. Images from folders [7 D1 images](https://github.com/Quantum-Biology-Institute/Weak-magnetic-field-effects-in-biology-are-measurable/tree/main/7%20D1%20images), [8 D2 images](https://github.com/Quantum-Biology-Institute/Weak-magnetic-field-effects-in-biology-are-measurable/tree/main/8%20D2%20images) and [9 D3 images](https://github.com/Quantum-Biology-Institute/Weak-magnetic-field-effects-in-biology-are-measurable/tree/main/9%20D3%20images) are manually copied into folders ‘(X+2) DX quantification/BY/BYC’ or ‘…/BYH’, with X in {1, 2, 3} and Y in {1, 2, …, 10}. For example, day 3 images for the first positive control run (*i.e.*, B8) are copied from their respective plate subfolders within 9 D3 images/B8 into folders 5 D3 quantification/B8/B8C and 5 D3 quantification/B8/B8H.
2. In their new folders, the images are resized to a common size for each batch using ResizeD1.py for day 1 or ResizeD2D3.py for days 2 and 3 p.f.;
3. The resized images are automatically analyzed using AnalysisAndPlot.py. This function generates two output.txt files per batch, one per condition, with the relevant parameters (*e.g.*, major axis, eccentricity, area, perimeter, …) for all images. These files are saved in folder ‘(X+2) DX quantification/Results/BY/BYCDX_analysis.txt’ and ‘…/BYHDX_analysis.txt’. For example, for day 2 quantification of batch B7, results for the hypomagnetic condition analysis are saved in folder 4 D2 quantification/Results/B7/B7HD2_analysis.txt;
4. The statistical analysis plots of a desired property, by day and for all batches, are generated by the file StatsAndPlots.py. For the progression of the quantities between days 2 and 3, file [StatsAndPlotsDeltaD2D3.py](https://github.com/Quantum-Biology-Institute/Weak-magnetic-field-effects-in-biology-are-measurable/blob/main/1%20Utilities/StatusAndPlotDeltaD2D3.py) is used instead. The analysis of each desired property is programatically set in the file SetupDays.py. If a plot-saving flag is set to True in one of the analysis functions, results are saved to folder ‘(X+2) DX quantification/Results/DX_{property name}’. For example, for day 1 quantification of eccentricity, plots are saved to the folder 3 D1 quantification/Results/D1_eccentricity. Outputs.txt files with results of the statistical tests are always saved to the above directory.

### SI74. Guide to statistical analysis plots

Our automated statistical analysis for each property, whose underlying code is described in Supplementary Section SI73, produces 10 pairs of plots, labeled a) through j). Here, we provide guidance on how to understand each plot.

Each plot title includes the following information: the day post-fertilization (1, 2, or 3) corresponding to the analyzed images; the property under study (along with units, if applicable); and a qualifier indicating the health status of the tadpoles (see details in Supplementary Section SI7). The qualifier is either ‘all states’ for tadpoles in all assessed health conditions, or ‘only healthy’ for tadpoles without any observed health conditions.

Our 7 experimental runs are called batches and labeled B1 through B7. We have three positive control runs labeled +1 through +3 in these plots. Values obtained for the hypomagnetic (respectively: control) condition tadpoles are labeled H (C). The label H in the positive control runs (*e.g.*, +2H) refers to tadpoles exposed to a field mimicking Earth’s generated inside the hypomagnetic chamber.

Details by plot are provided below:

a. presents the measured values of the property under study, grouped by batch and condition (control or hypomagnetic). Each tadpole’s value is represented by a horizontal line, color-coded as follows: green for healthy tadpoles, orange for tadpoles with one observed health condition, red for tadpoles with two or more health conditions, and black for tadpoles that were dead by day 1. The violin outlines display the kernel density estimation of the data distribution for each batch, with the width of the violin indicating the frequency of data points at corresponding values. For each batch, the control tadpoles are plotted in the leftmost violin, and the tadpoles under hypomagnetic conditions are plotted in rightmost violin, as indicated by the legend (C for control, H for hypomagnetic) in the top left, above the first batch (B1). A vertical dashed black line separates the violin pairs between different batches. Within each violin: The circular marker indicates the mean value; vertical lines extending from the mean show the standard deviation; and the median is denoted by an ‘x’ marker. In normalized plots, the mean value of the control violin is set to 1 for each batch. A red dashed horizontal line at 1 on the y-axis (ordinate) indicates this normalization. The labeled y-axis values and the y-axis range are manually chosen. The plot can toggle between showing violins for the entire tadpole population or only for the healthy subset, where only green horizontal lines (representing healthy tadpoles) are displayed.
b. presents two sets of violins comparing the aggregate values for control and hypomagnetic conditions across the 7 experimental runs, labeled B1–7C for control and B1–7H for hypomagnetic; each tadpole counts once. Additionally, two violins compare the aggregate values for control and positive control within the hypomagnetic chamber, labeled +1–3C for control and +1–3H for positive control. Each pair of violins is separated by a vertical dashed black line, and the total number of data points (one per tadpole) is displayed below each violin label. Within each violin: The circular marker indicates the mean value; vertical lines extending from the mean show the standard deviation; and the median is denoted by an ‘x’ marker. In normalized plots, the mean value of the control violin is set to 1 for each batch. A red dashed horizontal line at 1 on the y-axis (ordinate) indicates this normalization. The labeled y-axis values and the y-axis range are manually chosen, and correspond to those in plot a). The plot can toggle between showing violins for the entire tadpole population or only for the healthy subset, where only green horizontal lines (representing healthy tadpoles) are displayed.
c. illustrates the average value of the property under study for both the control and hypomagnetic populations, along with a 95% confidence interval (calculated as described in Supplementary Section SI16). The average for the control population is represented by a circular marker, while that of the hypomagnetic population is represented by a square marker. The experimental data and positive control data are separated by a vertical dashed black line. The data is presented by batch, and after batch B7, the aggregated average value, with an equal weight for each tadpole, and its corresponding confidence interval are plotted above the label B1–7. Similarly, to the right of the datasets for the three positive control runs, we display the aggregated average value and confidence interval above the label +1–3. In normalized plots, the mean value for the control population across all health states is set to 1. A red dashed horizontal line at 1 on the y-axis indicates this normalization. The labeled y-axis values and the y-axis range are automatically chosen by our code. The plot can toggle between showing the average for the entire tadpole population and for the healthy subset only. Blue markers are used for the entire population, and green markers are used for the healthy subset.
d. shows the bootstrapped distribution (as explained in Supplementary Section SI16) of the median difference between control and hypomagnetic populations, expressed as a percentage of the control median. The violins for the resampled populations are plotted by batch, with a circular marker indicating the bootstrapped median and a vertical line, its 95% confidence interval (calculated as described in Supplementary Section SI16). A vertical dashed black line separates the experimental data from the positive control data. After batch B7, the aggregated median value, weighted equally for each tadpole, along with its corresponding confidence interval, is plotted above the label B1–7. Similarly, to the right of the datasets for the three positive control runs, the aggregated median value and confidence interval are plotted above the label +1–3. The total number of data points (one per tadpole) is displayed below each batch label. In normalized plots, the median of the control population is set to 0, and a red dashed horizontal line at 0 on the y-axis indicates this normalization. The labeled y-axis values are automatically chosen by the code, with the y-axis range set between the minimum value and 80% of the maximum of all bootstrapped median differences. The plot can toggle between displaying the results for the entire tadpole population or only the healthy subset. Blue markers denote the entire population, while green markers represent the healthy subset.
e. is a dual plot. The top y-axis displays the absolute values of Cohen’s *d* (in orange diamond markers) and Cliff’s Δ (in purple triangle markers) by batch, representing two measures of effect size (explained in Supplementary Section SI14). Three solid orange lines at 20%, 50%, and 80% denote the conventional thresholds for small, medium, and large effect sizes, according to Cohen’s *d*. Similarly, three dotted purple lines at 15%, 33%, and 47% represent the thresholds for small, medium, and large effects according to Cliff’s Δ. On the bottom y-axis, the base-10 logarithm of the p value for each batch is displayed by circular markers on a log scale. Each p value is calculated as follows: A separate Kolmogorov-Smirnov normality test [17] is performed on both the control and hypomagnetic populations for each batch. If the p value from the normality test is greater than 0.01 for both populations (*i.e.*, both populations are normally distributed), the p value from a T-test [17] between the populations is plotted. If at least one population is not normally distributed (which is usually the case), the p value from a non-parametric Mann-Whitney test [17] is plotted instead. In the bottom plot, a dashed red line at y = −2 corresponds to p = 0.01; points above this threshold are not statistically significant at the 0.01 level. A vertical dashed black line separates the experimental data from the positive control data. After batch B7, the aggregated median value, equally weighted for each tadpole, along with its corresponding confidence interval, is plotted above the label B1–7. Similarly, the aggregated median value and confidence interval for the three positive control runs are plotted above the label +1–3. The total number of data points (one per tadpole) is displayed beneath each batch label. Both the y-axis labels and ranges are automatically determined by the code for both plots. The plot can toggle between displaying data for the entire tadpole population and only the healthy subset. While the top plot remains unchanged in color for different conditions, the bottom plot uses blue markers for the entire population and green markers for the healthy subset.
f. is a dual plot. The top y-axis displays the absolute values of Binhi’s *E* (in square markers) and the visibility *ν* (in ‘x’ markers) by batch, representing two measures of effect size (explained in Supplementary Section SI14). On the bottom y-axis, the base-10 logarithm of the p value for each batch is displayed by circular markers on a log scale. Each p value is calculated as follows: A separate Kolmogorov-Smirnov normality test [17] is performed on both the control and hypomagnetic populations for each batch. If the p value from the normality test is greater than 0.01 for both populations (*i.e.*, both populations are normally distributed), the p value from a T-test [17] between the populations is plotted. If at least one population is not normally distributed (which is usually the case), the p value from a non-parametric Mann-Whitney test [17] is plotted instead. In the bottom plot, a dashed red line at y = −2 corresponds to p = 0.01; points above this threshold are not statistically significant at the 0.01 level. A vertical dashed black line separates the experimental data from the positive control data. After batch B7, the aggregated median value, equally weighted for each tadpole, along with its corresponding confidence interval, is plotted above the label B1–7. Similarly, the aggregated median value and confidence interval for the three positive control runs are plotted above the label +1–3. The total number of data points (one per tadpole) is displayed beneath each batch label. Both the y-axis labels and ranges are automatically determined by the code for both plots. The plot can toggle between displaying data for the entire tadpole population and only the healthy subset. Both the top and bottom plots use blue markers for the entire population and green markers for the healthy subset. The bottom log plot in plot f) is identical to that in plot e). We chose to separate the plotting of Cohen’s *d* and Cliff’s Δ from Binhi’s *E* and the visibility *ν* because, in our data, we observed that each pair of effect size measures is more similar in magnitude within its pair, whereas the first pair (Cohen’s *d* and Cliff’s Δ) tends to be larger in magnitude.
g. presents the graphical representation of the Dunn post-hoc test results following a Kruskal-Wallis test which identified significant differences *within* the control or hypomagnetic condition groups (as explained in Supplementary Section SI16) [17], for all tadpole health states (represented by the blue color palette). The Kruskal-Wallis statistic (‘K.-W. stat.’) and the corresponding p value (which must be less than 0.01 to justify the Dunn test) are displayed beneath the plot title. The numbers within the individual matrix entries represent the base-10 logarithm of the p value between two different batches (within the same condition). The darker the blue in a matrix entry, the greater the difference between the two batches, *for the same condition*. For batch pairs with a Dunn p value less than 0.01 (*i.e.*, matrix values less than -2), the corresponding matrix entries are framed with a black border, indicating significant differences at the 0.01 level. This threshold is also visually marked by a black line at log(Dunn p value) = -2 on the rightmost color scale. p values smaller than the computational threshold for zero are represented by a white square with a black border. The plot can toggle between displaying data for the control and hypomagnetic populations (with labels containing a C or an H, respectively). The first seven rows of the matrix correspond to the experimental runs (labeled with a B), while the last three rows represent the positive control runs (labeled with a +).
h. presents the graphical representation of the Dunn post-hoc test results following a Kruskal-Wallis test which identified significant differences *within* the control or hypomagnetic condition groups (as explained in Supplementary Section SI16) [17], for the healthy tadpole subset only (represented by the green color palette). The Kruskal-Wallis statistic (‘K.-W. stat.’) and the corresponding p value (which must be less than 0.01 to justify the Dunn test) are displayed beneath the plot title. The numbers within the individual matrix entries represent the base-10 logarithm of the p value between two different batches (within the same condition). The darker the green in a matrix entry, the greater the difference between the two batches, *for the same condition*. For batch pairs with a Dunn p value less than 0.01 (*i.e.*, matrix values less than -2), the corresponding matrix entries are framed with a black border, indicating significant differences at the 0.01 level. This threshold is also visually marked by a black line at log(Dunn p value) = -2 on the rightmost color scale. p values smaller than the computational threshold for zero are represented by a white square with a black border. The plot can toggle between displaying data for the control and hypomagnetic populations (with labels containing a C or an H, respectively). The first seven rows of the matrix correspond to the experimental runs (labeled with a B), while the last three rows represent the positive control runs (labeled with a +).
i. presents the graphical representation of the Tukey post-hoc test results following an ART-ANOVA non-parametric test [17] which identified significant differences *within* the control or hypomagnetic condition groups (as explained in Supplementary Section SI16), for all tadpole health states (represented by the blue color palette). The ART-ANOVA statistics (‘ART-A. stat.’) and the corresponding p values (which must all be less than 0.01 to justify the Tukey test) are displayed beneath the plot title, for: the condition (control or hypomagnetic, ‘cond.’); the batch (‘batch’); and the interaction between the condition and the batch (‘cond.:batch’). The numbers within the individual matrix entries represent the base-10 logarithm of the adjusted p value between two different batches (within the same condition). The darker the blue in a matrix entry, the greater the difference between the two batches, *for the same condition*. For batch pairs with an adjusted p value less than 0.01 (*i.e.*, matrix values less than -2), the corresponding matrix entries are framed with a black border, indicating significant differences at the 0.01 level. This threshold is also visually marked by a black line at log(p value adj.) = -2 on the rightmost color scale. p values smaller than the computational threshold for zero are represented by a white square with a black border. The plot can toggle between displaying data for the control and hypomagnetic populations (with labels containing a C or an H, respectively). The first seven rows of the matrix correspond to the experimental runs (labeled with a B), while the last three rows represent the positive control runs (labeled with a +).
j. presents the graphical representation of the Tukey post-hoc test results following an ART-ANOVA non-parametric test [17] which identified significant differences *within* the control or hypomagnetic condition groups (as explained in Supplementary Section SI16), for the healthy tadpole subset only (represented by the green color palette). The ART-ANOVA statistics (‘ART-A. stat.’) and the corresponding p values (which must all be less than 0.01 to justify the Tukey test) are displayed beneath the plot title, for: the condition (control or hypomagnetic, ‘cond.’); the batch (‘batch’); and the interaction between the condition and the batch (‘cond.:batch’). The numbers within the individual matrix entries represent the base-10 logarithm of the adjusted p value between two different batches (within the same condition). The darker the green in a matrix entry, the greater the difference between the two batches, *for the same condition*. For batch pairs with an adjusted p value less than 0.01 (*i.e.*, matrix values less than -2), the corresponding matrix entries are framed with a black border, indicating significant differences at the 0.01 level. This threshold is also visually marked by a black line at log(p value adj.) = -2 on the rightmost color scale. p values smaller than the computational threshold for zero are represented by a white square with a black border. The plot can toggle between displaying data for the control and hypomagnetic populations (with labels containing a C or an H, respectively). The first seven rows of the matrix correspond to the experimental runs (labeled with a B), while the last three rows represent the positive control runs (labeled with a +).

### SI75. Guide to calibration data

In this folder, we present the raw calibration data and the code used to plot calibration curves.

Subfolder [Degaussing door test](https://github.com/Quantum-Biology-Institute/Weak-magnetic-field-effects-in-biology-are-measurable/tree/main/11%20Calibration%20data/Degaussing%20door%20test) contains data for the calibration of how magnetic fields change inside the hypomagnetic chamber as a function of its door opening and closing, and of the degaussing procedure, as plotted in Supplementary Fig. SI4.

Subfolder [Field uniformity](https://github.com/Quantum-Biology-Institute/Weak-magnetic-field-effects-in-biology-are-measurable/tree/main/11%20Calibration%20data/Field%20uniformity) contains the 3D interactive HTML plot of mapped residual fields inside the hypomagnetic chamber, as plotted in Supplementary Fig. SI5. The HTML file needs to be downloaded and opened with an internet browser.

Subfolder [Environment calibration](https://github.com/Quantum-Biology-Institute/Weak-magnetic-field-effects-in-biology-are-measurable/tree/main/11%20Calibration%20data/Environment%20calibration) contains data for the calibration of temperature, humidity, pressure, and light levels (with a sensor of sensitivity 0.01 Lux) in both the hypomagnetic chamber and the control box, as plotted in Supplementary Fig. SI7.

Subfolder [Spectra](https://github.com/Quantum-Biology-Institute/Weak-magnetic-field-effects-in-biology-are-measurable/tree/main/11%20Calibration%20data/Spectra) contains data for the calibration of spectral properties and light intensity levels (with a sensor of sensitivity 188 *µ*lux) in both the hypomagnetic chamber and the control box, as plotted in Supplementary Fig. SI8.

### SI76. Statistical comparisons across control and hypomagnetic populations

Given the quantitatively established high variability between individual batches, we are hesitant to include statistical comparisons *across* control and hypomagnetic conditions. As a point of curiosity, we cautiously present the *full* Tukey post-hoc matrix of adjusted p values following a non-parametric ART-ANOVA test for many of the properties we investigate. These plots extend the statistical plots of types i) and j), incorporating the adjusted p values for comparing control and hypomagnetic results.

It is important to emphasize that many of the studied biological properties, when not normalized by batch, exhibit statistically significant differences across batches, even within the same condition.

Note that we cannot extend plots of types g) and h): Dunn’s post-hoc test is specifically designed for pairwise comparisons between groups after a Kruskal-Wallis test and does not account for interaction terms. This is because the Kruskal-Wallis and Dunn tests focus on comparing group medians without modeling interactions between factors. Interaction effects, such as those between control and hypomagnetic conditions across different batches, require tests that support interaction terms, such as the non-parametric ART-ANOVA.

**Fig. SI60:**
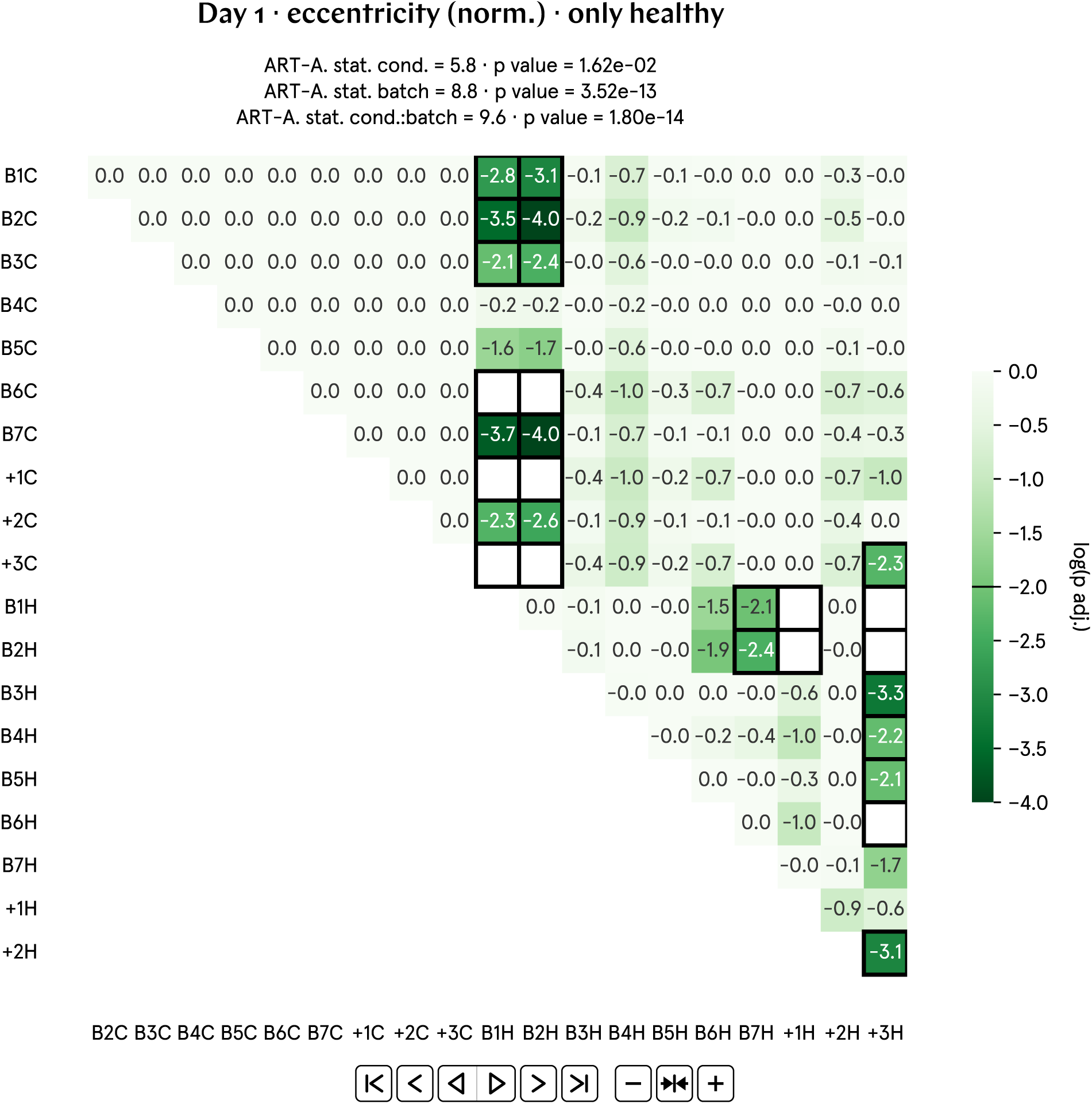
Full post-hoc matrix of adjusted p values for day 1 eccentricity (norm.).

**Fig. SI61:**
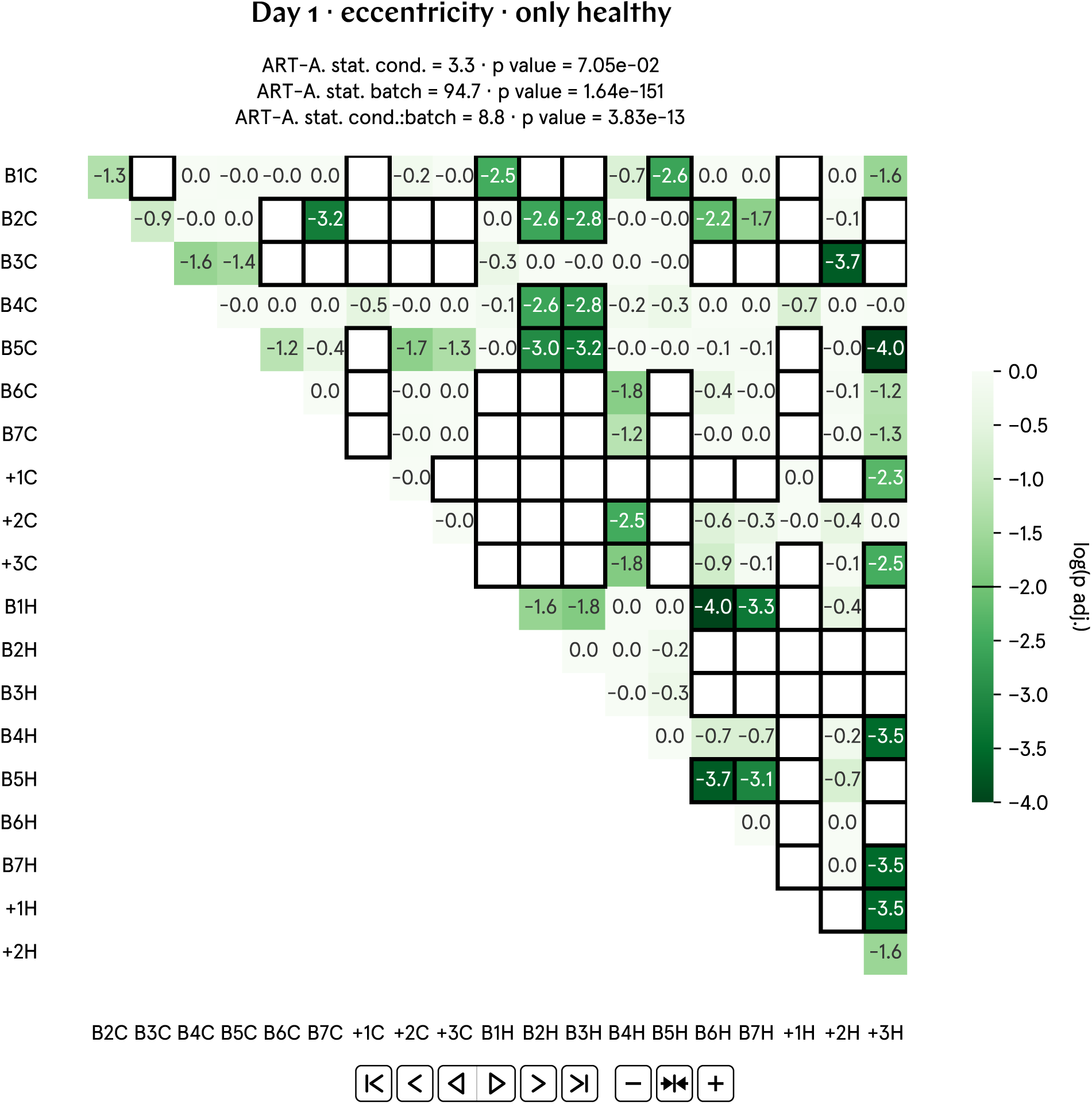
Full post-hoc matrix of adjusted p values for day 1 eccentricity.

**Fig. SI62:**
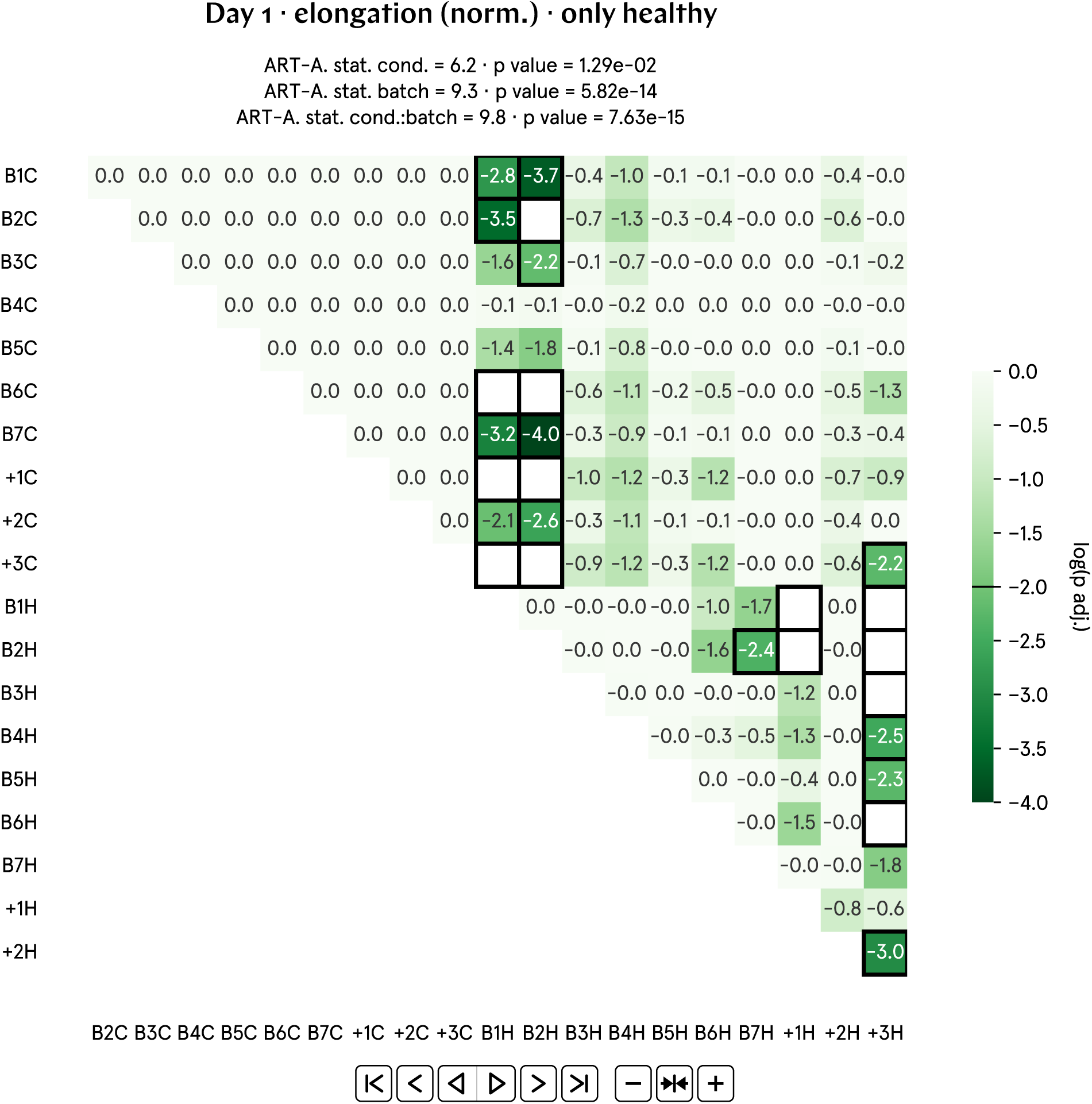
Full post-hoc matrix of adjusted p values for day 1 elongation (norm.).

**Fig. SI63:**
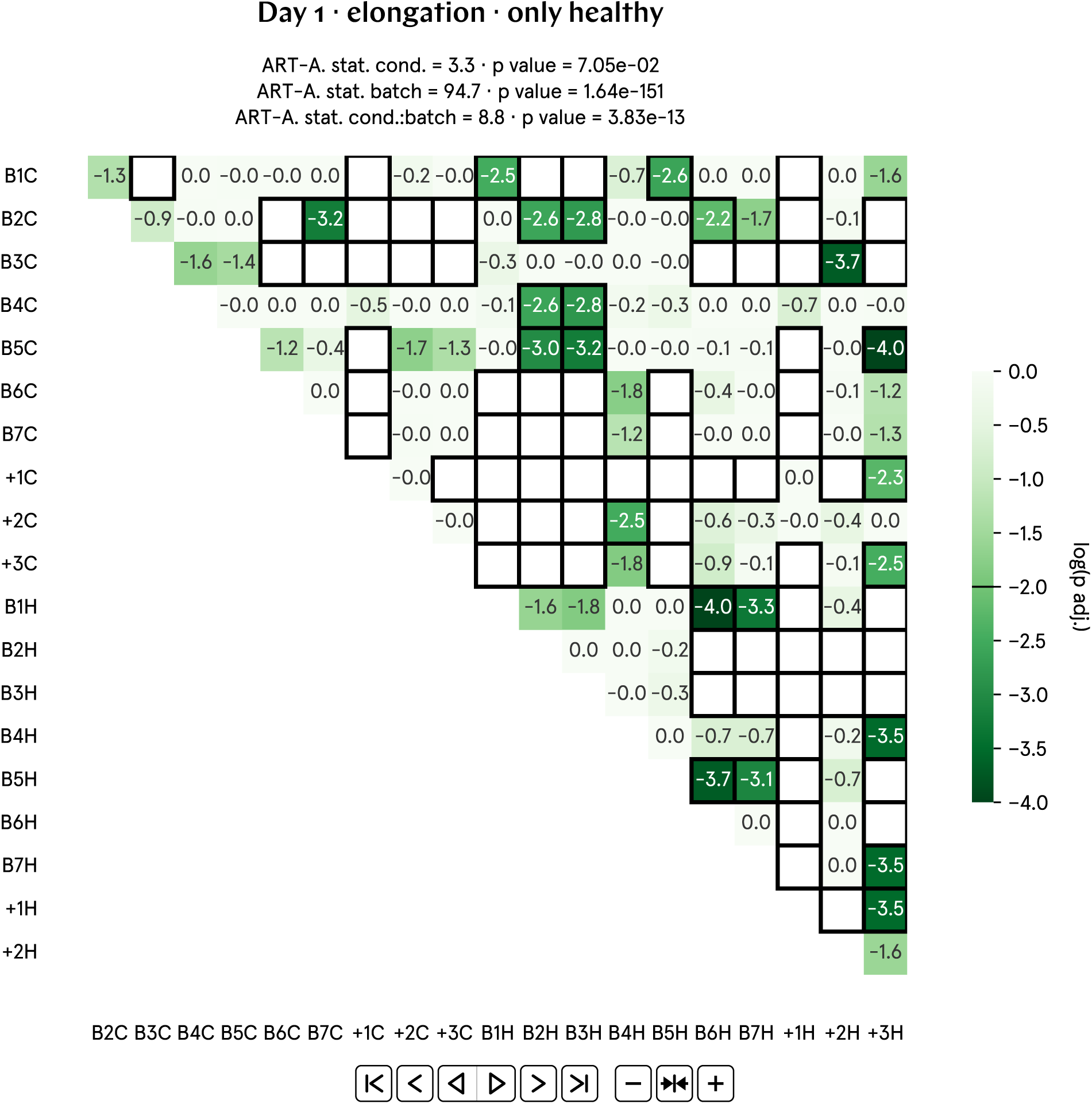
Full post-hoc matrix of adjusted p values for day 1 elongation.

**Fig. SI64:**
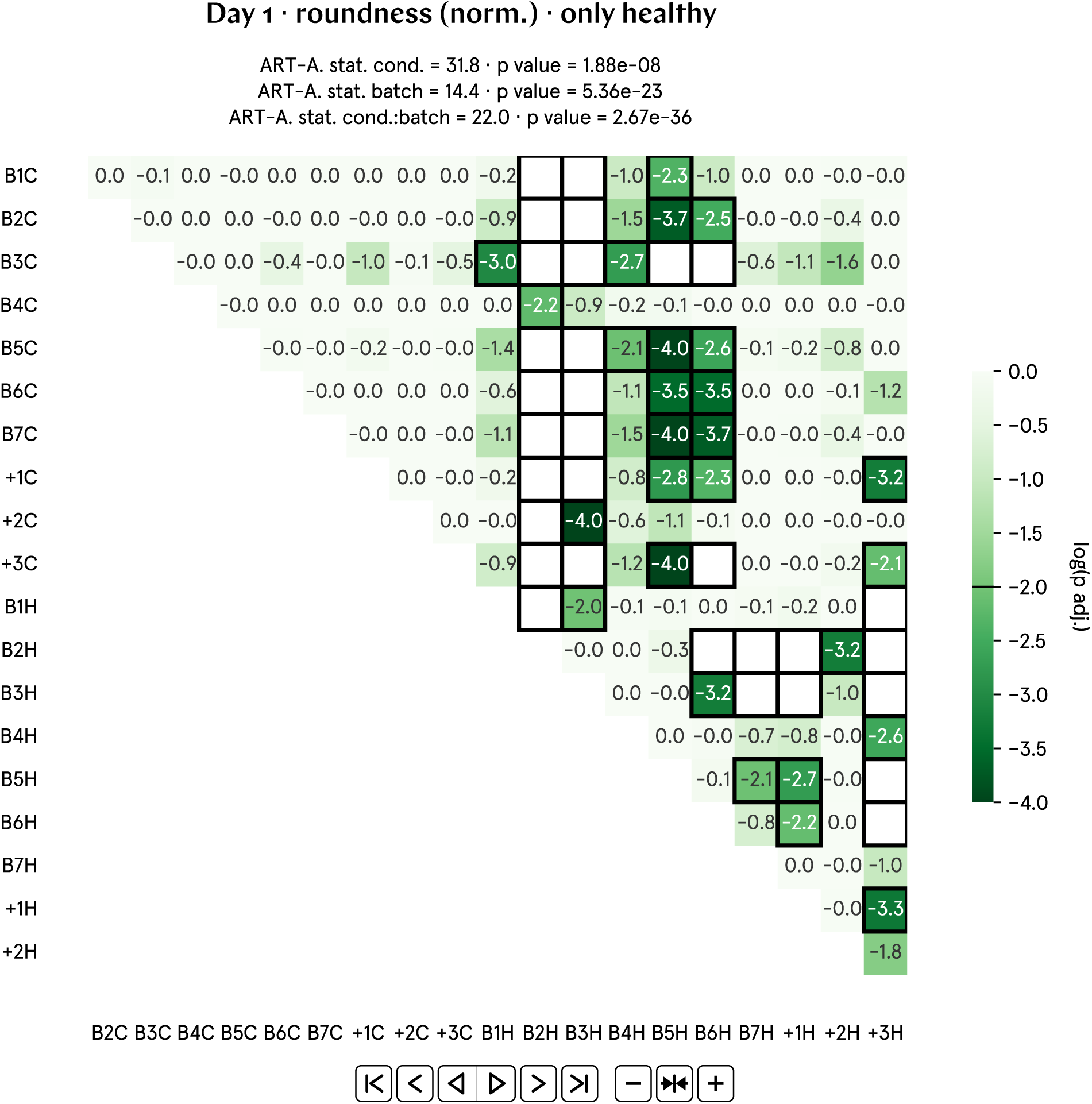
Full post-hoc matrix of adjusted p values for day 1 roundness (norm.).

**Fig. SI65:**
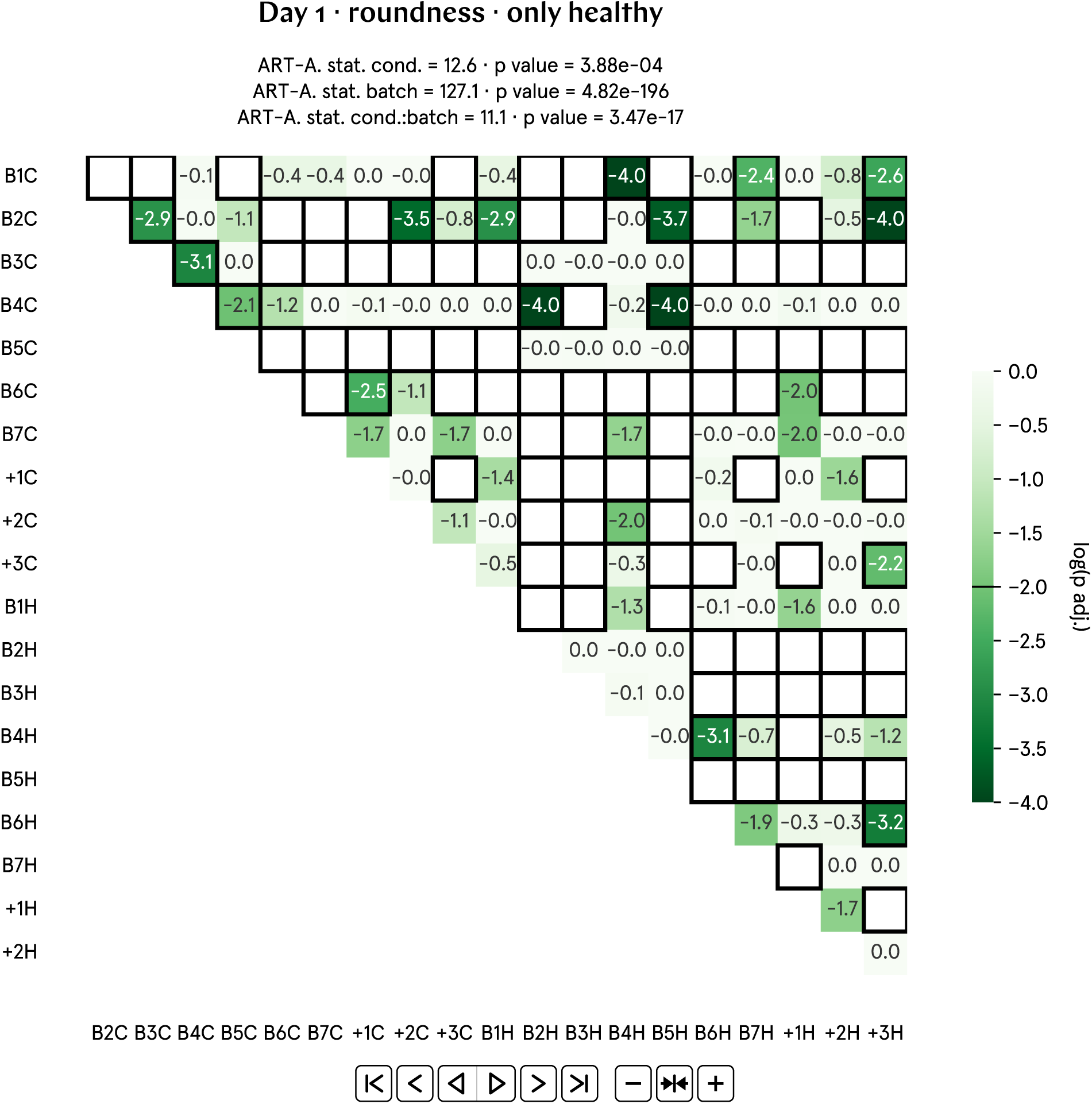
Full post-hoc matrix of adjusted p values for day 1 roundness.

**Fig. SI66:**
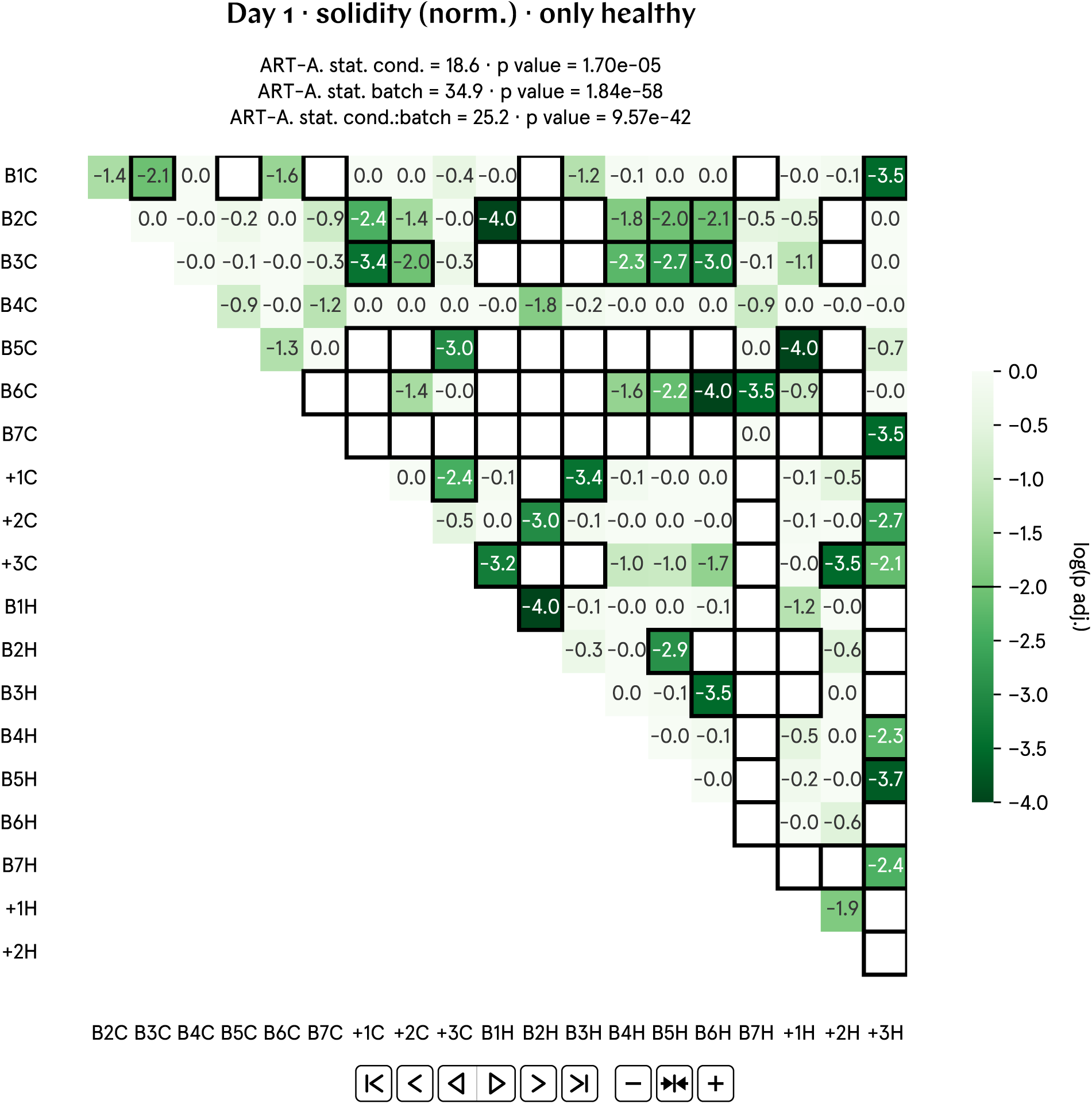
Full post-hoc matrix of adjusted p values for day 1 solidity (norm.).

**Fig. SI67:**
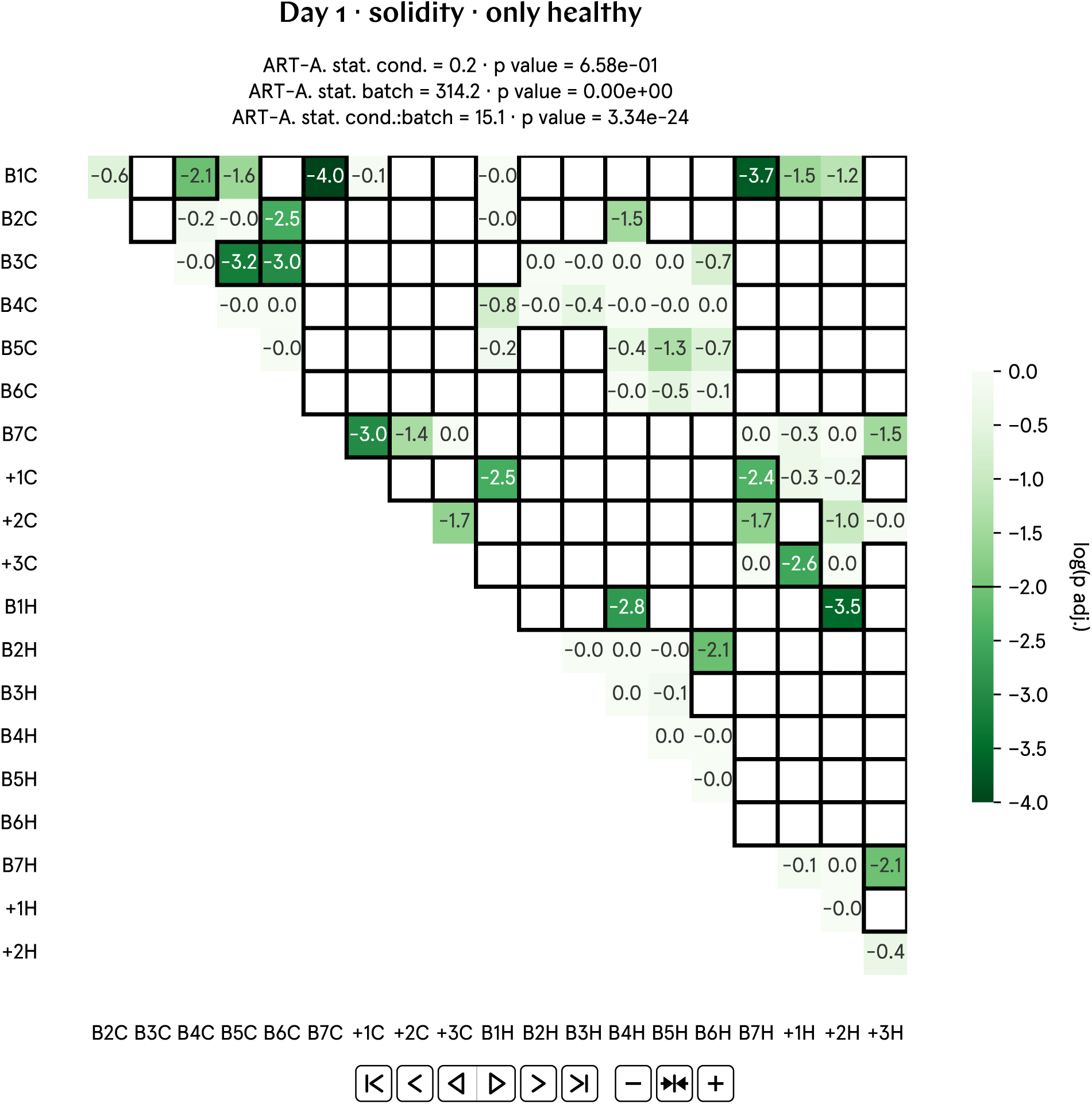
Full post-hoc matrix of adjusted p values for day 1 solidity.

**Fig. SI68:**
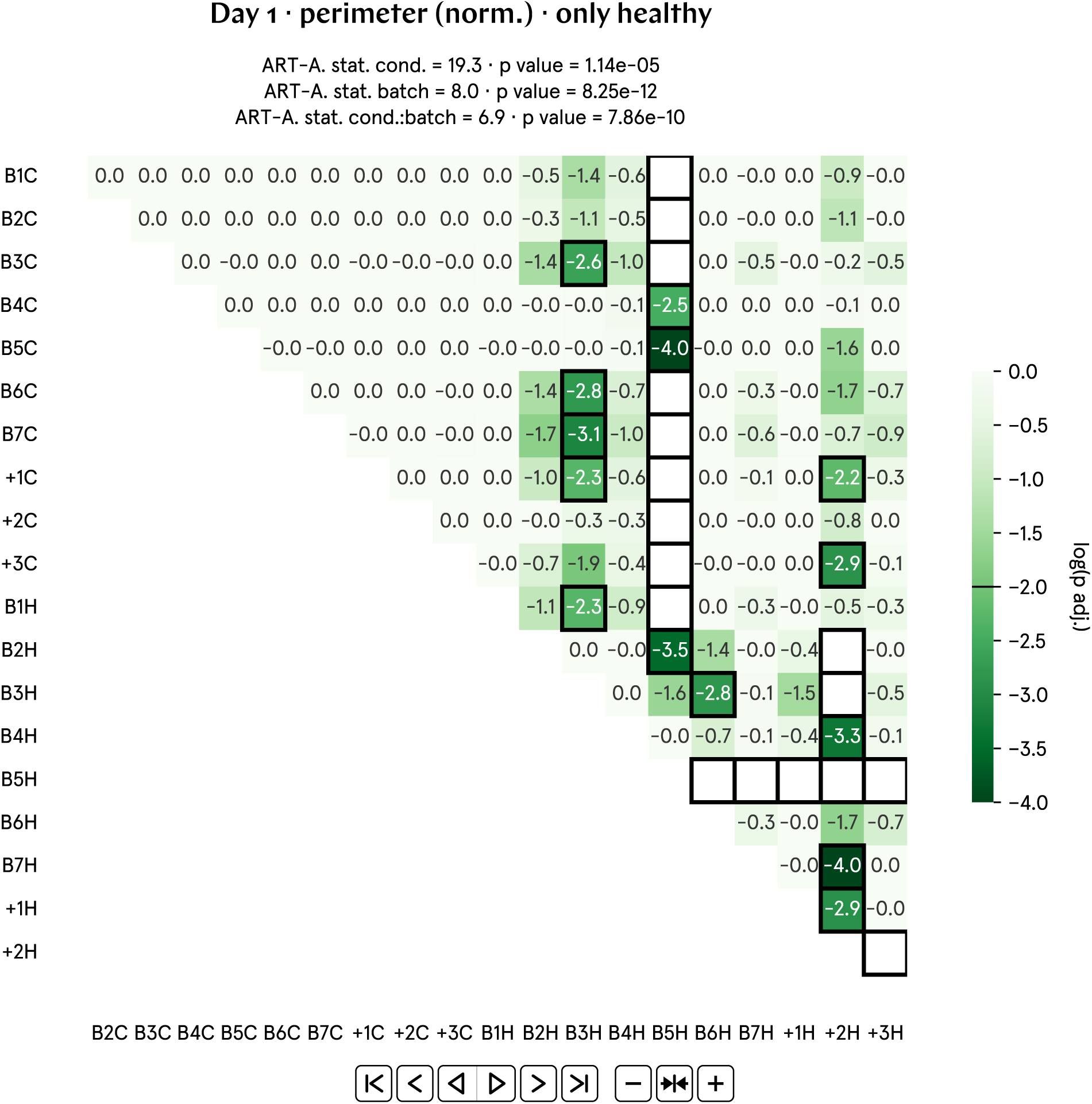
Full post-hoc matrix of adjusted p values for day 1 perimeter (norm.).

**Fig. SI69:**
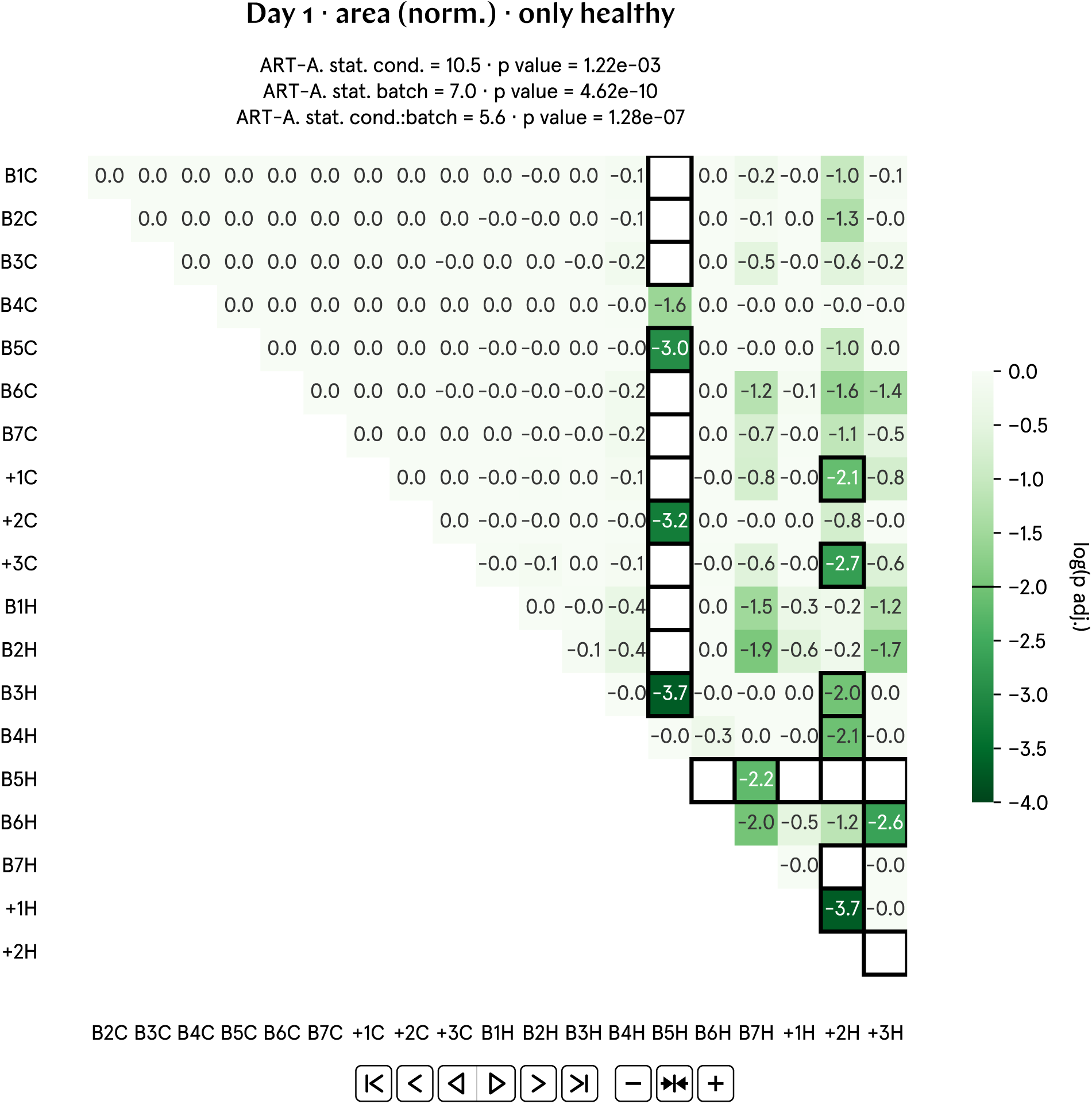
Full post-hoc matrix of adjusted p values for day 1 area (norm.).

**Fig. SI70:**
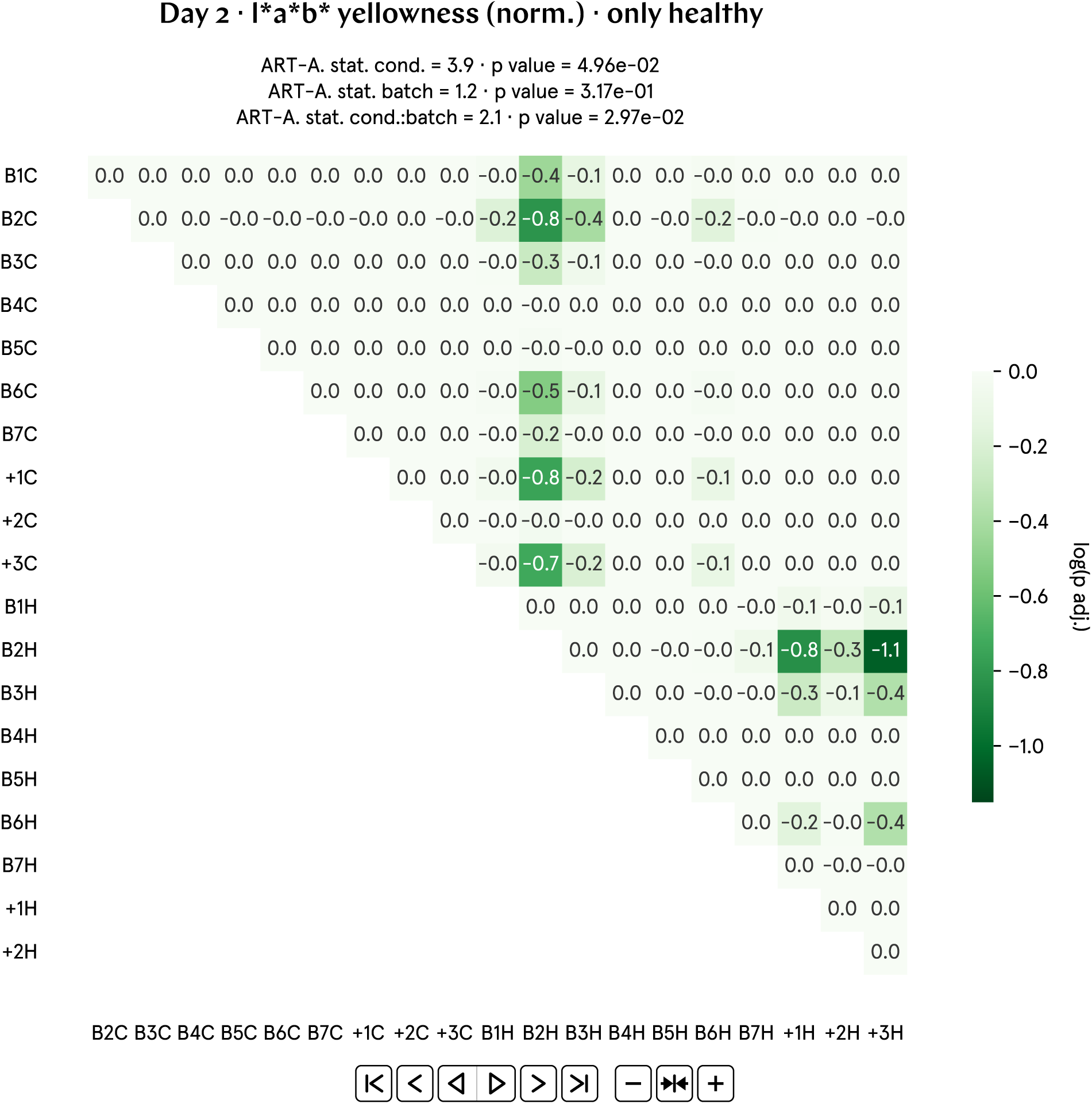
Full post-hoc matrix of adjusted p values for day 2 L*a*b* yellowness (norm.).

**Fig. SI71:**
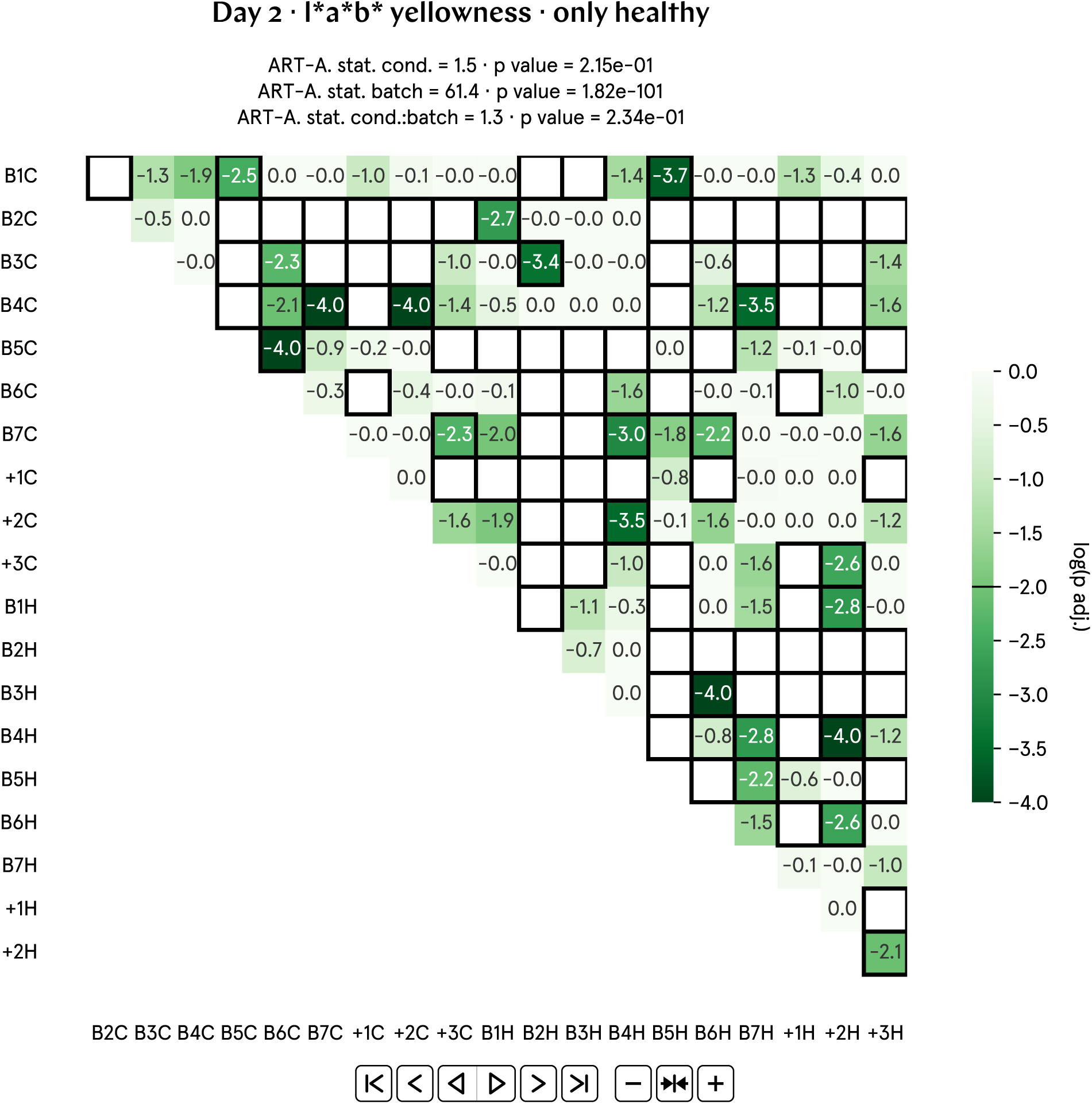
Full post-hoc matrix of adjusted p values for day 2 L*a*b* yellowness.

**Fig. SI72:**
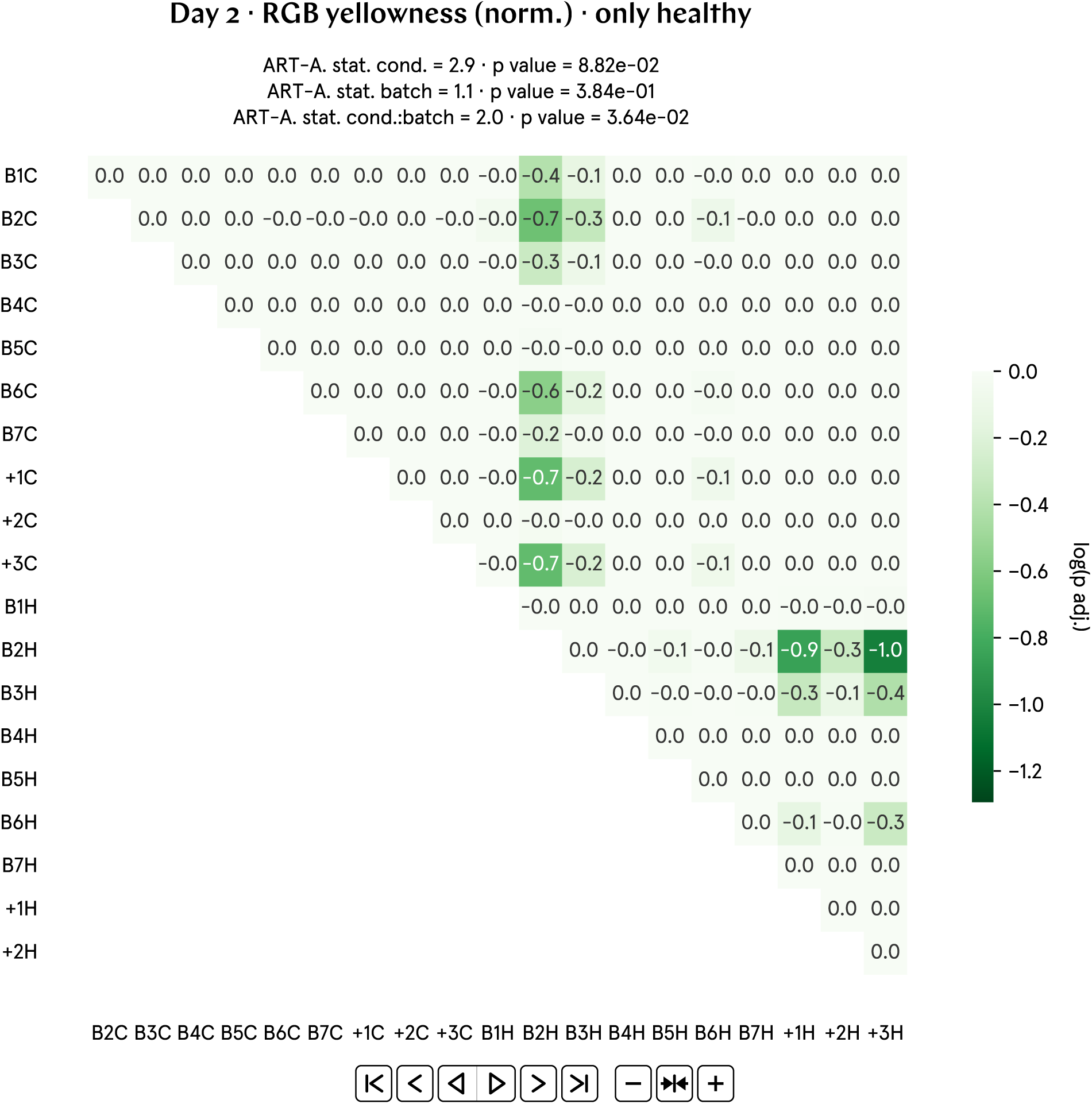
Full post-hoc matrix of adjusted p values for day 2 RGB yellowness (norm.).

**Fig. SI73:**
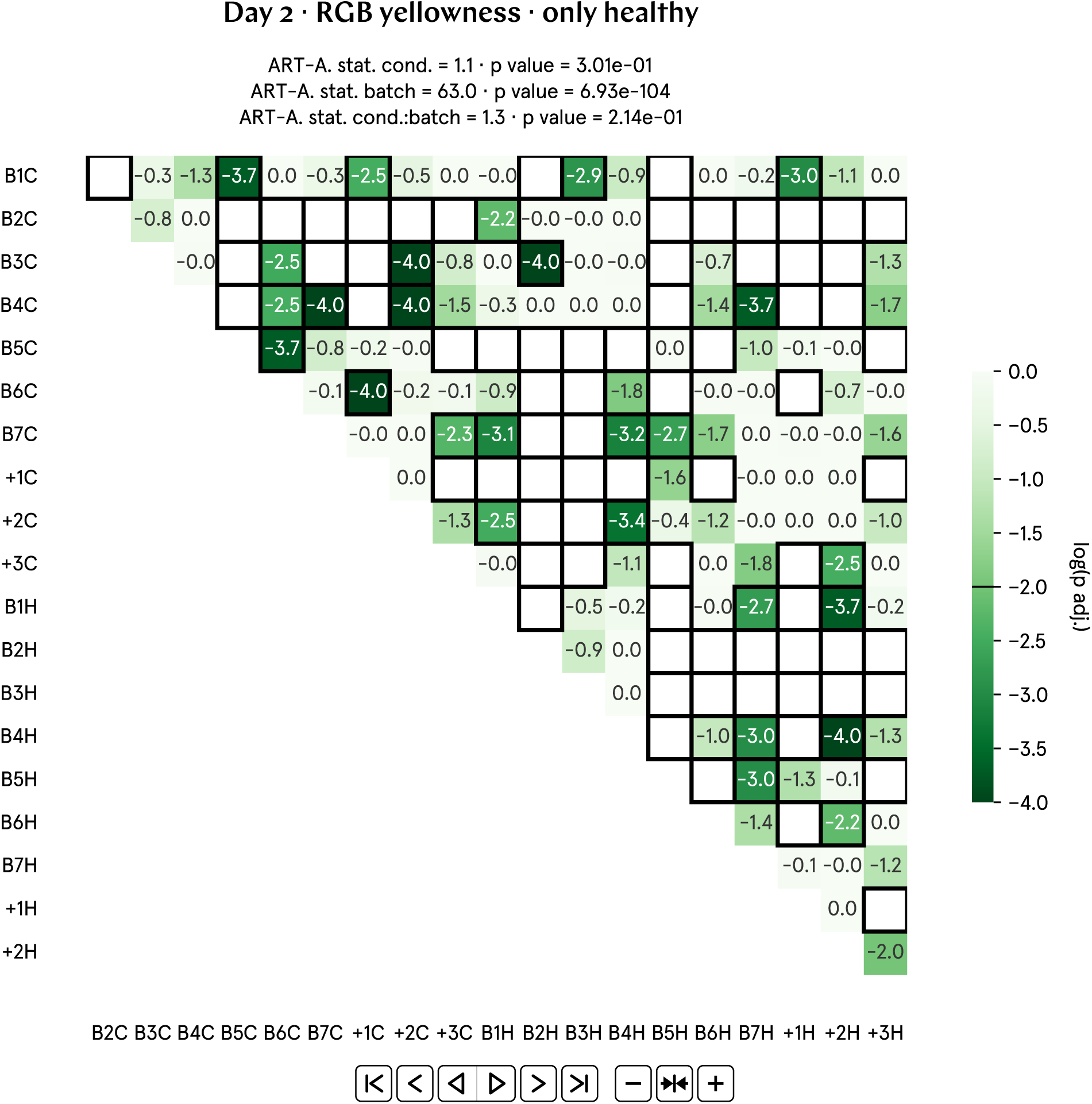
Full post-hoc matrix of adjusted p values for day 2 RGB yellowness.

**Fig. SI74:**
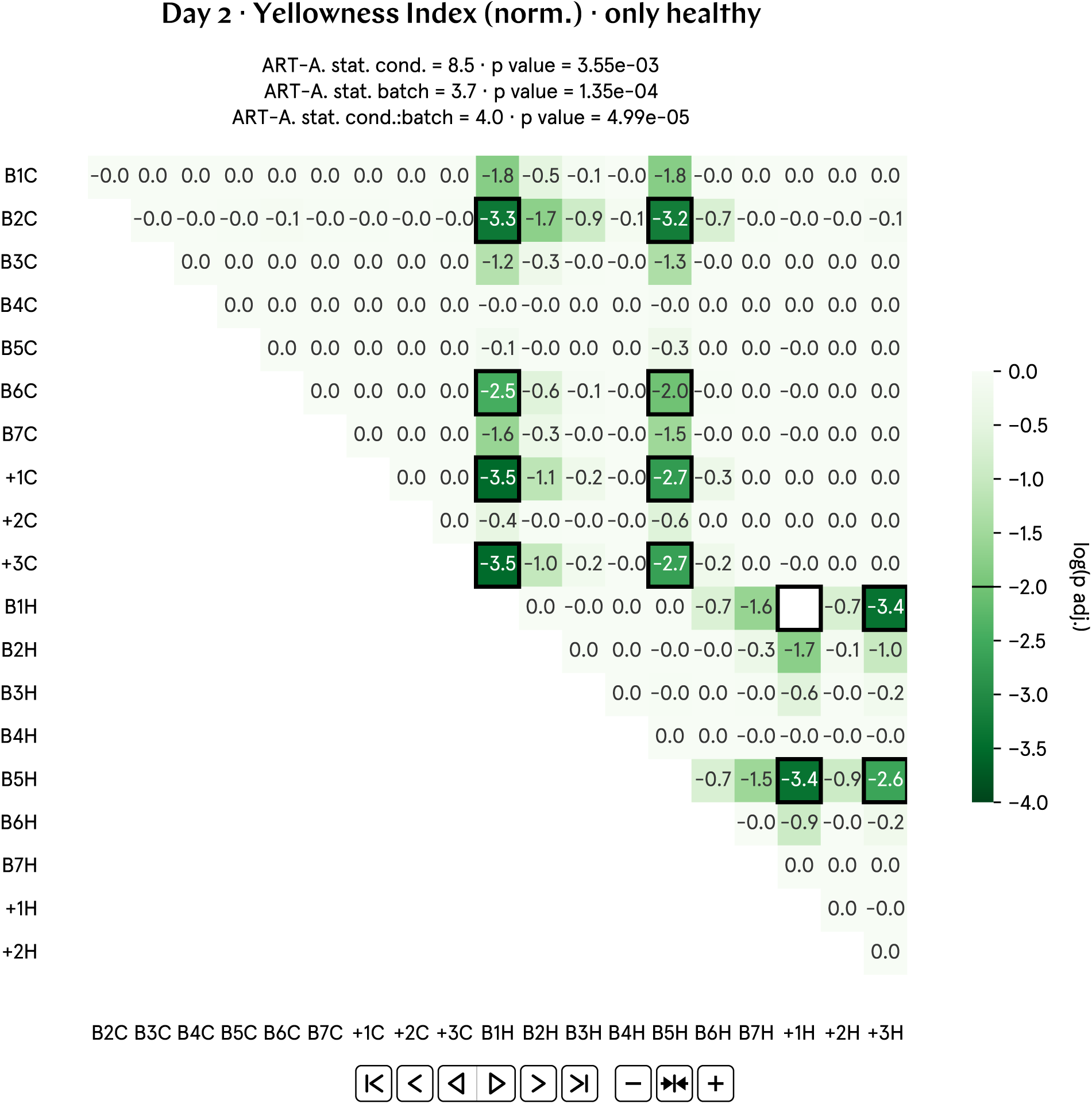
Full post-hoc matrix of adjusted p values for day 2 Yellowness Index (norm.).

**Fig. SI75:**
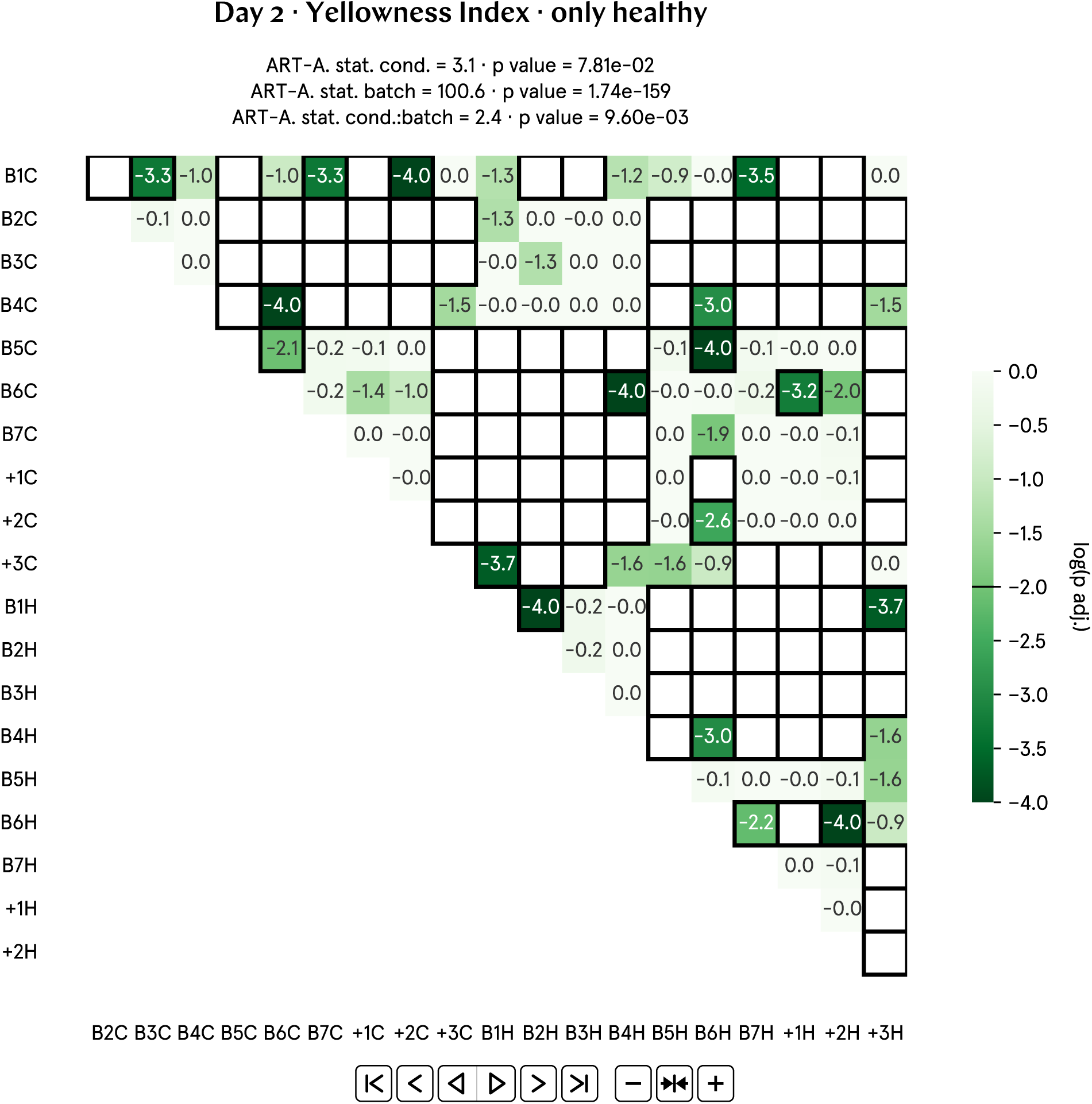
Full post-hoc matrix of adjusted p values for day 2 Yellowness Index.

**Fig. SI76:**
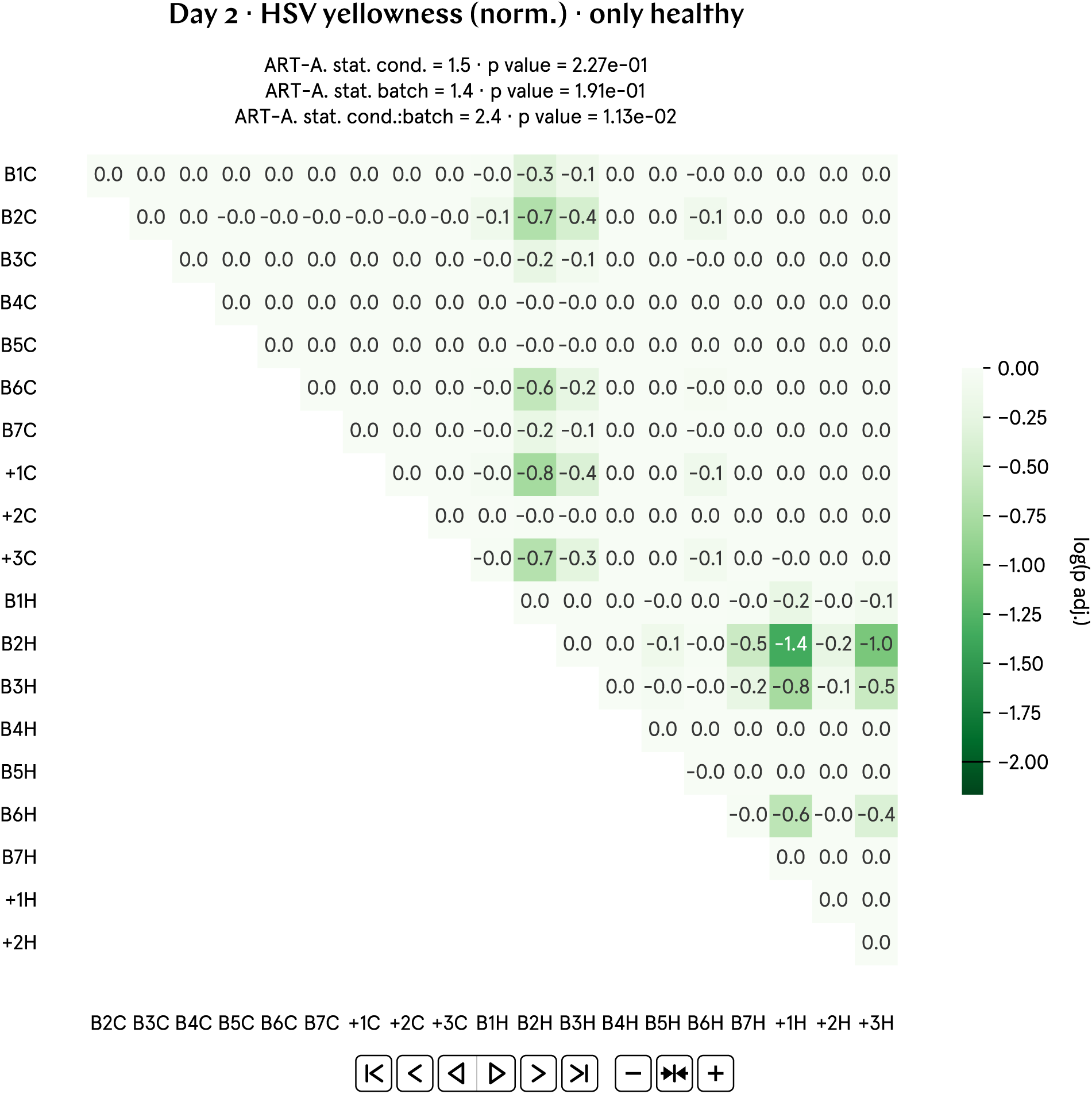
Full post-hoc matrix of adjusted p values for day 2 HSV yellowness (norm.).

**Fig. SI77:**
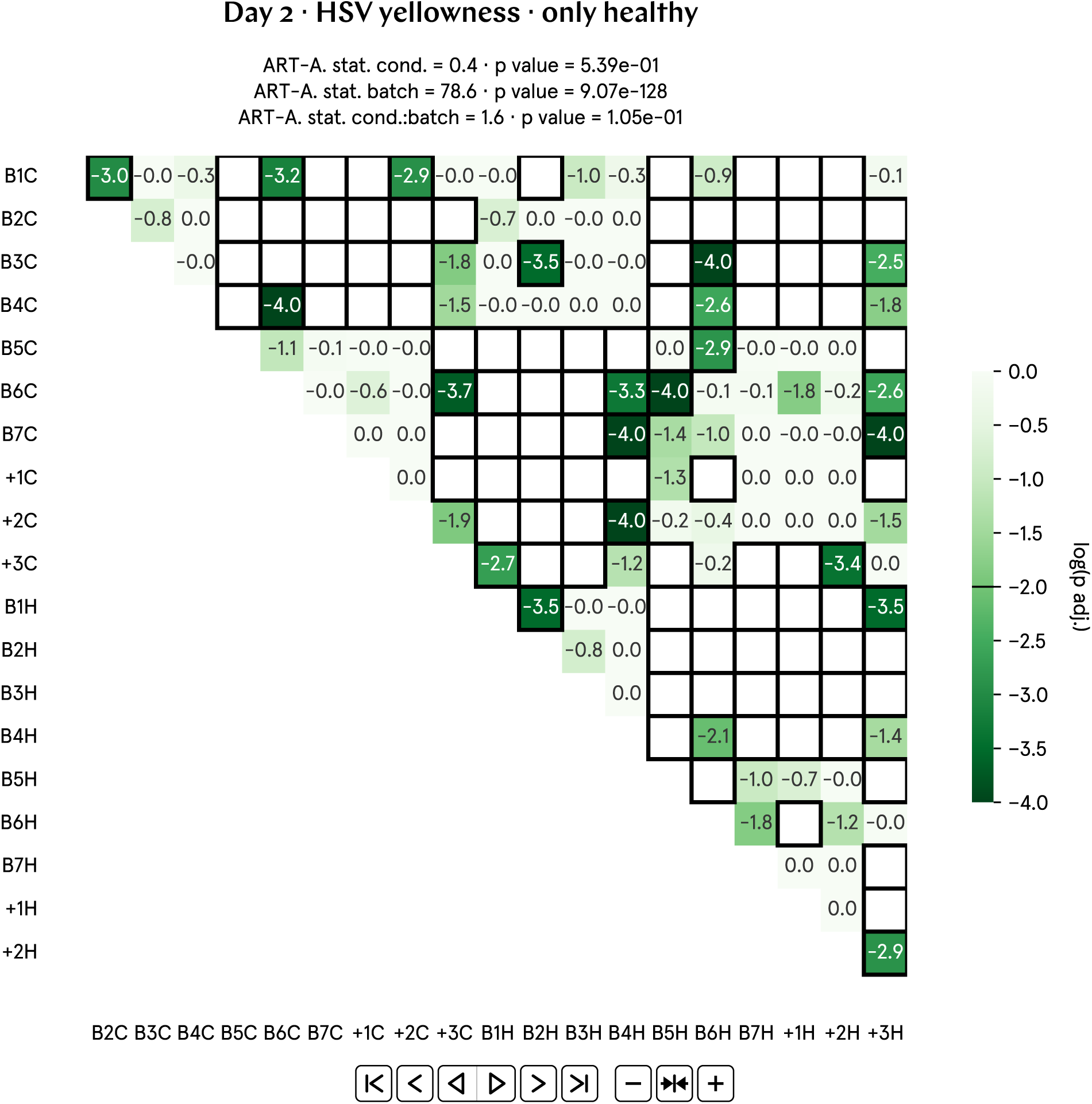
Full post-hoc matrix of adjusted p values for day 2 HSV yellowness.

**Fig. SI78:**
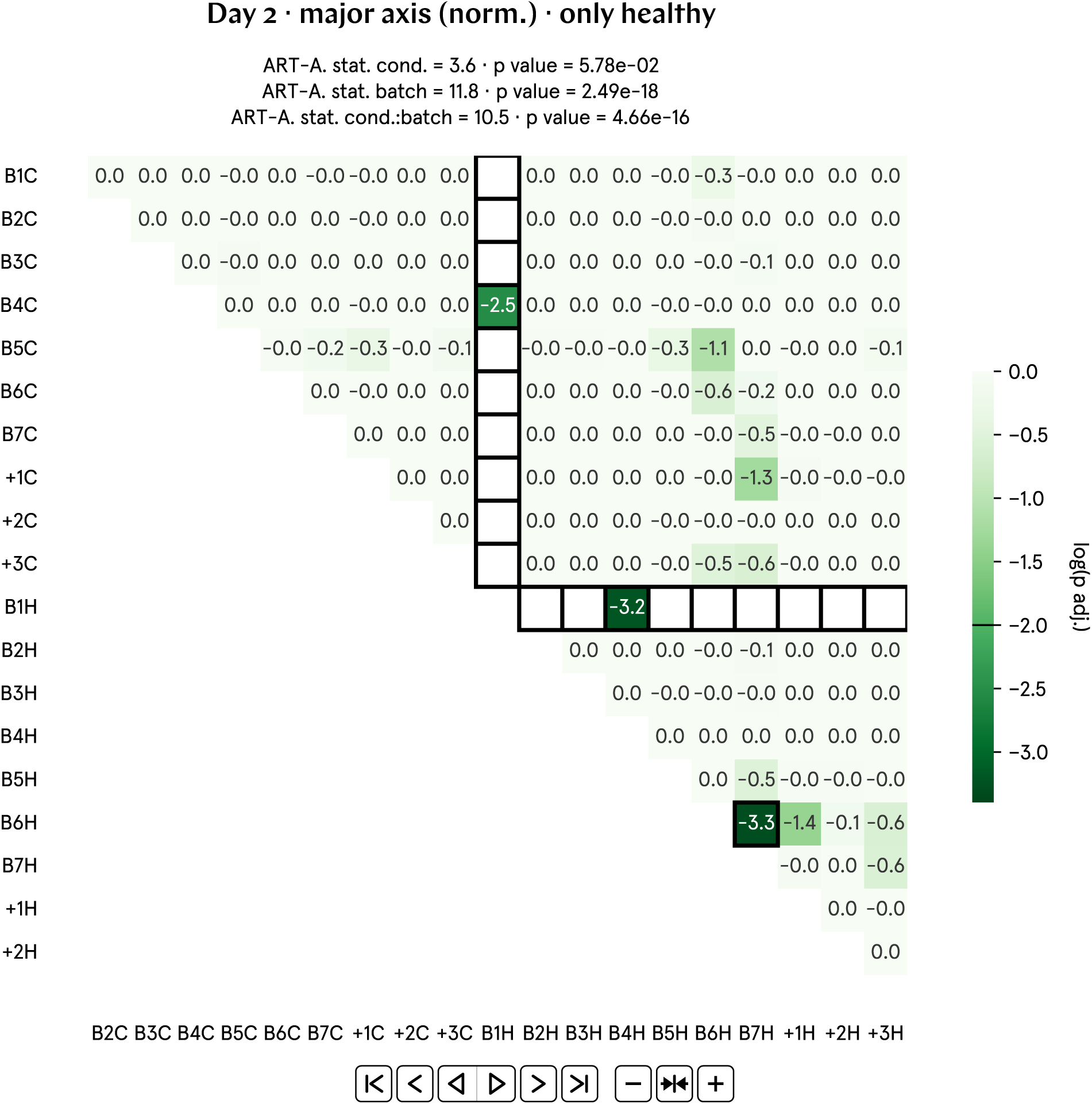
Full post-hoc matrix of adjusted p values for day 2 major axis (norm.).

**Fig. SI79:**
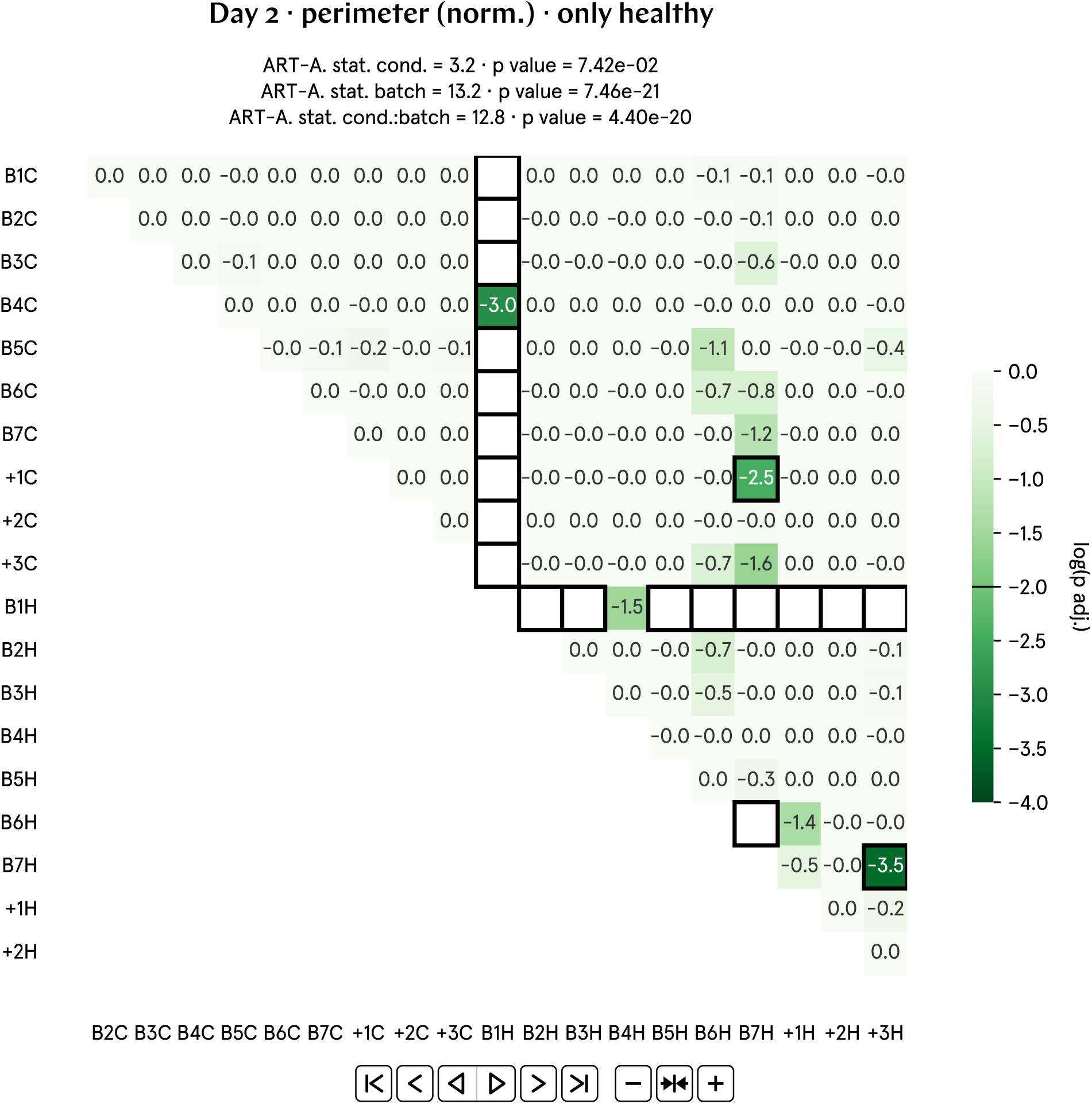
Full post-hoc matrix of adjusted p values for day 2 perimeter (norm.).

**Fig. SI80:**
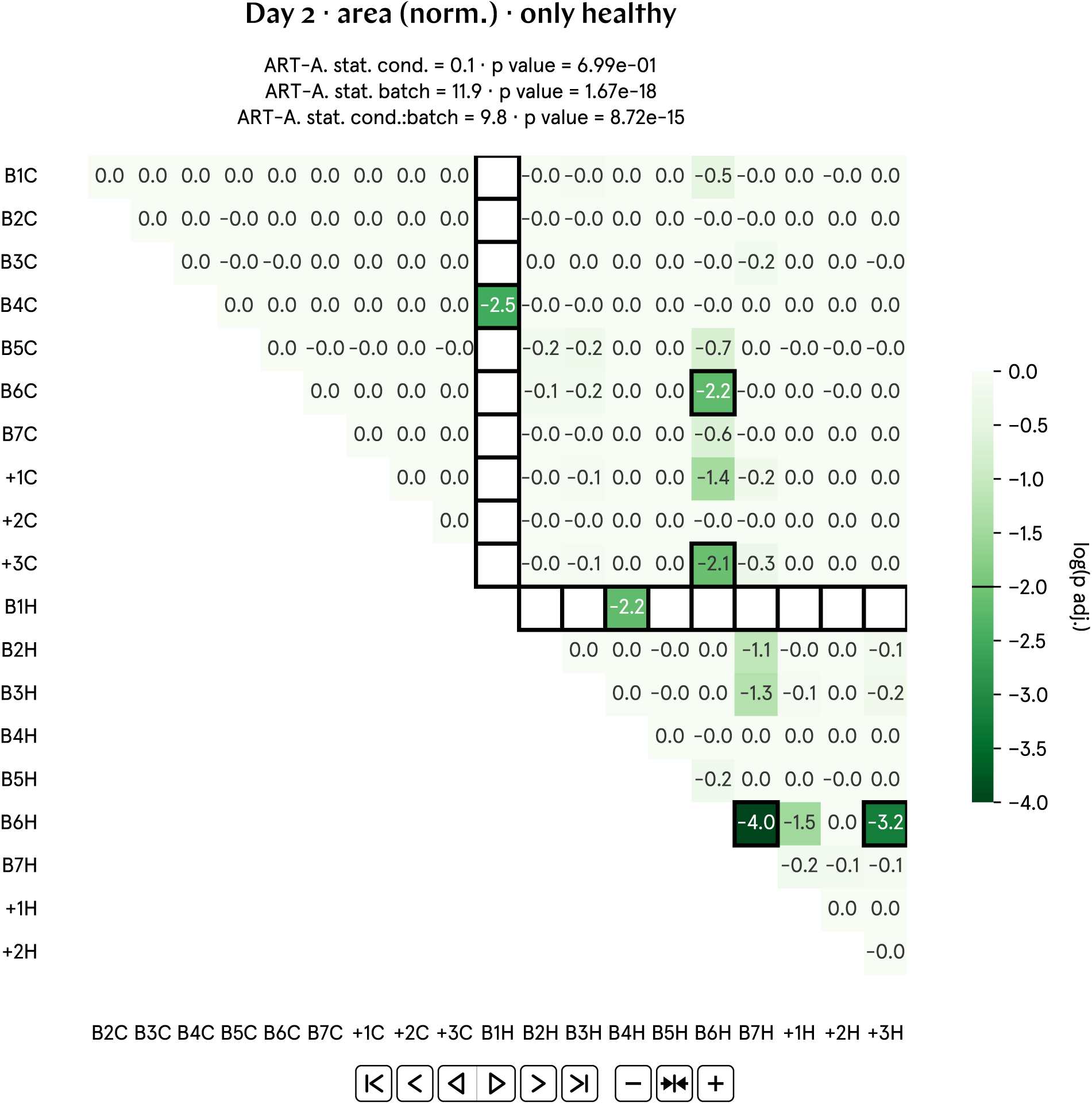
Full post-hoc matrix of adjusted p values for day 2 area (norm.).

**Fig. SI81:**
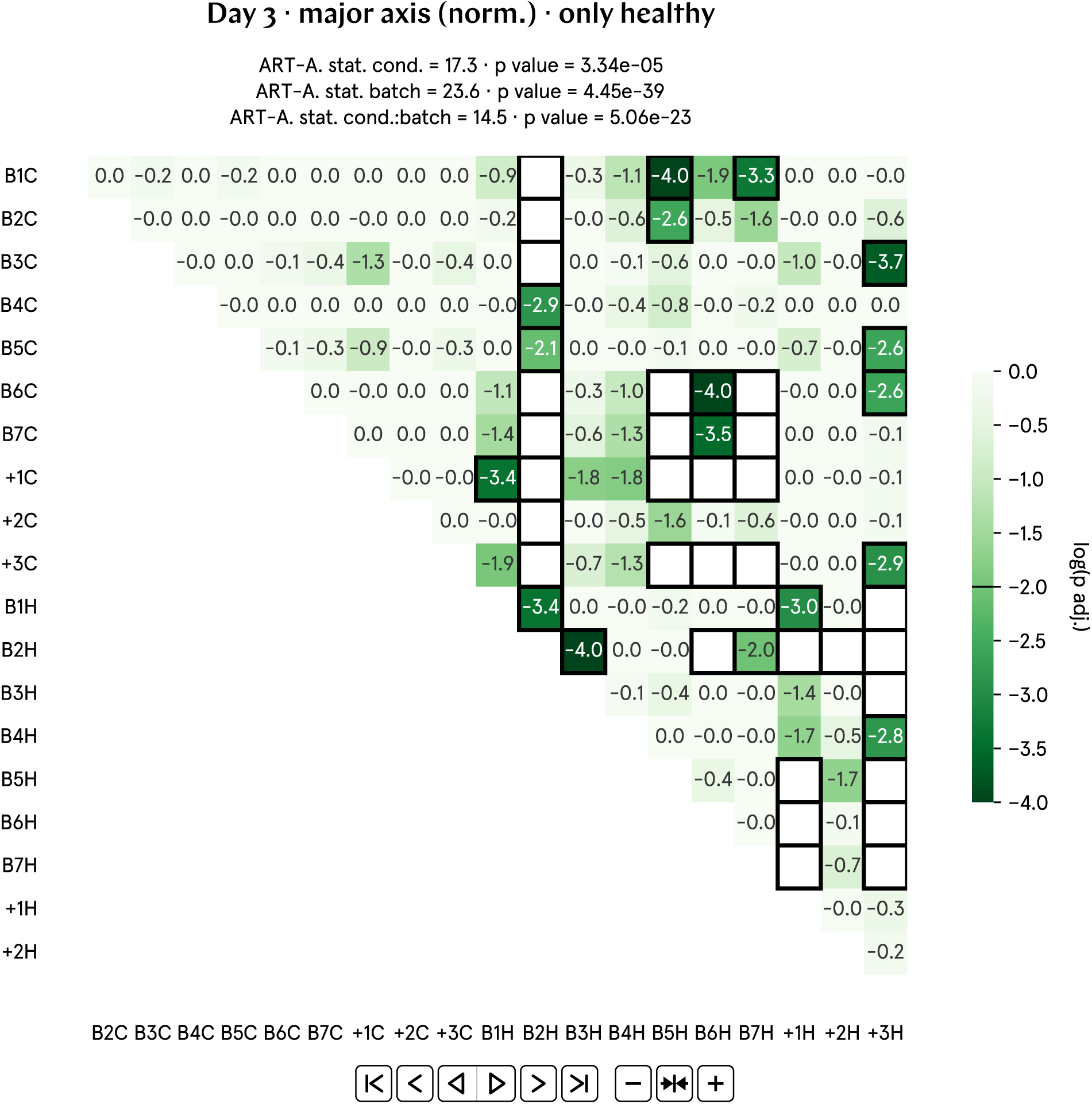
Full post-hoc matrix of adjusted p values for day 3 major axis (norm.).

**Fig. SI82:**
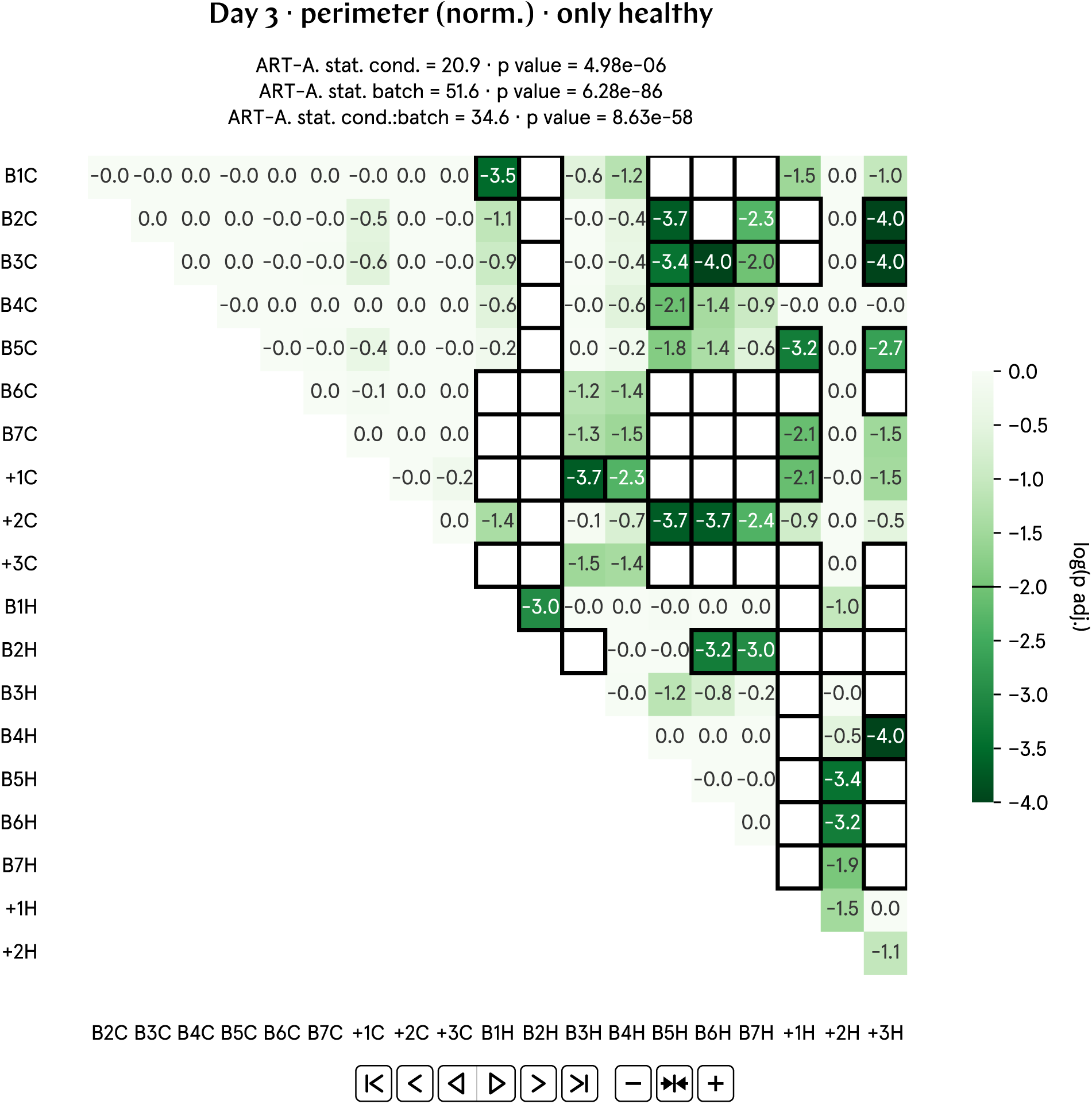
Full post-hoc matrix of adjusted p values for day 3 perimeter (norm.).

**Fig. SI83:**
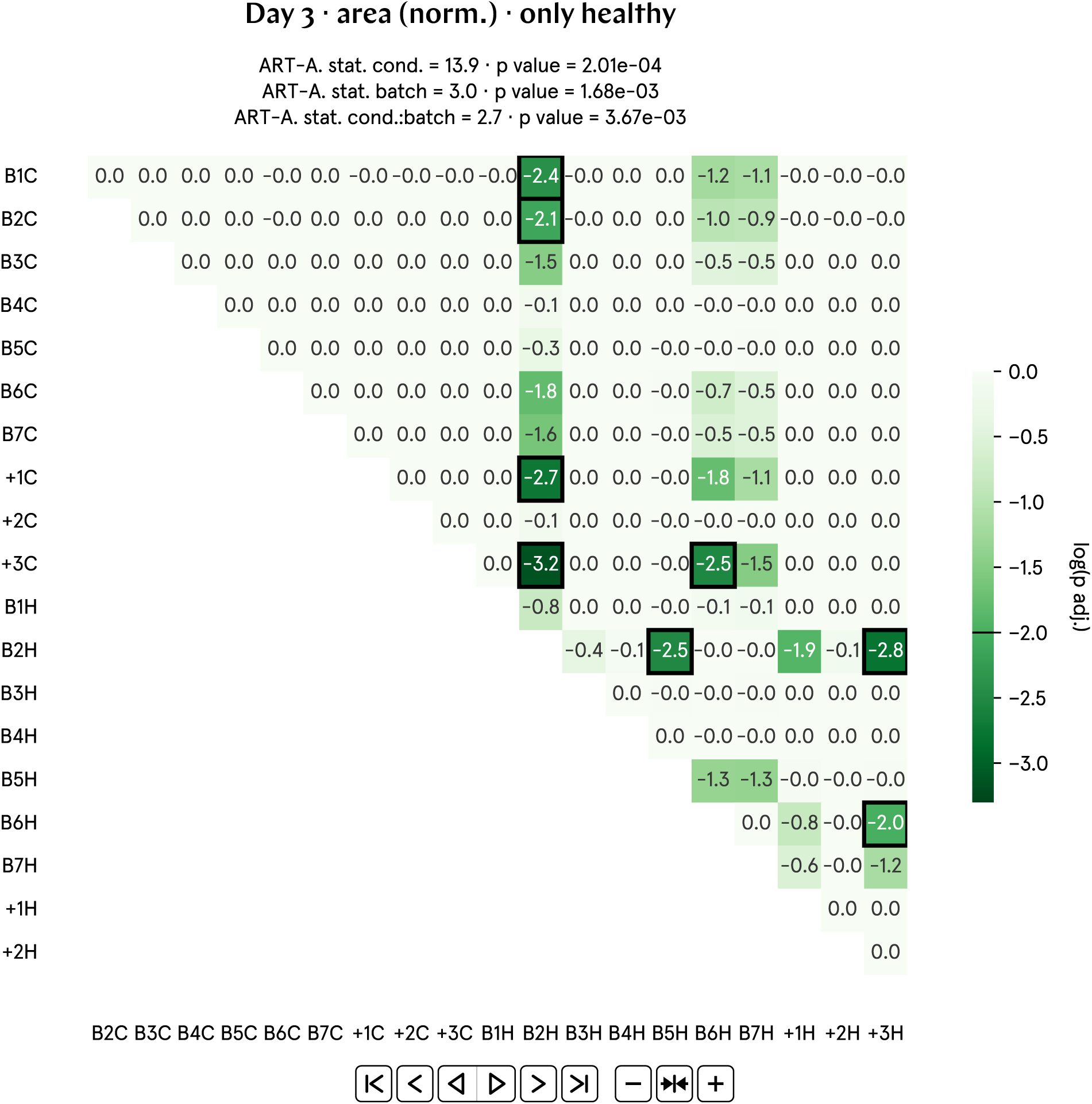
Full post-hoc matrix of adjusted p values for day 3 area (norm.).

**Fig. SI84:**
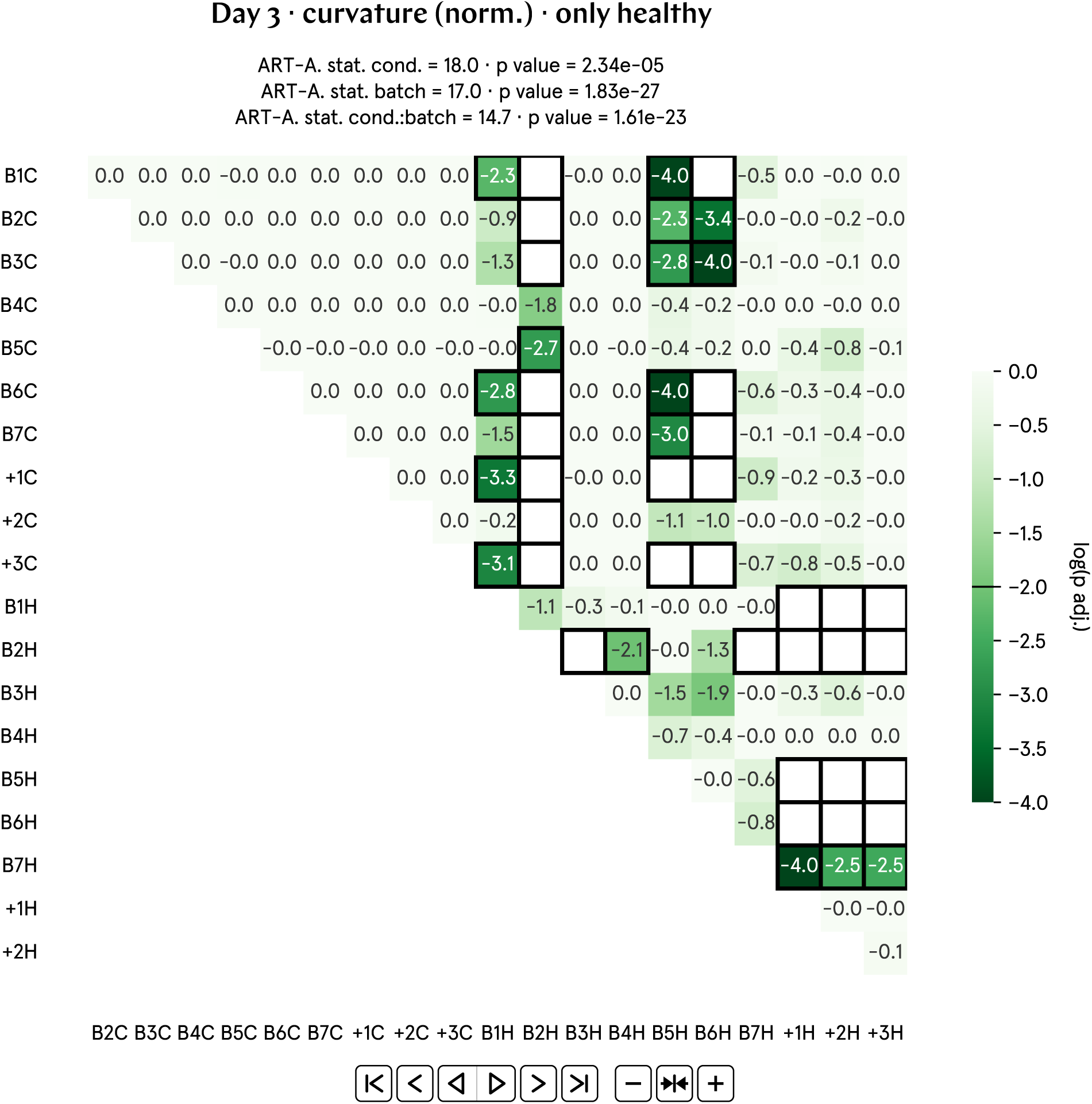
Full post-hoc matrix of adjusted p values for day 3 curvature (norm.).

**Fig. SI85:**
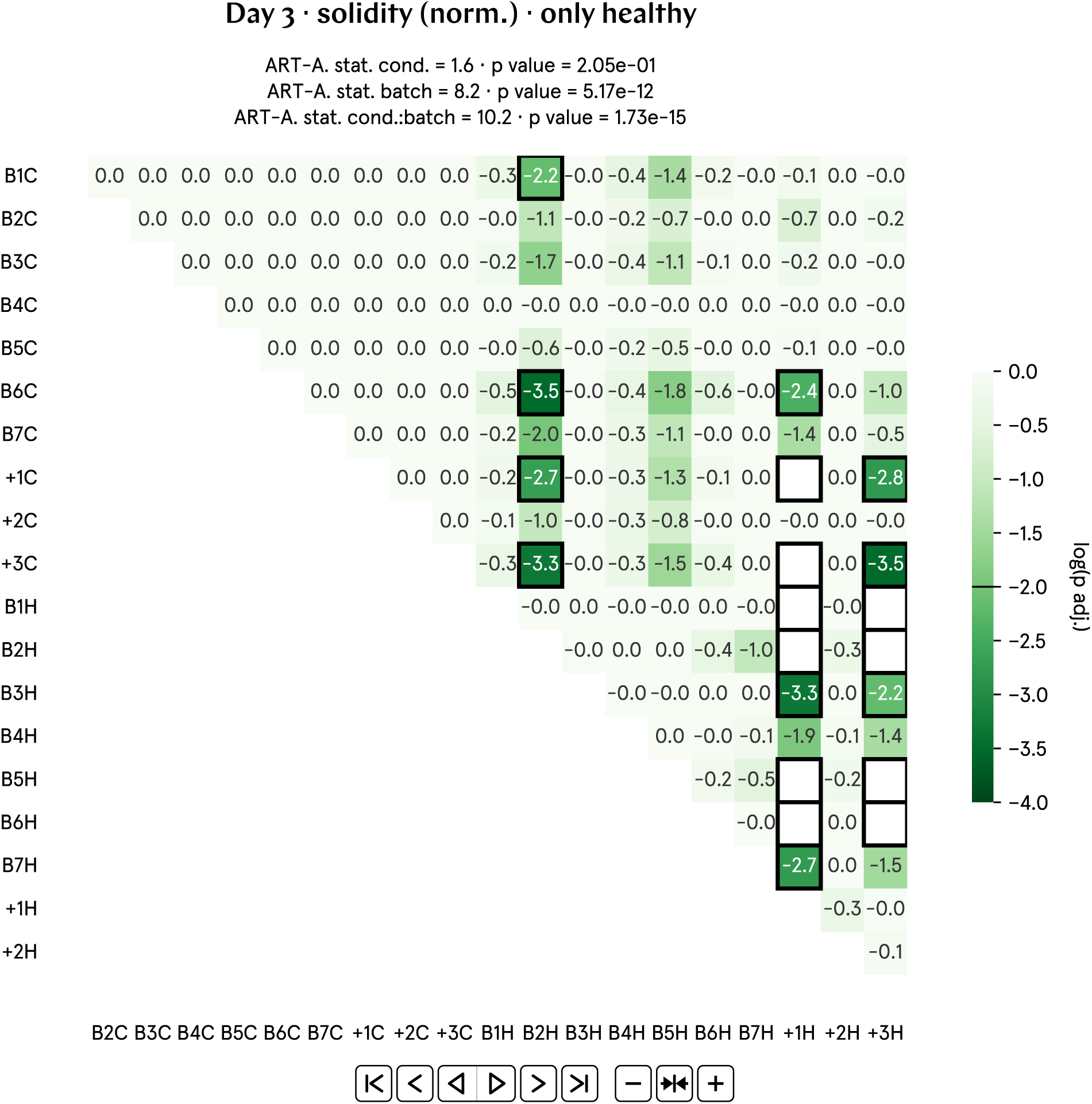
Full post-hoc matrix of adjusted p values for day 3 solidity (norm.).

**Fig. SI86:**
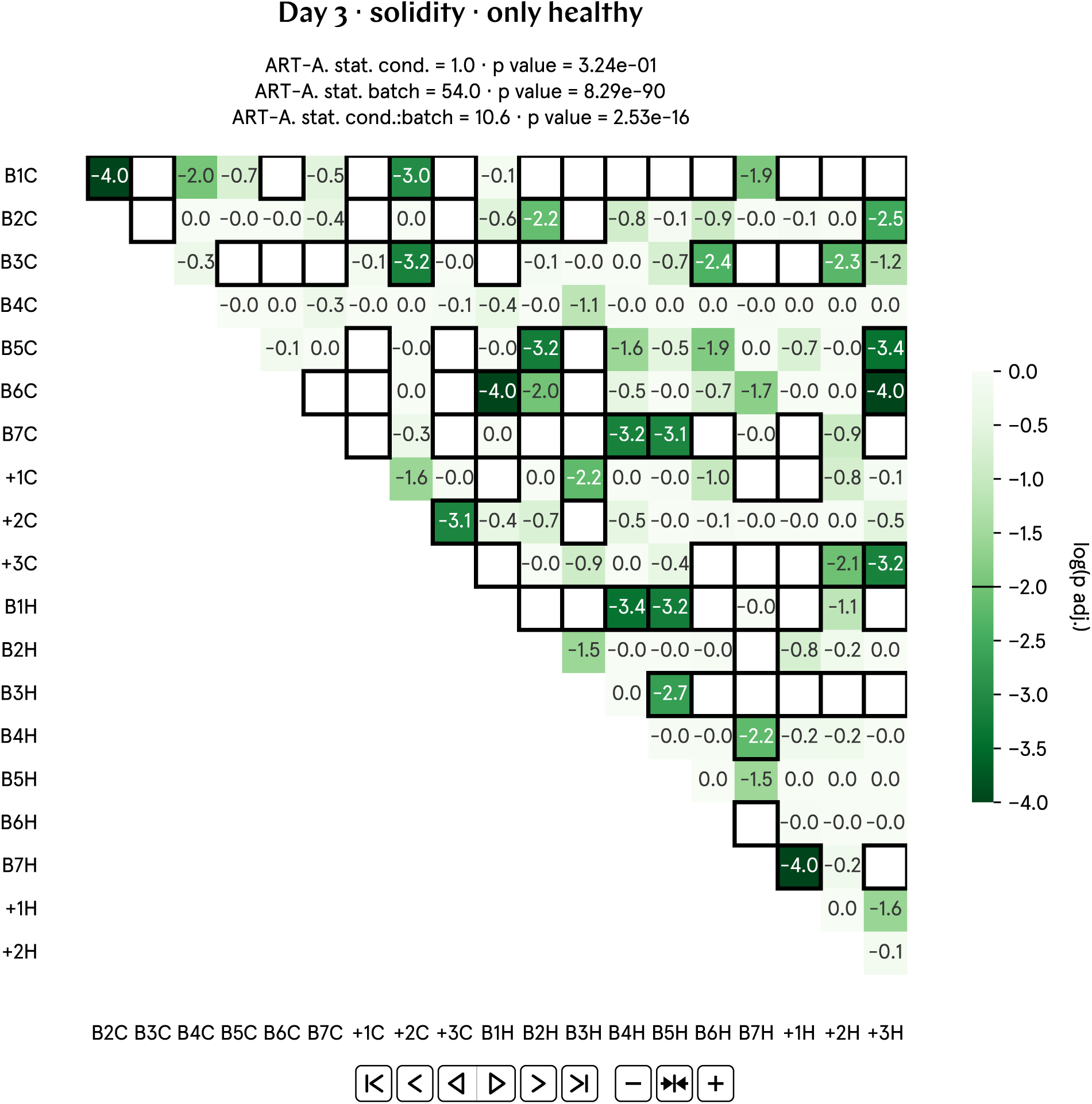
Full post-hoc matrix of adjusted p values for day 3 solidity.

**Fig. SI87:**
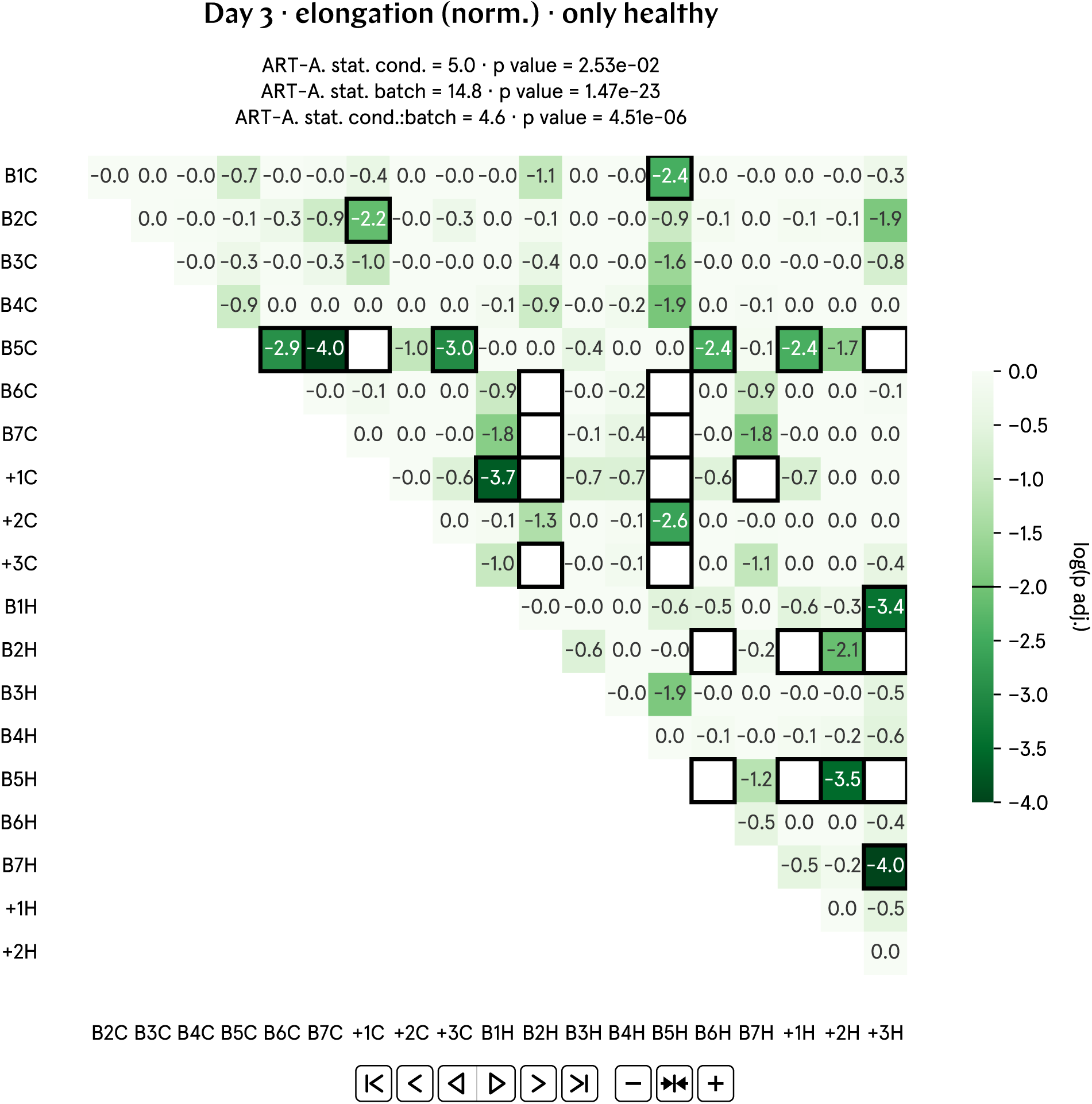
Full post-hoc matrix of adjusted p values for day 3 elongation (norm.).

**Fig. SI88:**
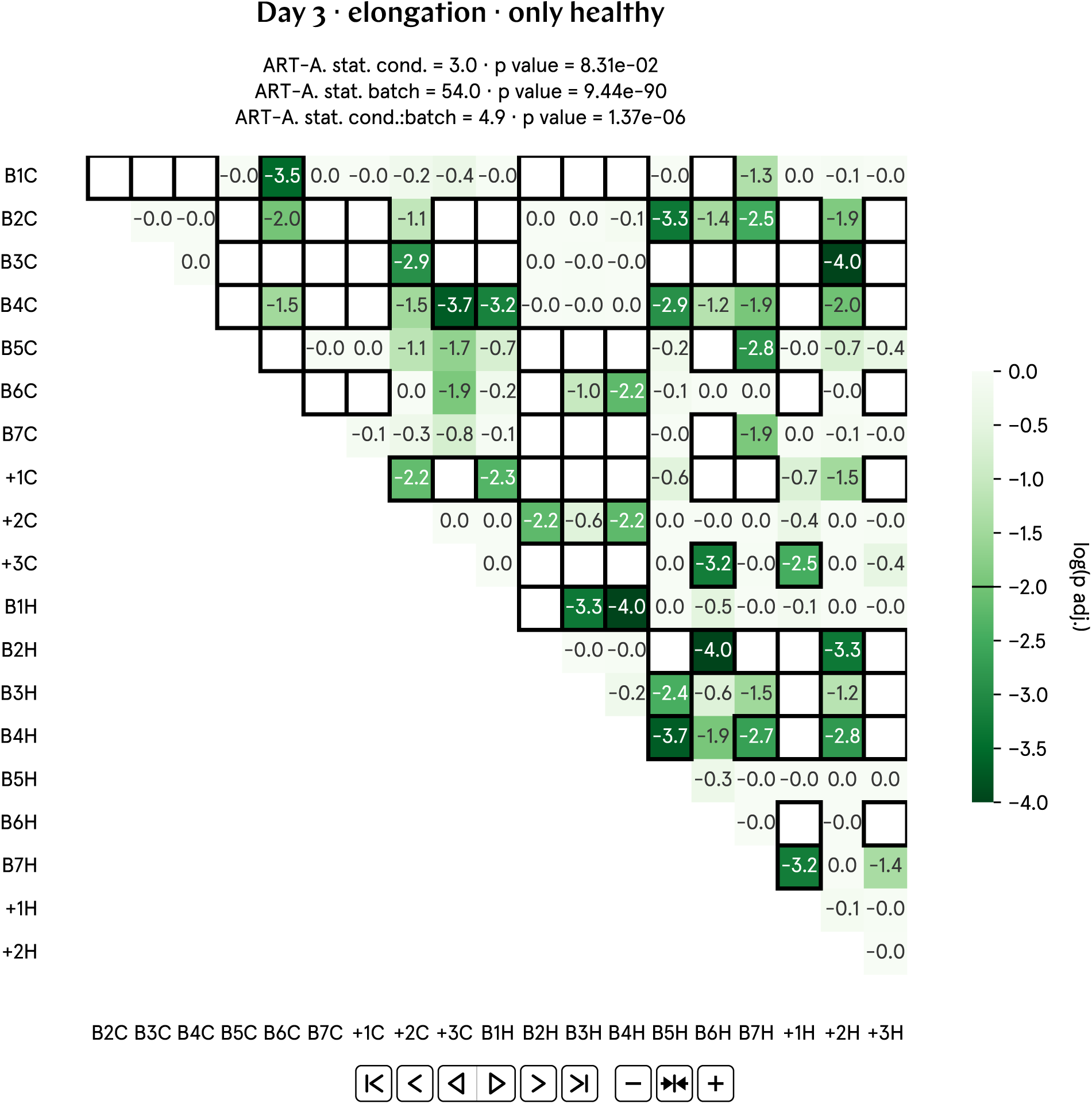
Full post-hoc matrix of adjusted p values for day 3 elongation.

**Fig. SI89:**
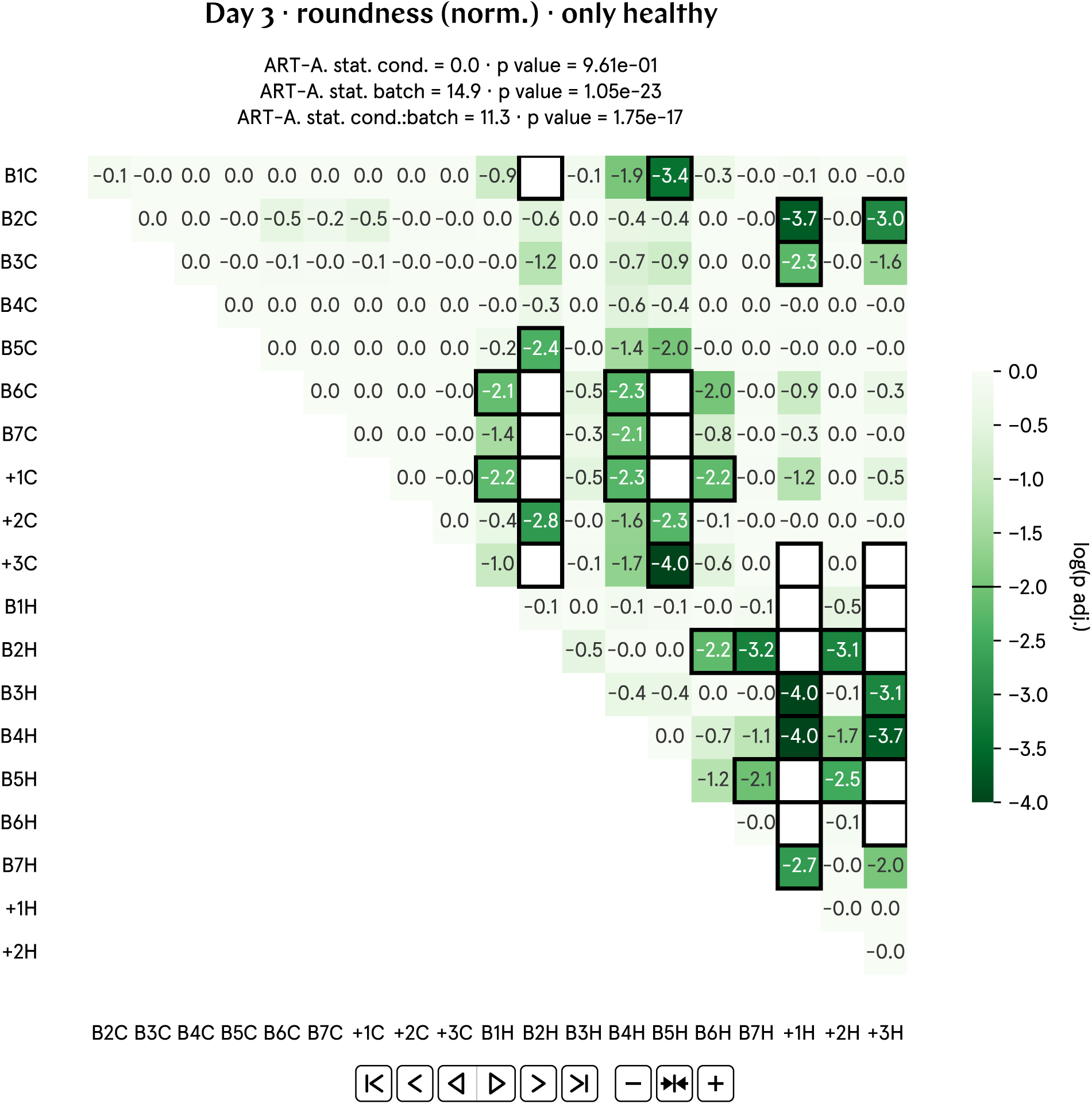
Full post-hoc matrix of adjusted p values for day 3 roundness (norm.).

**Fig. SI90:**
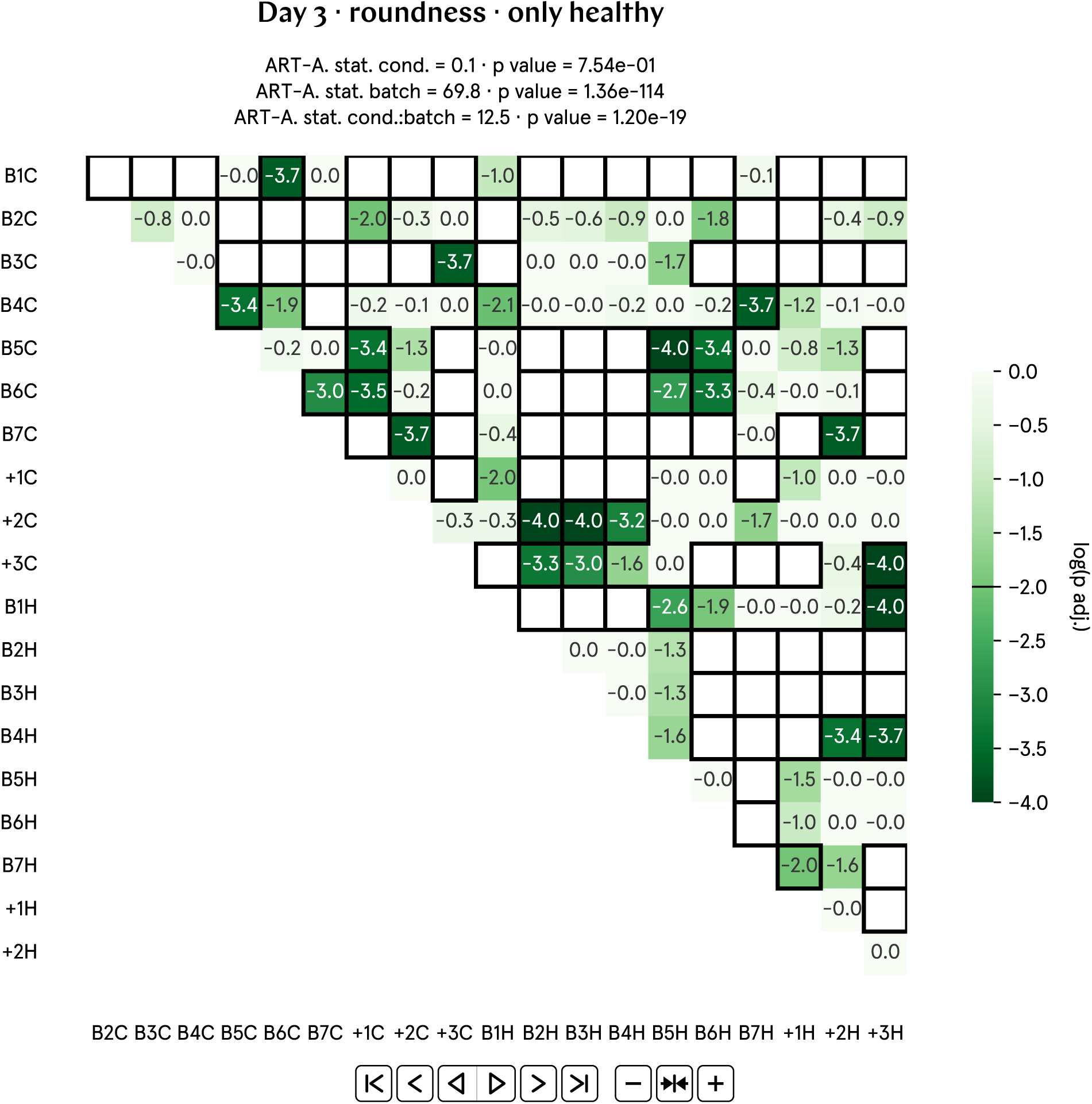
Full post-hoc matrix of adjusted p values for day 3 roundness.

### SI77. Discussion: Competing biological magnetic field sensing hypotheses favor quantum mechanisms

We have demonstrated that sensing Earth’s minute magnetic field regulates basal tadpole physiology as early as one day p.f., for purposes likely *not* related to orientation, navigation, and homing. To explain our findings, we note that, traditionally, organisms have been proposed to sense weak DC magnetic fields through three different mechanisms (for less discussed models, see [1]).

First, through electromagnetic induction [37]. Elasmobranchs such as sharks and rays have a specialized organ, the ‘ampullae of Lorenzini’, schematically consisting of channels filled with conductive material; upon crossing magnetic field lines, a voltage is induced according to Faraday’s law. It has been estimated that said channels might be sensitive enough to sense Earth’s magnetic field, but definitive proof is lacking. We find it unlikely that frog embryos possess similar organs. In addition, the detection of DC magnetic fields through this mechanism requires the organism to move and cross magnetic field lines—which is not the case for tadpole embryos at least until day 2 p.f.

Second, through biogenic magnetite (or maghemite–which is oxidated magnetite–, or greigite) clumps that act as ferrimagnetic compass needles. A key parameter in this model of organismal magnetic field sensing is the size of the ferrimagnetic clumps, which need to be large enough so that their magnetic energy (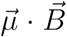, where 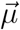 is the magnetic moment of the ferrimagnetic domain) is significantly stronger than the randomizing effects of thermal energy (*k_B_T* , where *k_B_* is the Boltzmann constant) [61]. At room temperature, single-domain magnetite crystals need to be roughly 35 nm or larger in size to have a permanent magnetic moment [35, 62]; a 35 nm-diameter sphere of magnetite has a magnetic moment of approximately 0.01 fA *·* m2, for a tiny ratio of 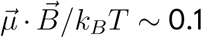 under the geofield and at room temperature. Modeling the magnetite crystal as a thermally-driven harmonic oscillator, its magnetic moment *µ* will be scattered around the direction of 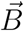 with a root-mean-square angular deviation of 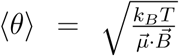; in this case, ⟨*θ*⟩ ∼ 159°. This means that biogenic magnetite that can correctly react to Earth’s magnetic field need occur as much bigger single clumps, or as smaller clumps that are so densely packed in the cell so that spin exchange interactions are strong enough to align the moments of the different magnetic domains; a ratio of 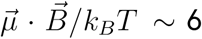 or larger, equivalently ⟨θ⟩ ∼ 23° or smaller, has been suggested as sufficient [34, 37]. The classic example here are magnetotactic bacteria that synthesize beads of single-domain magnetite enveloped by a lipid bilayer and positioned in a string known as the magnetosome; the beads, together, can exert enough torque to align a bacterium with the geofield [33]. The reported magnetic moments of magnetotactic bacteria range from 0.2 to 60 fA *·* m2 [63], or 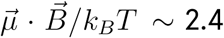 to 725, or ⟨θ⟩ ∼ 34 to 2°. It is hypothesized, but not demonstrated, that magnetic signal transduction could also be mediated by mechanosensitive ion channels that respond to the beads’ change in alignment [64], or by the incorporation of magnetite as an active component of the cells’ electric circuit [65]. The genes used by magnetotactic bacteria to absorb iron from their surroundings and synthesize magnetite seem to have few evident animal homologues [66, 67]. Biogenic magnetic nanoparticles have, however, been putatively identified across the tree of life [67]; some argue that, in the studies that use the Prussian Blue stain to label ferric iron, iron-rich macrophages are conflated with magnetite [68]. There is one isolated report of natural remanent magnetization consistent with magnetite in an amphibian, the newt [36]. A comprehensive study [68], enabled by an advance in instrumentation [63], performed large-scale cell screening for biogenic magnetite in over 10 million cells from the pigeon and the trout, organisms that had previously been singled-out through other methods [69–71] as having cells rich in magnetite for migration-related geofield sensing; of these 10 million cells, only 1 per almost half a million were found to have large magnetic moments consistent with the presence of biogenic magnetite; the smallest cellular magnetic moment reported in [63], namely 4 fA *·* m2 for a trout cell, would correspond to a sphere of magnetite of roughly 250 nm in diameter, or equivalent magnetic material sufficiently packed, for a resulting ratio 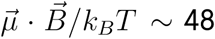 and angular deviation ⟨θ⟩ ∼ 6°; 250 nm is still below the diffraction limit of most common optical microscopes, and hence such a particle could, if present, evade simple optical detection. In tadpoles, iron, in addition to the endogenous quantity of approximately 50 ng/embryo that is maternal in origin [72] and that remains constant until developmental stage 50 [73], could come from contaminants. The MMR solution in which the tadpoles are raised (see Supplementary Section SI1) contains no iron. Iron contamination through laboratory plastics that are injection-molded into metal forms has been reported [74]; in our experiments, we find it implausible that large quantities of iron can be obtained through this scenario, since the embryos mostly just sit in plasticware after being manipulated with a plastic dropper, a practice that involves no agitation or centrifugation; during our fertilization protocol, however, the testes are macerated using a plastic piston. A *Xenopus laevis* embryo at stage 22 (roughly the stage at which our embryos are imaged on day 1 p.f., when geofield sensing is already occurring) has approximately only 30,000 cells [38]; even if enough iron for biomineralization is available endogenously or through contaminants, we argue that it is unlikely, but not impossible, that biogenic magnetite is being deployed by this mostly non-migratory species to interact with the geofield so early in the embryo development and in such a nonspecific way, given that only 0.0002% of tested cells belonging to two migratory species were positive for the presence of magnetite.

Third, organisms have been proposed to sense weak DC magnetic fields through electron spin-dependent chemical reactions. In what follows, we explain the basic elements of this hypothesis; argue that our present data is consistent with it; and expound on its features and limitations.

The net effect of a magnetic field is to change the macroscopic final products of electron spin-dependent chemical reactions [75], and the theoretical proposal is that organisms can sense a magnetic field to the extent that they can sense different physiological concentrations of such products [39]. On the one hand, in test-tube chemistry, such reactions have been unambiguously established in liquids, gases, and solids alike; at room temperature; for magnetic field strengths as weak as the Earth’s; and for a broad class of biologically relevant compounds—the most researched of which, for historical reasons related to bird magnetoreception studies, is a family of flavoproteins known as cryptochromes [9, 40] (incidentally, human cryptochrome in solution is also magnetosensitive [76]). We stress that many other proteins have also been observed to be magnetosensitive [41]; the yellow pigments in the tadpoles themselves—carotenoids [30], pteridines [77], and possibly also flavins [78]—all have rich spin-radical physics. As a general rule, magnetosensitive proteins tend to be redox-active. Compounds that can sustain electron-spin dependent chemical reactions are sensitive to varying magnetic field strengths and frequencies, even if isotropically tumbling; compounds that are, for example, anchored to a membrane could, in addition, overcome thermal randomization and be sensitive to varying magnetic field directions. Counter-intuitively, it is well-understood that the magnetic field effect size does not monotonically increase with field strength [9]; fields that are much stronger than typical hyperfine interactions (which are quoted, for example in [9], to be *∼* 0.1–5 mT) will not strongly change the outcome of the chemical reactions considered here, which make this a good hypothesis to explain our findings of biological sensitivity to a field of Earth’s strength. On the other hand, there is abundant correlative data that organisms react to magnetic fields in a way that is consistent with such electron spin-dependent chemical reactions being active inside their cells. However, at present there is no direct evidence of this fact.

Similarly to technological quantum sensing, *e.g.*, with a spin in diamond [42, 43], in electron spin-dependent chemical reactions the spins are almost always initialized (or optically pumped [79]) via photoexcitation (quantum details of the model are found in Supplementary Section SI78). (We do not believe thermally generated spin radicals could sense magnetic fields in this way.) This means that this modality of magnetic sensing can only happen in the presence of light of appropriate wavelengths and that, in biologically relevant compounds, these reactions involve chromophores; note that chromophores (*e.g.*, flavin) exist in possibly most cells, also in those that are not in contact with external light stimuli (*e.g.*, eye and skin). The tadpoles in our experiments have been raised in the dark. We note, however, that there is no such thing as absolute darkness: In Supplementary Fig. SI8 we reported the background light intensity and spectrum inside the hypomagnetic chamber and the control box, for an average illuminance of 0.0751 *±* 0.0002 lux and 0.0873 *±* 0.0003 lux, respectively. For comparison, migratory birds, which have long been proposed to sense the geofield through electron spin-dependent chemical reactions happening in their eyes, can orient themselves at dusk with light intensities of *∼* 0.0003 lux [80]; after losses and the assumption of uniform illumination of the visual field, this is estimated to correspond to just 1 photon/s per bird cell involved in magnetoreception [81]. If one posits that birds are, indeed, sensing magnetic fields in such a light-dependent way, an extraordinarily low photon flux could be influencing physiological function. This begs the question, not typically considered, of whether endogenous sources of photons could kick-start electron spin-dependent chemical reactions for magnetic modulation of basal biological function, *i.e.*, function that is *not* related to migration and homing. Photons are, for example, emitted as by-products of cellular oxidative metabolism and stress [82, 83]; in intensity, such ultra-weak, spontaneous photon emissions are way above thermal background, having been recorded to range up to several thousands of photons/s/cm2 (arguably more, if inefficiencies in the optical collection apparatus are taken into account). They are, moreover, broad in wavelength, including the blue portion of the spectrum that photoexcites flavin chromophores (for example, emissions from triplet excited carbonyls and from singlet-excited melanin are found, respectively, at 350 and 360 nm and higher wavelengths). Our background spectra, positively, and endogenous ultra-weak photon emission, perhaps, could thus provide a photon flux that warrants spin-based magnetic field sensing in both the hypomagnetic chamber and the control box.

Thus, in this model, magnetic sensing in biology has features matching decades of experimental observations across the tree of life. In particular, magnetic sensing: should not have biological specificity (unless spin physics information is used to perform magnetic engineering), which is consistent with effects that have been indiscriminately reported across organisms and cell types; should, instead, be widespread across a vast gamut of biological compounds that are magnetosensitive; should present a low correlation between effect magnitudes and magnetic field strengths or exposure durations, as it is indeed reported across many experiments [1]; and should not require a dedicated sensory organ (except maybe in migratory species). One important shortfall of the hypothesis is that the size of magnetic field effects on electron spin-dependent chemical reactions is typically small (at most a single-digit percent change of final products); moreover, it is at present unknown how magnetic signals could be amplified and/or transduced into statistically significant physiological and phenotypical differences as seen in our data and elsewhere.

If a subset of all observed weak magnetic field effects in biology are a result of the above described quantum sensing-like chemical reactions happening inside cells, the implications will be tremendous. This will mean, first, that electron spin superpositions survive the strongly decohering environment of the cell *for long enough to be used for function* (*i.e.*, to sense and physiologically react to a magnetic field; see timescale estimates in Supplementary Section SI78); and second, that knowledge of the endogenous spin physics properties of biological matter can be used to engineer weak magnetic fields to up-and down-regulate the whole machinery of the cell.

### SI78. Quantum features of magnetic field sensing via electron spin-dependent chemical reactions

#### SI78.1. Electron spin-dependent chemical reactions as quantum sensing

Electron spin-dependent chemical reactions should be understood as *bona fide* quantum sensing processes: In them, an electron spin radical or, more commonly, pairs of spin radicals are driven into a superposition by a combination of the externally applied magnetic field and the internal magnetic field due to hyperfine couplings to nearby nuclear spins. *The superposition performs the magnetic field sensing*—similarly to the simplest modality of technological quantum sensing known to physicists, namely magnetic field sensing via Ramsey interferometry [84] with, *e.g.*, an electron spin in diamond [85]. Different magnetic fields alter the time evolution of the superposition, thus also altering the dynamic probability that the electron spins be measured by the environment in each superposition state. To each collapsed electron spin state there corresponds one chemical pathway taken; in practice, it is believed that different electron spin states influence chemical conformation or electron transport rates.

#### SI78.2. Time and energy scales

Magnetic fields affect the electron spin superposition. Even after the superposition is long decohered, its magnetosensing effects can still be macroscopically relevant, since, on average (for many compounds concomitantly sensing the magnetic field, or for one compound sensing the magnetic field repeated times), the superposition determined how often each diverging chemical pathway was taken. Here, thermal energies *k_B_T* are indeed much larger than typical magnetic energies involved (for example, the energy difference between the electron spin superposition states); the crux of the issue is that *quantum-assisted magnetic sensing only occurs before thermalization: The argument relies on timescales, not energy scales*. For a conservative estimate, a single electron spin in a superposition can be understood to sense a magnetic field if it can Larmor precess around said field once before undergoing decoherence. The Larmor period of a free electron around Earth’s magnetic field is 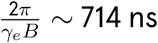, with *γ_e_* the gyromagnetic ratio of the electron; in this estimate, the electron spin superposition coherence time (analogous to T* in Ramsey interferometry [84, 85]) needs to be at least τ ∼ 714 ns in order for it to sense the geofield.

Using the Nyquist criterion instead yields a less conservative estimate 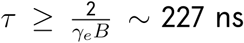. That such coherence times are relatively long and have never been directly measured for biocompounds inside the wet, warm environment of a single cell are the main contention points for this hypothesis, for which corroborating, but not causal, biological data abound. If further experimentation suggests that our present results are indeed explained through this hypothesis, the profound implication of that eventuality will be that electron spin superpositions are maintained inside the tadpoles’ cells for at least the above estimated coherence times.

#### SI78.3. Sensitivity considerations

For the magnetite hypothesis, the sensitivity *η* (in magnetic field resolution per unit time) is given by [65, 86]:

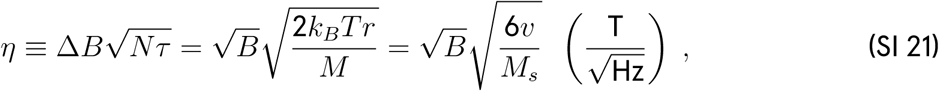

where *Nτ* is the total sensing time; 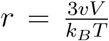is the characteristic rotational response time of the magnetite particle; *v* is the dynamic viscosity of the medium; *V* is the volume of the magnetite grain; *M* = *M_s_V* is the magnetic moment of the sphere; and *M_s_ ∼* 4.46 *·* 105 A/m is the saturation magnetization of magnetite. We assume here that the cellular cytoplasm is about 5.5 more viscous than water, as suggested elsewhere [87]. This yields *η ∼* 5.7 *·* 10^*−*2^ 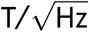 for a grain of magnetite.

In contrast, a freely evolving single-electron spin superposition can sense a magnetic field with a (Ramsey-like) sensitivity [85]

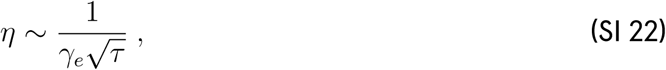

where *γ_e_ ∼* 1.7 *·* 1011 rad/(s *·* T) for the electron, assumed uncoupled, and *τ* is the sensing time. To sense Earth’s magnetic field with an electron spin superposition, a time of *∼* 714 ns is required as roughly estimated above; this yields 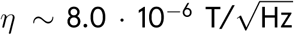, a sensitivity improvement by several orders of magnitude compared to the magnetite model.

#### SI78.4. Magnitude of effect as a function of field strength

Differently from quantum sensing with a single electron spin, quantum sensing with a pair of electron spins presents the counter-intuitive feature that *externally applied magnetic fields alter the final products of the chemical reaction by a proportion that is not monotonically increasing with field strength*. There is a well-understood competition between the external magnetic field and the endogenous hyperfine field; fields that are much stronger than typical hyperfine interactions will not strongly change the outcome of the chemical reaction.

As a rule of thumb, external magnetic fields larger than tens of mT will not influence electron spin-dependent chemical reactions. However, very strong external magnetic fields (usually greater than a few tesla) can trigger another spin mechanism, known as ‘delta g’, to once again influence the distribution of chemical reaction products [88].

## Bibliography

[1] V. N. Binhi and F. S. Prato, Biological effects of the hypomagnetic field: An analytical review of experiments and theories, PLoS ONE 12, e0179340 (2017).

[2] Z.-H. Hadi and C. Simon, Magnetic field effects in biology from the perspective of the radical pair mechanism, J. R. Soc. Interface. 19, 20220325 (2022).

[3] T. J. Zwang, E. C. M. Tse, D. Zhong, and J. K. Barton, A compass at weak magnetic fields using thymine dimer repair, ACS Cent. Sci. 4, 329 (2018).

[4] R. J. Usselman, C. Chavarriaga, P. R. Castello, M. Procopio, T. Ritz, E. A. Dratz, D. J. Singel, and C. F. Martino, The quantum biology of reactive oxygen species partitioning impacts cellular bioenergetics, Sci. Rep. 6, 38543 (2016).

[5] H. Wang and X. Zhang, Magnetic fields and reactive oxygen species, Int. J. Mol. Sci. 18, 2175 (2017).

[6] W.-C. Mo, Y. Liu, H. M. Cooper, and R.-Q. He, Altered development of *Xenopus* embryos in a hypogeomagnetic field, Bioelectromagnetics 33, 238 (2012).

[7] A. Bahrami, L. Y. Tanaka, R. C. Massucatto, F. R. M. Laurindo, and C. D. Aiello, Automated 1D Helmholtz coil design for cell biology: Weak magnetic fields alter cytoskeleton dynamics (2024), arXiv:2406.19555.

[8] F. Bertagna, R. Lewis, S. R. P. Silva, J. McFadden, and K. Jeevaratnam, Effects of electromagnetic fields on neuronal ion channels: A systematic review, Ann. N. Y. Acad. Sci. 1499, 82 (2021).

[9] P. J. Hore and H. Mouritsen, The radical-pair mechanism of magnetoreception, Annu. Rev. Biophys. 45, 299 (2016).

[10] L. Zastko, L. Makinistian, A. Tvarožná, et al., Mapping of static magnetic fields near the surface of mobile phones, Sci. Rep. 11, 19002 (2021).

[11] F. Hauksbee, An account of an experiment touching the production of light *in vacuo*, Philos. Trans. R. Soc. Lond. 26, 149 (1709).

[12] L. Landler and G. Gollmann, Magnetic orientation of the common toad: Establishing an arena approach for adult anurans, Front. Zool. 8, 6 (2011).

[13] M. K. Khokha, C. Chung, E. L. Bustamante, L. W. Gaw, K. A. Trott, J. Yeh, N. Lim, J. C. Lin, N. Taverner, E. Amaya, and N. Papalopulu, Techniques and probes for the study of *Xenopus tropicalis* development, Dev. Dyn. 225, 499 (2002).

[14] A. Leibovich, T. Edri, S. L. Klein, S. A. Moody, and A. Fainsod, Natural size variation among embryos leads to the corresponding scaling in gene expression, Dev. Biol. 462, 165 (2020).

[15] P. D. Nieuwkoop and J. Faber, Normal Table of Xenopus laevis (Daudin): A Systematical and Chronological Survey of the Development from the Fertilized Egg till the End of Metamorphosis, 2nd ed. (Garland Pub., 1994).

[16] Xenbase, Xenbase Anatomy and Developmental Stages (2024), accessed: October 12, 2024.

[17] J. H. Zar, Biostatistical Analysis, 5th ed. (Pearson, Upper Saddle River, NJ, 2010).

[18] J. E. Mir, A. Nasrallah, N. Thézé, M. Cario, H. Fayyad-Kazan, P. Thiébaud, and H.-R. Rezvani, *Xenopus* as a model system for studying pigmentation and pigmentary disorders, Pigment Cell Melanoma Res. 00, 1 (2024).

[19] J. T. Bagnara, M. E. Hadley, and J. H. Taylor, Regulation of bright-colored pigmentation of amphibians, Gen. Comp. Endocrinol. 2, 425 (1969).

[20] M. D. Fairchild, Color Appearance Models, 3rd ed. (John Wiley & Sons, 2013).

[21] J. T. Bagnara and M. E. Hadley, Chromatophores and Color Change: The Comparative Physiology of Animal Pigmentation (Prentice Hall, Englewood Cliffs, NJ, 1973).

[22] P. Jorgensen, J. A. J. Steen, H. Steen, and M. W. Kirschner, The mechanism and pattern of yolk consumption provide insight into embryonic nutrition in *Xenopus*, Development 136, 1539 (2009).

[23] J. D. Romano and T. R. Ziegler, Lipoproteins and carotenoids: Implications in embryogenesis and tissue differentiation, in Carotenoids: Volume 4: Natural Functions, edited by G. Britton, S. Liaaen-Jensen, and H. Pfander (Birkhäuser Verlag, 2002).

[24] G. E. Bertolesi, N. Debnath, H. R. Malik, L. L. H. Man, and S. McFarlane, Type II opsins in the eye, the pineal complex and the skin of *Xenopus laevis*: Using changes in skin pigmentation as a readout of visual and circadian activity, Front. Neuroanat. 15, 784478 (2021).

[25] H. Liedtke, K. Lopez-Hervas, I. Galván, and, et al., Background matching through fast and reversible melanin-based pigmentation plasticity in tadpoles comes with morphological and antioxidant changes, Sci. Rep. 13, 12064 (2023).

[26] M. Asashima, K. Shimada, and C. J. Pfeiffer, Magnetic shielding induces early developmental abnormalities in the newt, *Cynops pyrrhogaster*, Bioelectromagnetics 12, 215 (1991).

[27] S. Grimaldi, D. Pozzi, A. Lisi, S. Rieti, V. Manni, G. Ravagnan, L. Giuliani, T. Eremenko, and P. Volpe, Influence of the magnetic field on the tadpole metamorphosis, Int. J. Radiat. Med. 1, 96 (2000).

[28] M. Severini, A. M. Dattilo, and A. D. Gaetano, Sublethal effect of a weak intermittent magnetic field on the development of *Xenopus laevis* (daudin) tadpoles, Int. J. Biometeorol. 48, 91 (2003).

[29] H. Z. Haghighi, G. Bertolesi, C. Simon, and S. McFarlane, Weak magnetic field effects on pigmentation in tadpoles, in APS March Meeting (American Physical Society, Minneapolis, 2024) Session BB02: Biological Physics at the Molecular Scale.

[30] G. Britton, S. Liaaen-Jensen, and H. Pfander, eds., Carotenoids: Volume 4: Natural Functions (Birkhäuser Verlag, 2008).

[31] R. Rishabh, H. Zadeh-Haghighi, D. Salahub, and C. Simon, Radical pairs may explain reactive oxygen species-mediated effects of hypomagnetic field on neurogenesis, PLoS Comput. Biol. 18, e1010198 (2022).

[32] L. Tian, Y. Luo, A. Zhan, J. Ren, H. Qin, and Y. Pan, Hypomagnetic field induces the production of reactive oxygen species and cognitive deficits in mice hippocampus, Int. J. Mol. Sci. 23, 3622 (2022).

[33] C. T. Lefevre and D. A. Bazylinski, Ecology, diversity, and evolution of magnetotactic bacteria, Microbiol. Mol. Biol. Rev. 77, 497 (2013).

[34] J. Kirschvink and M. Walker, Particle size considerations for magnetite-based magnetoreceptors, in Magnetite Biomineralization and Magnetoreception in Organisms: A New Biomagnetism, Topics in Geobiology, Vol. 5, edited by J. Kirschvink, D. Jones, and B. McFadden (Plenum Press, New York, 1985).

[35] M. Meister, Physical limits to magnetogenetics, eLife 5, e17210 (2016).

[36] J. Brassart, J. L. Kirschvink, J. B. Phillips, and S. C. Borland, Ferromagnetic material in the eastern red-spotted newt *Notophthalmus viridescens*, J. Exp. Biol. 202, 3155– (1999).

[37] S. Johnsen and K. J. Lohmann, Magnetoreception in animals, Phys. Today 61, 29 (2008).

[38] J. Cooke, Properties of the primary organization field in the embryo of *Xenopus laevis*. IV. Pattern formation and regulation following early inhibition of mitosis, J. Embryol. Exp. Morphol. 30, 49 (1973).

[39] K. Schulten, C. E. Swenberg, and A. Weller, A biomagnetic sensory mechanism based on magnetic field modulated coherent electron spin motion, Z. Phys. Chem. 111, 1 (1978).

[40] T. Ritz, S. Adem, and K. Schulten, A model for photoreceptor-based magnetoreception in birds, Biophys. J. 78, 707 (2000).

[41] A. R. Jones, Magnetic field effects in proteins, Mol. Phys. 114, 1691 (2016).

[42] J. R. Maze, P. L. Stanwix, J. S. Hodges, S. Hong, J. M. Taylor, P. Cappellaro, L. Jiang, M. V. G. Dutt, E. Togan, A. S. Zibrov, A. Yacoby, R. L. Walsworth, and M. D. Lukin, Nanoscale magnetic sensing with an individual electronic spin in diamond, Nature 455, 644 (2008).

[43] J. M. Taylor, P. Cappellaro, L. Childress, L. Jiang, D. Budker, P. R. Hemmer, A. Yacoby, R. Walsworth, and M. D. Lukin, High-sensitivity diamond magnetometer with nanoscale resolution, Nat. Phys. 7, 270 (2008).

[44] H. Lee, N. Yang, and A. E. Cohen, Mapping nanomagnetic fields using a radical pair reaction, Nano Lett. 11, 5367 (2011).

[45] A. V. V. Huizen, J. M. Morton, L. J. Kinsey, D. G. V. Kannon, M. A. Saad, T. R. Birkholz, J. M. Czajka, J. Cyrus, F. S. Barnes, and W. S. Beane, Weak magnetic fields alter stem cell–mediated growth, Sci. Adv. 5, eaau7201 (2019).

[46] N. Ikeya and J. R. Woodward, Cellular autofluorescence is magnetic field sensitive, Proc. Natl. Acad. Sci. U.S.A. 118, e2018043118 (2021).

[47] Contributor role taxonomy (CRediT), https://credit.niso.org/ (2024), accessed: October 12, 2024.

[48] A. Marblestone, A. Gamick, T. Kalil, C. Martin, M. Cvitkovic, and S. G. Rodriques, Unblock research bottlenecks with non-profit start-ups, Nature 601, 188 (2022).

[49] S. Mcnamara, M. Wlizla, and M. E. Horb, Husbandry, general care, and transportation of *Xenopus laevis* and *Xenopus tropicalis*, Methods Mol. Biol. 1865, 1 (2018).

[50] M. Wlizla, S. McNamara, and M. E. Horb, Generation and care of *Xenopus laevis* and *Xenopus tropicalis* embryos, Methods Mol. Biol. 1865, 19 (2018).

[51 ] D. Budker, personal communication (2019).

[52] P. Fierlinger, personal communication (2019).

[53] A. E. Ruark and M. F. Peters, Helmholtz coils for producing uniform magnetic fields, J. Opt. Soc. Am. 13, 205 (1926).

[54] C. Kindermann and J.-M. Hero, Pigment cell distribution in a rapid colour changing amphibian (*Litoria wilcoxii*), Zoomorphology 135, 197 (2016).

[55] C. T. Rueden, J. Schindelin, M. C. Hiner, B. E. DeZonia, A. E. Walter, E. T. Arena, and K. W. Eliceiri, ImageJ2: ImageJ for the next generation of scientific image data, BMC Bioinformatics 18, 529 (2017).

[56] J. Cohen, Statistical Power Analysis for the Behavioral Sciences, 2nd ed. (Lawrence Erlbaum Associates, Hillsdale, NJ, 1988).

[57] N. Cliff, Dominance statistics: Ordinal analyses to answer ordinal questions, Psychol. Bull. 114, 494 (1993).

[58] J. Goedhart, Quantification of differences as an alternative to p values, Blog post on The Node (2018), accessed: October 12, 2024.

[59] The American Society for Testing and Materials, Test method for yellowness index of plastics, ASTM Standards D1925-70 (1970).

[60] A. R. Smith, Color gamut transform pairs, SIGGRAPH Comput. Graph. 12, 12 (1978).

[61] R. K. Adair, Constraints of thermal noise on the effects of weak 60-Hz magnetic fields acting on biological magnetite, Proc. Natl. Acad. Sci. U.S.A. 91, 2925 (1994).

[62] D. J. Dunlop, Magnetite: Behavior near the single-domain threshold, Science 176, 41 (1972).

[63] S. H. Eder, H. Cadiou, A. Muhamad, and M. Winklhofer, Magnetic characterization of isolated candidate vertebrate magnetoreceptor cells, Proc. Natl. Acad. Sci. U.S.A. 109, 12022 (2012).

[64] B. L. Clites and J. T. Pierce, Identifying cellular and molecular mechanisms for magnetosensation, Annu. Rev. Neurosci. 40, 231 (2017).

[65] J. L. Kirschvink and J. L. Gould, Biogenic magnetite as a basis for magnetic field detection in animals, Biosystems 13, 181 (1981).

[66] S. E. Greene and A. Komeili, Biogenesis and subcellular organization of the magnetosome organelles of magnetotactic bacteria, Curr. Opin. Cell Biol. 24, 490 (2012).

[67] O. Gorobets, S. Gorobets, and M. Koralewski, Physiological origin of biogenic magnetic nanoparticles in health and disease: From bacteria to humans, Int. J. Nanomedicine 12, 4371 (2017).

[68] N. B. Edelman, T. Fritz, S. Nimpf, and D. A. Keays, No evidence for intracellular magnetite in putative vertebrate magnetoreceptors identified by magnetic screening, Proc. Natl. Acad. Sci. U.S.A. 112, 262 (2015).

[69] G. Fleissner, B. Stahl, and P. Thalau, A novel concept of Fe-mineral-based magnetoreception: Histological and physicochemical data from the upper beak of homing pigeons, Naturwissenschaften 94, 631 (2007).

[70] G. Falkenberg, K. Schuchardt, M. Kuehbacher, P. Thalau, H. Mouritsen, D. Heyers, G. Wellenreuther, and G. Fleissner, Avian magnetoreception: Elaborate iron mineral containing dendrites in the upper beak seem to be a common feature of birds, PLoS ONE 5, e9231 (2010).

[71] M. Walker, C. Diebel, and C. Haugh, Structure and function of the vertebrate magnetic sense, Nature 390, 371– (1997).

[72] E. M. Pantelouris, B. Knox, and H. Wallace, Iron in amphibian oocytes and embryos, Exp. Cell Res. 32, 469 (1963).

[73] T. Nomizu, K. H. Falchuk, B. L. Vallee, and B. L. Vallee, Zinc, iron, and copper contents of *Xenopus laevis* oocytes and embryos, Mol. Reprod. Dev. 36, 419 (1993).

[74] J. L. Kirschvink, J. Brassart, and M. H. Nesson, Magnetite-based biological effects in animals: Biophysical, contamination, and sensory aspects. EPRI, Palo Alto, CA. Report TR–111901 (1998).

[75] U. E. Steiner and T. Ulrich, Magnetic field effects in chemical kinetics and related phenomena, Chem. Rev. 89, 51 (1989).

[76] Z. Zeng, J. Wei, Y. Liu, W. Zhang, and T. Mabe, Magnetoreception of photoactivated cryptochrome 1 in electrochemistry and electron transfer, ACS Omega 3, 4752 (2018).

[77] A. Ehrenberg, P. Hemmerich, F. Müller, and W. Pfleiderer, Electron spin resonance of pteridine radicals and the structure of hydropteridines, Eur. J. Biochem. 16, 584 (1970).

[78] F. Müller, ed., Chemistry and Biochemistry of Flavoenzymes (CRC Press, 1991).

[79] M. J. Leask, A physico-chemical mechanism for magnetic field detection by migratory birds and homing pigeons, Nature 267, 144 (1977).

[80] W. W. Cochran, H. Mouritsen, and M. Wikelski, Migrating songbirds recalibrate their magnetic compass daily from twilight cues, Science 304, 405 (2004).

[81] H. G. Hiscock, T. W. Hiscock, D. R. Kattnig, T. Scrivener, A. M. Lewis, D. E. Manolopoulos, and P. J. Hore, Navigating at night: Fundamental limits on the sensitivity of radical pair magnetoreception under dim light, Q. Rev. Biophys. 52, e9, 1 (2019).

[82] M. Cifra and P. Pospisil, Ultra-weak photon emission from biological samples: Definition, mechanisms, properties, detection and applications, J. Photochem. Photobiol. B 139, 2 (2014).

[83] R. van Wijk and C. G. J. van Wijk, Ultra-Weak Photon Emission: Characteristics, Origin, and Application (Springer, 2024).

[84] N. F. Ramsey, A molecular beam resonance method with separated oscillating fields, Phys. Rev. 78, 695 (1950).

[85] C. D. Aiello, M. Hirose, and P. Cappellaro, Composite-pulse magnetometry with a solid-state quantum sensor, Nat. Commun. 4, 1419 (2013).

[86] E. D. Yorke, A possible magnetic transducer in birds, J. Theor. Biol. 77, 101 (1979).

[87] J. Ashmore, Cochlear outer hair cell motility, Physiol. Rev. 88, 173 (2008).

[88] J. R. Woodward, T. J. Foster, A. R. Jones, A. T. Salaoru, and N. S. Scrutton, Time-resolved studies of radical pairs, Biochem. Soc. Trans. 37, 358 (2009).

